# Dysregulated cysteine metabolism leads to worsened liver pathology in diabetes-tuberculosis comorbid mice

**DOI:** 10.1101/2022.12.21.521387

**Authors:** Shweta Chaudhary, Falak Pahwa, Ranjan K. Nanda

## Abstract

Diabetes mellitus (DM) is a well-known risk factor for tuberculosis (TB). Interestingly, DM is growing to pandemic proportions in TB endemic South-East Asian countries. DM-TB comorbidity induced pathophysiological changes warrants a better understanding to develop effective therapeutics. Tissue metabolomic profiling of streptozotocin (STZ) induced diabetic animals, infected with *Mycobacterium tuberculosis* H37Rv, showed metabolic dysregulation in the lungs, liver, brain, kidney and thigh muscle. At 3 w.p.i., the tissue (lungs, spleen, liver) bacterial loads were similar between DM-TB and TB with worsened lung pathology. Enrichment analysis of the deregulated liver metabolites (n=20; log_2_DM-TB/TB>±1.0) showed major perturbation in the cysteine-methionine, glycine-serine, branched chain amino acid (BCAA) and fatty acid metabolism. Parallel relative quantification of liver proteome of DM-TB and control mice groups (TB, DM and healthy) identified 1833 proteins which showed group specific variations. Enrichment analysis of significantly altered proteins (n=60; log_2_DM-TB/TB>±1.0) showed major perturbations in cysteine-methionine metabolism corroborating the metabolomics data. In addition, amino acid biosynthesis, retinol metabolism and polyol biosynthetic process were also differentially enriched in DM-TB groups compared to controls. Furthermore, a global correlation analysis of liver metabolome and proteome data showed strong association between aspartic acid, pyruvic acid, leucine and isoleucine with Cyp450 enzymes (Cyp2a5, Cyp3a11, Cyp4a10, Cyp4a14) involved in retinol metabolism. Whereas iminodiacetic acid, isoleucine and γ-aminobutyric acid strongly correlated to enzymes (Cth, Ahcy, Kyat3, Mat1a) involved in the cysteine metabolism. So, targeting the perturbed liver cysteine and retinol metabolism in DM-TB comorbid condition might improve therapeutic outcomes and prevent organ damage.

## Introduction

Tuberculosis (TB), caused by *Mycobacterium tuberculosis* (Mtb), is a major global health threat. WHO South-East Asian region contributes to ∼46% of global TB burden and ∼36% of mortality due to TB.^1^ Post COVID, the TB prevalence in these regions have increased significantly.^1^ Globally, diabetes mellitus (DM) is also growing to epidemic proportions with 537 million adults affected in 2021.^2^ DM is a metabolic disorder encompassing multiple organs like brain, heart, kidney and liver, associated with a higher risk of stroke, end-stage renal disease and non-alcoholic fatty liver disease (NAFLD).^3,4^ DM also presents with a higher (3-fold) risk of active TB disease with higher odds of death during treatment (6-fold) as well as relapse (4-fold) after treatment completion.^5,6^

Chronic hyperglycemia in streptozotocin (STZ) induced DM mice, leads to an impaired adaptive immune response and a delayed dissemination of Mtb to draining lymph nodes.^7^ Impaired phagocytosis of alveolar macrophages, reduced IL-22 levels and increased cellular immunity were reported in DM-TB mice.^7,8^ Lung metabolome analysis of the Mtb infected mice showed dysregulation in metabolic and oxido-reductive pathways with high abundance of trimethylamine-N-oxide (TMAO).^9^ However, how DM-TB comorbidity impacts host tissue metabolism like liver, the major contributor of inflammatory mediators at molecular levels, remains unexplored.

In this study, we performed global metabolome and proteome analyses, using mass spectrometry, to identify tissue level changes in Mtb infected DM mice and controls. Exacerbated liver pathology in DM-TB mice with altered retinol and cysteine-methionine metabolism were observed. These metabolic pathways play a crucial role in oxidative stress response and excess of these intermediates induce hepatotoxicity.^10,11,12^ In addition, the Cyp450 family, involved in drug metabolism, showed deregulation in DM-TB which might impact drug metabolism, thereby highlighting the need to formulate specific personalized treatment strategies for patients with comorbid conditions like DM, in focus.

## Results

### Mtb infected diabetic and control C57BL/6 mice showed similar tissue bacterial load

Multiple low doses (50 mg/kg) of STZ resulted in hyperglycemia (random blood glucose: >250 mg/dL, p<0.005) in C57BL/6 female mice with ∼10% loss of body weight (**Figure 1A, 1B, 1C**). These hyperglycemic mice upon low aerosol dose of Mtb H37Rv (100-120 CFU) infection, showed marginal differences in gross tissue pathologies at 3 w.p.i. with similar tissue (lung, liver and spleen) bacterial burden to control (**Figure 1D, 1E**). Histology analysis of the lungs of DM, TB and DM-TB animals showed higher cellular infiltration compared to control (**Figure 1F, Supplementary Figure S1**). In the lungs of DM-TB animals, pulmonary edema was higher (**Figure 1F**). In DM and DM-TB groups, ectopic fat deposition in liver and nephropathy like glomerular and tubule-interstitial degeneration were also observed (**Figure 1F, Supplementary Figure S1**).

**Figure 1.**
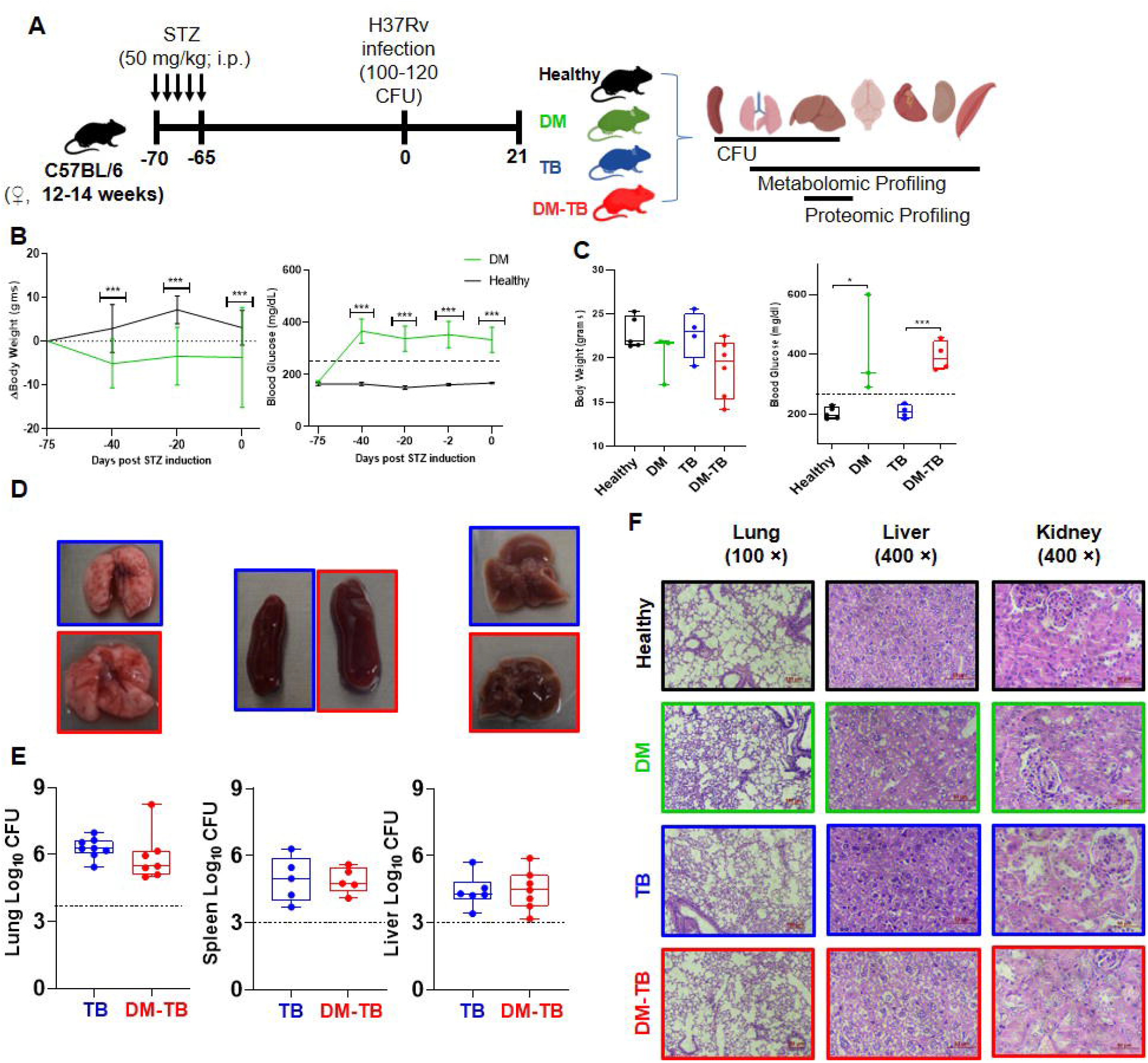
At early time points after Mtb H37Rv infection, DM-TB mice show similar tissue bacterial load but worsened tissue pathology. (A) Schematic showing method used for DM induction and Mtb infection for tissue bacterial load, metabolite and proteome profiling. (B) Change in body weight and blood glucose levels in DM induced mice. (C) Body weight and blood glucose levels of Mtb infected and control study groups (Healthy, DM, TB, DM-TB) at 21 days post infection (d.p.i.); dotted line represents >250 mg/dL. (D) Gross pathology of lungs, spleen and liver of TB and DM-TB groups. (E) Tissue bacterial load (lungs, spleen and liver) of the TB and DM-TB groups at 21 d.p.i.; dotted line represents limit of detection. (F) Hematoxylin and Eosin-stained lung, liver and kidney tissue sections showing histopathology differences between study groups, scale bar: lungs: 50 μm and liver/kidney: 100 μm. STZ: streptozotocin; H37Rv: *Mycobacterium tuberculosis* H37Rv (Mtb); CFU: Colony forming units; DM: streptozotocin induced diabetes C57BL/6 mice; TB/DM-TB: *Mycobacterium tuberculosis* H37Rv infected control/DM mice; *p ≤ 0.05, ***p≤0.005.

### Metabolic changes observed in multiple tissues of Mtb infected STZ induced diabetic mice

Polar metabolites extracted from multiple tissues (lungs, liver, brain, heart, thigh muscle, kidney) of the DM-TB and controls (TB, DM and healthy) were profiled using GC-MS and ∼160 molecular features were identified **(Figure 2A)**. Lung and liver metabolites of DM-TB, TB, DM and healthy showed separate clusters in the partial least square discriminant analysis (PLS-DA) plots **(Figure 2B, 2C, Supplementary Figure S2)**. Tissue metabolites of DM-TB and controls (TB, DM, healthy) showed significant differences (**Supplementary Figure S2**). All the QC samples (pooled from all the test samples) were clustered together, so the observed variations at metabolic components are due to the biological changes and not associated with the adopted processing and data acquisition methods **(Supplementary Figure S3)**. Out of 128 molecules identified in the lungs, 15 showed deregulation (−log_10_ p-value ≥ 0.05; log_2_ Fold change ≥ 1.0) between DM-TB and TB **(Figure 2D, Supplementary Table S1)**. In the liver, 20 deregulated molecules, (out of 160) were observed in DM-TB and TB **(Figure 2E, Supplementary Table S2)**. Identity of all the deregulated molecules were confirmed by running a mixture of commercial standards and by matching their retention time and fragmentation pattern (**Supplementary Figure S4**). Significantly higher liver pyruvic acid levels were observed in both DM-TB and DM groups as compared to healthy controls. Similarly, high levels of lactic acid were observed in the liver of Mtb infected groups (DM-TB and TB) from the controls (**Supplementary Figure S5**). In addition, liver cysteine levels of TB and DM groups were significantly low with respect to healthy control (**Supplementary Figure S5**). Interestingly, liver cysteine levels in DM-TB mice were comparable to healthy control which could be attributed to infection associated increase in the cysteine levels in DM-TB comorbid condition. Metabolite set enrichment analysis (MSEA) of the identified dysregulated liver metabolic features of the DM-TB and TB groups showed altered glutathione-homocysteine, amino acid, fatty acid and citric acid cycle metabolism **(Figure 2F)**.

**Figure 2.**
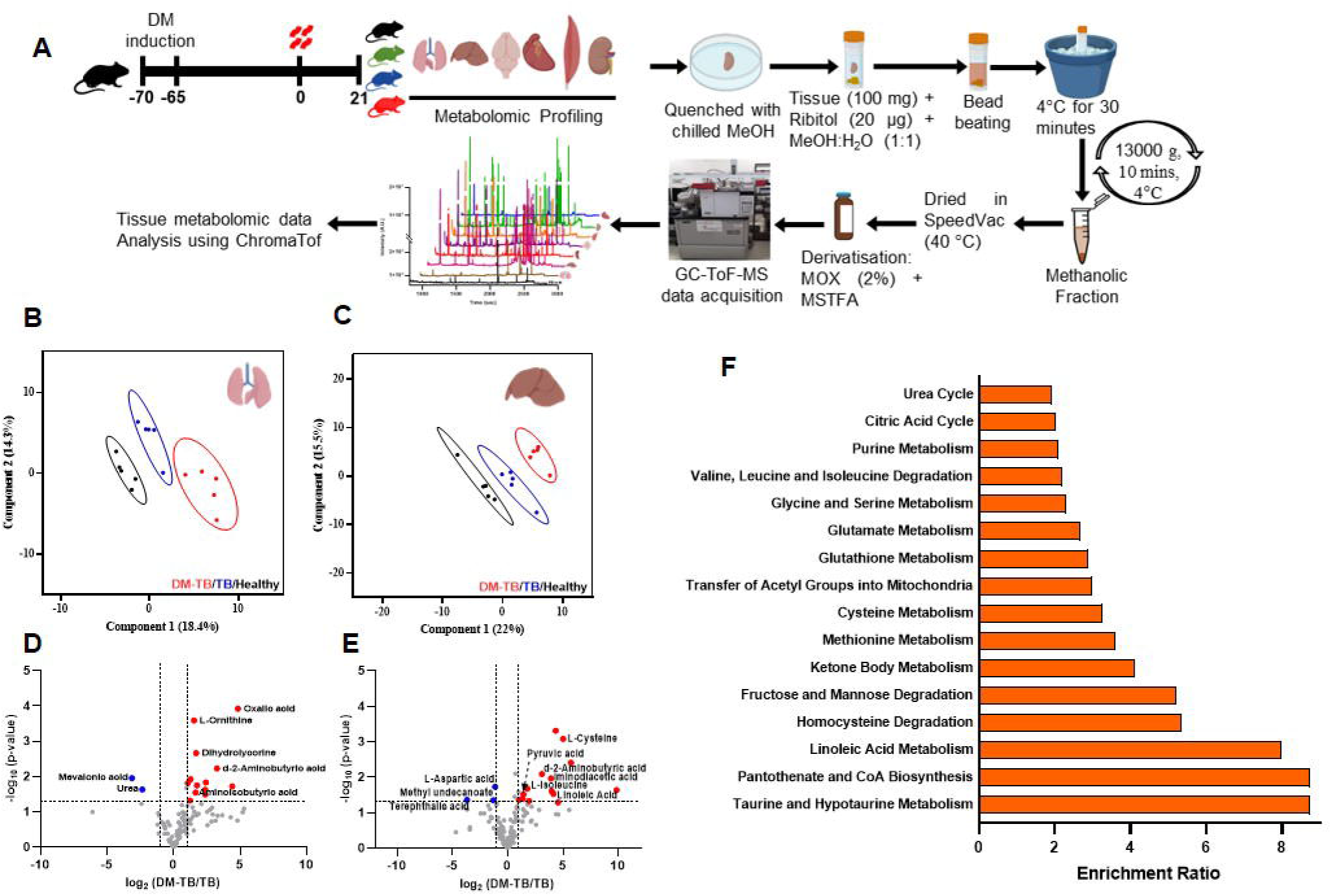
Tissue metabolic profiling of DM-TB and TB mice show marked differences from the healthy controls. (A) Schematic representation of method adopted for metabolite extraction, derivatization, GC-ToF-MS data acquisition and analysis. Partial Least Square-Determinant Analysis (PLS-DA) plots of lungs (B) and liver (C) metabolites of study groups (DM-TB, TB and healthy). Volcano plots of lung (D) and liver (E) metabolome showing significantly deregulated (−log_10_ p-value ≥ 0.05; log_2_ Fold change ≥ 1.0) metabolic features; 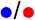: up-/down-regulated. (F) Metabolite set enrichment analysis (MSEA) of deregulated liver metabolites showing differentially enriched metabolic pathways in DM-TB. TB/DM-TB: *Mycobacterium tuberculosis* H37Rv infected control/streptozotocin induced diabetes C57BL/6 mice.

### Liver proteome alters in Mtb infected STZ induced diabetic mice

From the multiplex TMT quantitative proteomics experiment, equal amount of liver proteins were compared and a set of 1833 unique liver proteins were identified in the DM-TB and control groups (TB, DM, healthy) **(Figure 3A, 3B, Supplementary Figure S6, S7, Table S3)**. A set of 60 proteins showed deregulation (log_2_fold change ≥1; FDR adjusted p ≤ 0.05, 31/29: up-/down-regulated) in the DM-TB group with respect to TB **(Figure 3B, 3C)**. Functional analysis of these important liver proteins using Metascape showed perturbations in 20 functional modules including monocarboxylic acid metabolism, amino acids biosynthesis, cysteine and methionine metabolism, retinol metabolism, monocarboxylic acid metabolic process, polyol biosynthetic process, steroid hormone biosynthesis, small molecule catabolic and biosynthetic process **(Figure 3D)**. Cysteine and methionine metabolism modules had one of the highest numbers (n=11) of identified contributors and an inverse expression of these was observed between DM-TB and TB groups. Interestingly, proteins involved in retinoic acid production (Cyp4a10, Cyp4a14, Cyp3a11, Cyp3a16) showed significant up-regulation in DM-TB than TB.

**Figure 3:**
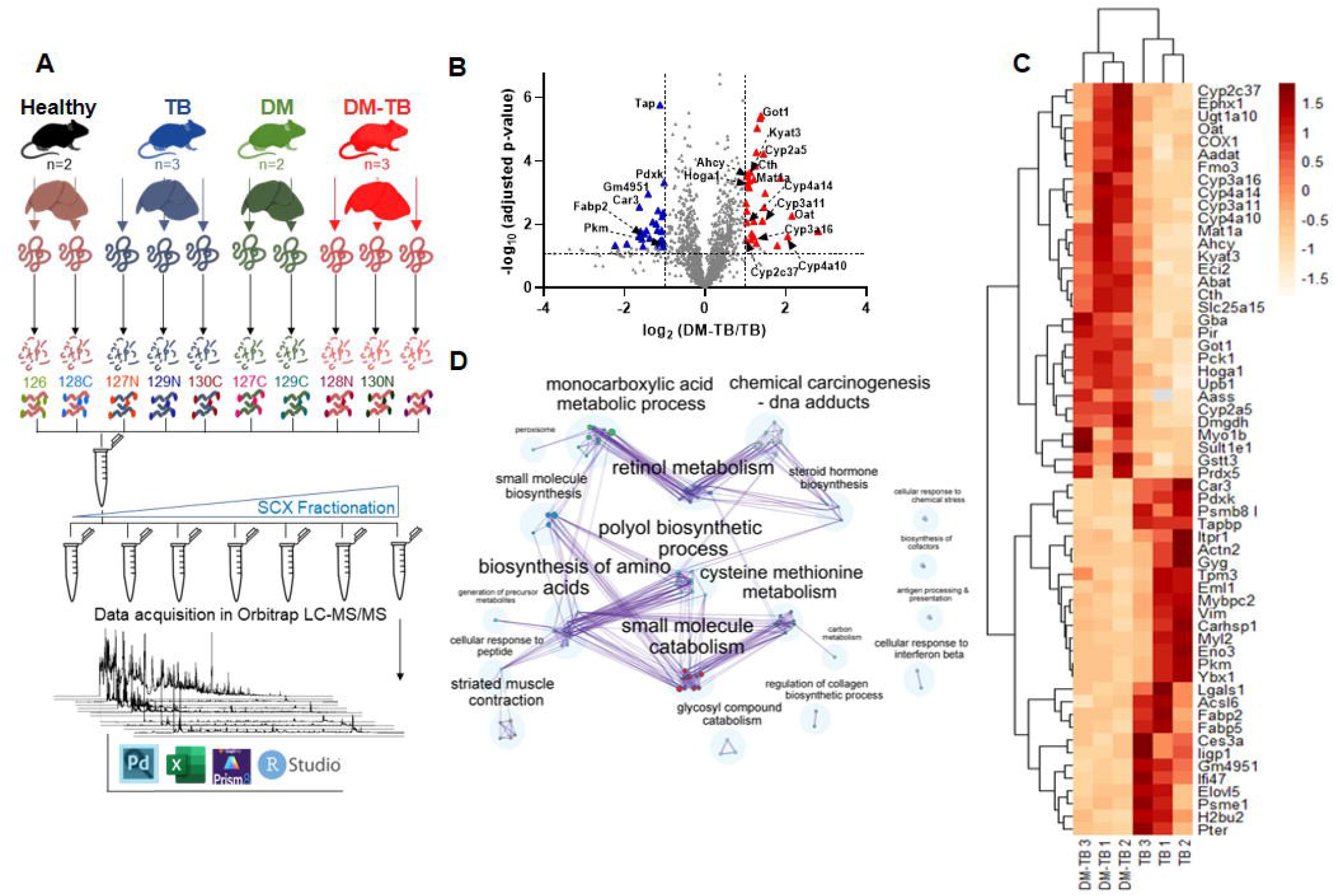
Liver proteome profile of DM-TB and TB groups showed an altered cysteine-methionine and retinol metabolism. (A) Schematic representation of method adapted for TMT based liver proteome analysis of DM-TB and control groups. (B) Volcano plot of liver proteome showing significantly deregulated proteins in (−log_10_ adjusted p-value ≥ 0.05; log_2_ Fold change DM-TB/TB ≥ 1.0) 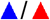: up-/down-regulation. (C) Metascape gene enrichment analysis of gene ontology (GO) terms of significantly deregulated proteins. (D) Heatmap showing abundance of significantly deregulated proteins in DM-TB vs. TB group. DM: streptozotocin induced diabetes C57BL/6 mice; TB/DM-TB: *Mycobacterium tuberculosis* H37Rv infected control/DM mice; SCX: strong cation exchange.

### Correlation analysis of the deregulated liver metabolome and proteome of DM-TB

Pairwise correlation analysis of the important liver proteins (n=60) and metabolites (n=21) clustered the co-varying molecules together and 25 molecules showed high degree of correlation (R^2^>0.9±0.1; p ≤ 0.05) **(Figure 4A, 4B, Supplemental Figure S7)**. These molecules were grouped in three major clusters involved in retinoic acid, aspartic acid, cysteine and methionine metabolism **(Figure 4C)**. Cyp4a and Cyp3a subfamily members (Cyp4a10, Cyp4a14, Cyp3a11, Cyp3a16) showed a strong positive correlation with leucine, isoleucine and 9,12-octadecadienoic acid (Linoleic acid). A set of Cyp450 important proteins (Cyp4a10, Cyp4a14, Cyp2a5 and Cyp3a11) exhibited a positive correlation (R^2^ = 0.7-0.95) with pyruvic acid and a negative correlation (R^2^ = 0.76-0.98) with aspartic acid. Liver Got1 (Aspartate amino transferase), was found to be positively correlated (R^2^>0.8) to liver acetone, isoleucine, valine and cysteine while negatively (R^2^ = 0.95) correlated to γ-aminobutyric acid (GABA). Iminodiacetic acid and leucine levels in the liver of DM-TB groups showed a strong positive correlation (R^2^ = 0.55-0.96) with enzymes (Cth, Ahcy, Kyat3, Mat1a) involved in the cysteine-methionine metabolism. In addition, liver GABA level was negatively correlated (R^2^ = 0.55-0.97) with all these enzymes except Cth.

**Figure 4:**
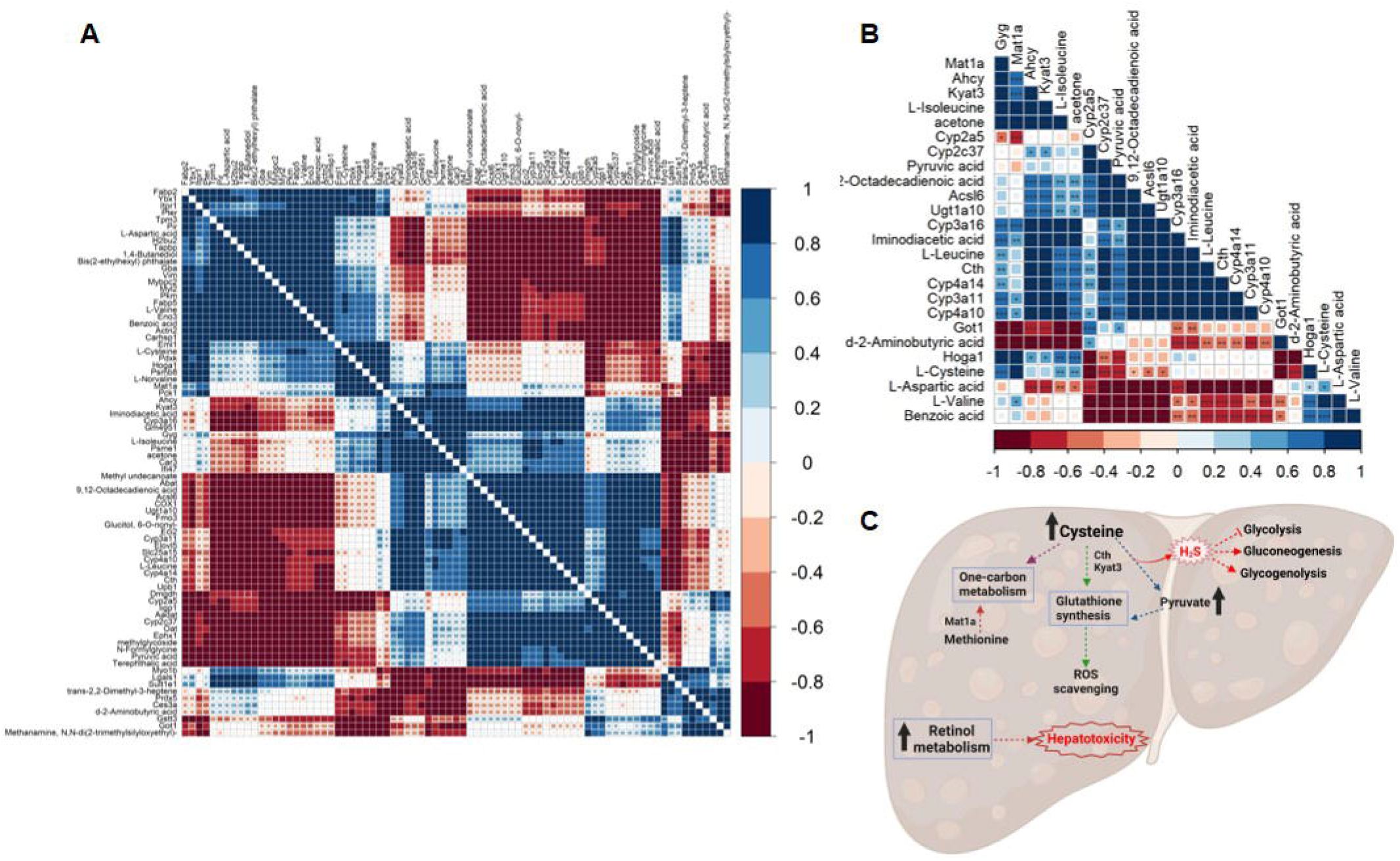
Global correlation analysis of deregulated liver proteins and metabolites demonstrated a strong positive correlation between pyruvic acid and retinol metabolism in DM-TB mice. (A) Pairwise correlation of significantly deregulated (p-value ≤ 0.05; log_2_ Fold change DM-TB/TB ≥ ±1.0) proteins and metabolites in DM-TB group. 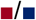: negative/ positive correlation. (B) A section of global correlation map showing correlation of proteins involved in cysteine-methionine and retinol metabolism with important metabolites in DM-TB group. (C) Metabolic consequences of Mtb infected diabetic liver at protein and metabolite levels. DM-TB: *Mycobacterium tuberculosis* H37Rv infected streptozotocin induced C57BL/6 mice.

## Discussion

Higher circulatory blood sugar in diabetes mellitus (DM) influences major host tissue metabolism leading to metabolic disorders.^3,4^ Perturbed tissue metabolic pathways in DM impact the overall host homeostasis making it susceptible to different pathogenic attacks.^13^ In rodent and non-human primate models, upon Mtb introduction, the pathogen disseminates from the lungs to the lymph nodes, spleen and liver.^14,15^ Involvement of liver as a part of disseminated TB is seen in ∼50-80% of pulmonary TB patients, however, hepatobiliary TB is rare.^16^ Majority of the hepatic TB cases are reported in immunocompromised conditions like HIV co-infection.^17^ Diabetic patients often present with liver steatosis (∼30%) which may progress to NAFLD.^4^

STZ induced diabetic mice models are used commonly and, in this study, upon low dose of Mtb infection in DM mice, tissue specific metabolic and protein changes were monitored. At 3 w.p.i., DM-TB animals showed similar lungs, spleen and liver bacterial burden with the Mtb infected animals corroborating earlier reports.^7^ As expected, lungs of DM-TB mice showed higher neutrophil infiltration and inflammation compared to TB controls. Ectopic fat deposition observed in the liver of DM-TB and DM mice indicates development of insulin resistance which was absent in TB mice. Nephropathy, a common complication reported in DM, was also observed in the DM-TB mice with higher glomerular and tubulointerstitial degeneration.

Continuous hyperglycemia in uncontrolled diabetes influences major cellular biochemical pathways leading to organ dysfunction.^3,4,18^ Multiple tissues (lungs, liver, brain, kidney, heart and thigh muscle) of DM-TB, TB and healthy mice groups showed significant variation in their metabolome components. MSEA analysis of liver metabolites showed differential enrichment of pathways involving cysteine such as the taurine, hypotaurine and homocysteine metabolism. In the current study, liver proteome of DM-TB and TB mice showed a set of 60 deregulated molecules involving in small molecule catabolic process, retinol metabolism, polyol biosynthetic process, cysteine and methionine metabolism. Similar observations were reported in the liver proteome of diabetic Chinese hamsters which showed enriched proteins related to metabolism of glucose, lipid, linoleic acid, amino acid, retinol, purine and bile secretion.^19^

Cysteine and methionine are responsible for glutathione production, thereby contributing to oxidative stress control.^10^ Methionine is a substrate for S-adenosylmethionine (SAM), a universal methyl donor which in turn is used for >85% of all transmethylation reactions.^20^ Cystathionine-γ-lyase (Cth) catalyzes break down of cysteine to pyruvate and H_2_S and accumulation of these byproducts contributes to hepatic dysfunction.^21^ Mice with deficient H_2_S production capacity showed higher resistance to Mtb infection.^22^ We observed higher liver cysteine in the DM-TB and healthy groups, with respect to TB and DM. Higher H_2_S level influences cellular energetics by lowering glycolysis, increasing gluconeogenesis and glycogenolysis.^11^ So, higher liver cysteine levels and Cth abundance in the DM-TB mice might be contributing to the observed exacerbated liver pathology in this group. Liver is the primary site of methionine metabolism and higher liver homocysteine level is associated with NAFLD and liver disorders.^20^ Adenosylhomocysteinase (Ahcy) converts S-adenosylhomocysteine to homocysteine and adenosine. Significantly higher liver Ahcy abundance in DM-TB mice and an enriched homocysteine metabolism observed from the metabolomics data corroborate earlier findings.^23^

Higher serum Got1 levels have been reported in dementia, stroke, colorectal adenoma, frailty, disability, sarcopenia, metabolic syndrome, and liver injury.^24^ Higher liver alanine aminotransferase (ALT) levels in diabetic ob/ob mice was reported but limited literature on Got1 is available.^24,25,26,27,28^ Elevated serum aspartate aminotransferase (AST/Got1) level is used as a marker for NAFLD and liver injury.^29,30^ Higher abundance of liver Got1, which catalyzes conversion of aspartate to a gluconeogenic substrate i.e. oxaloacetate, in the DM-TB mice might have contributed lower liver aspartic acid. The reduced aspartic acid levels might be indicative of increased gluconeogenesis in liver-a hallmark of diabetes, which contributes to increased circulating glucose levels and multiple organ dysfunction.^31^

Retinol acts as an antioxidant and its’ metabolism generates retinoic acid which regulates expression of multiple genes involved in insulin release and energy homeostasis.^32^ Retinoic acid also regulates the checkpoints in inflammation and immune tolerance, thereby playing a critical role in host defense against pathogens.^33^ Multiple enzymes (Cyp2a5, Cyp2c37, Cyp3a11, Cyp3a16, Cyp4a10, Cyp4a14) of the retinol metabolism involved in the conversion of trans retinoate to all trans-hydroxyretinoic acid, were highly abundant in the liver of DM-TB group compared to TB. Similar high liver Cyp2a5 levels were also reported in *Plasmodium berghei* and *Citrobacter rodentium* infections, liver fluke infestation and in alcohol induced liver injuries.^34,35^ Liver Cyp3a subfamily metabolizes multiple drugs and TLR-4 dependent inflammation lowers Cyp3a11 abundance.^36^ In the DM-TB mice, we observed higher liver Cyp3a11 levels whereas lower in the TB group with respect to healthy. Altered Cyp3a11 expression in DM-TB mice might influence metabolism of the TB drugs which warrants further evaluation. Cyp4a protein family in the liver is involved in ω-fatty acid hydroxylation and the deregulated Cyp4a10 and Cyp4a14 metabolizes medium and long chain fatty acids, respectively. Murine Cyp4a10 and Cyp4a14 convert arachidonic acid to 20-hydroxyeicosatetraenoic acid, which regulates inflammatory processes through the generation of ROS.^37^ Higher Cyp4a10 levels in the liver of DM-TB mice might be indicative of steatosis development.

Disrupted pyruvic acid metabolism has been reported in cancer, heart failure, neurodegeneration and DM.^38^ We observed high liver pyruvic acid in DM and DM-TB groups indicating that the increase is primarily due to DM, not TB. Increased PDK activity inhibits PDH and decreases pyruvate flux into TCA cycle.^38^ In the current study, low liver PDH abundance was observed in the DM-TB mice and PDK was not detected. Pyruvate replenishes glutathione to reduce ROS levels and higher abundance of it in DM-TB mice might be useful to handle the accumulated ROS.^38^ Low serum aspartic acid levels in DM patients was reported and its’ supplementation benefits in diabetic kidney disease.^39^ In this study, strong correlations between aspartic acid and pyruvic acid with Cyp450 enzymes was observed and needs further investigation.

In conclusion, the current study showed that Mtb infection in DM mice significantly affects liver patho-physiology. Deregulated liver retinol metabolism observed in DM-TB condition might be accelerating ROS production leading to the progression of liver disease. Increased cysteine metabolism in DM-TB may be contributing higher H_2_S production worsening glycemic control. Thus, screening of the DM-TB comorbid conditions at the time of case presentation might be useful to introduce appropriate interventions to minimize liver injury.

## Materials and Methods

### Animal model for diabetes induction and Mtb infection for harvesting tissues

The animal experimental protocols followed in this study are approved by the institute animal ethics committee (vide reference ICGEB/IAEC/07032020/TH-13Ext). **Female** C57BL/6 mice (12-14 weeks) were subjected to multiple low STZ dose (MLD, 5 doses, 50 mg/kg) for diabetes induction following earlier reported method.^40^ Mice with random plasma glucose >250 mg/dL (measured using a glucometer of Dr. Morepen Gluco-One BG-03) were grouped as diabetic (DM) and parameters like body weight and random blood glucose were monitored weekly till experiment completion. After 60 days of hyperglycemia, animals were infected with low dose (100-120 CFU/animal) Mtb H37Rv using a madison chamber in tuberculosis aerosol challenge BSL-III facility at the host institution. After 1- and 21-days post infection (d.p.i.), animals were anaesthetized for whole blood collection via retro-orbital puncture and serum samples were isolated. At 21 d.p.i., organs (lung, liver, spleen, kidney, heart, thigh muscle and brain) were harvested for bacterial CFU assay, histology, metabolite and protein extraction from specific tissues. Part of the lung, liver and kidney stored in buffered formalin solution were used for histology analysis using standard hematoxylin and eosin staining methods. For CFU enumeration, spleen, lung (left lobe) and liver (2/3rd part) were homogenized in phosphate buffer saline (PBS) and plated on 7H11 media supplemented with OADC (10 %) in triplicates and CFU counting was carried out 21 days later. For metabolite analysis, tissues (heart, brain, thigh muscle, part of kidney, lung and liver) were treated with chilled methanol (−80 °C) immediately after harvesting and stored at -80 °C. Part of these tissues were snap-freezed immediately and stored at -80 °C until further protein analysis.

### Tissue metabolite extraction, trimethylsilyl derivatization, GC-Tof-MS mass spectrometry data acquisition and analysis

Tissues (∼100 mg) were chopped finely (10 × 10 mm) and transferred to bead beating tubes (2 ml) with zirconium beads (2 mm, 250 mg). Chilled methanol (80%, 1 ml) was added to the tubes with Ribitol (5 μl, 2 mg/ml) as a spike in standard. Bead beating was done using Biospec Mini-Beadbeater-16 at 3650 oscillations/minute, 30 seconds on/off cycles for six times. Samples were kept on ice for 30 minutes before centrifuging at 10,000 *g* for 10 minutes at 4 °C to collect the supernatant (∼800 μl). The extracted metabolites were filtered (0.2 μm, cellulose acetate filter) and stored at -20 °C until further processing. Equal volume (∼122 μl) of extracted metabolites from all of the test samples (n=148) were pooled to prepare a quality control (QC) sample. For trimethylsilyl derivatization, all samples were randomized using randomizer (www.randomizer.org) and processed in a batch of 20 samples per day with 3 QC samples to run at the beginning, middle and end of the sequence. Briefly, samples (400 μl) were dried at 40 °C using vacuum concentrator (Labconco Centrivap, USA) and incubated with Methoxamine hydrochloride (MeOX-HCl, 2%, 40 μl) at 60 °C for 2 hours at 400 rpm in a thermomixer (Eppendorf, USA). After adding N-methyl-N-(trimethylsilyl) trifluoroacetamide (MSTFA, 70 μl), it was incubated at 60 °C for 30 minutes at 400 rpm in a thermomixer. After centrifuging at 16,000 g for 10 minutes at 23 °C, the supernatant was transferred to GC vial inserts for GC-MS analysis using a 7890 Gas Chromatograph (Agilent Technologies, Santa Clara, CA) coupled to a Pegasus 4D GC × GC time-of-flight mass spectrometer (LECO Corporation). The derivatized samples (1 μL) were injected to an HP-5ms column (30 m length, 250μm internal diameter) in splitless mode using Helium as a carrier gas at a constant flow rate (1 ml/min). The secondary column was Rxi-17 of 1.5 m length and 250μmdiameter. Electron ionization (EI) mode was fixed at −70 eV to scan ions of 33 to 600 m/z range at acquisition rate of 20 spectra/second. Ion source temperature was set at 220 °C. GC method parameters used for acquisition were: 50 °C for 1 min, ramp of 8.5 °C/min to 200 °C, ramp of 6 °C/ min to 280 °C, hold for 5 minutes. Secondary oven temperature offset was set at 5 °C relative to GC oven temperature. Similarly, the modulator temperature offset was set at 15 °C relative to the secondary oven temperature. Transfer line temperature was set at 225 °C. A solvent delay of 600 seconds was used and all data acquisition were completed within 24 hours of derivatization. A commercial standard mix of organic acids and amino acids were derivatized and run using the same GC-MS method to confirm the identity of the important metabolic features.

All GC-MS raw data files of the study groups (Healthy, DM, TB and DM-TB) were aligned using the “Statistical Compare” feature of ChromaTOF (4.50.8.0, Leco, CA, USA). Peaks with a width of 1.3 seconds and the signal to noise ratio (S/N) threshold of 50 were selected. Putative identity of the peaks was assigned using a NIST (National Institute of Standards and Technology, USA) library (version 11.0; 243,893 spectra) search qualifying a similarity match of >600 at a mass threshold of 100. The .csv file, generated from the above analysis, was manually curated to check the peak identity. Analytes with similarity match <600, silanes and S/N <75 were excluded and peak areas of the same analytes, eluting at different retention time, were merged to prepare the final data matrix. Analytes present in > 50% of the samples of a study group were selected for statistical analysis. Ribitol peak area from the solvent blank GC-MS run was used to normalize tissue metabolite data. Metabolites qualifying the parameters-log_2_ Fold change ≥ ±1.0 and -log_10_ p-value ≥ 0.05 between study groups were selected as deregulated molecules.

### Liver proteome analysis in DM-TB and TB

Liver tissues (∼100 mg) were chopped finely (3 × 3 mm) and after adding extraction buffer (500 μL of 8M urea, 100 mM Tris, pH 8 and protease inhibitor cocktail; ThermoFisher Scientific; Cat# A32965) were sonicated at 30% amplitude with 3 cycles of 10 second pulses. After incubating the tissue lysates in ice for 20 minutes, were centrifuged at 16,000 *g* for 30 minutes at 4 °C and the supernatant was collected for protein estimation using Pierce™ BCA Protein Assay Kit (ThermoFisher Scientific; Cat# 23227). Equal amounts of liver protein (100 μg) from the four study groups (Healthy, DM, TB and DM-TB) with at least two biological replicates were taken for multiplex TMT™ isobaric tagging (**Figure 1A**). Extracted protein (5 μg) of all liver samples used in proteomics experiments were separated using SDS-PAGE and silver stained to monitor their distribution (**Supplemental Figure S6**). The protein samples taken for TMT experiment were adjusted to a final volume of 100 μl with TEAB (100 mM, ThermoFisher Scientific; cat# 90110) and reduced by incubating with TCEP (5 μl, 200 mM) at 55 °C for 1 hour followed by iodoacetamide (5 μl, 375 mM) for 30 minutes at room temperature. These samples were treated with methanol-chloroform, centrifuged to collect and the pellet was stored at -80 °C. These protein pellets were resuspended in TEAB (100 μl, 100 mM) and after adding trypsin (2.5 μg/100 μg protein) were incubated for 16 hours at 37 °C. After resuspending the supplied TMT tags (0.8 mg) in acetonitrile (41 μl), the sample tryptic peptides were added to the respective TMT reagent vials and incubated at 22 °C for 1 hour. Labelling of peptides was quenched using hydroxylamine (5%, 8 μl) and incubated at 22 °C for 15 minutes. TMT labelled tryptic peptides from all samples were pooled and dried using a SpeedVac at 40°C for 1.5 hours. An aliquot of labelled tryptic peptides (∼300 μg) was taken for strong cation exchange chromatography (SCX). Briefly, labelled peptides were dissolved in ammonium formate buffer (5 mM, 2 ml) and loaded onto a pre-equilibrated SCX column ICAT™ cartridge kit (AB Sciex, USA). Peptides were eluted using different concentrations (30, 50, 80, 120, 250, 300, 400, and 500 mM) of ammonium formate buffer and collected as fractions (n=8). Initial two fractions eluted with ammonium formate (30 mM and 50 mM) were pooled and all the fractions were dried in the SpeedVac at 40°C. These SCX fractions (n=7) were cleaned up using C18 spin columns (89870, Pierce, ThermoFisher, USA), dried using SpeedVac at 40 °C and resuspended in formic acid (0.1 %, 20 μl) before proceeding with LC-MS/MS analysis.

### LC-MS/MS data acquisition and analyses

Each tryptic peptide fractions (∼2 μg) were resuspended in formic acid (0.1 %, 20 μl) and analysed using an Orbitrap Fusion Lumos Tribrid Mass Spectrometer connected to a nano-LC Easy nLC 1200 (ThermoFisher Scientific, Singapore). The peptides were separated at a flow rate of 300 nL/minute on C18 pre-column (Acclaim PepMap™ 100, 75 μm□×□2 cm, nanoViper, P/N 164946, ThermoFisher Scientific Incorporation) followed by analytical column (Acclaim PepMap™ RSLC C18, 75 μm□×□50 cm, 2 μm, 100 Å, P/N ES803) using a gradient of solvent B (95% acetonitrile in 0.1% formic acid) from 5% - 10% for 0-5 minutes followed by 35% for 5-95 minutes and finally, 95% for 95-130 minutes and solvent A (0.1% formic acid). The eluted peptides were injected into the mass spectrometer and the MS1 data were acquired in positive ion mode at 120,000 orbitrap resolution with mass range from 400 to 1600 Da. Precursor ions were allowed to fragment using higher-energy C-trap dissociation (HCD) in ion trap (IT) detector with collision energy of 38% in a data dependent MSn Scan acquisition. Precursor ions with +2 to +6 charge and monoisotopic ions were selected. Parent ions once fragmented were excluded for 60 s with exclusion mass width of ±□10 ppm.

All these 7 raw files were searched against the *Mus musculus* uniprot proteome database (ID: UP000000589; accessed on 29/09/2022; 51,315 protein count) and contaminant protein database (PD_Contaminants_2015_5.fasta, provided by the manufacturer) in Proteome Discoverer software version 2.3 (Thermo Scientific). All proteomics raw data files are accessible from the ProteomeXchange database with accession number PXD038960. A maximum of two trypsin cleavages were allowed with a precursor mass tolerance of 10 ppm and fragment mass tolerance of 0.8 Da. TMT10plex +229.163 Da (Lys) at N-termini of peptides and carbamidomethylation +57.021 Da (Cys) at C termini were selected as static modifications. Oxidation +15.995 Da (Met) and acetylation +42.011 Da at N-terminus were selected as dynamic modifications and proteins were identified based on at least two peptides at an FDR < 0.05. Proteins qualifying the parameters-log_2_ Fold change ≥ 1.0; -log_10_ adjusted p-value ≥ 0.05 between study groups were selected as important molecules.

### Statistical Analysis

Changes in protein abundance (log_2_fold change) were calculated by taking the ratio of the average abundance between two study groups. Student t-test was carried out using absolute abundance values of samples between groups. Proteins qualifying criteria of log_2_fold change ≥ 1.0 and a p-value ≤ 0.05 were selected as significantly deregulated proteins. Enrichment analysis for protein samples was performed using Metascape. MetaboAnalyst 5.0 was used for multivariate and metabolite set enrichment analysis of metabolomics data. Deregulated metabolites and proteins were taken for correlation analysis using Rstudio (2022.07.2). R functions pheatmap and corrplot were used to prepare protein heatmap and global correlation map respectively.

## Acknowledgments

S.C. is supported by Council of Scientific and Industrial Research-Shyama Prasad Mukherjee Fellowship (CSIR-SPM) fellowship, F.P. is supported by Department of Biotechnology-Senior Research Fellowship (DBT-SRF). Financial supports from the Department of Biotechnology, India and core budget of International Centre for Genetic Engineering and Biotechnology, New Delhi to R.K.N. are highly acknowledged. Department of Biotechnology supported Tuberculosis Aerosol Challenge Facility (TACF) at International Centre for Genetic Engineering and Biotechnology is kindly acknowledged. We thank ICGEB animal house staffs, TACF staffs, Dr(s) Girish H Rajacharya, Hossain Md. Faruquee and Ms. Haripriya Priyadarsini for their help during the experiments.

## Author Contributions

Author contributions: S.C., and R.K.N. designed research; S.C., and F.P. performed research; S.C. analyzed data; and S.C. and R.K.N. wrote the paper.

## Legends of supplementary figures and tables

**Supplementary Figure S1:** Hematoxylin and Eosin-stained lung, liver and kidney tissue sections showing histopathology differences between study groups (Healthy, DM: Streptozotocin induced diabetic C57BL/6 mice, TB: Mycobacterium tuberculosis H37Rv infected control mice, DM-TB: Mycobacterium tuberculosis H37Rv infected DM mice). Scale bar: lungs/liver/kidney: 100 μm.

**Supplementary Figure S2:** Partial Least Square-Determinant Analysis (PLS-DA) plots generated from the identified global metabolites of heart (A), liver (B), lungs (C), kidney (D), thigh muscle (E) and brain (F) of animals belonging to all study groups (H: Healthy, DM: Streptozotocin induced diabetic C57BL/6 mice, TB: Mycobacterium tuberculosis H37Rv infected control mice, DM-TB: Mycobacterium tuberculosis H37Rv infected DM mice). Volcano plots showing the significantly deregulated (−log10 p-value ≥ 0.05; log2 Fold change ≥ ±1.0) metabolic features in each tissue of study groups. 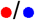: up-/down-regulated.

**Supplementary Figure S3**: Partial Least Square-Determinant Analysis (PLS-DA) plot generated from the global metabolite analysis of representative tissue i.e. lungs of all study groups (Healthy, DM: Streptozotocin induced diabetic C57BL/6 mice, TB: *Mycobacterium tuberculosis* H37Rv infected control mice, DM-TB: *Mycobacterium tuberculosis* H37Rv infected DM mice) including quality control (QC) i.e. mixture of all studied tissue samples from lungs, liver, heart, kidney, thigh muscle and brain.

**Supplemental Figure S4**: Total ion chromatogram (TIC) obtained by running the trimethylsilyl derivatized commercial standards including organic and amino acids used for molecular feature identification. The fragmentation pattern of the test molecules obtained from the run were matched with the available spectral library.

**Supplemental Figure S5**: Relative abundance of individual analytes involved in amino acid and acetone metabolism in the liver of Healthy controls and *Mycobacterium tuberculosis* H37Rv infected control C5BL/6 animals (TB) and streptozotocin induced diabetic animals (DM-TB). A.U.: arbitrary unit.

**Supplemental Figure S6**: Silver-stained gel image of equal amount (4 μg) of liver proteins isolated from the healthy (H), Streptozotocin induced diabetic (DM), *Mycobacterium tuberculosis* H37Rv infected C57BL/6 mice (TB) and DM mice infected with *Mycobacterium tuberculosis* H37Rv (DM-TB). 1,2,3 shows the biological replicates.

**Supplemental Figure S7:** Heatmap showing abundance of identified liver proteins (n=1833), obtained from the TMT multiplex experiment, using the healthy (H), Streptozotocin induced diabetic (DM), *Mycobacterium tuberculosis* H37Rv infected C57BL/6 mice (TB) and DM mice infected with *Mycobacterium tuberculosis* H37Rv (DM-TB). Each row presents abundance of individual protein. 1,2,3 shows the biological replicates. Colour intensity indicates the log_2_fold change values.

**Supplemental Figure S8:** Global correlation analysis of identified liver proteins and metabolites in *Mycobacterium tuberculosis* H37Rv infected C57BL/6 mice (TB). (A) Pairwise correlation of significantly deregulated (p-value ≤ 0.05; log_2_ Fold change DM-TB/TB ≥ 1.0) proteins and metabolites in TB group. 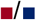: negative/ positive correlation. (B) A section of global correlation map showing correlation of proteins involved in cysteine-methionine and retinol metabolism proteins and their correlation with important metabolites in TB group.

