## Supplemental Figure S1, S2, S3, S4, S5, S6, S7, S8, Table S1, S2, S3 for "Dysregulated cysteine metabolism leads to worsened liver pathology in diabetes-tuberculosis comorbid mice"

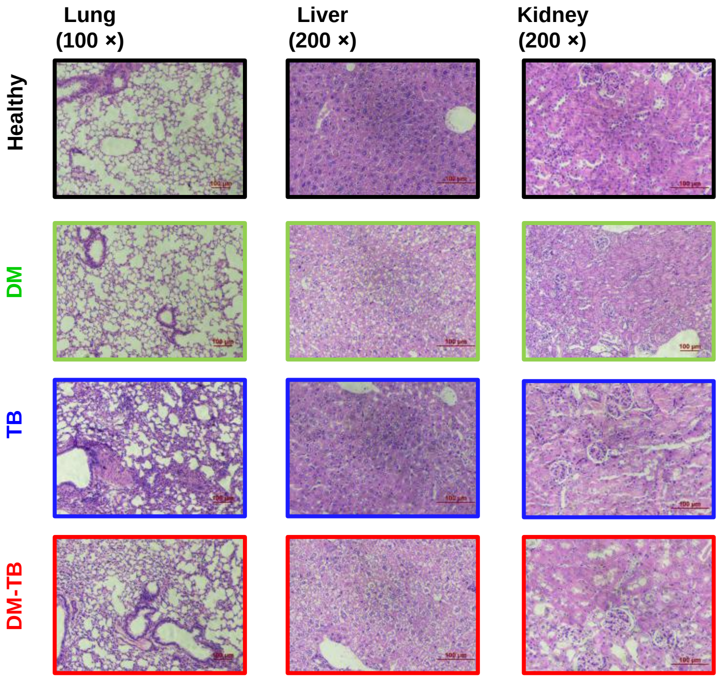


**Supplemental Figure S1**: Hematoxylin and Eosin-stained lung, liver and kidney tissue sections showing histopathology differences between study groups (Healthy, DM: Streptozotocin induced diabetic C57BL/6 mice, TB: *Mycobacterium tuberculosis* H37Rv infected control mice, DM-TB: *Mycobacterium tuberculosis* H37Rv infected DM mice). Scale bar: lungs/liver/kidney: 100 µm.

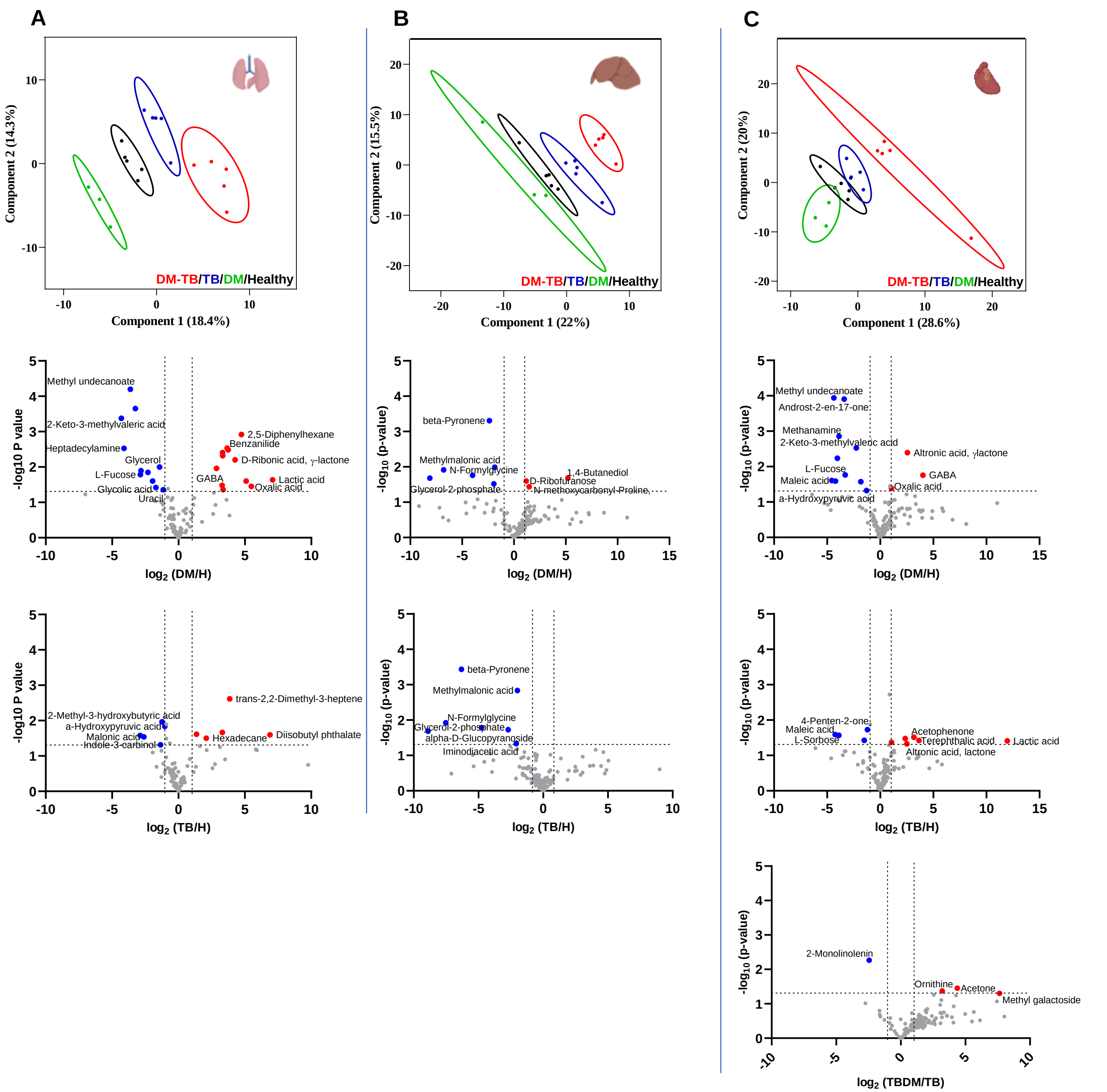


**Supplemental Figure S2**: Partial Least Square-Determinant Analysis (PLS-DA) plots generated from the identified global metabolites of heart (A), liver (B), lungs (C), kidney (D), thigh muscle (E) and brain (F) of animals belonging to all study groups (H: Healthy, DM: Streptozotocin induced diabetic C57BL/6 mice, TB: *Mycobacterium tuberculosis* H37Rv infected control mice, DM-TB: *Mycobacterium tuberculosis* H37Rv infected DM mice). Volcano plots showing the significantly deregulated ($-\log_{10}$ p-value ≥ 0.05; $\log_2$ Fold change ≥ ±1.0) metabolic features in each tissue of study groups. ●/●: up-/down-regulated.

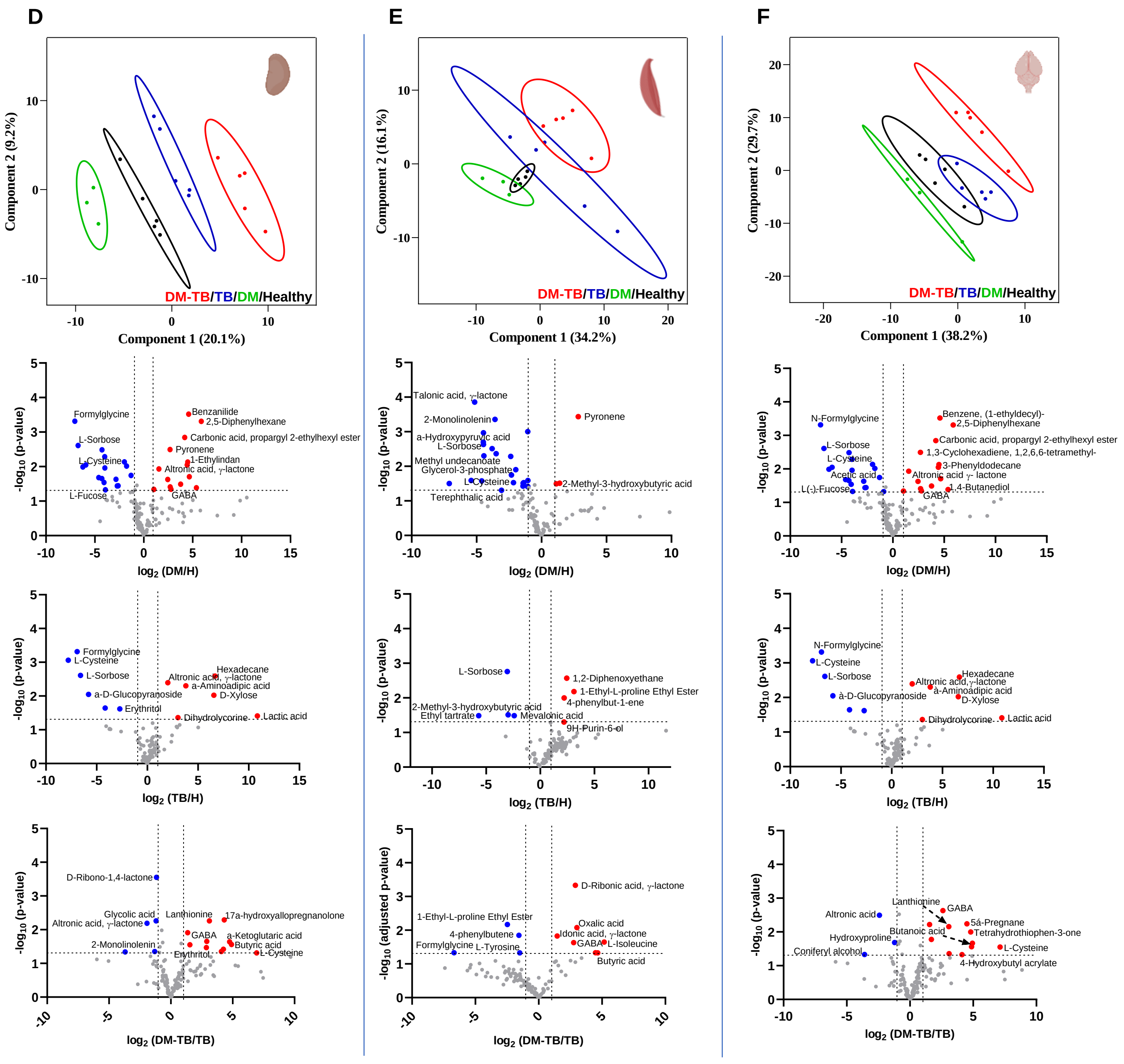


Supplemental Figure S2 (continued)

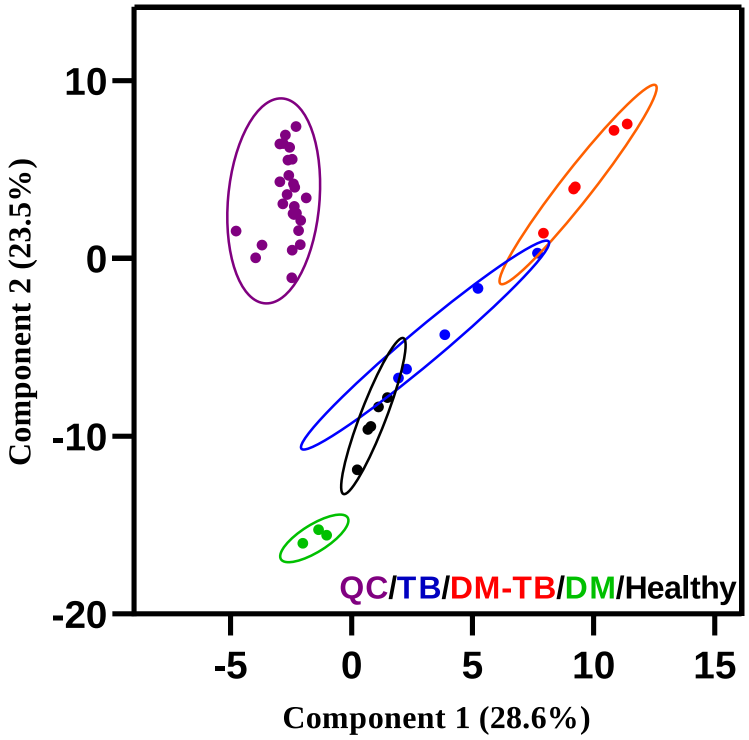


**Supplemental Figure S3**: Partial Least Square-Determinant Analysis (PLS-DA) plot generated from the global metabolite analysis of representative tissue i.e. lungs of all study groups (Healthy, DM: Streptozotocin induced diabetic C57BL/6 mice, TB: *Mycobacterium tuberculosis* H37Rv infected control mice, DM-TB: *Mycobacterium tuberculosis* H37Rv infected DM mice) including quality control (QC) i.e. mixture of all studied tissue samples from lungs, liver, heart, kidney, thigh muscle and brain.

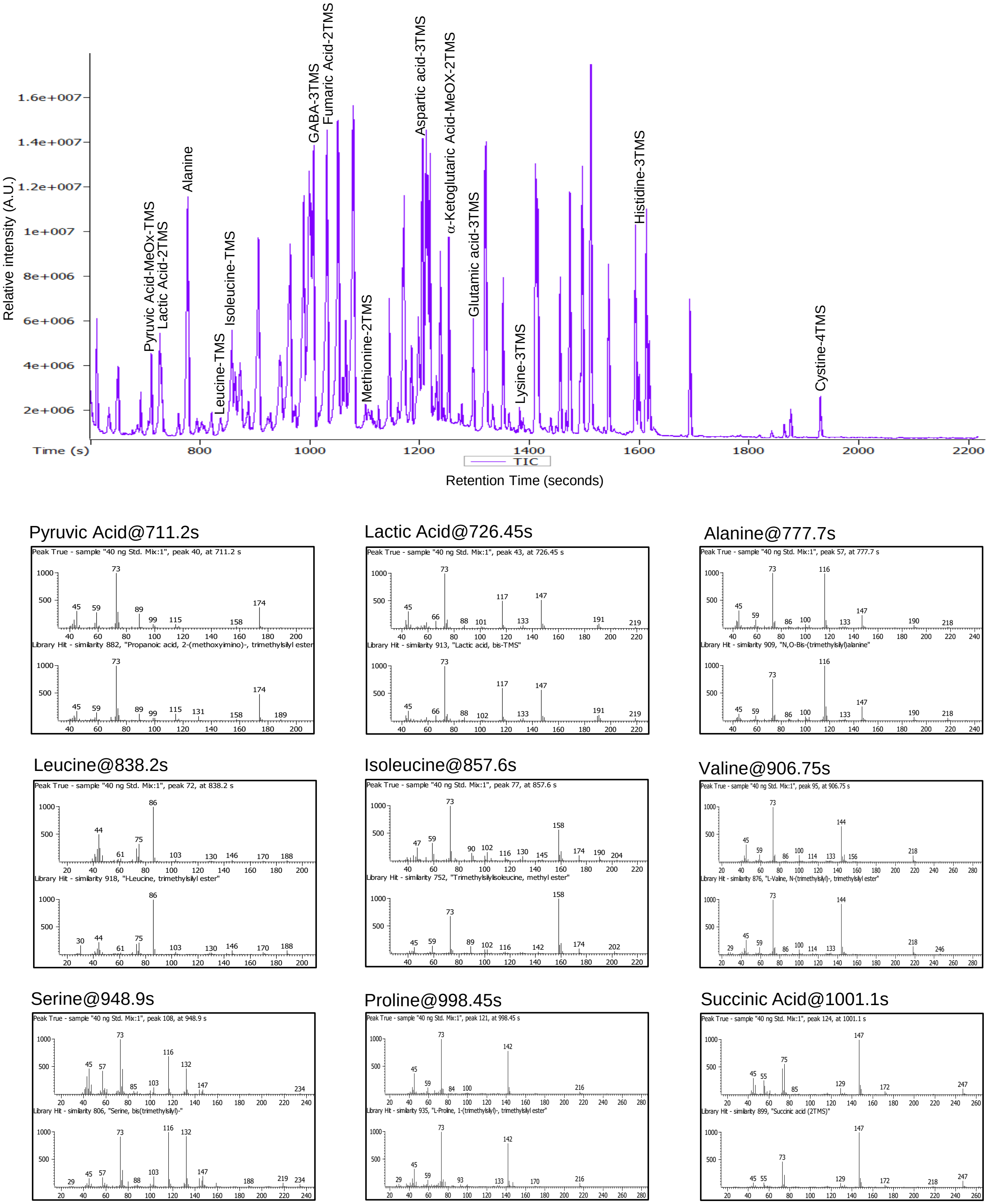


**Supplemental Figure S4**: Total ion chromatogram (TIC) obtained by running the trimethylsilyl derivatized commercial standards including organic and amino acids used for molecular feature identification. The fragmentation pattern of the test molecules obtained from the run were matched with the available spectral library.

Aminobutyric Acid@1007.1s

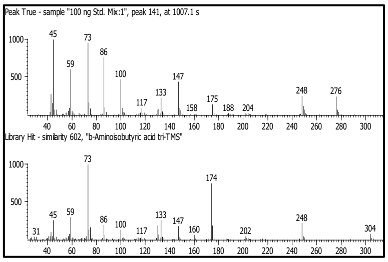


Fumaric Acid@1031.1s

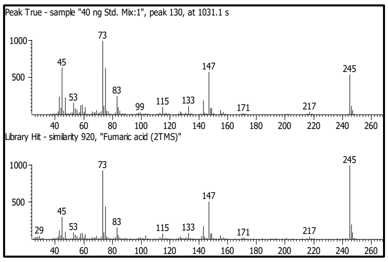


Threonine@1079.6s

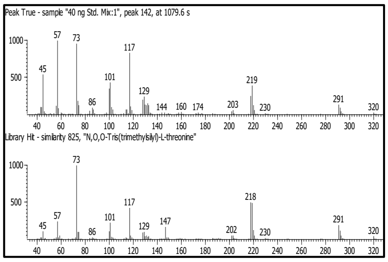


Methionine@1117.85s

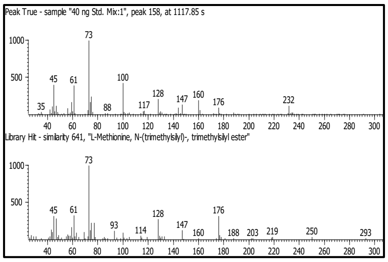


Malic Acid@1171.75s

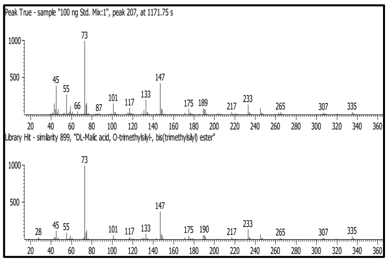


Aspartic acid@1204.1s

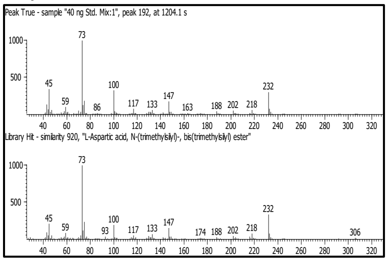


Cysteine@1243.6s

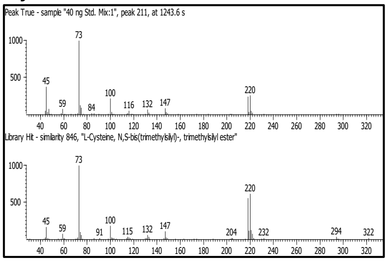


α-Ketoglutaric Acid@1252.7s

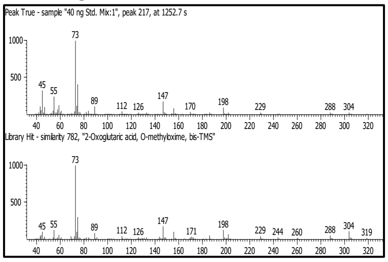


Ornithine@1293.9s

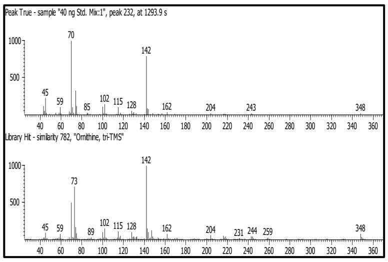


Glutamic acid@1297s

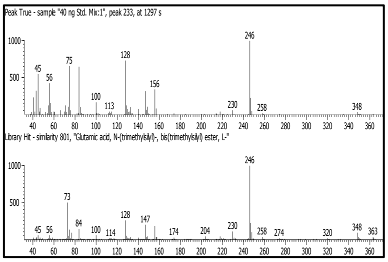


Phenylalanine@1320.25s

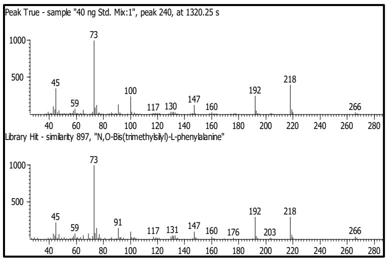


Lysine@1382.05s

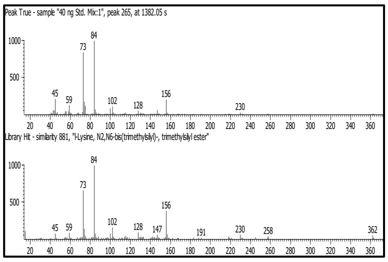


Citric Acid@1496s

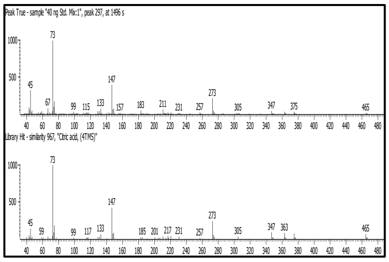


Histidine@1598.95s

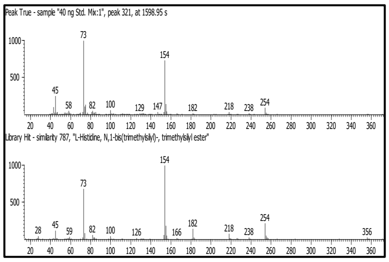


Tyrosine@1612.3s

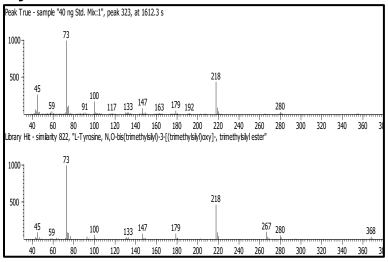


Tryptophan@1875.95s

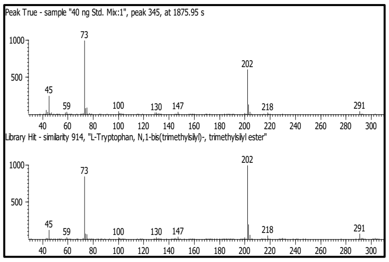


Cystine@1929.95s

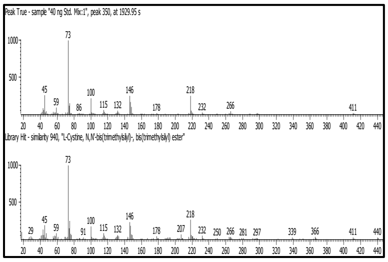




Supplemental Figure S4 (continued)

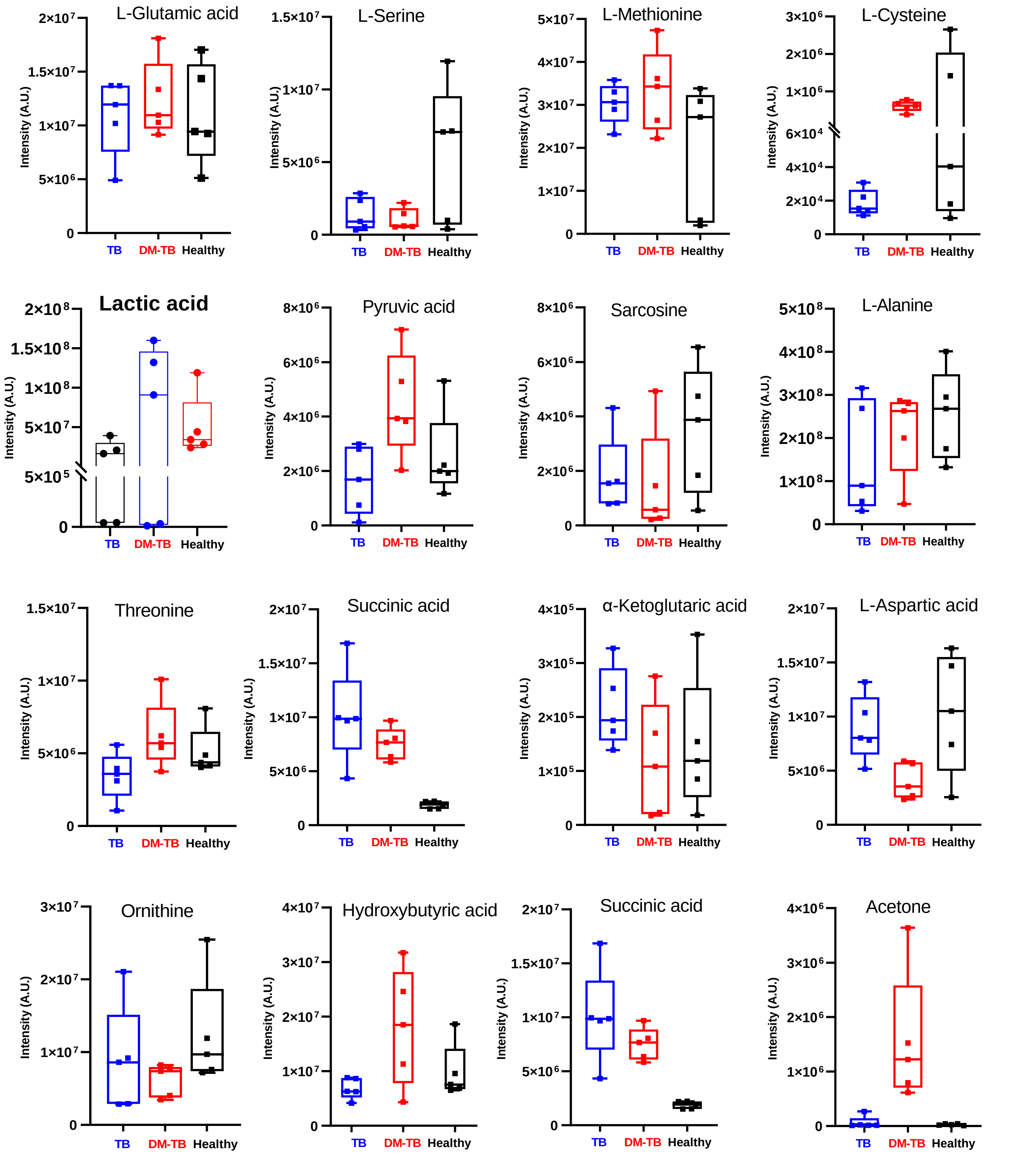


**Supplemental Figure S5**: Relative abundance of individual analytes involved in amino acid and acetone metabolism in the liver of Healthy controls and *Mycobacterium tuberculosis* H37Rv infected control C5BL/6 animals (TB) and streptozotocin induced diabetic animals (DM-TB). A.U.: arbitrary unit.

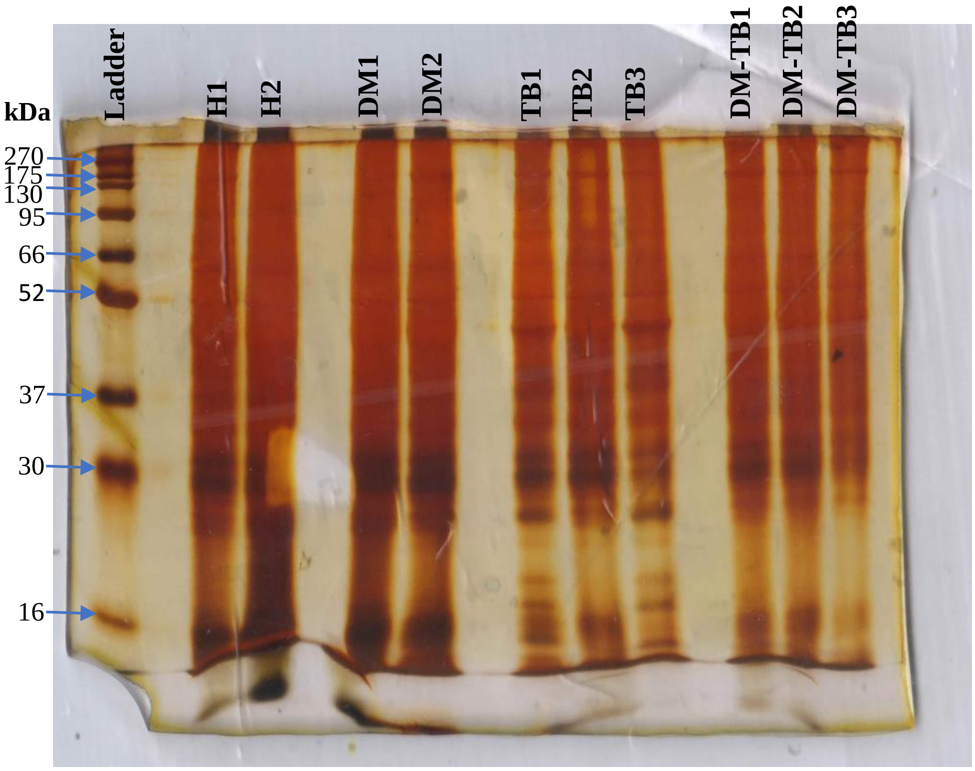


**Supplemental Figure S6**: Silver stained gel image of equal amount (4 µg) of liver proteins isolated from the healthy (H), Streptozotocin induced diabetic (DM), *Mycobacterium tuberculosis* H37Rv infected C57BL/6 mice (TB) and DM mice infected with *Mycobacterium tuberculosis* H37Rv (DM-TB). 1,2,3 shows the biological replicates.

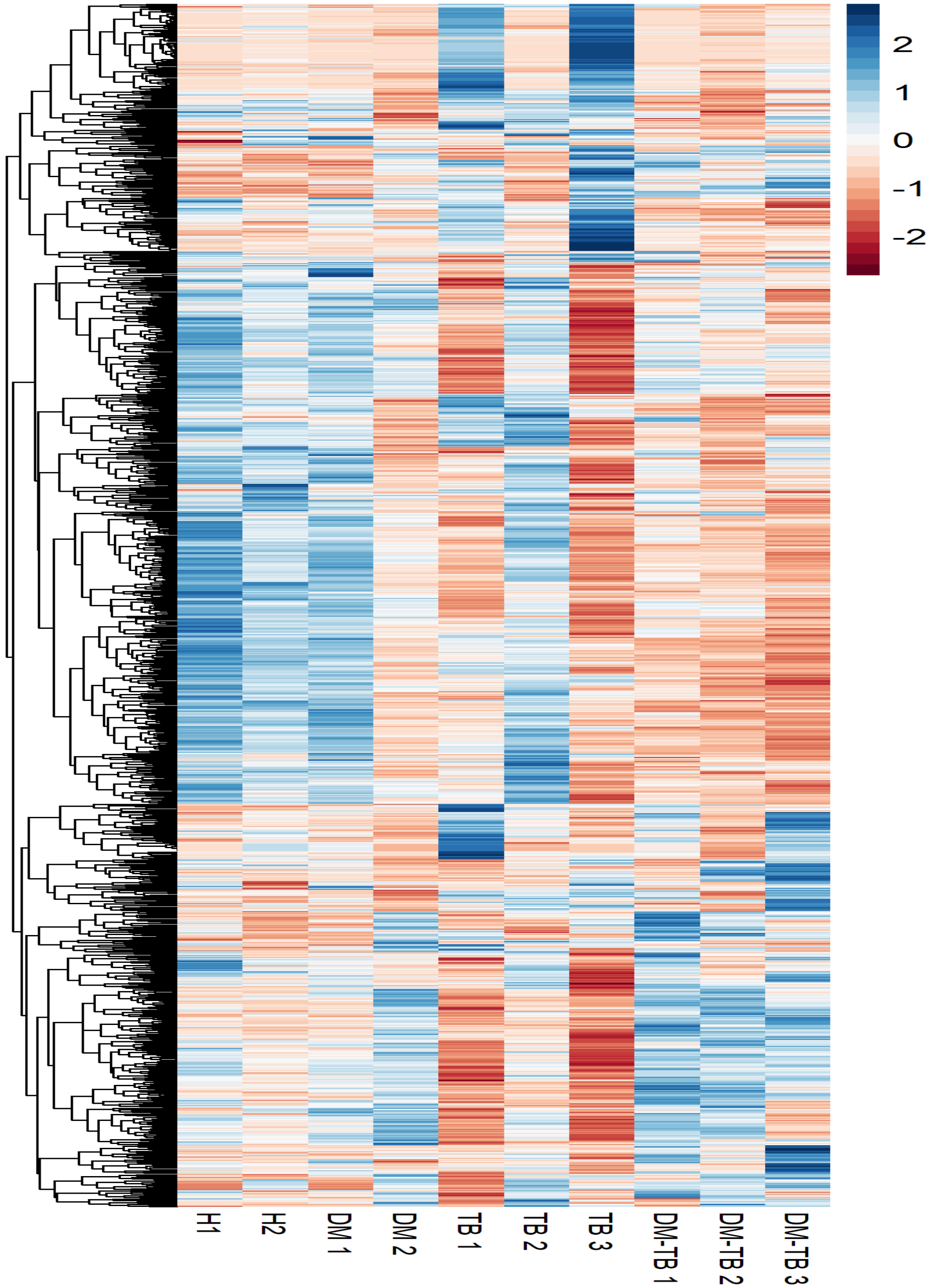


**Supplemental Figure S7:** Heatmap showing abundance of identified liver proteins (n=1833), obtained from the TMT multiplex experiment, using the healthy (H), Streptozotocin induced diabetic (DM), *Mycobacterium tuberculosis* H37Rv infected C57BL/6 mice (TB) and DM mice infected with *Mycobacterium tuberculosis* H37Rv (DM-TB). Each row presents abundance of individual protein. 1,2,3 shows the biological replicates. Colour intensity indicates the $log_2$fold change values.

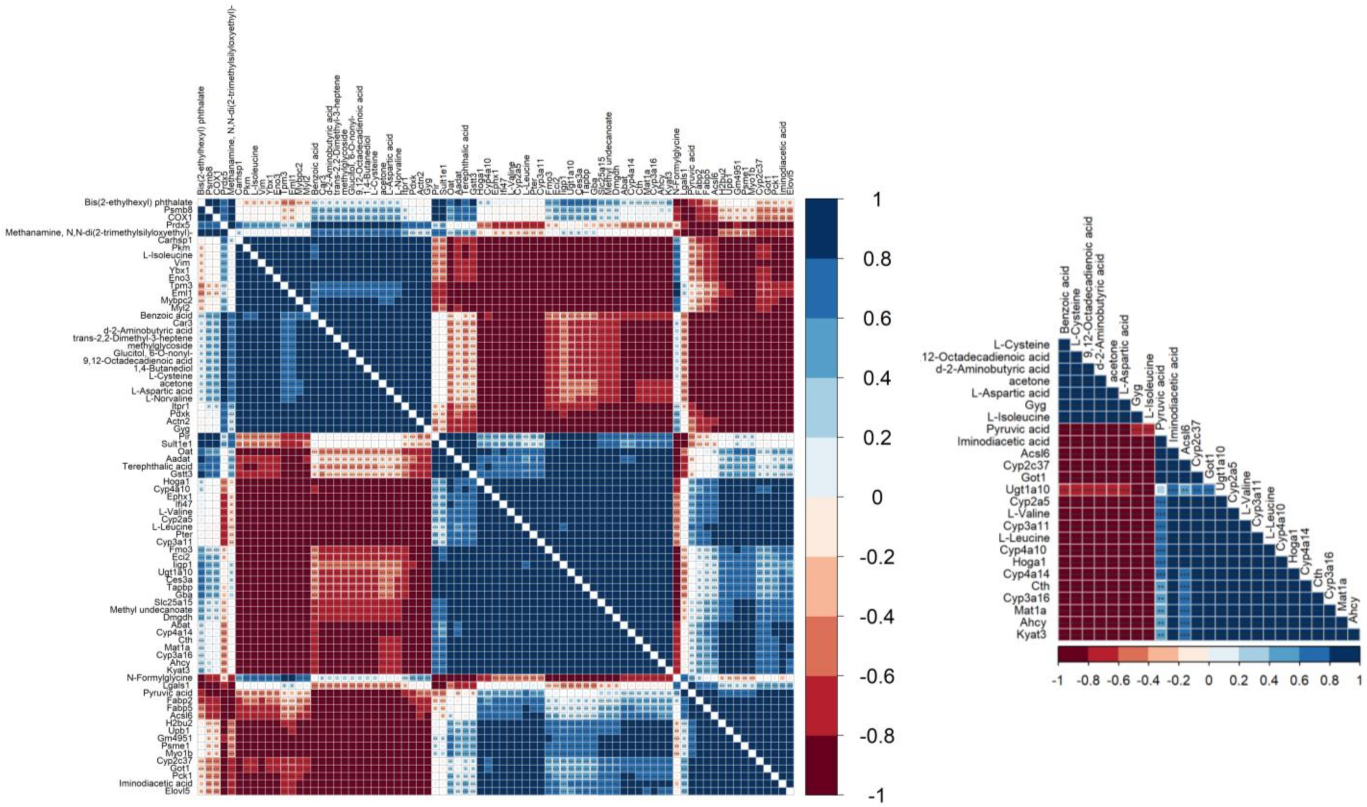


**Supplemental Figure S8:** Global correlation analysis of identified liver proteins and metabolites in *Mycobacterium tuberculosis* H37Rv infected C57BL/6 mice (TB). (A) Pairwise correlation of significantly deregulated (p-value $\leq 0.05$; $\log_2$ Fold change DM-TB/TB $\geq 1.0$) proteins and metabolites in TB group. ■/■: negative/ positive correlation. (B) A section of global correlation map showing correlation of proteins involved in cysteine-methionine and retinol metabolism proteins and their correlation with important metabolites in TB group.

**Supplementary Table S1:** List of identified lung metabolic features with their $log_2$ Fold change (DM-TB/TB) and $-log_{10}$ P-Value. TB/DM-TB: *Mycobacterium tuberculosis* H37Rv infected control/streptozotocin induced diabetes C57BL/6 mice.

| S.No. | Metabolic features | $log_2$(DM-TB/TB) | $-log_{10}$ P-value |
|---|---|---|---|
| 1 | Oxalic acid | 4.82 | 3.92 |
| 2 | L-Ornithine | 1.53 | 3.59 |
| 3 | Dihydrolycorine | 1.69 | 2.66 |
| 4 | d-2-Aminobutyric acid | 3.25 | 2.23 |
| 5 | Mevalonic acid | -3.14 | 1.96 |
| 6 | Xylonic acid, 1,5-lactone | 1.27 | 1.93 |
| 7 | Fenfluramine | 2.42 | 1.84 |
| 8 | Idonic acid, ?-lactone | 1.07 | 1.81 |
| 9 | 2-Bromomethyl-1,3-dioxolane | 1.75 | 1.75 |
| 10 | 1,4-Butanediol | 4.39 | 1.72 |
| 11 | Glycine | 0.95 | 1.68 |
| 12 | Urea | -2.36 | 1.64 |
| 13 | Malonic acid | 2.36 | 1.63 |
| 14 | Aminoisobutyric acid | 1.64 | 1.55 |
| 15 | Carbonic acid, propargyl 2-ethylhexyl ester | 2.38 | 1.49 |
| 16 | Phosphoric acid | 1.22 | 1.32 |
| 17 | Succinic acid | -2.43 | 1.23 |
| 18 | Arbutin | 2.07 | 1.22 |
| 19 | D- Erythrofuranose | 0.61 | 1.19 |
| 20 | Glyceric acid | 1.03 | 1.17 |
| 21 | Cl-5993 ditms | 1.42 | 1.13 |
| 22 | D-Ribonic acid, ?-lactone | 2.89 | 1.12 |
| 23 | Ile-pro-ile | 5.28 | 1.09 |
| 24 | 3-hydroxybutyric acid | 2.72 | 1.07 |
| 25 | D-Glucuronic acid c-lactone | 1.53 | 1.01 |
| 26 | 4-Hydroxybutyl acrylate | 2.52 | 1.00 |
| 27 | Formylglycine | -6.10 | 1.00 |
| 28 | Maleic acid | 0.66 | 0.99 |
| 29 | D-Lyxose | 5.19 | 0.99 |
| 30 | Citric acid | 0.81 | 0.98 |
| 31 | Hexadecane | -1.16 | 0.97 |
| 32 | dodecyl hexyl oxalate | 1.86 | 0.95 |
| 33 | D-Ribono-1,4-lactone | 0.38 | 0.94 |
| 34 | Butyl 2-ethylhexyl phthalate | 0.88 | 0.94 |
| 35 | D-Xylose | 2.93 | 0.93 |
| 36 | Triethylamine, 2-chloro-2'-(trimethyl)silyloxy- | 1.70 | 0.91 |
| 37 | Ethyl 4-ethoxybenzoate | 3.65 | 0.91 |
| 38 | L-Norvaline | 1.75 | 0.90 |
| 39 | Heptadecylamine | -1.10 | 0.90 |
| 40 | 2-Methyl-3-hydroxybutyric acid | 0.79 | 0.90 |
| 41 | L-Cysteine | 3.31 | 0.89 |
| 42 | [+]-2,4-Dihydroxy-3,3-dimethyl-N-[2-[N-(6-methoxy-8-quinolyl)sulfamyl]ethyl]butyramide | 1.45 | 0.88 |
| 43 | 6H-Purine-6-thione, 9-amino-1,9-dihydro- | 4.28 | 0.85 |

| | | | |
|---|---|---|---|
| 44 | Glycerol-3-phosphate | 0.62 | 0.83 |
| 45 | l-Leucine | 1.05 | 0.80 |
| 46 | L-Threonine | 1.28 | 0.79 |
| 47 | Indole-3-carbinol | 0.88 | 0.76 |
| 48 | Crotonic acid | 4.78 | 0.76 |
| 49 | acetone | 3.15 | 0.75 |
| 50 | 4-Ketoglucose | 0.77 | 0.75 |
| 51 | 2,5-Diphenylhexane | 2.84 | 0.73 |
| 52 | trans-2,2-Dimethyl-3-heptene | -0.46 | 0.72 |
| 53 | Oxalic acid, hexadecyl 2-isopropoxyphenyl ester | -0.91 | 0.72 |
| 54 | L-Phenylalanine | 0.36 | 0.70 |
| 55 | L-Isoleucine | 1.12 | 0.70 |
| 56 | L-Tyrosine | 0.45 | 0.68 |
| 57 | Lanthionine | 1.27 | 0.66 |
| 58 | 2-Keto-3-methylvaleric acid | -0.78 | 0.66 |
| 59 | Benzamide, N-(3-phenyl-2H-benzo[b]1,4-thiazin-2-ylideno)- | -1.21 | 0.64 |
| 60 | L-Valine | 0.81 | 0.63 |
| 61 | Pyruvic acid | 2.54 | 0.62 |
| 62 | Uracil | -0.30 | 0.62 |
| 63 | Benzenemethanol, a-[(butylamino)methyl]-4-hydroxy- | -0.70 | 0.61 |
| 64 | Indane-1,3-dione, 2-benzylaminomethylene- | -0.70 | 0.61 |
| 65 | 3-Phenyldodecane | -0.70 | 0.61 |
| 66 | Pyronene | -0.70 | 0.61 |
| 67 | 1-Ethylindan | -0.70 | 0.61 |
| 68 | Benzanilide | -0.70 | 0.61 |
| 69 | a-Aminoadipic acid | -0.70 | 0.61 |
| 70 | 1-Phenylethylamine, N-(4-bromobenzylidene) | -0.70 | 0.61 |
| 71 | Decanal, O-methyloxime | -0.70 | 0.61 |
| 72 | 2-Monolinolenin | -0.70 | 0.61 |
| 73 | 3-Pyridinecarbonitrile | 0.86 | 0.60 |
| 74 | L-Aspartic acid | -0.68 | 0.60 |
| 75 | L-Glutamic acid | 0.51 | 0.59 |
| 76 | Methylmalonic acid | -0.65 | 0.53 |
| 77 | Adenine | 0.98 | 0.53 |
| 78 | Ethyl tartrate | 0.42 | 0.50 |
| 79 | Talonic acid, ?-lactone | 1.49 | 0.48 |
| 80 | 3-Methoxy-4-methylheptane | 0.75 | 0.47 |
| 81 | L-Methionine | 0.45 | 0.46 |
| 82 | N-Acetyl aspartic acid | 0.85 | 0.46 |
| 83 | L-Serine | 0.27 | 0.46 |
| 84 | 2-Trifluoromethylbenzylamine, N,N-diundecyl- | 0.64 | 0.45 |
| 85 | N-formyl-Glycine | 0.93 | 0.44 |
| 86 | L-Alanine | 0.56 | 0.43 |
| 87 | Hexadecanoic acid | -1.13 | 0.39 |
| 88 | Methanamine, N,N-di(2-trimethylsilyloxyethyl)- | -0.46 | 0.39 |
| 89 | L-Lysine | 0.30 | 0.39 |
| 90 | L-Asparagine | 0.32 | 0.38 |
| 91 | Ethyl 2-hexenoate | -0.68 | 0.38 |

| | | | |
|---|---|---|---|
| 92 | Sebacic acid, 2-phenylphenyl undecyl ester | 0.98 | 0.37 |
| 93 | Niacinamide | 0.20 | 0.36 |
| 94 | 2-Hydroxyglutaric acid | 0.35 | 0.33 |
| 95 | Phosphoric acid, monomethyl ester | 0.28 | 0.33 |
| 96 | 4-Penten-2-one, 3-methyl-, O-methyloxime | 0.37 | 0.33 |
| 97 | Pent-4-enoic acid, 2-(4-hydroxybenzyl)- | -0.12 | 0.29 |
| 98 | Ornithine | 0.46 | 0.28 |
| 99 | Creatinine enol | 0.33 | 0.28 |
| 100 | Fumaric acid, 2,4-dimethylpent-3-yl octyl ester | 0.41 | 0.26 |
| 101 | allyl octadecyl oxalate | 0.33 | 0.25 |
| 102 | D-Mannopentadecane | 0.33 | 0.23 |
| 103 | Methyl undecanoate | -0.37 | 0.21 |
| 104 | Androst-2-en-17-amine, N-(2-phenylethyl)-, (5a)- | -0.29 | 0.20 |
| 105 | Oleic acid | -0.32 | 0.16 |
| 106 | 9H-Purin-6-ol | 0.19 | 0.15 |
| 107 | 2-Pyrrolidone-5-carboxylic acid | -0.19 | 0.15 |
| 108 | Cyclopentane-R1,(trans)-2-dicarboxylic acid, 3,3-dimethyl-4-methylene-(cis)-5-trimethylsilyl-, dimethyl ester | 0.16 | 0.14 |
| 109 | Hydroxyproline | 0.11 | 0.14 |
| 110 | Glycolic acid | -0.19 | 0.14 |
| 111 | Glycerol | 0.17 | 0.14 |
| 112 | Diisobutyl phthalate | -0.23 | 0.13 |
| 113 | Benzoic acid trimethylsilyl ester | 0.21 | 0.12 |
| 114 | Altronic acid, lactone | -0.17 | 0.11 |
| 115 | Lauric acid | 0.12 | 0.11 |
| 116 | Isobutyric acid | 0.18 | 0.11 |
| 117 | Dihydroobscurinervidinediol | 0.09 | 0.10 |
| 118 | Terephthalic acid | 0.16 | 0.09 |
| 119 | Methanephosphonothioic acid, O,O'-bis(trimethylsilyl) ester | -0.12 | 0.09 |
| 120 | 1,2-Diphenoxyethane | 0.15 | 0.08 |
| 121 | Glucitol, 6-O-nonyl- | -0.13 | 0.05 |
| 122 | Tetracyclo[5.1.0.0(2,4).0(3,5)]octane, 8,8-dichloro-exo-6-[(trimethylsilyl)oxy]- | -0.10 | 0.05 |
| 123 | a-Hydroxypyruvic acid | -0.07 | 0.04 |
| 124 | Lactic acid | 0.13 | 0.04 |
| 125 | 5-Methyl-2-propyldecahydroquinoline | -0.04 | 0.03 |
| 126 | Succinic acid, 3,5-difluorophenyl 2-(dimethylamino)ethyl ester | 0.02 | 0.02 |
| 127 | L-Fucose | 0.01 | 0.01 |
| 128 | Fumaric acid | 0.0008 | 0.001 |

**Supplementary Table S2:** List of identified liver metabolic features with their $log_2$ Fold change (DM-TB/TB) and $-log_{10}$ P-Value. TB/DM-TB: *Mycobacterium tuberculosis* H37Rv infected control/streptozotocin induced diabetes C57BL/6 mice.

| S.No. | Metabolic features | $log_2$(DM-TB/TB) | $-log_{10}$ P-value |
|---|---|---|---|
| 1 | trans-2,2-Dimethyl-3-heptene | 4.34 | 3.31 |
| 2 | L-Cysteine | 5.02 | 3.08 |
| 3 | 1,4-Butanediol | 5.73 | 2.41 |
| 4 | L-Leucine | 0.69 | 2.10 |
| 5 | d-2-Aminobutyric acid | 3.10 | 2.09 |
| 6 | Iminodiacetic acid | 3.91 | 1.97 |
| 7 | L-Aspartic acid | -1.15 | 1.73 |
| 8 | L-Isoleucine | 1.78 | 1.68 |
| 9 | methylglycoside | 9.86 | 1.64 |
| 10 | Glucitol, 6-O-nonyl- | 3.99 | 1.61 |
| 11 | L-Norvaline | 0.97 | 1.59 |
| 12 | Linoleic Acid | 4.14 | 1.53 |
| 13 | Pyruvic acid | 1.41 | 1.52 |
| 14 | L-Valine | 0.63 | 1.45 |
| 15 | N-Formylglycine | 1.37 | 1.39 |
| 16 | Terephthalic acid | -3.68 | 1.36 |
| 17 | Bis(2-ethylhexyl) phthalate | 1.02 | 1.36 |
| 18 | Methyl undecanoate | -1.30 | 1.35 |
| 19 | Benzoic acid | 1.94 | 1.33 |
| 20 | acetone | 4.55 | 1.29 |
| 21 | Methanamine, N,N-di(2-trimethylsilyloxyethyl)- | -0.98 | 1.26 |
| 22 | 1-Naphthoic acid, tridecynyl ester | 1.79 | 1.23 |
| 23 | Oxalic acid, hexadecyl 2-isopropoxyphenyl ester | -1.04 | 1.22 |
| 24 | N,N-diundecyl-2-Trifluoromethylbenzylamine | -1.77 | 1.20 |
| 25 | L-Threonine | 0.85 | 1.18 |
| 26 | L-Fucose | -3.99 | 1.17 |
| 27 | beta-Pyronene | 2.17 | 1.17 |
| 28 | sarcosine | -5.67 | 1.15 |
| 29 | Carbonic acid, propargyl 2-ethylhexyl ester | 0.79 | 1.13 |
| 30 | alpha-Hydroxypyruvic acid | 0.89 | 1.11 |
| 31 | alpha-Hydroxybutyric acid | 1.40 | 1.10 |
| 32 | D-Ribono lactone | -1.61 | 1.10 |
| 33 | D-Xylose | -3.67 | 1.09 |
| 34 | Oxalic acid, dodecyl hexyl ester | 2.85 | 1.08 |
| 35 | Pentanoic acid | 5.57 | 1.08 |
| 36 | Adenine | 1.44 | 1.08 |
| 37 | Fenfluramine | 1.64 | 1.01 |
| 38 | Pyroglutamic acid | 0.62 | 1.00 |
| 39 | n-Decanal | 1.74 | 0.99 |
| 40 | Tryptophol | -2.47 | 0.98 |
| 41 | Sebacic acid, di(3-hexyl) ester | -2.13 | 0.98 |
| 42 | Tryptamine | 4.38 | 0.96 |
| 43 | N-Acetyl aspartic acid | 4.56 | 0.95 |

| | | | |
|---|---|---|---|
| 44 | 3-Ethoxy-1,1,1,7,7,7-hexamethyl-3,5,5-tris(trimethylsiloxy)tetrasiloxane | 1.49 | 0.92 |
| 45 | Maleic acid | 1.44 | 0.91 |
| 46 | L-Serine | 0.74 | 0.90 |
| 47 | 1,5-Octadiene, 7-methyl-3-(1-methylethyl)- | -1.51 | 0.88 |
| 48 | Occlesterone | -0.87 | 0.86 |
| 49 | 4-Mercaptophenol | -1.43 | 0.85 |
| 50 | Indole-3-carbinol | -0.81 | 0.85 |
| 51 | Phosphoethanolamine | 0.75 | 0.83 |
| 52 | Uracil | 0.26 | 0.80 |
| 53 | Fumaric acid | 0.96 | 0.79 |
| 54 | Glycerophosphoric acid | -0.33 | 0.79 |
| 55 | trans-Vaccenic acid | 2.17 | 0.78 |
| 56 | hydroxyallopregnanolone | 0.84 | 0.78 |
| 57 | (2S,4aS,5R,8aR)-5-Methyl-2-propyldecahydroquinoline | -0.92 | 0.75 |
| 58 | Undecylenic acid | -1.69 | 0.75 |
| 59 | D-Ribofuranose | 1.34 | 0.75 |
| 60 | Citric acid | 2.10 | 0.75 |
| 61 | Glyceric acid | 0.61 | 0.74 |
| 62 | 3-Pyridinecarbonitrile | 0.53 | 0.74 |
| 63 | Ethyl 2-hexenoate | -0.72 | 0.72 |
| 64 | Mannonic acid, gamma-lactone | 2.57 | 0.66 |
| 65 | alpha-Hydroxyglutaric acid | -0.58 | 0.63 |
| 66 | 2,4-dimethyl-3-Pentanol | 0.35 | 0.59 |
| 67 | xylo-Hexos-5-ulose | -3.42 | 0.59 |
| 68 | D-Glucuronic acid gamma-lactone | -0.45 | 0.58 |
| 69 | 2-Bromomethyl-1,3-dioxolane | -1.27 | 0.57 |
| 70 | Succinic acid | -0.43 | 0.57 |
| 71 | Methylmalonic acid | -0.71 | 0.54 |
| 72 | Talonic acid | -0.91 | 0.54 |
| 73 | Arabinonic acid, gamma-lactone | -3.43 | 0.53 |
| 74 | methyl-2-O-acetyl-3,4-O-octylidene-alpha-D-Arabinopyranoside | -0.80 | 0.52 |
| 75 | Butanoic acid | 1.28 | 0.51 |
| 76 | 3,5,5-Trimethylhexene-1 | -0.75 | 0.50 |
| 77 | Glycine | 0.27 | 0.48 |
| 78 | Methanephosphonothioic acid | 0.55 | 0.47 |
| 79 | L-Asparagine | -0.31 | 0.47 |
| 80 | Dihydrolycorine | 0.40 | 0.46 |
| 81 | Phosphoric acid, monomethyl ester | 0.49 | 0.46 |
| 82 | Myristoleic acid | -0.99 | 0.45 |
| 83 | Cyclopentane-R1,(trans)-2-dicarboxylic acid, 3,3-dimethyl-4-methylene-(cis)-5-trimethylsilyl-, dimethyl ester | -0.56 | 0.45 |
| 84 | d-Galactose | -4.68 | 0.45 |
| 85 | Urea | -0.98 | 0.42 |
| 86 | Niacinamide | 0.27 | 0.42 |
| 87 | Hypoxanthine | -0.81 | 0.41 |
| 88 | L-Phenylalanine | 0.40 | 0.41 |

| | | | |
|---|---|---|---|
| 89 | Idonic acid, gamma-lactone | 0.38 | 0.41 |
| 90 | 2-Keto-3-methylvaleric acid | -0.65 | 0.40 |
| 91 | Creatinine | -0.67 | 0.38 |
| 92 | L-alanine | 0.51 | 0.38 |
| 93 | Galactopyranose | 0.71 | 0.35 |
| 94 | Norleucine | -0.69 | 0.35 |
| 95 | L-Histidine | -0.57 | 0.34 |
| 96 | Ornithine | -0.54 | 0.33 |
| 97 | Acetic acid | -0.42 | 0.33 |
| 98 | 4-Oxononanal | -0.66 | 0.33 |
| 99 | D-Ribonic acid-gamma-lactone | 0.33 | 0.31 |
| 100 | 2-(4-hydroxybenzyl)-Pent-4-enoic acid | 0.16 | 0.30 |
| 101 | Lactic acid | -0.62 | 0.30 |
| 102 | Ribitol | 0.00 | 0.29 |
| 103 | 2-Ketoisovaleric acid | 0.49 | 0.28 |
| 104 | 4-Penten-2-one, 3-methyl-, O-methyloxime | -0.72 | 0.28 |
| 105 | 4,2-Cresotic acid, 6-methoxy-, bimol. ester, methyl ester, 4,6-dimethoxy-o-toluate | -0.52 | 0.27 |
| 106 | L-Glutamic acid | 0.18 | 0.27 |
| 107 | Glycerol | 0.13 | 0.25 |
| 108 | Hexadecane | -0.45 | 0.24 |
| 109 | Dihydroobscurinervidinediol | 0.26 | 0.24 |
| 110 | Succinic acid, 3,5-difluorophenyl 2-(dimethylamino)ethyl ester | 0.15 | 0.23 |
| 111 | Altronic acid, gamma-lactone | -0.41 | 0.23 |
| 112 | Oleic acid | -0.39 | 0.22 |
| 113 | 2-Methyl-3-hydroxybutyric acid, | 0.20 | 0.21 |
| 114 | 1-Heptadecanamine | 0.36 | 0.20 |
| 115 | Hydroxyproline | -0.23 | 0.20 |
| 116 | D-Lyxose | 0.26 | 0.20 |
| 117 | Isobutyric acid | -0.18 | 0.19 |
| 118 | 4-Hydroxybutyl acrylate | -0.38 | 0.19 |
| 119 | D-Mannopentadecane-1,2,3,4,5-pentaol | -0.13 | 0.17 |
| 120 | 2-Methyl-3-oxo-N-(2-oxo-2-[(trimethylsilyl)oxy]ethyl)-N-(trimethylsilyl)-beta-alanine# | 0.45 | 0.17 |
| 121 | Fumaric acid, 2,4-dimethylpent-3-yl octyl ester | 0.27 | 0.16 |
| 122 | D- Erythrofuranose | -0.14 | 0.16 |
| 123 | 4-ethyl-2-methyl-2,4-Hexadiene | -0.24 | 0.16 |
| 124 | 1-ethyl-, ethylPyrrolidine-2-carboxylate | 0.24 | 0.16 |
| 125 | Hydroxynorleucine | 0.35 | 0.16 |
| 126 | Arbutin | 0.54 | 0.15 |
| 127 | L-Proline | 0.30 | 0.14 |
| 128 | Lauric acid | -0.13 | 0.14 |
| 129 | Keto-phenylpyruvic acid | -0.40 | 0.14 |
| 130 | 3-Methylbutyl S-thioacetate | 0.29 | 0.14 |
| 131 | alpha-D-Glucopyranoside | -0.16 | 0.13 |
| 132 | Glycerol-2-phosphate | -0.16 | 0.13 |
| 133 | l-Proline, N-methoxycarbonyl-, heptadecyl ester | -0.16 | 0.13 |
| 134 | alpha-Ketoglutaric acid | -0.16 | 0.13 |

| | | | |
|---|---|---|---|
| 135 | 1-Butanamine | -0.16 | 0.13 |
| 136 | 4,5-Pyrimidinediamine | -0.16 | 0.13 |
| 137 | Ile-pro-ile | -0.16 | 0.13 |
| 138 | Piperidine, 1,1'-(dithiodicarbonothioyl)bis- | -0.16 | 0.13 |
| 139 | Acetophenone | -0.16 | 0.13 |
| 140 | L-Sorbose | -0.16 | 0.13 |
| 141 | Oxalic acid, allyl octadecyl ester | -0.16 | 0.13 |
| 142 | 3-Aminobutyric acid | -0.16 | 0.13 |
| 143 | 1-Ethylindan | -0.16 | 0.13 |
| 144 | Octanoic acid | -0.16 | 0.13 |
| 145 | Aminoadipic acid | -0.28 | 0.12 |
| 146 | Sebacic acid, 2-phenylphenyl undecyl ester | -0.29 | 0.11 |
| 147 | Ethyl tartrate | -0.29 | 0.09 |
| 148 | 1-diethylamino-4,4-dimethyl-Pent-1-en-3-one | 0.18 | 0.08 |
| 149 | 4-Ketoglucose | 0.10 | 0.08 |
| 150 | 2,5,5,8a-Tetramethyl-3-oxo-3,4,4a,5,6,7,8,8a-octahydronaphthalene-1-carboxylic acid | 0.11 | 0.06 |
| 151 | Cl-5993 | 0.11 | 0.05 |
| 152 | 9-amino-1,9-dihydro-6H-Purine-6-thione | 0.11 | 0.05 |
| 153 | Oxalic acid | -0.12 | 0.04 |
| 154 | 4-phenyl-1-butene | 0.13 | 0.04 |
| 155 | L-Tyrosine | -0.05 | 0.04 |
| 156 | L-Methionine | -0.04 | 0.04 |
| 157 | 2-Oxopentanoic acid | -0.04 | 0.03 |
| 158 | Diisobutyl phthalate | -0.08 | 0.02 |
| 159 | D-Xylono- 1,5-lactone | -0.01 | 0.01 |
| 160 | L-Lysine | -0.01 | 0.01 |

**Supplementary Table S3:** Identified liver proteins with their absolute abundance in each sample obtained from the multiplex TMT experiment using Proteome Discoverer analysis. DM: streptozotocin induced diabetes C57BL/6 mice; TB/DM-TB: *Mycobacterium tuberculosis* H37RV infected control/DM mice.

| S.No. | Description | Absolute Abundance | | | | | | | | | |
|---|---|---|---|---|---|---|---|---|---|---|---|
| | | H1 | H2 | DM 1 | DM 2 | TB 1 | TB 2 | TB 3 | DM-TB 1 | DM-TB 2 | DM-TB 3 |
| 1 | Immunoglobulin kappa variable 6-13 OS=Mus musculus OX=10090 GN=Igkv6-13 PE=1 SV=7 | | 125.3 | | 121.8 | | 122.4 | 188.4 | 153.2 | 120.9 | 168 |
| 2 | Immunoglobulin heavy constant gamma 2B (Fragment) OS=Mus musculus OX=10090 GN=Ighg2b PE=1 SV=1 | 79.3 | 99.1 | 118.4 | 68.3 | 112 | 70.5 | 167.5 | 130.9 | 63.5 | 90.5 |
| 3 | Immunoglobulin heavy variable 1-22 (Fragment) OS=Mus musculus OX=10090 GN=Ighv1-22 PE=1 SV=1 | 107.9 | 123.1 | 109.1 | 94.1 | 88.2 | 128.4 | 86.9 | 101.8 | 83.2 | 77.4 |
| 4 | Immunoglobulin heavy constant alpha (Fragment) OS=Mus musculus OX=10090 GN=Igha PE=1 SV=1 | 85.9 | 109.4 | 105 | 139.7 | 61.7 | 84.7 | 71.6 | 127.5 | 112.8 | 101.7 |
| 5 | Mitochondrial amidoxime-reducing component 1 OS=Mus musculus OX=10090 GN=Mtarc1 PE=1 SV=1 | 102.1 | 85.7 | 97.8 | 94.2 | 108.3 | 99.7 | 141.1 | 95.5 | 86.9 | 88.7 |
| 6 | Actin-like protein 3 (Fragment) OS=Mus musculus OX=10090 GN=Actr3 PE=1 SV=1 | | 136 | 140.4 | 105.8 | 94 | 155.4 | 64 | 90.4 | 110.8 | 103.2 |
| 7 | Alcohol dehydrogenase iron-containing protein 1 (Fragment) OS=Mus musculus OX=10090 GN=Adhfe1 PE=1 SV=1 | 99.1 | 99.2 | 103.7 | 117.8 | 88.3 | 94.4 | 84.9 | 109.1 | 95.5 | 108.1 |
| 8 | Eukaryotic translation elongation factor 1 beta 2 OS=Mus musculus OX=10090 GN=Eef1b2 PE=1 SV=1 | 90 | 99 | 83.8 | 54.9 | 180.9 | 126.2 | 45.3 | 102.1 | 77.2 | 140.6 |
| 9 | Arf-GAP domain and FG repeat-containing protein 1 OS=Mus musculus OX=10090 GN=Agfg1 PE=1 SV=1 | 90.9 | 85.9 | 103.6 | 96.1 | 107.8 | 112.4 | 93.1 | 90.3 | 94.5 | 125.6 |
| 10 | Glutaryl-CoA dehydrogenase, mitochondrial OS=Mus musculus OX=10090 GN=Gcdh PE=1 SV=1 | 85.2 | 82.3 | 94 | 128.7 | 73.2 | 95.1 | 44.4 | 119.4 | 136.6 | 141.1 |
| 11 | Eukaryotic translation initiation factor 5A (Fragment) OS=Mus musculus OX=10090 GN=Eif5a PE=1 SV=1 | 90.7 | 91.2 | 98.7 | 89.6 | 154.6 | 99.2 | 99.9 | 99.4 | 74.9 | 101.9 |
| 12 | Complement factor H (Fragment) OS=Mus musculus OX=10090 GN=Cfh PE=1 SV=1 | 97.9 | 109.8 | 125.1 | 77 | 101.7 | 101.2 | 96.7 | 116.2 | 83 | 91.4 |
| 13 | 60S ribosomal protein L31 OS=Mus musculus OX=10090 GN=Rpl31 PE=1 SV=1 | 115.5 | 90.9 | 98.2 | 88 | 133.7 | 108.5 | 96.3 | 104.8 | 76 | 88.1 |
| 14 | Complex I-49kD (Fragment) OS=Mus musculus OX=10090 GN=Ndufs2 PE=1 SV=5 | 98.4 | 85.2 | 95.8 | 110.7 | 80.2 | 87.3 | 126.3 | 110.2 | 106.3 | 99.7 |
| 15 | Mitochondrial pyruvate carrier 2 OS=Mus musculus OX=10090 GN=Mpc2 PE=1 SV=1 | 118.3 | 123 | 112.2 | 123.2 | 87.4 | 97 | 62.6 | 100.8 | 92.5 | 82.9 |
| 16 | Glutamate--cysteine ligase regulatory subunit (Fragment) OS=Mus musculus OX=10090 GN=Gclm PE=1 SV=1 | 97.4 | 106.8 | 101 | 80.3 | 196.4 | 96.2 | 111.7 | 74.7 | 70.9 | 64.6 |
| 17 | ATP-binding cassette sub-family D member 3 OS=Mus musculus OX=10090 GN=Abcd3 PE=1 SV=1 | 131.8 | 120.5 | 133.4 | 103.6 | 76.6 | 100.7 | 69.8 | 91.6 | 100.6 | 71.4 |
| 18 | Glucosylceramidase OS=Mus musculus OX=10090 GN=Gba PE=1 SV=1 | | 112.7 | 123.9 | 116.4 | 54.6 | 88.3 | 63.9 | 145.6 | 120.2 | 174.4 |
| 19 | 40S ribosomal protein S27 (Fragment) OS=Mus musculus OX=10090 GN=Rps27 PE=1 SV=1 | 130.3 | 106.2 | 94.9 | 87.6 | 94.6 | 121.8 | 72.2 | 113.7 | 81.4 | 97.2 |
| 20 | Mevalonate kinase OS=Mus musculus OX=10090 GN=Mvk PE=1 SV=1 | 134.9 | 113.4 | 139.1 | 101.7 | 83.4 | 107.6 | 63.7 | 90.2 | 87.4 | 78.6 |
| 21 | Methylcrotonoyl-CoA carboxylase subunit alpha, mitochondrial OS=Mus musculus OX=10090 GN=Mccc1 PE=1 SV=1 | 96.9 | 91.6 | 95 | 112.7 | 71 | 91.9 | 87.9 | 111.8 | 128.9 | 112.4 |
| 22 | Cysteine-S-conjugate beta-lyase 2 (Fragment) OS=Mus musculus OX=10090 GN=Kyat3 PE=1 SV=1 | 82.3 | 77.2 | 80.6 | 152 | 55.5 | 81.6 | 54.2 | 161.8 | 136.2 | 118.6 |
| 23 | Glycogen debrancher OS=Mus musculus OX=10090 GN=Agl PE=1 SV=1 | 135 | 115.1 | 115 | 87.6 | 90 | 94.9 | 114.9 | 85.8 | 81.6 | 80 |
| 24 | Gc-globulin (Fragment) OS=Mus musculus OX=10090 GN=Gc PE=1 SV=1 | 89.9 | 138.2 | 96.7 | 107.6 | 95.2 | 124.4 | 84.2 | 89.9 | 79.2 | 94.6 |
| 25 | Annexin A5 (Fragment) OS=Mus musculus OX=10090 GN=Anxa5 PE=1 SV=1 | 123.1 | 98.9 | 120.8 | 105.7 | 112.4 | 141.4 | 72.1 | 75 | 85.3 | 65.5 |
| 26 | ATP synthase subunit b (Fragment) OS=Mus musculus OX=10090 GN=Atp5pb PE=1 SV=1 | 122 | 90.5 | 106.1 | 98.2 | 108.5 | 91.8 | 106.4 | 95.1 | 100.9 | 80.7 |
| 27 | Dihydropteridine reductase OS=Mus musculus OX=10090 GN=Qdpr PE=1 SV=1 | 20.6 | 145.3 | 153.4 | 123.8 | 83.3 | 127.3 | 74 | 103.6 | 97 | 71.6 |
| 28 | Ketohexokinase OS=Mus musculus OX=10090 GN=Khk PE=1 SV=1 | 125.5 | 119.1 | 108.8 | 106.2 | 76.1 | 110.2 | 73.5 | 95.2 | 85.8 | 99.6 |
| 29 | Mitochondrial glutamate carrier 1 OS=Mus musculus OX=10090 GN=Slc25a22 PE=1 SV=1 | 81.4 | 73.1 | 98.7 | 151.8 | 66.5 | 71.4 | 62.9 | 145 | 140.7 | 108.5 |
| 30 | Eukaryotic translation initiation factor 4 gamma 1 OS=Mus musculus OX=10090 GN=Eif4g1 PE=1 SV=1 | 109.1 | 109.9 | 126.1 | 95.2 | 84.3 | 114.6 | 77.7 | 102.7 | 90.9 | 89.5 |

| | | | | | | | | | | | |
|---|---|---|---|---|---|---|---|---|---|---|---|
| 31 | Electron transfer flavoprotein subunit beta OS=Mus musculus OX=10090 GN=Etfbl PE=3 SV=1 | 119.8 | 106.9 | 112.5 | 127.1 | 92 | 99 | 75.7 | 93.3 | 93.5 | 80.3 |
| 32 | Inositol 1,4,5-trisphosphate receptor type 1 (Fragment) OS=Mus musculus OX=10090 GN=Itpr1 PE=1 SV=1 | 87.6 | 75.7 | 87 | 73.2 | 140.4 | 100 | 218.1 | 84.3 | 56.8 | 76.9 |
| 33 | Cytochrome c oxidase subunit NDUFA4 OS=Mus musculus OX=10090 GN=Ndufa4 PE=1 SV=1 | 82.3 | 82 | 82.6 | 101.5 | 119.9 | 79.7 | 119 | 126.6 | 102 | 104.3 |
| 34 | Cytochrome P450 OS=Mus musculus OX=10090 GN=Cyp2d26 PE=1 SV=1 | | 123 | 152.2 | 162.6 | 48.7 | 91.8 | 48.5 | 142.4 | 160.9 | 70 |
| 35 | Histidine-rich glycoprotein OS=Mus musculus OX=10090 GN=Hrg PE=1 SV=1 | 86.9 | 77.4 | 119.6 | 105.8 | 95.5 | 99.9 | 51.9 | 134.6 | 127.1 | 101.3 |
| 36 | Antigen-presenting glycoprotein CD1d1 OS=Mus musculus OX=10090 GN=Cd1d1 PE=1 SV=1 | 124.4 | 125.3 | 128.6 | 106.2 | 80.7 | 115.1 | 72.5 | 89.4 | 90.5 | 67.1 |
| 37 | Phospholipase B-like OS=Mus musculus OX=10090 GN=Plbd1 PE=1 SV=1 | 107.8 | 95.7 | 100.2 | 91.2 | 92.3 | 107.8 | 67.9 | 110.5 | 101.1 | 125.6 |
| 38 | CMP-N-acetylneuraminic acid synthase OS=Mus musculus OX=10090 GN=Cmas PE=1 SV=1 | 114.6 | 94.8 | 119.2 | 92.7 | 80.5 | 110.7 | 81.1 | 105 | 96.1 | 105.5 |
| 39 | Prolow-density lipoprotein receptor-related protein 1 OS=Mus musculus OX=10090 GN=Lrp1 PE=1 SV=1 | 112.9 | 100.6 | 109.9 | 84.9 | 115 | 122.6 | 99.6 | 93.9 | 74 | 86.4 |
| 40 | Acyl-peptide hydrolase (Fragment) OS=Mus musculus OX=10090 GN=Apeh PE=1 SV=1 | 128.5 | 112.5 | 121.8 | 86 | 110.9 | 122 | 90.2 | 79.5 | 78.9 | 69.6 |
| 41 | NADH dehydrogenase [ubiquinone] 1 alpha subcomplex subunit 12 OS=Mus musculus OX=10090 GN=Ndufa12 PE=1 SV=1 | 108.2 | 109.6 | 89.6 | 104.1 | 98.1 | 82.9 | 108.6 | 126.3 | 98.5 | 74 |
| 42 | Fatty acid synthase OS=Mus musculus OX=10090 GN=Fasn PE=1 SV=1 | 199.9 | 119.1 | 159.2 | 83 | 69.6 | 130.8 | 53.4 | 66.1 | 66.2 | 52.6 |
| 43 | Isocitrate dehydrogenase [NADP], mitochondrial (Fragment) OS=Mus musculus OX=10090 GN=Idh2 PE=1 SV=1 | 71.4 | 68.7 | 92 | 106.9 | 138.5 | 81.2 | 177.6 | 89.2 | 82.3 | 92.3 |
| 44 | MICOS complex subunit MIC60 OS=Mus musculus OX=10090 GN=Immt PE=1 SV=1 | 89 | 90.5 | 93.2 | 101.1 | 94.2 | 104.9 | 74.4 | 119.6 | 113.1 | 120 |
| 45 | Myosin regulatory light chain 2, skeletal muscle isoform OS=Mus musculus OX=10090 GN=Mylpf PE=1 SV=1 | 26.7 | 40.7 | 19.1 | 27.2 | 342.8 | 24.2 | 414.6 | 29.1 | 21.5 | 54 |
| 46 | NADH dehydrogenase [ubiquinone] 1 subunit C2 OS=Mus musculus OX=10090 GN=Ndufc2 PE=1 SV=1 | 72.8 | 74.1 | 74.4 | 92.6 | 107.8 | 75.3 | 159.8 | 99.1 | 91.1 | 153 |
| 47 | Flavin reductase (NADPH) OS=Mus musculus OX=10090 GN=Blvrb PE=1 SV=1 | 80.6 | 120.1 | 121.1 | 109.2 | 90.2 | 112 | 64.4 | 93.4 | 106.3 | 102.7 |
| 48 | ADP-ribosylation factor-like protein 6-interacting protein 1 (Fragment) OS=Mus musculus OX=10090 GN=Arl6ip1 PE=1 SV=1 | 121.1 | 108.9 | 102.1 | 124.4 | 85.3 | 71.1 | 74.8 | 109.1 | 123.2 | 80.2 |
| 49 | Methylmalonyl-CoA epimerase, mitochondrial OS=Mus musculus OX=10090 GN=Mcee PE=1 SV=1 | | 144 | | 152.9 | | 147.1 | 107.5 | 152.8 | 141.3 | 154.5 |
| 50 | 40S ribosomal protein S13 OS=Mus musculus OX=10090 GN=Rps13 PE=1 SV=1 | 118.6 | 114.7 | 125.4 | 93.6 | 107.2 | 113.4 | 64 | 104.7 | 91.4 | 67.1 |
| 51 | 40S ribosomal protein S3 OS=Mus musculus OX=10090 GN=Rps3 PE=1 SV=1 | 142.2 | 112 | 121.1 | 89.8 | 86 | 120.7 | 66.4 | 107.4 | 91.7 | 62.6 |
| 52 | Cytochrome c oxidase subunit 6B1 OS=Mus musculus OX=10090 GN=Cox6b1 PE=1 SV=1 | 82.5 | 84.1 | 86.5 | 103.1 | 142.8 | 93 | 100.9 | 117.8 | 93.6 | 95.7 |
| 53 | 60S ribosomal protein L9 OS=Mus musculus OX=10090 GN=Rpl9-ps6 PE=3 SV=1 | 122 | 119.5 | 110.7 | 84.7 | 110.8 | 110.6 | 109 | 100.4 | 71.3 | 61 |
| 54 | L-lactate dehydrogenase C chain OS=Mus musculus OX=10090 GN=Ldhc PE=1 SV=1 | 150 | 112.4 | 123.5 | 87.1 | 89.6 | 150 | 54.5 | 88.1 | 75.8 | 68.9 |
| 55 | Carbonyl reductase (NADPH) OS=Mus musculus OX=10090 GN=Gm5678 PE=3 SV=1 | 105.5 | 95.8 | 84.1 | 109.2 | 93.7 | 80.9 | 67.2 | 141.9 | 101.4 | 120.1 |
| 56 | 40S ribosomal protein S11 OS=Mus musculus OX=10090 GN=Rps11 PE=1 SV=1 | 128.9 | 110.4 | 124.9 | 90.3 | 86.9 | 124.7 | 74 | 101.7 | 84.6 | 73.6 |
| 57 | Pre-mRNA-processing factor 40 homolog A OS=Mus musculus OX=10090 GN=Prpf40a PE=1 SV=1 | 92.6 | 100.9 | 92.3 | 81 | 112.6 | 103.8 | 102.8 | 117.4 | 81.2 | 115.4 |
| 58 | ATP-binding cassette sub-family C member 6 (Fragment) OS=Mus musculus OX=10090 GN=Abcc6 PE=1 SV=1 | 138.8 | 113 | 123.7 | 95.7 | 70.9 | 119.3 | 65.3 | 95.3 | 94.6 | 83.3 |
| 59 | NAD-dependent protein deacetylase OS=Mus musculus OX=10090 GN=Sirt3 PE=1 SV=1 | 100.7 | 91.3 | 93.5 | 100.3 | 106 | 95.1 | 109.7 | 103.7 | 96.3 | 103.3 |
| 60 | 60S ribosomal protein L18 (Fragment) OS=Mus musculus OX=10090 GN=Rpl18 PE=1 SV=1 | 100.5 | 121.9 | 107.9 | 64.8 | 95.9 | 109.9 | 113.4 | 108.7 | 85.8 | 91.2 |
| 61 | 40S ribosomal protein S11 (Fragment) OS=Mus musculus OX=10090 GN=Rps11 PE=1 SV=1 | 126.4 | 119.3 | 125.6 | 92.8 | 87.4 | 125.4 | 59.2 | 102.4 | 87.9 | 73.5 |
| 62 | 60S ribosomal protein L13a OS=Mus musculus OX=10090 GN=Rpl13a PE=3 SV=1 | 126 | 124.4 | 119.6 | 96.8 | 64.9 | 116.6 | 56.4 | 114.2 | 90.8 | 90.3 |
| 63 | Aldehyde dehydrogenase family 16 member A1 OS=Mus musculus OX=10090 GN=Aldh16a1 PE=1 SV=1 | 127.6 | 110.7 | 104.6 | 91.1 | 93.1 | 118.4 | 80 | 104.3 | 85.4 | 84.7 |
| 64 | Peptide-methionine (S)-S-oxide reductase OS=Mus musculus OX=10090 GN=Msra PE=1 SV=1 | 109 | 133.5 | 93.1 | 109.4 | 88.1 | 115.3 | 73.9 | 100.1 | 97.3 | 80.3 |
| 65 | 60S ribosomal protein L10 (Fragment) OS=Mus musculus OX=10090 GN=Rpl10 PE=4 SV=1 | 134.4 | 104.7 | 108.1 | 89.7 | 95.6 | 132 | 76.4 | 104.1 | 77.9 | 77.1 |

| | | | | | | | | | | | |
|---|---|---|---|---|---|---|---|---|---|---|---|
| 66 | Glyceraldehyde-3-phosphate dehydrogenase OS=Mus musculus OX=10090 GN=Gm10358 PE=1 SV=1 | 99.3 | 90.5 | 106.8 | 99.8 | 119.1 | 105.3 | 119.5 | 91.8 | 93.2 | 74.9 |
| 67 | Metallothionein-1 OS=Mus musculus OX=10090 GN=Mt1 PE=1 SV=1 | 19.3 | 7.6 | 17 | 31.6 | 34.3 | 38.2 | 26.4 | 351.4 | 33.3 | 440.9 |
| 68 | Coactosin-like protein OS=Mus musculus OX=10090 GN=Cotl1 PE=1 SV=1 | 125.5 | 102.9 | 119.3 | 89.3 | 107.9 | 194.4 | 49.2 | 71.9 | 61.8 | 77.8 |
| 69 | Adenine phosphoribosyltransferase OS=Mus musculus OX=10090 GN=Aprt PE=1 SV=1 | 80.1 | 69.8 | 75.3 | 138.6 | 75.2 | 109.4 | 68.6 | 151.5 | 133.1 | 98.5 |
| 70 | 60S ribosomal protein L18a (Fragment) OS=Mus musculus OX=10090 GN=Rpl18a PE=1 SV=1 | 160.9 | 136.8 | 112.7 | 89.7 | 84.8 | 112.3 | 57 | 116.6 | 89.4 | 39.8 |
| 71 | Hypoxia up-regulated protein 1 (Fragment) OS=Mus musculus OX=10090 GN=Hyou1 PE=1 SV=1 | 119.7 | 110.9 | 140.1 | 88.1 | 89.9 | 131.5 | 82.4 | 98.4 | 70.1 | 69 |
| 72 | 40S ribosomal protein S25 OS=Mus musculus OX=10090 GN=Rps25 PE=1 SV=1 | 116.5 | 98.4 | 104 | 101.9 | 108.2 | 115.4 | 103.2 | 93.4 | 90.3 | 68.5 |
| 73 | N-acyl-L-amino-acid amidohydrolase OS=Mus musculus OX=10090 GN=Acy1 PE=1 SV=1 | 108.8 | 107.8 | 117.7 | 98.8 | 80.8 | 113.7 | 90.6 | 94.2 | 89.8 | 97.8 |
| 74 | Peptidyl-prolyl cis-trans isomerase OS=Mus musculus OX=10090 GN=Ppia PE=1 SV=1 | 120 | 110 | 100 | 88.5 | 108.1 | 114.7 | 85.6 | 100.2 | 84.1 | 88.7 |
| 75 | Leucine--tRNA ligase OS=Mus musculus OX=10090 GN=Lars2 PE=1 SV=1 | 101.8 | 93 | 103.7 | 108 | 48.2 | 96 | 87.9 | 124.9 | 114.7 | 121.7 |
| 76 | Isocitrate dehydrogenase [NAD] subunit, mitochondrial OS=Mus musculus OX=10090 GN=Idh3a PE=1 SV=1 | 49.5 | 78 | 70.2 | 90.3 | 149.8 | 78.7 | 208.1 | 90.6 | 83.7 | 101 |
| 77 | Adrenodoxin, mitochondrial OS=Mus musculus OX=10090 GN=Fdx1 PE=1 SV=1 | 94.2 | 113.9 | 91.7 | 73.5 | 126.6 | 99.2 | 85.6 | 95.7 | 84.1 | 135.4 |
| 78 | 60S ribosomal protein L29 (Fragment) OS=Mus musculus OX=10090 GN=Rpl29 PE=1 SV=1 | 118.7 | 91.7 | 111.4 | 81.7 | 106.7 | 110.6 | 67 | 128.2 | 89.5 | 94.6 |
| 79 | Alpha-actinin-4 OS=Mus musculus OX=10090 GN=Actn4 PE=1 SV=1 | 111.1 | 100.4 | 95.8 | 88.9 | 101.2 | 109.1 | 93.4 | 114.8 | 84.4 | 100.9 |
| 80 | Glutathione S-transferase theta-2 OS=Mus musculus OX=10090 GN=Gstt2 PE=1 SV=1 | 81.2 | 58.3 | 138.1 | 134.7 | 58 | 121.4 | 64.8 | 159.4 | 114.8 | 69.4 |
| 81 | Histone H3.2 OS=Mus musculus OX=10090 GN=H3c14 PE=1 SV=1 | 80.6 | 131.9 | 107.1 | 62.3 | 197.6 | 59.8 | 73.3 | 88.3 | 52.8 | 146.3 |
| 82 | 4a-hydroxytetrahydrobiopterin dehydratase OS=Mus musculus OX=10090 GN=Pcbd1 PE=1 SV=1 | 109.9 | 132 | 95.4 | 76.5 | 134.1 | 125.9 | 103.9 | 102.1 | 57.3 | 63 |
| 83 | Acid phosphatase OS=Mus musculus OX=10090 GN=Acp1 PE=1 SV=1 | 134.2 | 116.4 | 112.6 | 98.9 | 86.9 | 113.2 | 72.2 | 92.8 | 84.7 | 88.1 |
| 84 | CCT-beta (Fragment) OS=Mus musculus OX=10090 GN=Cct2 PE=1 SV=1 | 114.1 | 111.6 | 115.9 | 94.5 | 93.4 | 116.3 | 92.4 | 91.5 | 89.2 | 81.3 |
| 85 | Glutamine amidotransferase-like class 1 domain-containing 3A (Fragment) OS=Mus musculus OX=10090 GN=Gatd3a PE=1 SV=1 | | 135.8 | | 142 | | 129.3 | 152.1 | 157.3 | 143.6 | 139.9 |
| 86 | Methylmalonate-semialdehyde dehydrogenase [acylating], mitochondrial (Fragment) OS=Mus musculus OX=10090 GN=Aldh6a1 PE=1 SV=1 | 114.6 | 93.2 | 110.7 | 112.6 | 64.1 | 99.1 | 78.4 | 125.3 | 96.7 | 105.2 |
| 87 | Flavoprotein subunit of complex II (Fragment) OS=Mus musculus OX=10090 GN=Sdha PE=1 SV=1 | 114 | 96 | 91.9 | 120.5 | 103.2 | 105.5 | 80.8 | 101.8 | 102.6 | 83.7 |
| 88 | Cofilin-2 OS=Mus musculus OX=10090 GN=Cfl2 PE=1 SV=1 | 105.5 | 103.8 | 102.2 | 89.5 | 128.3 | 91 | 144.3 | 75.2 | 73.1 | 87.1 |
| 89 | Isoamyl acetate-hydrolyzing esterase 1 homolog (Fragment) OS=Mus musculus OX=10090 GN=Iah1 PE=1 SV=1 | 142.8 | 134.5 | 134.8 | 96.5 | 86.2 | 102.1 | 52.2 | 80.4 | 88.7 | 81.8 |
| 90 | Maleylacetoacetate isomerase OS=Mus musculus OX=10090 GN=Gstz1 PE=1 SV=1 | 156.8 | 111.2 | 114.7 | 100.6 | 71.9 | 98.6 | 75.7 | 80.7 | 91.6 | 98.1 |
| 91 | Estradiol 17 beta-dehydrogenase 5 OS=Mus musculus OX=10090 GN=Akr1c6 PE=1 SV=1 | 158.2 | 146.4 | 135.8 | 81.1 | 95.2 | 151.7 | 52.9 | 60.8 | 63.1 | 55 |
| 92 | APOBEC1 complementation factor OS=Mus musculus OX=10090 GN=A1cf PE=1 SV=1 | 147.1 | 119.7 | 128.7 | 79.5 | 118.7 | 121.6 | 49.3 | 85.9 | 77.9 | 71.7 |
| 93 | Inosine phosphorylase (Fragment) OS=Mus musculus OX=10090 GN=Gm49342 PE=1 SV=1 | 136.5 | 100.3 | 123.7 | 104.5 | 72.9 | 106.9 | 73.4 | 104.1 | 94.7 | 83.1 |
| 94 | Polyadenylate-binding protein 1 (Fragment) OS=Mus musculus OX=10090 GN=Pabpc1 PE=1 SV=1 | 132 | 115.7 | 127.9 | 90.4 | 78.4 | 114.1 | 46 | 97.2 | 98.5 | 99.7 |
| 95 | Acyl-CoA-binding domain-containing protein 5 OS=Mus musculus OX=10090 GN=Acbd5 PE=1 SV=1 | 88.6 | 79.6 | 84.9 | 65.1 | 179 | 80.3 | 176.2 | 87.6 | 66.6 | 92 |
| 96 | Inosine-guanosine phosphorylase (Fragment) OS=Mus musculus OX=10090 GN=Pnp PE=1 SV=1 | 144 | 97.9 | 127.9 | 104.7 | 70.8 | 109.5 | 68.7 | 106.3 | 94 | 76.2 |
| 97 | Galectin-1 OS=Mus musculus OX=10090 GN=Lgals1 PE=1 SV=1 | 56 | 69.2 | 86.7 | 81.8 | 271.3 | 141 | 122 | 46.1 | 56.3 | 69.4 |
| 98 | Alanine transaminase (Fragment) OS=Mus musculus OX=10090 GN=Gpt PE=1 SV=1 | 116.8 | 140.7 | 125.9 | 104.1 | 72.3 | 91.2 | 89.6 | 94.3 | 90 | 75 |
| 99 | La-related protein 4 OS=Mus musculus OX=10090 GN=Larp4 PE=1 SV=1 | 97.2 | 88.7 | 93.9 | 75.4 | 120.8 | 104.6 | 138.2 | 96.8 | 80 | 104.4 |
| 100 | Cysteine sulfinic acid decarboxylase OS=Mus musculus OX=10090 GN=Csad PE=1 SV=1 | 53.9 | 224.6 | 113.7 | 65.1 | 105.7 | 141.4 | 93.4 | 68.8 | 65.2 | 68.3 |

| | | | | | | | | | | | |
|---|---|---|---|---|---|---|---|---|---|---|---|
| 101 | Aconitate hydratase, mitochondrial (Fragment) OS=Mus musculus OX=10090 GN=Aco2 PE=1 SV=1 | 76.1 | 78.8 | 80.8 | 77.9 | 121.1 | 74.2 | 155.9 | 125.5 | 88.8 | 120.8 |
| 102 | Cysteine sulfinic acid decarboxylase (Fragment) OS=Mus musculus OX=10090 GN=Csad PE=1 SV=1 | | 272.7 | 119 | 59.6 | 119.7 | 155.6 | 94.5 | 60.3 | 55.2 | 63.5 |
| 103 | Methyltransferase-like 7A1 OS=Mus musculus OX=10090 GN=Mettl7a1 PE=1 SV=1 | 102.6 | 82.5 | 78.9 | 114.9 | 75.6 | 113.8 | 68.3 | 150 | 129.7 | 83.6 |
| 104 | Myoglobin OS=Mus musculus OX=10090 GN=Mb PE=1 SV=1 | 14.3 | 33.5 | 15.3 | 15.7 | 392.5 | 12.7 | 448.4 | 15.7 | 13.3 | 38.5 |
| 105 | E3 ubiquitin-protein ligase RBX1 OS=Mus musculus OX=10090 GN=Rbx1 PE=1 SV=1 | 86.4 | 78.8 | 76.2 | 66.9 | 142.4 | 86.1 | 231.9 | 80.9 | 70.2 | 80.2 |
| 106 | Carbonyl reductase [NADPH] 1 OS=Mus musculus OX=10090 GN=Cbr1 PE=1 SV=1 | 104.6 | 93.4 | 82.2 | 112.9 | 107.5 | 84.3 | 50.9 | 135.5 | 119.5 | 109.2 |
| 107 | Omega-amidase NIT2 OS=Mus musculus OX=10090 GN=Nit2 PE=1 SV=1 | 141.8 | 125.3 | 132.6 | 109.6 | 65.5 | 88.3 | 90.2 | 69.4 | 92.7 | 84.9 |
| 108 | ATP synthase peripheral stalk subunit OSCP (Fragment) OS=Mus musculus OX=10090 GN=Atp5o PE=1 SV=1 | 97 | 85.3 | 104.3 | 111.5 | 115.6 | 91.5 | 109.4 | 103.7 | 97.1 | 84.7 |
| 109 | High mobility group protein HMG-I/HMG-Y OS=Mus musculus OX=10090 GN=Hmga1 PE=1 SV=1 | 36.2 | 79.4 | 72.6 | 51.6 | 262.6 | 157.6 | 63.1 | 82 | 53 | 142 |
| 110 | 60S ribosomal protein L10a (Fragment) OS=Mus musculus OX=10090 GN=Rpl10a PE=1 SV=1 | 116.8 | 89.5 | 127.7 | 79.8 | 98.4 | 139.6 | 97.6 | 94.1 | 87.8 | 68.7 |
| 111 | pre-mRNA 3' end-processing protein WDR33 OS=Mus musculus OX=10090 GN=Wdr33 PE=1 SV=1 | 122.7 | 100.7 | 109 | 104.8 | 58.1 | 109.4 | 50.2 | 112.6 | 120.7 | 111.7 |
| 112 | 3-hydroxyanthranilate 3,4-dioxygenase OS=Mus musculus OX=10090 GN=Haao PE=1 SV=1 | 123.7 | 113.7 | 120.9 | 94.4 | 105.1 | 104.8 | 100.8 | 79.2 | 82.7 | 74.9 |
| 113 | Cysteine dioxygenase OS=Mus musculus OX=10090 GN=Cdo1 PE=1 SV=1 | 115.3 | 149.2 | 93.9 | 97.7 | 79.1 | 107.6 | 79.7 | 97.7 | 81.1 | 98.6 |
| 114 | Enoyl-CoA delta isomerase 1, mitochondrial OS=Mus musculus OX=10090 GN=Eci1 PE=1 SV=1 | 102.1 | 105.7 | 103.7 | 107.5 | 114.8 | 107.8 | 92 | 96.4 | 91.9 | 78.1 |
| 115 | Asparaginyl-tRNA synthetase OS=Mus musculus OX=10090 GN=Nars PE=1 SV=1 | 94.9 | 101.4 | 85.4 | 90.6 | 146.1 | 103.1 | 97.3 | 100.8 | 72.7 | 107.6 |
| 116 | 3-ketoacyl-CoA thiolase, mitochondrial (Fragment) OS=Mus musculus OX=10090 GN=Acaa2 PE=1 SV=1 | 123 | 110.4 | 108.6 | 100.8 | 98.5 | 113.4 | 81.7 | 89.3 | 93.4 | 80.9 |
| 117 | Glycerol-3-phosphate acyltransferase 1, mitochondrial OS=Mus musculus OX=10090 GN=Gpam PE=1 SV=1 | 169.6 | 118.3 | 140 | 102.5 | 64.4 | 118.9 | 31.9 | 81.4 | 101.2 | 71.8 |
| 118 | Glutathione S-transferase omega OS=Mus musculus OX=10090 GN=Gsto1 PE=1 SV=1 | 118.7 | 116.7 | 105.8 | 116.6 | 67.7 | 104.4 | 69.7 | 119.3 | 87.1 | 93.9 |
| 119 | Carnitine O-palmitoyltransferase (Fragment) OS=Mus musculus OX=10090 GN=Cpt1a PE=1 SV=1 | 122 | 107.5 | 120.5 | 113.3 | 63.1 | 111.8 | 57.5 | 95.1 | 114.5 | 94.6 |
| 120 | ADP-ribosylation factor-like protein 3 OS=Mus musculus OX=10090 GN=Arl3 PE=1 SV=1 | 111.3 | 110.7 | 102.1 | 101.7 | 102.5 | 108.5 | 85.7 | 96.2 | 100 | 81.2 |
| 121 | GST class-pi OS=Mus musculus OX=10090 GN=Gstp1 PE=1 SV=1 | 75.6 | 75 | 93 | 87.7 | 175 | 118.1 | 74.2 | 111.9 | 89.1 | 100.5 |
| 122 | HECT-type E3 ubiquitin transferase OS=Mus musculus OX=10090 GN=Nedd4l PE=1 SV=1 | 108.9 | 107 | 104.1 | 95.1 | 79.1 | 95.6 | 91.5 | 107.8 | 94.7 | 116.2 |
| 123 | Bifunctional 3'-phosphoadenosine 5'-phosphosulfate synthase 2 OS=Mus musculus OX=10090 GN=Papss2 PE=1 SV=1 | 118.6 | 104.3 | 121.5 | 107.1 | 75.8 | 112.8 | 70 | 87.4 | 110.9 | 91.6 |
| 124 | Cystathionine beta-synthase (Fragment) OS=Mus musculus OX=10090 GN=Cbs PE=1 SV=1 | 123.5 | 116.9 | 114.4 | 102 | 82.8 | 117.8 | 77.2 | 92.6 | 88.7 | 84.2 |
| 125 | Dipeptidyl peptidase 3 OS=Mus musculus OX=10090 GN=Dpp3 PE=1 SV=1 | 124.8 | 117 | 114 | 93.2 | 103.5 | 117.2 | 73 | 92.2 | 82.7 | 82.3 |
| 126 | Acyl-CoA (8-3)-desaturase OS=Mus musculus OX=10090 GN=Fads1 PE=1 SV=1 | 150.3 | 111.2 | 123.4 | 100.8 | 93.2 | 118.7 | 68.2 | 83.5 | 88.2 | 62.5 |
| 127 | Lactoylglutathione lyase OS=Mus musculus OX=10090 GN=Glo1 PE=1 SV=1 | 144 | 154.8 | 119.3 | 73.3 | 111.3 | 109.4 | 100.1 | 79 | 59.7 | 49.2 |
| 128 | Guanine nucleotide-binding protein subunit alpha-14 OS=Mus musculus OX=10090 GN=Gna14 PE=1 SV=1 | 103.1 | 93.7 | 99.2 | 103.6 | 73.3 | 91.9 | 64.2 | 154.5 | 112 | 104.5 |
| 129 | Asparagine--tRNA ligase OS=Mus musculus OX=10090 GN=Nars PE=1 SV=1 | 94.9 | 101.4 | 85.4 | 90.6 | 146.1 | 103.1 | 97.3 | 100.8 | 72.7 | 107.6 |
| 130 | Coiled-coil domain-containing protein 30 (Fragment) OS=Mus musculus OX=10090 GN=Ccdc30 PE=1 SV=1 | 12.5 | 36.7 | 11.3 | 14.9 | 288.1 | 10.3 | 562.8 | 17.2 | 14.9 | 31.3 |
| 131 | Myosin-binding protein C, slow-type OS=Mus musculus OX=10090 GN=Mybpc1 PE=1 SV=1 | 32.6 | 48.2 | 39.1 | 44.2 | 239 | 32.1 | 446.2 | 35.9 | 31.5 | 51.3 |
| 132 | Integrin alpha-6 OS=Mus musculus OX=10090 GN=Itga6 PE=1 SV=1 | | 82.7 | 105 | 83.2 | 145.1 | 99.9 | 200.5 | 101.1 | 93.6 | 88.8 |
| 133 | Oxysterol-binding protein (Fragment) OS=Mus musculus OX=10090 GN=Osbpl8 PE=1 SV=1 | | 167 | | 126 | | 151.6 | 137.9 | 157.4 | 155.7 | 104.3 |
| 134 | Phosphatidylinositol-3-phosphatase SAC1 OS=Mus musculus OX=10090 GN=Sacm1l PE=1 SV=1 | 127.7 | 110.6 | 116 | 97.2 | 77.6 | 114.4 | 93.7 | 94.1 | 96.5 | 72.2 |
| 135 | L-lactate dehydrogenase OS=Mus musculus OX=10090 GN=Ldhb PE=1 SV=1 | 30.8 | 66.7 | 30.9 | 46.2 | 282.2 | 28.5 | 396.1 | 31.6 | 29.6 | 57.4 |

| | | | | | | | | | | | |
|---|---|---|---|---|---|---|---|---|---|---|---|
| 136 | Leucine-rich repeat and calponin homology domain-containing protein 1 OS=Mus musculus OX=10090 GN=Lrch1 PE=1 SV=1 | 83.9 | 77.3 | 94.1 | 89.4 | 112.8 | 112.9 | 125.2 | 106.9 | 86 | 111.4 |
| 137 | 60 kDa lysophospholipase OS=Mus musculus OX=10090 GN=Aspg PE=1 SV=1 | 110.1 | 103.1 | 120.5 | 94.2 | 96.8 | 113.9 | 89.4 | 101.8 | 83.5 | 86.7 |
| 138 | Gephyrin OS=Mus musculus OX=10090 GN=Gphn PE=1 SV=1 | | 161.8 | | 138.1 | | 186.4 | 128.7 | 131.7 | 126.7 | 126.5 |
| 139 | Keratin, type I cytoskeletal 10 OS=Mus musculus OX=10090 GN=Krt10 PE=1 SV=1 | 84.2 | 61 | 60.4 | 120.9 | 182 | 103.5 | 104.5 | 130.4 | 54.7 | 98.4 |
| 140 | Maillard deglycase (Fragment) OS=Mus musculus OX=10090 GN=Park7 PE=1 SV=8 | 108.8 | 128.2 | 101.7 | 101.3 | 98.5 | 99.7 | 63.8 | 103.5 | 94.1 | 100.4 |
| 141 | Oxysterol-binding protein-related protein 9 OS=Mus musculus OX=10090 GN=Osbpl9 PE=1 SV=1 | 93.2 | 93.5 | 119.7 | 81.7 | 88.7 | 129.8 | 62.7 | 124.4 | 110.3 | 96.1 |
| 142 | Cytochrome P450 4A10 OS=Mus musculus OX=10090 GN=Cyp4a10 PE=1 SV=1 | 94.5 | 85.7 | 108.8 | 147.6 | 30.3 | 57.8 | 21.7 | 209.9 | 181.1 | 62.6 |
| 143 | Obscurin OS=Mus musculus OX=10090 GN=Obscn PE=1 SV=3 | 22.5 | 49.3 | 23.8 | 25.8 | 251.4 | 26.2 | 510.1 | 25.4 | 15.2 | 50.2 |
| 144 | CTP:phosphoethanolamine cytidylyltransferase OS=Mus musculus OX=10090 GN=Pcyt2 PE=1 SV=1 | 150.7 | 119 | 126.8 | 93.9 | 97 | 109 | 65.5 | 88.9 | 78.5 | 70.7 |
| 145 | L-xylulose reductase OS=Mus musculus OX=10090 GN=Dcxr PE=1 SV=1 | 100.7 | 101.5 | 130.7 | 134.1 | 86.2 | 101.4 | 67.4 | 99.7 | 97.6 | 80.7 |
| 146 | Dolichyl-diphosphooligosaccharide--protein glycosyltransferase subunit 2 OS=Mus musculus OX=10090 GN=Rpn2 PE=1 SV=1 | 91.6 | 94.4 | 97.7 | 115.7 | 67.1 | 132.3 | 86.6 | 82.7 | 116.1 | 115.7 |
| 147 | Eukaryotic translation initiation factor 3 subunit I (Fragment) OS=Mus musculus OX=10090 GN=Eif3i PE=1 SV=1 | 112 | 111.9 | 107.2 | 91.1 | 94.2 | 116 | 80.4 | 95.4 | 90.5 | 101.2 |
| 148 | Mitochondrial carrier homolog 2 OS=Mus musculus OX=10090 GN=Mtch2 PE=1 SV=1 | 132.9 | 106.6 | 121 | 112.3 | 51 | 93.1 | 73.9 | 98.6 | 118 | 92.6 |
| 149 | Pirin (Fragment) OS=Mus musculus OX=10090 GN=Pir PE=1 SV=1 | 73.4 | 71.2 | 73.2 | 130.2 | 46.2 | 75.3 | 67.5 | 151.4 | 146.1 | 165.4 |
| 150 | Alpha-tocopherol transfer protein OS=Mus musculus OX=10090 GN=Ttpa PE=1 SV=2 | 116.6 | 103.4 | 124.1 | 105.3 | 68.1 | 108.7 | 70.4 | 108.8 | 102.5 | 92.2 |
| 151 | Alpha-MPP OS=Mus musculus OX=10090 GN=Pmpca PE=1 SV=1 | 115.1 | 105.4 | 113.2 | 91.2 | 101.5 | 100 | 85.7 | 99.9 | 84.4 | 103.7 |
| 152 | ATP synthase, H+-transporting, mitochondrial F1 complex, gamma polypeptide 1 (Fragment) OS=Mus musculus OX=10090 GN=Atp5c1 PE=1 SV=1 | 43.3 | 83.9 | 72.8 | 138.9 | 77.2 | 140.1 | 121.8 | 132.8 | 87.9 | 101.4 |
| 153 | Catalase OS=Mus musculus OX=10090 GN=Cat PE=1 SV=1 | 120.7 | 123.7 | 117.4 | 91.1 | 92.3 | 116.6 | 73.1 | 95.6 | 100.2 | 69.4 |
| 154 | Aspartyl/asparaginyl beta-hydroxylase OS=Mus musculus OX=10090 GN=Asph PE=1 SV=1 | 125.4 | 83.1 | 132.9 | 101 | 98.2 | 148.2 | 62.1 | 97.2 | 82.3 | 69.8 |
| 155 | DnaJ homolog subfamily C member 25 OS=Mus musculus OX=10090 GN=Dnajc25 PE=1 SV=1 | | 147.7 | 158.3 | 93.2 | 96.1 | 132.9 | 87.9 | 105.3 | 99 | 79.6 |
| 156 | F-actin-capping protein subunit beta OS=Mus musculus OX=10090 GN=Capzb PE=1 SV=1 | 115.9 | 113 | 117.3 | 98.9 | 96.4 | 121.9 | 76.2 | 89.3 | 86.7 | 84.4 |
| 157 | E3 ubiquitin-protein ligase UBR4 OS=Mus musculus OX=10090 GN=Ubr4 PE=1 SV=1 | 121.4 | 108.6 | 127.1 | 94.5 | 83.2 | 112.9 | 85.4 | 87.2 | 97.1 | 82.6 |
| 158 | Complex I-B17 OS=Mus musculus OX=10090 GN=Ndufb6 PE=1 SV=1 | 92.7 | 80 | 87.9 | 109.8 | 93.9 | 95.2 | 101.5 | 131.5 | 101.1 | 106.4 |
| 159 | Nebulin OS=Mus musculus OX=10090 GN=Neb PE=1 SV=1 | 52.4 | 67 | 54.5 | 49 | 206.6 | 54.1 | 358.4 | 47.8 | 45.5 | 64.6 |
| 160 | COP9 signalosome complex subunit 2 OS=Mus musculus OX=10090 GN=Cops2 PE=1 SV=1 | 111.2 | 96.7 | 110.5 | 101.5 | 83.4 | 108.2 | 89.2 | 96.9 | 97.6 | 104.9 |
| 161 | Propionate--CoA ligase OS=Mus musculus OX=10090 GN=Acss2 PE=1 SV=1 | 159.8 | 127.5 | 136 | 81.4 | 108.7 | 103.3 | 102.6 | 68.2 | 56.9 | 55.6 |
| 162 | Glutathione synthase (Fragment) OS=Mus musculus OX=10090 GN=Gss PE=1 SV=1 | 98.3 | 102.6 | 100.2 | 97.2 | 118.8 | 86.2 | 98.8 | 103.5 | 89.9 | 104.4 |
| 163 | Glycerol-3-phosphate dehydrogenase OS=Mus musculus OX=10090 GN=Gpd2 PE=1 SV=1 | 112.8 | 109.9 | 121.5 | 96.1 | 89.9 | 109.2 | 95.3 | 83.1 | 87.1 | 94.9 |
| 164 | Kynureninase OS=Mus musculus OX=10090 GN=Kynu PE=1 SV=1 | 13.3 | 110.3 | 108.9 | 114.5 | 123.9 | 118.4 | 127.4 | 99.6 | 92 | 91.7 |
| 165 | Agmatinase, mitochondrial OS=Mus musculus OX=10090 GN=Agmat PE=1 SV=1 | 88.7 | 79.9 | 95.4 | 130.5 | 59.5 | 96.4 | 65.9 | 99.7 | 146 | 137.9 |
| 166 | Probable 2-oxoglutarate dehydrogenase E1 component DHKTD1, mitochondrial OS=Mus musculus OX=10090 GN=Dhtkd1 PE=1 SV=1 | 77.6 | 74.5 | 72.3 | 105.1 | 77.4 | 99 | 95.8 | 149.3 | 113.4 | 135.6 |
| 167 | Kelch-like protein 41 OS=Mus musculus OX=10090 GN=Klhl41 PE=1 SV=1 | 55.2 | 108.1 | 58 | 69 | 186.9 | 63.3 | 292.8 | 63.7 | 33.1 | 70 |
| 168 | Parathion hydrolase-related protein (Fragment) OS=Mus musculus OX=10090 GN=Pter PE=1 SV=2 | 156.9 | 146.2 | 145.8 | 76.7 | 114.1 | 142.7 | 62.3 | 50.1 | 47 | 58.1 |
| 169 | Major urinary protein 11 OS=Mus musculus OX=10090 GN=Mup9 PE=3 SV=1 | 167.6 | 194.2 | 143.4 | 69.2 | 56.5 | 76.2 | 49.6 | 82.8 | 82.3 | 78.1 |

| | | | | | | | | | | | |
|---|---|---|---|---|---|---|---|---|---|---|---|
| 170 | Phosphoglucomutase-1 OS=Mus musculus OX=10090 GN=Pgm1 PE=1 SV=1 | 128.4 | 112.6 | 112.9 | 106.3 | 98.1 | 110.9 | 86.8 | 94.6 | 73.6 | 75.8 |
| 171 | Damage-control phosphatase ARMT1 OS=Mus musculus OX=10090 GN=Armt1 PE=1 SV=1 | 126.8 | 98.4 | 118.6 | 111.8 | 62.6 | 75.1 | 80 | 123.7 | 108.7 | 94.4 |
| 172 | Inter alpha-trypsin inhibitor, heavy chain 4 OS=Mus musculus OX=10090 GN=Itih4 PE=1 SV=2 | 105.6 | 105.1 | 127.5 | 86.9 | 96.7 | 112.6 | 86.2 | 104.2 | 76.6 | 98.5 |
| 173 | Fructose-bisphosphate aldolase OS=Mus musculus OX=10090 GN=Aldoart2 PE=1 SV=1 | 14.3 | 21.8 | 12.7 | 44.8 | 85.8 | 40.6 | 660.5 | 19.8 | 23.5 | 76.2 |
| 174 | Hemoglobin X, alpha-like embryonic chain in Hba complex (Fragment) OS=Mus musculus OX=10090 GN=Hba-x PE=1 SV=1 | 118.2 | 102.8 | 197.1 | 78.2 | 66 | 83.2 | 46.9 | 102.7 | 113.8 | 91.2 |
| 175 | Beta-globin OS=Mus musculus OX=10090 GN=Hbb-bs PE=1 SV=1 | 82.8 | 120 | 136.7 | 103.6 | 47.4 | 102.4 | 68.6 | 111.6 | 118.4 | 108.5 |
| 176 | Adenosine 5'-monophosphoramidase HINT1 OS=Mus musculus OX=10090 GN=Hint1 PE=1 SV=1 | 106.1 | 124 | 103.8 | 95.3 | 86.8 | 101.4 | 65.9 | 104.8 | 91.3 | 120.7 |
| 177 | CDGSH iron-sulfur domain-containing protein 3, mitochondrial OS=Mus musculus OX=10090 GN=Cisd3 PE=1 SV=1 | 91.8 | 81.4 | 95 | 123.5 | 89.7 | 97.3 | 65.7 | 135.9 | 111.2 | 108.5 |
| 178 | Myosin, heavy polypeptide 13, skeletal muscle OS=Mus musculus OX=10090 GN=Myh13 PE=1 SV=1 | 13.1 | 39.1 | 12.3 | 23.4 | 249.4 | 12.7 | 589.8 | 17.8 | | 42.5 |
| 179 | 60S ribosomal protein L26 (Fragment) OS=Mus musculus OX=10090 GN=Rpl26 PE=1 SV=1 | 109 | 122.5 | 120.8 | 92 | 101.1 | 124 | 64.7 | 106.4 | 87.4 | 72.1 |
| 180 | Phosphoribosyl pyrophosphate synthase-associated protein 1 OS=Mus musculus OX=10090 GN=Prpsap1 PE=1 SV=1 | 142.4 | 115.2 | 109.2 | 111.4 | 78.6 | 116.2 | 48.3 | 106 | 88 | 84.8 |
| 181 | Cytosolic malate dehydrogenase (Fragment) OS=Mus musculus OX=10090 GN=Mdh1 PE=1 SV=8 | 112.5 | 107.1 | 122.7 | 101.9 | 99.5 | 77.8 | 114.1 | 85.8 | 92.1 | 86.5 |
| 182 | Apoptosis-inducing factor 1, mitochondrial OS=Mus musculus OX=10090 GN=Aifm1 PE=1 SV=1 | 94.2 | 98.5 | 97.6 | 113.9 | 82.2 | 105.6 | 74.3 | 120.4 | 118.7 | 94.7 |
| 183 | Aldehyde dehydrogenase OS=Mus musculus OX=10090 GN=Aldh3a2 PE=1 SV=1 | 126.7 | 107.2 | 138.7 | 121.2 | 68.2 | 103.5 | 66.2 | 88 | 116.4 | 63.9 |
| 184 | Hydroxysteroid dehydrogenase-like protein 2 OS=Mus musculus OX=10090 GN=Hsdl2 PE=1 SV=1 | 102.6 | 95.1 | 97 | 107.3 | 81 | 97.2 | 69.8 | 117.1 | 116.4 | 116.4 |
| 185 | Cordon-bleu protein-like 1 OS=Mus musculus OX=10090 GN=Cobll1 PE=1 SV=1 | 76.4 | 93.8 | 84.1 | 96.6 | 84.6 | 94.8 | 74.9 | 106.8 | 103.3 | 184.8 |
| 186 | Palmitoyl-protein hydrolase 1 OS=Mus musculus OX=10090 GN=Ppt1 PE=1 SV=1 | 84.2 | 103.4 | 85.7 | 75 | 146.2 | 96.4 | 131.5 | 97.3 | 83 | 97.3 |
| 187 | Poly(rC)-binding protein 2 OS=Mus musculus OX=10090 GN=Pcbp2 PE=1 SV=1 | 101.4 | 98.1 | 99.5 | 87 | 114.2 | 105 | 114.4 | 115.5 | 81.4 | 83.6 |
| 188 | Myo18a protein OS=Mus musculus OX=10090 GN=Myo18a PE=1 SV=1 | 128.4 | 118 | 111.6 | 100.2 | 81.5 | 121.3 | 69.3 | 89.4 | 94.3 | 86.1 |
| 189 | CDGSH iron sulfur domain 3 OS=Mus musculus OX=10090 GN=Cisd3 PE=1 SV=1 | 91.8 | 81.4 | 95 | 123.5 | 89.7 | 97.3 | 65.7 | 135.9 | 111.2 | 108.5 |
| 190 | Filamin, alpha OS=Mus musculus OX=10090 GN=Flna PE=1 SV=1 | | 168.9 | | 118.7 | | 159.8 | 164.2 | 123.4 | 124.5 | 140.6 |
| 191 | Peroxin-7 OS=Mus musculus OX=10090 GN=Pex7 PE=1 SV=1 | 122.9 | 109.4 | 130.2 | 95 | 99.5 | 111.9 | 103.3 | 87.7 | 80.7 | 59.3 |
| 192 | C3/C5 convertase OS=Mus musculus OX=10090 GN=Gm20547 PE=4 SV=1 | 122.1 | 119.9 | 137.9 | 86.7 | 106.5 | 108.5 | 87.4 | 76.7 | 78.1 | 76.2 |
| 193 | Heterogeneous nuclear ribonucleoprotein D-like OS=Mus musculus OX=10090 GN=Hnrnpdl PE=1 SV=1 | 85.6 | 85.8 | 85.1 | 74.1 | 133.5 | 101.1 | 45.6 | 114.6 | 86.5 | 188.2 |
| 194 | Acyl-CoA dehydrogenase family member 11 OS=Mus musculus OX=10090 GN=Acad11 PE=1 SV=1 | 78.3 | 88.3 | 88.2 | 101.8 | 78.3 | 102.5 | 111.8 | 103.8 | 122.9 | 124.1 |
| 195 | Adenylyl cyclase-associated protein OS=Mus musculus OX=10090 GN=Cap2 PE=1 SV=1 | 113.9 | 110.3 | 109.2 | 95.5 | 106.9 | 125.9 | 88.4 | 86 | 86.4 | 77.5 |
| 196 | Isobutyryl-CoA dehydrogenase, mitochondrial OS=Mus musculus OX=10090 GN=Acad8 PE=1 SV=1 | 116.7 | 83.6 | 112.6 | 109.6 | 67.1 | 94.1 | 83.2 | 112.3 | 115.6 | 105.2 |
| 197 | Kininogen-1 OS=Mus musculus OX=10090 GN=Kng1 PE=1 SV=1 | 70 | 82.2 | 108.5 | 83.5 | 114.5 | 77.3 | 101.2 | 116.4 | 105.5 | 140.7 |
| 198 | 40S ribosomal protein S19 (Fragment) OS=Mus musculus OX=10090 GN=Rps19 PE=1 SV=8 | 146.5 | 106.4 | 122.1 | 95.1 | 92.7 | 121.7 | 59.3 | 95.5 | 85.3 | 75.6 |
| 199 | Complex I-PDSW (Fragment) OS=Mus musculus OX=10090 GN=Ndufb10 PE=1 SV=1 | 87.1 | 92 | 90.9 | 79.8 | 157.8 | 98.4 | 168.7 | 64.8 | 88 | 72.6 |
| 200 | 40S ribosomal protein S9 (Fragment) OS=Mus musculus OX=10090 GN=Rps9 PE=1 SV=8 | 116.5 | 128.6 | 126 | 85.5 | 81 | 106.2 | 89 | 115.1 | 83 | 68.9 |
| 201 | 40S ribosomal protein S2 (Fragment) OS=Mus musculus OX=10090 GN=Rps2 PE=1 SV=1 | 127.2 | 106.3 | 124.7 | 90.6 | 84.7 | 118.9 | 74.2 | 107.4 | 83.8 | 82.2 |
| 202 | Glutathione S-transferase Mu 6 OS=Mus musculus OX=10090 GN=Gstm6 PE=1 SV=1 | 92.3 | 87 | 104 | 132.3 | 76.3 | 122.1 | 78.8 | 92 | 110.1 | 105 |
| 203 | Mitochondrial pyruvate carrier OS=Mus musculus OX=10090 GN=Mpc1 PE=1 SV=1 | 132.5 | 107.2 | 104.4 | 99.5 | 89.5 | 97.5 | 101.9 | 82.2 | 93.8 | 91.6 |
| 204 | Arp2/3 complex 34 kDa subunit OS=Mus musculus OX=10090 GN=Arpc2 PE=1 SV=1 | 115.1 | 92.4 | 101.3 | 86.3 | 85.9 | 113.2 | 61.1 | 88.3 | 83.4 | 172.9 |

| | | | | | | | | | | | |
|---|---|---|---|---|---|---|---|---|---|---|---|
| 205 | Complex I-49kD OS=Mus musculus OX=10090 GN=Ndufs2 PE=1 SV=1 | 95 | 84.8 | 94.8 | 109.4 | 86.1 | 88.1 | 129.5 | 108.3 | 105.4 | 98.7 |
| 206 | DET1- and DDB1-associated protein 1 OS=Mus musculus OX=10090 GN=Dda1 PE=1 SV=1 | 89.8 | 96.5 | 94.3 | 70.3 | 103.9 | 95.6 | 114.8 | 108.5 | 98.5 | 127.7 |
| 207 | Phosphoglycerate mutase (Fragment) OS=Mus musculus OX=10090 GN=Bpgm PE=1 SV=1 | 106.1 | 98.1 | 110.3 | 87.6 | 92.5 | 109.4 | 102.9 | 110.2 | 86.5 | 96.3 |
| 208 | 40S ribosomal protein S5 (Fragment) OS=Mus musculus OX=10090 GN=Rps5 PE=1 SV=1 | 98.3 | 92.9 | 86.3 | 83.1 | 96.1 | 110.4 | 66.6 | 96.2 | 82.2 | 187.7 |
| 209 | Heat shock protein beta-1 OS=Mus musculus OX=10090 GN=Hspb1 PE=1 SV=1 | 56.7 | 77.7 | 49.7 | 55.3 | 225 | 56 | 262.2 | 67.6 | 56.4 | 93.6 |
| 210 | Arachidonate--CoA ligase OS=Mus musculus OX=10090 GN=Acsl1 PE=1 SV=1 | 123 | 115.2 | 111.2 | 97.3 | 96.7 | 114.5 | 119.1 | 83.2 | 81.6 | 58.2 |
| 211 | MICOS complex subunit Mic19 (Fragment) OS=Mus musculus OX=10090 GN=Chchd3 PE=1 SV=1 | 77 | 79.6 | 82.4 | 92.8 | 125.3 | 85.7 | 172.7 | 101.5 | 91.1 | 91.9 |
| 212 | Acyl-protein thioesterase 1 OS=Mus musculus OX=10090 GN=Lypla1 PE=1 SV=1 | 108.7 | 107.7 | 107.6 | 117.3 | 75.6 | 105.9 | 71 | 114.3 | 106.8 | 85 |
| 213 | Glutaminyl-tRNA synthetase OS=Mus musculus OX=10090 GN=Qars PE=1 SV=2 | 138.4 | 105.4 | 123.5 | 93.4 | 87.4 | 129.7 | 63.9 | 97.5 | 87.5 | 73.3 |
| 214 | Corrinoid adenosyltransferase OS=Mus musculus OX=10090 GN=Mmab PE=1 SV=1 | 92.2 | 105.7 | 84.6 | 92.9 | 115.2 | 92.1 | 95.1 | 111 | 94.8 | 116.3 |
| 215 | Phosphatidylethanolamine-binding protein 1 OS=Mus musculus OX=10090 GN=Pebp1 PE=1 SV=1 | 151.6 | 113.4 | 120.9 | 100 | 115.4 | 82.1 | 71.5 | 101.6 | 70.4 | 73.1 |
| 216 | L-serine ammonia-lyase (Fragment) OS=Mus musculus OX=10090 GN=Sds PE=1 SV=1 | 98.2 | 95.6 | 93.3 | 99.3 | 88 | 109.7 | 78.8 | 111.4 | 104.2 | 121.2 |
| 217 | Aspartate dehydrogenase domain-containing protein (Fragment) OS=Mus musculus OX=10090 GN=Aspdh PE=1 SV=8 | 141 | 112.6 | 123.6 | 94.9 | 66.8 | 115.9 | 82.8 | 83.2 | 92.2 | 87.1 |
| 218 | Acyl-Coenzyme A dehydrogenase family, member 12 OS=Mus musculus OX=10090 GN=Acad12 PE=1 SV=1 | 94.2 | 91 | 90.4 | 90.7 | 124.2 | 102.8 | 134 | 106.4 | 86.9 | 79.4 |
| 219 | Actin-related protein 2/3 complex subunit 3 OS=Mus musculus OX=10090 GN=Arpc3 PE=1 SV=1 | 112.6 | 99.5 | 103.8 | 91.3 | 97.2 | 129.7 | 80 | 95.1 | 91.1 | 99.6 |
| 220 | CDGSH iron-sulfur domain-containing protein 2 OS=Mus musculus OX=10090 GN=Cisd2 PE=1 SV=1 | 128.8 | 109.2 | 122.2 | 89.5 | 91.5 | 128.6 | 92.3 | 83 | 83.4 | 71.6 |
| 221 | Ceramide synthase 2 (Fragment) OS=Mus musculus OX=10090 GN=Cers2 PE=1 SV=1 | 106.4 | 110.7 | 110.9 | 94.3 | 123.3 | 107.2 | 87.4 | 92.3 | 85.9 | 81.7 |
| 222 | Complex I-B9 OS=Mus musculus OX=10090 GN=Ndufa3 PE=1 SV=1 | 92.8 | 78.9 | 100.5 | 122.8 | 82.6 | 88 | 106.4 | 133.8 | 118.2 | 76 |
| 223 | Formyltetrahydrofolate dehydrogenase OS=Mus musculus OX=10090 GN=Aldh1l2 PE=1 SV=1 | 104.8 | 93.9 | 92.2 | 137 | 75.2 | 111 | 65.3 | 132.3 | 105 | 83.4 |
| 224 | 3-oxoacyl-[acyl-carrier-protein] reductase (Fragment) OS=Mus musculus OX=10090 GN=Cbr4 PE=1 SV=1 | 114.1 | 99.8 | 93.5 | 107.9 | 78.2 | 95.8 | 62.9 | 124.9 | 101.9 | 121 |
| 225 | Mitogen-activated protein kinase 3 OS=Mus musculus OX=10090 GN=Mapk3 PE=1 SV=1 | 108.2 | 113.7 | 115.6 | 79.5 | 80.3 | 98.6 | 96 | 113.8 | 85.7 | 108.8 |
| 226 | 40S ribosomal protein S19 OS=Mus musculus OX=10090 GN=Rps19 PE=1 SV=1 | 146.5 | 106.4 | 122.1 | 95.1 | 92.7 | 121.7 | 59.3 | 95.5 | 85.3 | 75.6 |
| 227 | Acidic leucine-rich nuclear phosphoprotein 32 family member A OS=Mus musculus OX=10090 GN=Anp32a PE=1 SV=1 | 102.2 | 99.9 | 96.5 | 80.3 | 121.8 | 110.8 | 93.9 | 83.3 | 66.4 | 144.9 |
| 228 | Mitochondrial fission regulator 1-like OS=Mus musculus OX=10090 GN=Mtfr1l PE=1 SV=1 | 105.6 | 101.4 | 94.9 | 97 | 117.4 | 104 | 95.6 | 86.7 | 96.3 | 101.2 |
| 229 | Glyoxylate reductase/hydroxypyruvate reductase OS=Mus musculus OX=10090 GN=Grhpr PE=1 SV=1 | 126 | 112.1 | 121.3 | 85.9 | 95.2 | 143.9 | 53.9 | 98.9 | 89.8 | 73 |
| 230 | Proline dehydrogenase OS=Mus musculus OX=10090 GN=Prodh2 PE=1 SV=1 | 112.4 | 102 | 106 | 119.5 | 64.9 | 107.7 | 67.5 | 116.9 | 114.9 | 88.2 |
| 231 | DCS-1 OS=Mus musculus OX=10090 GN=Dcps PE=1 SV=1 | 136.6 | 128.8 | 121.1 | 87.7 | 73.8 | 103.4 | 68.1 | 88.5 | 88.2 | 103.9 |
| 232 | Choline kinase alpha OS=Mus musculus OX=10090 GN=Chka PE=4 SV=1 | 107.3 | 74.1 | 95.6 | 104.7 | 100.1 | 86.8 | 115.9 | 106.3 | 90.1 | 118.9 |
| 233 | NADH dehydrogenase [ubiquinone] flavoprotein 1, mitochondrial OS=Mus musculus OX=10090 GN=Ndufv1 PE=1 SV=1 | 86.9 | 82.3 | 79.2 | 102.5 | 104.7 | 78.8 | 124.5 | 116 | 103.8 | 121.4 |
| 234 | Coatomer subunit alpha OS=Mus musculus OX=10090 GN=Copa PE=1 SV=1 | 121.7 | 121 | 108.8 | 77.9 | 92.7 | 129.9 | 99.1 | 93.2 | 78.6 | 77.2 |
| 235 | Peroxisomal membrane protein PEX14 OS=Mus musculus OX=10090 GN=Pex14 PE=1 SV=1 | 122.1 | 107.6 | 118.2 | 95.2 | 90.7 | 104 | 73.9 | 94.3 | 99.7 | 94.4 |
| 236 | Peroxisomal trans-2-enoyl-CoA reductase OS=Mus musculus OX=10090 GN=Pecr PE=1 SV=1 | 103.6 | 92.5 | 106.8 | 123.8 | 84.8 | 120.9 | 61.9 | 118 | 108.4 | 79.5 |
| 237 | ENH isoform 1b OS=Mus musculus OX=10090 GN=Pdlim5 PE=1 SV=1 | 129.5 | 117.6 | 121.6 | 88.6 | 98.5 | 111.9 | 61.7 | 94.2 | 78 | 98.4 |
| 238 | ENH isoform 3a OS=Mus musculus OX=10090 GN=Pdlim5 PE=1 SV=1 | | 40.1 | 20.8 | 36.8 | 259.6 | 34.7 | 433.9 | 60 | 35.8 | 78.3 |
| 239 | Delta(14)-sterol reductase TM7SF2 OS=Mus musculus OX=10090 GN=Tm7sf2 PE=1 SV=1 | 222.2 | 138.6 | 134.5 | 81.1 | 80.7 | 78 | 38.6 | 117.7 | 76.8 | 31.8 |

| | | | | | | | | | | | |
|---|---|---|---|---|---|---|---|---|---|---|---|
| 240 | Alpha N-catenin OS=Mus musculus OX=10090 GN=Ctnna2 PE=1 SV=1 | 124.4 | 132.2 | 128.8 | 85.9 | 82.7 | 119.8 | 86 | 80.7 | 85.4 | 74.2 |
| 241 | Dynamin-like 120 kDa protein, mitochondrial OS=Mus musculus OX=10090 GN=Opa1 PE=1 SV=1 | | 116.6 | | 111.9 | | 103.4 | 322.1 | 108.2 | 111 | 126.9 |
| 242 | Collagen alpha-2(I) chain (Fragment) OS=Mus musculus OX=10090 GN=Col1a2 PE=1 SV=1 | 61 | 122.3 | 79.2 | 67.1 | 183.7 | 49.9 | 217.1 | 80.8 | 64 | 74.9 |
| 243 | Nucleolar protein 56 OS=Mus musculus OX=10090 GN=Nop56 PE=1 SV=1 | 131.4 | 106.7 | 122.9 | 94.8 | 64.2 | 120.7 | 72.8 | 102.7 | 90.4 | 93.5 |
| 244 | H3.3 histone A (Fragment) OS=Mus musculus OX=10090 GN=H3f3a PE=3 SV=1 | 83.8 | 132.8 | 109.9 | 63.2 | 179.7 | 60.6 | 71.1 | 90.8 | 57.2 | 150.8 |
| 245 | NudC domain-containing protein 2 OS=Mus musculus OX=10090 GN=Nudcd2 PE=1 SV=1 | | 125 | | 110.3 | | 156.3 | 101.5 | 104.3 | 191.5 | 211.2 |
| 246 | Methionine adenosyltransferase 2 subunit beta OS=Mus musculus OX=10090 GN=Mat2b PE=1 SV=1 | 124.1 | 106.3 | 137.7 | 102.5 | 76.4 | 108.3 | 70.1 | 94.8 | 100.6 | 79.1 |
| 247 | Amylo-1,6-glucosidase, 4-alpha-glucanotransferase OS=Mus musculus OX=10090 GN=Agl PE=1 SV=1 | 155.3 | 129 | 128.6 | 96.2 | 66.8 | 93.4 | 81.9 | 82.9 | 88 | 77.8 |
| 248 | Histone H3 (Fragment) OS=Mus musculus OX=10090 GN=H3f3a PE=3 SV=1 | 82.6 | 133 | 108.5 | 63.1 | 184.7 | 60.8 | 72.4 | 89 | 56.4 | 149.5 |
| 249 | N-acetyltransferase 8 (GCN5-related) family member 1 OS=Mus musculus OX=10090 GN=Nat8f1 PE=1 SV=1 | 139 | 125.8 | 100.4 | 112.4 | 84.3 | 162.9 | 46.1 | 89.2 | 90.5 | 49.4 |
| 250 | Alpha-1,4 glucan phosphorylase OS=Mus musculus OX=10090 GN=Pygm PE=1 SV=1 | 25.3 | 60.7 | 23.2 | 72 | 81.3 | 59.5 | 462.6 | 53.5 | 59.3 | 102.6 |
| 251 | Fibrinogen alpha chain OS=Mus musculus OX=10090 GN=Fga PE=1 SV=1 | 88.7 | 84.6 | 120.4 | 70.8 | 99.2 | 97.6 | 83.3 | 126.7 | 90.9 | 137.8 |
| 252 | eIF-2-alpha kinase activator GCN1 OS=Mus musculus OX=10090 GN=Gcn1 PE=1 SV=1 | 117 | 121.5 | 119.1 | 99 | 85.8 | 124.8 | 64.1 | 97 | 95.7 | 75.9 |
| 253 | Microtubule-actin cross-linking factor 1 OS=Mus musculus OX=10090 GN=Macf1 PE=1 SV=2 | 124.4 | 105.1 | 107 | 88.7 | 83.9 | 107.6 | 100.4 | 94.8 | 96 | 92 |
| 254 | Golgi autoantigen, golgin subfamily b, macrogolgin 1 OS=Mus musculus OX=10090 GN=Golgb1 PE=1 SV=1 | 105.7 | 104.7 | 116.8 | 89.6 | 100.6 | 110.2 | 81.8 | 102.7 | 79.9 | 108 |
| 255 | Isoleucyl-tRNA synthetase OS=Mus musculus OX=10090 GN=Iars2 PE=1 SV=1 | 98.1 | 86.4 | 90.2 | 112.6 | 48.2 | 93.1 | 101.1 | 121 | 129.4 | 119.9 |
| 256 | Coronin OS=Mus musculus OX=10090 GN=Coro1c PE=1 SV=1 | 106.3 | 105.5 | 105.9 | 85.9 | 98.2 | 122.1 | 87.1 | 93 | 94.6 | 101.3 |
| 257 | Dedicator of cytokinesis protein 7 OS=Mus musculus OX=10090 GN=Dock7 PE=1 SV=2 | | 99.3 | 91.2 | 124.7 | 125.6 | 140.7 | 98.2 | 103.4 | 105.5 | 111.5 |
| 258 | Cytochrome P450, family 2, subfamily c, polypeptide 69 OS=Mus musculus OX=10090 GN=Cyp2c69 PE=1 SV=1 | 161.9 | 205.8 | 177.4 | 81.4 | 35.3 | 94.4 | 57.6 | 63.2 | 71.3 | 51.8 |
| 259 | Equilibrative nucleoside transporter 1 OS=Mus musculus OX=10090 GN=Slc29a1 PE=1 SV=1 | 117.7 | 100.7 | 108.4 | 105.2 | 80.5 | 105.9 | 65.8 | 118.5 | 96.9 | 100.4 |
| 260 | Cullin-4A OS=Mus musculus OX=10090 GN=Cul4a PE=1 SV=1 | 116.9 | 111 | 120.9 | 104.2 | 89.4 | 112.4 | 79.5 | 92.1 | 91.6 | 82.1 |
| 261 | Bile acyl-CoA synthetase OS=Mus musculus OX=10090 GN=Slc27a5 PE=1 SV=1 | 119.9 | 104.5 | 102.2 | 98.7 | 62.6 | 116.8 | 95.3 | 114.4 | 89.9 | 95.8 |
| 262 | Cullin-4B OS=Mus musculus OX=10090 GN=Cul4b PE=1 SV=2 | 137.4 | 109.9 | 113.6 | 101.6 | 93.9 | 115.9 | 77.6 | 96.8 | 78.7 | 74.7 |
| 263 | Aminopeptidase B OS=Mus musculus OX=10090 GN=Rnpep PE=1 SV=1 | 107.8 | 104.7 | 120.3 | 91.5 | 94.4 | 121.7 | 94.3 | 97.1 | 83.9 | 84.3 |
| 264 | Ethanolamine-phosphate phospho-lyase OS=Mus musculus OX=10090 GN=Etnppl PE=1 SV=1 | 93.3 | 88.7 | 106.2 | 97.5 | 121.9 | 85.7 | 120.9 | 102.3 | 72.7 | 110.7 |
| 265 | LIM domain-binding protein 3 OS=Mus musculus OX=10090 GN=Ldb3 PE=1 SV=1 | 32.4 | 50 | 30.1 | 24.7 | 335.5 | 25.2 | 406.1 | 25.6 | 13.2 | 57 |
| 266 | Nucleoside diphosphate kinase OS=Mus musculus OX=10090 GN=Gm20390 PE=3 SV=1 | 152.9 | 116 | 107.4 | 95.1 | 101.1 | 100.8 | 67 | 100.3 | 87.2 | 72.1 |
| 267 | Hydroxy-delta-5-steroid dehydrogenase, 3 beta- and steroid delta-isomerase 9 OS=Mus musculus OX=10090 GN=Hsd3b9 PE=1 SV=1 | | 116.2 | 136 | 116.6 | 96.7 | 139.4 | 110.4 | 93.5 | 103.2 | 88.1 |
| 268 | Glyoxalase domain-containing protein 4 OS=Mus musculus OX=10090 GN=Glod4 PE=1 SV=1 | 157.3 | 140.7 | 118.5 | 100.4 | 66.5 | 133.9 | 34.9 | 87.6 | 82.1 | 78.1 |
| 269 | 4-hydroxy-2-oxoglutarate aldolase, mitochondrial OS=Mus musculus OX=10090 GN=Hoga1 PE=1 SV=1 | 92.7 | 86.2 | 84.5 | 126.2 | 61.1 | 87.3 | 50.1 | 142.9 | 131.3 | 137.7 |
| 270 | Hemoglobin, beta adult s chain (Fragment) OS=Mus musculus OX=10090 GN=Hbb-bs PE=1 SV=1 | 75.5 | 122.9 | 135.5 | 105.5 | 46.8 | 103 | 68.9 | 113 | 122.1 | 106.9 |
| 271 | Myosin, heavy chain 15 OS=Mus musculus OX=10090 GN=Myh15 PE=1 SV=1 | 56.9 | 68.5 | 53.6 | 49.6 | 195.9 | 56.8 | 347.1 | 55.1 | 43.4 | 73.1 |
| 272 | Hydroxyacylglutathione hydrolase, mitochondrial (Fragment) OS=Mus musculus OX=10090 GN=Hagh PE=1 SV=1 | 147.7 | 118.6 | 124.6 | 107.7 | 66.2 | 114.2 | 54.9 | 93.8 | 98.9 | 73.4 |
| 273 | C2 domain-containing protein 2 OS=Mus musculus OX=10090 GN=C2cd2 PE=1 SV=1 | 111 | 107.8 | 111.5 | 100.7 | 81.1 | 112.8 | 68.7 | 98.3 | 108.7 | 99.4 |
| 274 | Heat shock protein HSP 90-beta (Fragment) OS=Mus musculus OX=10090 GN=Hsp90ab1 PE=1 SV=1 | 146 | 129 | 141.7 | 84.7 | 87.8 | 134.2 | 70.7 | 78.7 | 75.7 | 51.6 |

| | | | | | | | | | | | |
|---|---|---|---|---|---|---|---|---|---|---|---|
| 275 | Phosphatidylinositol 4-kinase alpha OS=Mus musculus OX=10090 GN=Pi4ka PE=1 SV=2 | 107.7 | 97.3 | 119.6 | 128.4 | 55.9 | 84.4 | 55.3 | 88.8 | 144.7 | 118 |
| 276 | Major vault protein OS=Mus musculus OX=10090 GN=Mvp PE=1 SV=1 | 110.4 | 107.6 | 106.2 | 86.7 | 124.5 | 132.4 | 74.2 | 91.9 | 86.1 | 80 |
| 277 | Amine oxidase OS=Mus musculus OX=10090 GN=Maob PE=1 SV=1 | 115.3 | 111.1 | 128.3 | 124.1 | 78.9 | 96.4 | 81 | 82.7 | 114.4 | 67.9 |
| 278 | Coiled-coil domain-containing protein 58 OS=Mus musculus OX=10090 GN=Ccdc58 PE=1 SV=1 | | 132.2 | 123.8 | 101.9 | 91.1 | 133.4 | 92.9 | 120.6 | 99.6 | 104.5 |
| 279 | Hormone-sensitive lipase OS=Mus musculus OX=10090 GN=Lipe PE=1 SV=1 | 82 | 104 | 92 | 99.7 | 104.8 | 112.6 | 119.3 | 106.6 | 87.9 | 91 |
| 280 | KxDL motif-containing protein 1 (Fragment) OS=Mus musculus OX=10090 GN=Kxd1 PE=1 SV=1 | 85.1 | 101.9 | 93 | 72.1 | 139.6 | 94.7 | 109 | 108.3 | 112.5 | 83.7 |
| 281 | Cullin-3 OS=Mus musculus OX=10090 GN=Cul3 PE=1 SV=1 | 128.2 | 107.9 | 115.6 | 89.3 | 96.5 | 125.5 | 113 | 72.4 | 75.3 | 76.3 |
| 282 | Acetyl-CoA carboxylase 2 OS=Mus musculus OX=10090 GN=Acacb PE=1 SV=1 | 207.5 | 149.4 | 152.4 | 59.6 | 96 | 116.6 | 54.1 | 61.8 | 55.5 | 47.1 |
| 283 | Desmoplakin OS=Mus musculus OX=10090 GN=Dsp PE=1 SV=1 | 124.2 | 109.8 | 112.2 | 107.7 | 92.6 | 112.5 | 88.1 | 92.6 | 79.9 | 80.4 |
| 284 | Methyltransferase-like 26 OS=Mus musculus OX=10090 GN=Mettl26 PE=1 SV=1 | 105.1 | 130.3 | 108.8 | 84.8 | 103.1 | 126.3 | 82.7 | 95.1 | 74.5 | 89.2 |
| 285 | Cytochrome P450 2C44 OS=Mus musculus OX=10090 GN=Cyp2c23 PE=1 SV=1 | 127.5 | 137.8 | 119.7 | 132.4 | 66.8 | 95.7 | 92.4 | 81 | 91.5 | 55.2 |
| 286 | AHNAK nucleoprotein (desmoyokin) OS=Mus musculus OX=10090 GN=Ahnak PE=1 SV=1 | 43.5 | 70.6 | 55.1 | 46.6 | 376.9 | 72.7 | 189.6 | 42.2 | 23.2 | 79.6 |
| 287 | Catenin beta-1 OS=Mus musculus OX=10090 GN=Ctnnb1 PE=1 SV=1 | 107.8 | 108.6 | 99.6 | 86.7 | 108.9 | 116.5 | 68.7 | 96.7 | 89.7 | 116.9 |
| 288 | Phosphodiesterase OS=Mus musculus OX=10090 GN=Pde1c PE=1 SV=1 | 111 | 86.2 | 75.2 | 111.5 | 137 | 97.6 | 105.5 | 89.2 | 104.7 | 82.1 |
| 289 | DnaJ homolog subfamily C member 11 OS=Mus musculus OX=10090 GN=Dnajc11 PE=1 SV=1 | 98.9 | 100.3 | 111.8 | 107.2 | 96 | 95.1 | 97 | 89.8 | 96.5 | 107.5 |
| 290 | Myosin regulatory light chain 2, ventricular/cardiac muscle isoform OS=Mus musculus OX=10090 GN=Myl2 PE=1 SV=1 | 45.9 | 57.4 | 46.2 | 51 | 252 | 61.6 | 308 | 55.5 | 40.8 | 81.6 |
| 291 | NADH dehydrogenase [ubiquinone] 1 beta subcomplex subunit 8, mitochondrial (Fragment) OS=Mus musculus OX=10090 GN=Ndufb8 PE=1 SV=8 | 92.9 | 84.9 | 87.2 | 101.8 | 100.3 | 86.8 | 136.3 | 112.1 | 88.2 | 109.6 |
| 292 | CAD protein OS=Mus musculus OX=10090 GN=Cad PE=1 SV=1 | 136.6 | 155.7 | 106 | 80.4 | 77.4 | 163.3 | 59.3 | 105.8 | 47.2 | 68.4 |
| 293 | D-glutamate cyclase, mitochondrial OS=Mus musculus OX=10090 GN=Dglucy PE=1 SV=1 | 85.7 | 75.5 | 103.5 | 122.5 | 74 | 111.7 | 96 | 94.4 | 117.7 | 118.9 |
| 294 | Atlastin-2 OS=Mus musculus OX=10090 GN=Atl2 PE=1 SV=1 | 86.8 | 104 | 87.9 | 89.6 | 113.8 | 93.1 | 139.8 | 94 | 98.6 | 92.4 |
| 295 | DEAH box protein 9 OS=Mus musculus OX=10090 GN=Dhx9 PE=1 SV=1 | 121.1 | 101.8 | 109.3 | 85 | 82.6 | 119.6 | 62.8 | 103.2 | 101.5 | 113 |
| 296 | Platelet-activating factor acetylhydrolase OS=Mus musculus OX=10090 GN=Pafah2 PE=1 SV=1 | 113 | 96.6 | 116.5 | 101.3 | 72.8 | 90.2 | 88.2 | 125.8 | 114.2 | 81.4 |
| 297 | Collagen alpha-1(XVIII) chain OS=Mus musculus OX=10090 GN=Col18a1 PE=1 SV=1 | 87.4 | 89.7 | 94.3 | 88 | 96.1 | 97.8 | 175.3 | 91.8 | 77.9 | 101.7 |
| 298 | Non-specific serine/threonine protein kinase OS=Mus musculus OX=10090 GN=Obscn PE=1 SV=2 | 22.5 | 49.3 | 23.8 | 25.8 | 251.4 | 26.2 | 510.1 | 25.4 | 15.2 | 50.2 |
| 299 | Lysosomal alpha-glucosidase (Fragment) OS=Mus musculus OX=10090 GN=Gaa PE=1 SV=1 | 110.6 | 99.4 | 96.1 | 93.9 | 103.9 | 84.7 | 114.9 | 99 | 100.7 | 96.7 |
| 300 | Peptidylprolyl isomerase (Fragment) OS=Mus musculus OX=10090 GN=Fkbp4 PE=1 SV=9 | 120.1 | 109.6 | 113.8 | 83.9 | 96.8 | 112 | 100.8 | 86.8 | 86.1 | 90.1 |
| 301 | Glutaminase (Fragment) OS=Mus musculus OX=10090 GN=Gls2 PE=1 SV=1 | 96.7 | 79.5 | 72.3 | 116.1 | 81.1 | 91.2 | 125.8 | 140.1 | 103.1 | 93.9 |
| 302 | Centromere protein V (Fragment) OS=Mus musculus OX=10090 GN=Cenpv PE=1 SV=1 | 113.6 | 118.9 | 99.2 | 94 | 84.9 | 107.7 | 70.8 | 105.2 | 91.6 | 114.1 |
| 303 | Glutaminase liver isoform, mitochondrial (Fragment) OS=Mus musculus OX=10090 GN=Gls2 PE=1 SV=1 | 93.3 | 90 | 77.5 | 91.2 | 66.6 | 126.5 | 137.7 | 121 | 102 | 94 |
| 304 | Epoxide hydrolase 1 (Fragment) OS=Mus musculus OX=10090 GN=Ephx1 PE=1 SV=2 | 88.9 | 75.6 | 129.7 | 151.2 | 61.7 | 78.1 | 44.1 | 125.3 | 165 | 80.3 |
| 305 | 40S ribosomal protein S12 OS=Mus musculus OX=10090 GN=Rps12 PE=1 SV=1 | 54.8 | 63.9 | 56.8 | 43 | 151.1 | 99.4 | 175.2 | 73.5 | 42.8 | 239.4 |
| 306 | Aminopeptidase A (Fragment) OS=Mus musculus OX=10090 GN=Enpep PE=1 SV=1 | 116.4 | 103.2 | 109.4 | 106.2 | 86.9 | 99.1 | 68.4 | 106.5 | 94.6 | 109.3 |
| 307 | Clathrin light chain (Fragment) OS=Mus musculus OX=10090 GN=Cltb PE=1 SV=1 | 97.6 | 89.5 | 95 | 70.9 | 132.2 | 99.2 | 118.4 | 106.4 | 67.6 | 123.2 |
| 308 | Nucleoside-diphosphate kinase (Fragment) OS=Mus musculus OX=10090 GN=Ak2 PE=1 SV=1 | 64.8 | 121.9 | 61.9 | 132.6 | 56.1 | 126.5 | 74.2 | 119.9 | 127.5 | 114.4 |
| 309 | Mesencephalic astrocyte-derived neurotrophic factor (Fragment) OS=Mus musculus OX=10090 GN=Manf PE=1 SV=1 | 103.9 | 104.4 | 102.8 | 85.6 | 120.9 | 131.2 | 70.9 | 96 | 78.4 | 105.9 |

| | | | | | | | | | | | |
|---|---|---|---|---|---|---|---|---|---|---|---|
| 310 | Endoplasmin (Fragment) OS=Mus musculus OX=10090 GN=Hsp90b1 PE=1 SV=1 | 120 | 102.1 | 128.1 | 77.5 | 118.4 | 121.6 | 108.8 | 76.4 | 73.8 | 73.4 |
| 311 | Eukaryotic translation initiation factor 4 gamma 2 (Fragment) OS=Mus musculus OX=10090 GN=Eif4g2 PE=1 SV=1 | 112.9 | 113.2 | 116.9 | 72.2 | 117.3 | 89.7 | 149.9 | 77.8 | 68.9 | 81.2 |
| 312 | FERM, ARHGEF and pleckstrin domain-containing protein 1 OS=Mus musculus OX=10090 GN=Farp1 PE=1 SV=1 | 103.8 | 87.7 | 78.6 | 103.5 | 107.3 | 89.7 | 109.1 | 97.2 | 105.7 | 117.5 |
| 313 | Calcium-transporting ATPase OS=Mus musculus OX=10090 GN=Atp2b2 PE=1 SV=1 | 135.8 | 119.9 | 91.1 | 84 | 67.1 | 119.6 | 112 | 78.2 | 129.6 | 62.8 |
| 314 | Copper homeostasis protein cutC homolog OS=Mus musculus OX=10090 GN=Cutc PE=1 SV=1 | 94.7 | 93.5 | 101.4 | 95.4 | 77 | 90.1 | 86.1 | 121 | 94.9 | 145.8 |
| 315 | Histone H3 OS=Mus musculus OX=10090 GN=H3f3a PE=1 SV=1 | 80.6 | 131.9 | 107.1 | 62.3 | 197.6 | 59.8 | 73.3 | 88.3 | 52.8 | 146.3 |
| 316 | PDZ and LIM domain protein 5 OS=Mus musculus OX=10090 GN=Pdlim5 PE=1 SV=2 | | 40.1 | 20.8 | 36.8 | 259.6 | 34.7 | 433.9 | 60 | 35.8 | 78.3 |
| 317 | Myosin, heavy polypeptide 2, skeletal muscle, adult OS=Mus musculus OX=10090 GN=Myh2 PE=1 SV=1 | 32.4 | 54.5 | 27.5 | 32.6 | 246.9 | 34.5 | 458.2 | 38 | 24.5 | 50.9 |
| 318 | Outer mitochondrial membrane protein porin 2 OS=Mus musculus OX=10090 GN=Vdac2 PE=1 SV=1 | 92.6 | 83.8 | 87.9 | 110.2 | 101.6 | 95.2 | 86 | 118.3 | 121.3 | 103.1 |
| 319 | Heterogeneous nuclear ribonucleoprotein Q OS=Mus musculus OX=10090 GN=Syncrip PE=1 SV=1 | 96.8 | 89.5 | 102 | 79.5 | 89.6 | 107.3 | 98.2 | 90 | 113.8 | 133.2 |
| 320 | General transcription factor II-I OS=Mus musculus OX=10090 GN=Gtf2i PE=1 SV=1 | | 186 | | 139.2 | | 197.5 | 111.1 | 114.3 | 133.2 | 118.7 |
| 321 | Bifunctional UDP-N-acetylglucosamine 2-epimerase/N-acetylmannosamine kinase OS=Mus musculus OX=10090 GN=Gne PE=1 SV=1 | 106.5 | 98 | 103 | 90.4 | 79.7 | 103.1 | 115.9 | 118.1 | 88.2 | 97.2 |
| 322 | Aldehyde dehydrogenase (NAD(+)) OS=Mus musculus OX=10090 GN=Aldh7a1 PE=1 SV=1 | 91.9 | 84.1 | 92.5 | 155.9 | 60.1 | 111.2 | 43.1 | 113.7 | 145.2 | 102.3 |
| 323 | 40S ribosomal protein S28 (Fragment) OS=Mus musculus OX=10090 GN=Rps28 PE=1 SV=1 | 95.3 | 99.5 | 110 | 78.3 | 122 | 117 | 130.9 | 79.6 | 74.3 | 93 |
| 324 | Ceruloplasmin (Fragment) OS=Mus musculus OX=10090 GN=Cp PE=1 SV=1 | 87.4 | 89.3 | 94.9 | 80.5 | 131.7 | 96.8 | 123.7 | 105.3 | 88 | 102.3 |
| 325 | D-dopachrome decarboxylase OS=Mus musculus OX=10090 GN=Ddt PE=1 SV=1 | 105 | 114.5 | 106.9 | 107.6 | 74.4 | 92.7 | 117.2 | 110.5 | 92.5 | 78.7 |
| 326 | Casein kinase II subunit beta OS=Mus musculus OX=10090 GN=Csnk2b PE=1 SV=1 | 87.5 | 92.6 | 94 | 100.8 | 94.6 | 87.4 | 130.3 | 104.5 | 105.2 | 103.1 |
| 327 | Dynamin GTPase (Fragment) OS=Mus musculus OX=10090 GN=Dnm2 PE=1 SV=1 | 114.4 | 97.1 | 117.6 | 101.7 | 96.8 | 93.4 | 67.2 | 117.7 | 103.1 | 91 |
| 328 | Glycine hydroxymethyltransferase (Fragment) OS=Mus musculus OX=10090 GN=Shmt1 PE=1 SV=1 | 146.9 | 136.4 | 129.5 | 93.7 | 104 | 118.6 | 38.3 | 97.9 | 82 | 52.7 |
| 329 | Ceruloplasmin OS=Mus musculus OX=10090 GN=Cp PE=1 SV=1 | 102.8 | 97.8 | 110.1 | 80 | 99.9 | 98.2 | 95.9 | 108.4 | 94 | 112.9 |
| 330 | Carboxypeptidase OS=Mus musculus OX=10090 GN=Ctsa PE=1 SV=1 | 91.7 | 93.5 | 89.8 | 78.3 | 140.9 | 104 | 126.5 | 94.4 | 86 | 95 |
| 331 | Aldehyde oxidase 3 OS=Mus musculus OX=10090 GN=Aox3 PE=1 SV=1 | 144.4 | 130 | 129.2 | 103.2 | 84.5 | 119.4 | 59.8 | 75.4 | 88.5 | 65.6 |
| 332 | Aldo-keto reductase family 1, member C19 OS=Mus musculus OX=10090 GN=Akr1c19 PE=1 SV=1 | 106.7 | 106.5 | 106.8 | 109 | 94.3 | 114.6 | 81.9 | 90.6 | 103.7 | 85.9 |
| 333 | Isopentenyl-diphosphate Delta-isomerase OS=Mus musculus OX=10090 GN=Idi1 PE=1 SV=1 | 127.2 | 100.9 | 87 | 114.1 | 77 | 128.9 | 68.2 | 63.7 | 117.6 | 115.3 |
| 334 | Plasma membrane calcium-transporting ATPase 1 OS=Mus musculus OX=10090 GN=Atp2b1 PE=1 SV=1 | 135.8 | 119.9 | 91.1 | 84 | 67.1 | 119.6 | 112 | 78.2 | 129.6 | 62.8 |
| 335 | Periplakin OS=Mus musculus OX=10090 GN=Ppl PE=1 SV=1 | 89.1 | 93 | 102.8 | 103 | 109.8 | 91.2 | 101.7 | 92.7 | 101.3 | 115.4 |
| 336 | Heterogeneous nuclear ribonucleoprotein D0 OS=Mus musculus OX=10090 GN=Hnrnpd PE=1 SV=1 | 102 | 92.8 | 80.8 | 73.9 | 125.7 | 98.3 | 74.7 | 101.9 | 82.8 | 167.1 |
| 337 | Glutathione transferase OS=Mus musculus OX=10090 GN=Gstm6 PE=1 SV=1 | 92.3 | 87 | 104 | 132.3 | 76.3 | 122.1 | 78.8 | 92 | 110.1 | 105 |
| 338 | Methyltransferase hypoxia-inducible domain-containing 1 OS=Mus musculus OX=10090 GN=Methig1 PE=4 SV=1 | 145 | 85.6 | 92.7 | 94.7 | 72.9 | 83.1 | 65.2 | 164 | 95.3 | 101.5 |
| 339 | Esterase D OS=Mus musculus OX=10090 GN=Esd PE=1 SV=1 | 154.1 | 118 | 141.1 | 103.6 | 73.8 | 107.8 | 67.6 | 88.6 | 85.6 | 59.8 |
| 340 | Fumarate hydratase 1 (Fragment) OS=Mus musculus OX=10090 GN=Fh1 PE=1 SV=1 | 110.4 | 79.4 | 88.4 | 133.3 | 50.4 | 112.4 | 90.4 | 132.2 | 109.2 | 93.9 |
| 341 | NOL1/NOP2/Sun domain family member 2 OS=Mus musculus OX=10090 GN=Nsun2 PE=1 SV=1 | 124.9 | 93.3 | 108.6 | 105.7 | 73.8 | 106.6 | 53.5 | 114.3 | 106.2 | 113 |
| 342 | Carboxylic ester hydrolase OS=Mus musculus OX=10090 GN=Ces1e PE=1 SV=1 | | 138.4 | | 158.3 | | 157.3 | 112.7 | 140.2 | 157 | 136.2 |
| 343 | CCT-theta OS=Mus musculus OX=10090 GN=Cct8 PE=1 SV=1 | 136 | 107 | 123.2 | 112.4 | 68.3 | 108.4 | 78.2 | 93.4 | 89.4 | 83.8 |
| 344 | Paraoxonase OS=Mus musculus OX=10090 GN=Pon1 PE=1 SV=1 | 119.2 | 107.1 | 125.3 | 131.8 | 56.8 | 100.9 | 38.3 | 113.3 | 141.6 | 65.6 |

| | | | | | | | | | | | |
|---|---|---|---|---|---|---|---|---|---|---|---|
| 345 | Heterogeneous nuclear ribonucleoprotein K OS=Mus musculus OX=10090 GN=Hnrnpk PE=1 SV=1 | 63.3 | 108.3 | 63.5 | 99 | 71.9 | 132.3 | 100.9 | 120 | 104.4 | 136.5 |
| 346 | Alanine--glyoxylate aminotransferase 2, mitochondrial OS=Mus musculus OX=10090 GN=Agxt2 PE=1 SV=1 | 92.9 | 99.1 | 111.3 | 123.7 | 82.1 | 101.2 | 68.4 | 107.9 | 112.7 | 100.6 |
| 347 | Dimethylaniline monooxygenase [N-oxide-forming] OS=Mus musculus OX=10090 GN=Fmo6 PE=1 SV=1 | 82.3 | 85.9 | 89.1 | 138 | 15.1 | 58.5 | 34.6 | 183.1 | 182.6 | 130.9 |
| 348 | Asialoglycoprotein receptor 2 (Fragment) OS=Mus musculus OX=10090 GN=Asgr2 PE=1 SV=1 | 125.4 | 112.9 | 114.2 | 100.9 | 78.9 | 120.2 | 61.3 | 101.6 | 98 | 86.5 |
| 349 | ATP-dependent (S)-NAD(P)H-hydrate dehydratase OS=Mus musculus OX=10090 GN=Naxd PE=1 SV=1 | 106.7 | 96.3 | 101.1 | 83.5 | 114.4 | 102 | 151.1 | 89.3 | 84.3 | 71.4 |
| 350 | Bile salt export pump OS=Mus musculus OX=10090 GN=Abcb11 PE=1 SV=1 | 110.4 | 104.9 | 97.5 | 98.7 | 80.6 | 101.1 | 75.6 | 112.2 | 120.5 | 98.5 |
| 351 | NAD(P)HX dehydratase OS=Mus musculus OX=10090 GN=Naxd PE=1 SV=1 | 106.7 | 96.3 | 101.1 | 83.5 | 114.4 | 102 | 151.1 | 89.3 | 84.3 | 71.4 |
| 352 | Collagen, type VI, alpha 3 OS=Mus musculus OX=10090 GN=Col6a3 PE=1 SV=1 | 76.2 | 76.5 | 83.4 | 61.1 | 161.6 | 65.7 | 274.4 | 64.5 | 66.2 | 70.5 |
| 353 | Phenazine biosynthesis-like domain-containing protein 1 OS=Mus musculus OX=10090 GN=Pbld1 PE=1 SV=1 | 135.3 | 110.8 | 124.7 | 111 | 83.6 | 116.3 | 58 | 88.9 | 105.5 | 66 |
| 354 | Cytochrome c oxidase subunit 4 (Fragment) OS=Mus musculus OX=10090 GN=Cox4i1 PE=1 SV=1 | 108 | 79.9 | 108.2 | 103 | 97.9 | 93.6 | 108.3 | 99.4 | 87.7 | 114 |
| 355 | Dihydropyrimidinase-related protein 2 OS=Mus musculus OX=10090 GN=Dpysl2 PE=1 SV=2 | 122.8 | 119.2 | 117.9 | 90.1 | 94.4 | 111.2 | 93.8 | 79.4 | 84.4 | 86.8 |
| 356 | Galectin-9 OS=Mus musculus OX=10090 GN=Lgals9 PE=1 SV=1 | 140.2 | 100.2 | 119.8 | 101 | 76.6 | 116.5 | 79.9 | 92.1 | 85 | 88.8 |
| 357 | Endonuclease G, mitochondrial OS=Mus musculus OX=10090 GN=Endog PE=1 SV=1 | 98.8 | 93.1 | 111.2 | 91.4 | 119.9 | 116.8 | 106.3 | 97.9 | 85.4 | 79.1 |
| 358 | Microsomal triglyceride transfer protein large subunit OS=Mus musculus OX=10090 GN=Mttp PE=1 SV=2 | 151.1 | 109.9 | 143.4 | 87.7 | 79.4 | 110.2 | 63.7 | 99.7 | 87.6 | 67.5 |
| 359 | Myosin-11 OS=Mus musculus OX=10090 GN=Myh11 PE=1 SV=1 | 24.1 | 206.6 | 20.4 | 96.3 | 13.3 | 84.5 | 264.1 | 86.7 | 109 | 94.9 |
| 360 | Neutrophilic granule protein OS=Mus musculus OX=10090 GN=Ngp PE=1 SV=1 | | 105.9 | | 91.1 | | 165 | 141 | 204.3 | 107.8 | 184.8 |
| 361 | A-kinase anchor protein 1, mitochondrial OS=Mus musculus OX=10090 GN=Akap1 PE=1 SV=4 | 121.3 | 109 | 115.6 | 88.5 | 95.6 | 99.9 | 106.1 | 95.2 | 85.7 | 83.1 |
| 362 | Caspase-6 OS=Mus musculus OX=10090 GN=Casp6 PE=1 SV=1 | | 124.6 | 118.4 | 114.7 | 100.3 | 116.7 | 98.4 | 102.2 | 101.7 | 122.9 |
| 363 | Dihydrolipoyl dehydrogenase, mitochondrial OS=Mus musculus OX=10090 GN=Dld PE=1 SV=2 | 109.7 | 92.9 | 99 | 110.6 | 94.7 | 91 | 83.8 | 117.3 | 108.2 | 92.9 |
| 364 | 3-hydroxyacyl-CoA dehydrogenase type-2 OS=Mus musculus OX=10090 GN=Hsd17b10 PE=1 SV=4 | 65.9 | 96.3 | 70.6 | 123.1 | 48.4 | 95 | 78.1 | 129.1 | 134.7 | 158.9 |
| 365 | Glucosidase 2 subunit beta OS=Mus musculus OX=10090 GN=Prkcsh PE=1 SV=1 | 91 | 107.7 | 117.2 | 88.2 | 120.3 | 118.6 | 73.4 | 89 | 87.6 | 106.9 |
| 366 | Fatty-acid amide hydrolase 1 OS=Mus musculus OX=10090 GN=Faah PE=1 SV=1 | 140.2 | 118.7 | 122.4 | 95.6 | 90.2 | 105.9 | 87.4 | 85.6 | 88.7 | 65.3 |
| 367 | Dimethylargininase OS=Mus musculus OX=10090 GN=Ddah2 PE=1 SV=1 | 116.9 | 103.2 | 102.7 | 116.7 | 91.3 | 110.1 | 73 | 102.4 | 106.9 | 76.7 |
| 368 | Copper transport protein ATOX1 OS=Mus musculus OX=10090 GN=Atox1 PE=1 SV=1 | 68.4 | 101.8 | 71.9 | 64.2 | 184.5 | 84.3 | 106.3 | 126.1 | 69.2 | 123.2 |
| 369 | Histone deacetylase 1 OS=Mus musculus OX=10090 GN=Hdac1 PE=1 SV=1 | | 152.4 | | 108.7 | | 182.7 | 144.9 | 129 | 118.8 | 163.5 |
| 370 | NADH dehydrogenase [ubiquinone] 1 beta subcomplex subunit 11, mitochondrial OS=Mus musculus OX=10090 GN=Ndufb11 PE=1 SV=2 | 84.8 | 91.8 | 82.5 | 111.3 | 104.1 | 86.3 | 121.6 | 110.6 | 100.2 | 106.8 |
| 371 | Glutathione S-transferase omega-1 OS=Mus musculus OX=10090 GN=Gsto1 PE=1 SV=2 | 112.7 | 119.4 | 108.1 | 111.9 | 62.4 | 103.2 | 79.1 | 115.8 | 91.7 | 95.7 |
| 372 | Lysosomal alpha-mannosidase OS=Mus musculus OX=10090 GN=Man2b1 PE=1 SV=4 | 111.7 | 99.1 | 121.9 | 104.8 | 51.2 | 104.1 | 58.4 | 111.9 | 109.7 | 127.1 |
| 373 | Calsequestrin-2 OS=Mus musculus OX=10090 GN=Casq2 PE=1 SV=3 | 8.6 | 36.9 | 10.7 | 10.3 | 327.6 | 8.7 | 555.7 | 8 | 6 | 27.6 |
| 374 | Calsequestrin-1 OS=Mus musculus OX=10090 GN=Casq1 PE=1 SV=3 | 18.1 | 52.6 | 20.6 | 23.3 | 276.2 | 29.5 | 498.6 | 26.3 | 14.4 | 40.4 |
| 375 | 60S ribosomal protein L21 OS=Mus musculus OX=10090 GN=Rpl21 PE=1 SV=3 | 120 | 101.6 | 121.7 | 101.3 | 75.5 | 116.8 | 89.3 | 106.9 | 80.3 | 86.6 |
| 376 | Glutamate--cysteine ligase regulatory subunit OS=Mus musculus OX=10090 GN=Gclm PE=1 SV=1 | 116.6 | 109.1 | 107.8 | 88.7 | 145.8 | 111.5 | 92 | 82.8 | 74.5 | 71.3 |
| 377 | Homogentisate 1,2-dioxygenase OS=Mus musculus OX=10090 GN=Hgd PE=1 SV=2 | 105.7 | 100.6 | 112.9 | 121.9 | 65.8 | 105.8 | 61.3 | 115.6 | 117.1 | 93.4 |
| 378 | Alpha-methylacyl-CoA racemase OS=Mus musculus OX=10090 GN=Amacr PE=1 SV=4 | 88 | 92.5 | 107.1 | 105.6 | 93.6 | 113.7 | 68.9 | 116.5 | 105.8 | 108.4 |
| 379 | Lysosome membrane protein 2 OS=Mus musculus OX=10090 GN=Scarb2 PE=1 SV=3 | 111.8 | 106.3 | 112.7 | 92 | 87.4 | 121.7 | 100.6 | 102.1 | 84.2 | 81.2 |
| 380 | Prohibitin-2 OS=Mus musculus OX=10090 GN=Phb2 PE=1 SV=1 | 110.9 | 95.9 | 108 | 108.4 | 85.1 | 97.5 | 79 | 106.9 | 104.9 | 103.5 |

| | | | | | | | | | | | |
|---|---|---|---|---|---|---|---|---|---|---|---|
| 381 | Cleavage and polyadenylation specificity factor subunit 2 OS=Mus musculus OX=10090 GN=Cpsf2 PE=1 SV=1 | 116.6 | 92.9 | 104.2 | 106 | 94.7 | 100.3 | 91.8 | 81.2 | 116.5 | 95.8 |
| 382 | Importin subunit alpha-4 OS=Mus musculus OX=10090 GN=Kpna3 PE=1 SV=1 | 130.4 | 111.6 | 113.1 | 91.3 | 111 | 106.8 | 86.9 | 88.5 | 87.7 | 72.7 |
| 383 | Phytanoyl-CoA dioxygenase, peroxisomal OS=Mus musculus OX=10090 GN=Phyh PE=1 SV=1 | 92.6 | 91.4 | 90 | 105.3 | 89.4 | 120.6 | 64.8 | 121.9 | 115.8 | 108.3 |
| 384 | HCLS1-associated protein X-1 OS=Mus musculus OX=10090 GN=Hax1 PE=1 SV=1 | 150.6 | 106.5 | 116.1 | 114.3 | 84.2 | 86.9 | 55.4 | 99.2 | 106.9 | 80 |
| 385 | Delta(3,5)-Delta(2,4)-dienoyl-CoA isomerase, mitochondrial OS=Mus musculus OX=10090 GN=Ech1 PE=1 SV=1 | 129.5 | 112.5 | 119.8 | 111.7 | 68.2 | 90 | 61.1 | 105.2 | 110.5 | 91.5 |
| 386 | Peptidyl-prolyl cis-trans isomerase FKBP8 OS=Mus musculus OX=10090 GN=Fkbp8 PE=1 SV=2 | 109.8 | 105.5 | 116 | 93.6 | 98 | 113.1 | 95 | 93.7 | 92.3 | 82.9 |
| 387 | Betaine--homocysteine S-methyltransferase 1 OS=Mus musculus OX=10090 GN=Bhmt PE=1 SV=1 | 88.7 | 86.8 | 114.6 | 163.4 | 73.2 | 93.2 | 46.9 | 95.2 | 136.4 | 101.5 |
| 388 | NPC intracellular cholesterol transporter 1 OS=Mus musculus OX=10090 GN=Npc1 PE=1 SV=2 | 96 | 84.8 | 98.6 | 94.2 | 105.7 | 123.6 | 96.4 | 113.9 | 88 | 98.8 |
| 389 | AP-1 complex subunit beta-1 OS=Mus musculus OX=10090 GN=Ap1b1 PE=1 SV=2 | 132 | 101.5 | 114.4 | 91.1 | 74.9 | 127.8 | 80.1 | 104.1 | 91.9 | 82.2 |
| 390 | Cytochrome P450 4A14 OS=Mus musculus OX=10090 GN=Cyp4a14 PE=1 SV=1 | 71.6 | 71.1 | 101.6 | 123.3 | 66.9 | 76 | 65.3 | 166.1 | 157.6 | 100.4 |
| 391 | Chitinase-like protein 3 OS=Mus musculus OX=10090 GN=Chil3 PE=1 SV=2 | 84.9 | 129.7 | 63.6 | 64.2 | 75.5 | 139.1 | 45.2 | 139.5 | 97.9 | 160.4 |
| 392 | Mitochondrial import inner membrane translocase subunit TIM44 OS=Mus musculus OX=10090 GN=Timm44 PE=1 SV=2 | 101 | 96.9 | 98 | 94.2 | 98.8 | 109.7 | 102.4 | 98.5 | 91.7 | 108.8 |
| 393 | Aldehyde dehydrogenase, cytosolic 1 OS=Mus musculus OX=10090 GN=Aldh1a7 PE=1 SV=1 | 106.7 | 110.1 | 128 | 120.6 | 84.3 | 102.6 | 50.1 | 110.4 | 97.9 | 89.3 |
| 394 | Caveolae-associated protein 1 OS=Mus musculus OX=10090 GN=Cavin1 PE=1 SV=1 | 62.3 | 65.9 | 66.9 | 66.7 | 238.6 | 61.8 | 237.8 | 75 | 53.3 | 71.9 |
| 395 | Dolichyl-diphosphooligosaccharide--protein glycosyltransferase 48 kDa subunit OS=Mus musculus OX=10090 GN=Ddost PE=1 SV=2 | 135.2 | 125.3 | 142.1 | 91.4 | 83.6 | 130.5 | 65.6 | 80.6 | 77.8 | 67.9 |
| 396 | Cytochrome P450 2J5 OS=Mus musculus OX=10090 GN=Cyp2j5 PE=1 SV=1 | 137.7 | 105.5 | 117.5 | 68.7 | 146.5 | 98.6 | 122.4 | 104 | 55.5 | 43.6 |
| 397 | Membrane-associated progesterone receptor component 1 OS=Mus musculus OX=10090 GN=Pgrmc1 PE=1 SV=4 | 124.9 | 110.5 | 116.4 | 101.7 | 117 | 101.8 | 72.5 | 93.8 | 86.1 | 75.3 |
| 398 | Coatomer subunit beta' OS=Mus musculus OX=10090 GN=Copb2 PE=1 SV=2 | 133.7 | 109.4 | 130 | 91.9 | 73.8 | 124.6 | 74 | 98.5 | 88.7 | 75.4 |
| 399 | Desmoglein-2 OS=Mus musculus OX=10090 GN=Dsg2 PE=1 SV=3 | 107.1 | 107.9 | 112.4 | 92.6 | 96.4 | 108.2 | 85.3 | 90.2 | 92.5 | 107.3 |
| 400 | Eukaryotic translation initiation factor 6 OS=Mus musculus OX=10090 GN=Eif6 PE=1 SV=2 | 88.8 | 96.6 | 100 | 99.1 | 112.5 | 96.4 | 63.7 | 127.9 | 80.4 | 134.7 |
| 401 | 60S ribosomal protein L35a OS=Mus musculus OX=10090 GN=Rpl35a PE=1 SV=2 | 138 | 121.3 | 110.6 | 94.3 | 80.4 | 123 | 49 | 106.3 | 85.9 | 91 |
| 402 | Integrin-linked protein kinase OS=Mus musculus OX=10090 GN=Ilk PE=1 SV=2 | 74.9 | 94.9 | 76.8 | 85.8 | 118.5 | 102.3 | 134.5 | 96.7 | 96.9 | 118.7 |
| 403 | Eukaryotic translation initiation factor 3 subunit D OS=Mus musculus OX=10090 GN=Eif3d PE=1 SV=2 | 113.9 | 103.3 | 96.8 | 83.7 | 109.4 | 101.9 | 126.1 | 88 | 85.4 | 91.5 |
| 404 | Elongation factor 1-beta OS=Mus musculus OX=10090 GN=Eef1b PE=1 SV=5 | 87.7 | 98.4 | 83.3 | 66 | 182.8 | 124.3 | 57.1 | 95.6 | 69.3 | 135.5 |
| 405 | Phospholipid hydroperoxide glutathione peroxidase OS=Mus musculus OX=10090 GN=Gpx4 PE=1 SV=4 | 99.6 | 88.9 | 92.2 | 111.8 | 91.5 | 87.8 | 86.3 | 115.1 | 114.9 | 111.8 |
| 406 | PDZ and LIM domain protein 1 OS=Mus musculus OX=10090 GN=Pdlim1 PE=1 SV=4 | 85.1 | 90.1 | 77.7 | 62.1 | 205.2 | 89.7 | 115.5 | 85.5 | 59.6 | 129.3 |
| 407 | Alpha-N-acetylglucosaminidase OS=Mus musculus OX=10090 GN=Naglu PE=1 SV=1 | | 130 | 134.5 | 97.5 | 117.9 | 129.5 | 88.3 | 93.5 | 106.5 | 102.3 |
| 408 | Metaxin-2 OS=Mus musculus OX=10090 GN=Mtx2 PE=1 SV=1 | | 122.6 | 116.1 | 106.7 | 92.1 | 112 | 114 | 116.2 | 103.3 | 116.9 |
| 409 | 7-dehydrocholesterol reductase OS=Mus musculus OX=10090 GN=Dhcr7 PE=1 SV=1 | 151.5 | 129.9 | 119.3 | 86.6 | 102.8 | 110.9 | 75.3 | 92.7 | 79 | 51.9 |
| 410 | Palmitoyl-protein thioesterase 1 OS=Mus musculus OX=10090 GN=Ppt1 PE=1 SV=2 | 106.6 | 113.2 | 96.3 | 73.6 | 118.7 | 117.5 | 86.9 | 100.8 | 86.9 | 99.6 |
| 411 | Aromatic-L-amino-acid decarboxylase OS=Mus musculus OX=10090 GN=Ddc PE=1 SV=1 | 102.3 | 107.7 | 114 | 124.5 | 86.7 | 95.4 | 67.7 | 100.2 | 108.6 | 92.9 |
| 412 | COP9 signalosome complex subunit 3 OS=Mus musculus OX=10090 GN=Cops3 PE=1 SV=3 | 107.3 | 102.7 | 98.8 | 98.1 | 98.2 | 108.6 | 93.2 | 109.8 | 94.6 | 88.8 |
| 413 | Heterogeneous nuclear ribonucleoproteins A2/B1 OS=Mus musculus OX=10090 GN=Hnrnpa2b1 PE=1 SV=2 | 49.5 | 48.8 | 49 | 46.5 | 48.7 | 56.4 | 120.3 | 53.5 | 202.8 | 324.4 |
| 414 | Catechol O-methyltransferase OS=Mus musculus OX=10090 GN=Comt PE=1 SV=2 | 133.1 | 98.8 | 119.2 | 100.6 | 93.5 | 114.7 | 92.7 | 96.5 | 76.5 | 74.4 |
| 415 | Isocitrate dehydrogenase [NADP] cytoplasmic OS=Mus musculus OX=10090 GN=Idh1 PE=1 SV=2 | 50.8 | 149.6 | 140 | 106.4 | 92.9 | 142.5 | 78.5 | 84.7 | 83.3 | 71.3 |

| | | | | | | | | | | | |
|---|---|---|---|---|---|---|---|---|---|---|---|
| 416 | 7-alpha-hydroxycholest-4-en-3-one 12-alpha-hydroxylase OS=Mus musculus OX=10090 GN=Cyp8b1 PE=1 SV=1 | 141.9 | 102.5 | 150.9 | 168.3 | 55.9 | 85.6 | 52.6 | 71.5 | 123.5 | 47.3 |
| 417 | Legumain OS=Mus musculus OX=10090 GN=Lgmn PE=1 SV=1 | 97.5 | 95 | 97.5 | 89.1 | 102.4 | 122.8 | 57.7 | 98.1 | 98.2 | 141.7 |
| 418 | Coronin-1A OS=Mus musculus OX=10090 GN=Coro1a PE=1 SV=5 | 97.9 | 88.6 | 101 | 81.9 | 105.3 | 171.2 | 93.7 | 114.1 | 64.9 | 81.3 |
| 419 | Cytochrome c, testis-specific OS=Mus musculus OX=10090 GN=Cyct PE=1 SV=3 | 47.3 | 76.9 | 67.6 | 87 | 185.9 | 81.8 | 81.4 | 139.5 | 78.4 | 154.1 |
| 420 | Cytochrome P450 1A2 OS=Mus musculus OX=10090 GN=Cyp1a2 PE=1 SV=1 | 169.9 | 140.8 | 168.9 | 119.6 | 61.4 | 104.3 | 38.2 | 71.7 | 77.1 | 48.3 |
| 421 | Alcohol dehydrogenase 1 OS=Mus musculus OX=10090 GN=Adh1 PE=1 SV=2 | 160.8 | 118.7 | 142.3 | 99.2 | 74.3 | 111.7 | 53.6 | 89.5 | 79.7 | 70.2 |
| 422 | Dihydrofolate reductase OS=Mus musculus OX=10090 GN=Dhfr PE=1 SV=3 | 119.6 | 110.4 | 125.4 | 105.7 | 84 | 128.5 | 82.4 | 83.2 | 78.2 | 82.7 |
| 423 | Cytochrome c oxidase subunit 1 OS=Mus musculus OX=10090 GN=Mtco1 PE=1 SV=2 | | 115.9 | 104.1 | 155 | 49.7 | 77 | 80.7 | 150.6 | 160.9 | 106 |
| 424 | Cytochrome c oxidase subunit 2 OS=Mus musculus OX=10090 GN=Mtco2 PE=1 SV=1 | 81.8 | 84.8 | 89.6 | 104 | 115.5 | 93.6 | 122 | 104.6 | 104 | 100.3 |
| 425 | Alpha-amylase 1 OS=Mus musculus OX=10090 GN=Amy1 PE=1 SV=2 | 244.7 | 36.2 | 257.7 | 19.7 | 67.7 | 64.9 | 103.3 | 39.9 | 57.7 | 107.9 |
| 426 | Carbonic anhydrase 2 OS=Mus musculus OX=10090 GN=Ca2 PE=1 SV=4 | 96 | 101.8 | 132.2 | 75.9 | 119.4 | 69.6 | 163.4 | 78.6 | 82.6 | 80.6 |
| 427 | Complement C3 OS=Mus musculus OX=10090 GN=C3 PE=1 SV=3 | 121.8 | 123.9 | 146.5 | 94.2 | 87.3 | 123 | 70.9 | 95.9 | 72.8 | 63.6 |
| 428 | Complement C4-B OS=Mus musculus OX=10090 GN=C4b PE=1 SV=3 | 114.7 | 94.7 | 122.4 | 86.5 | 79.3 | 121.6 | 121 | 89.1 | 89.9 | 80.8 |
| 429 | Immunoglobulin kappa constant OS=Mus musculus OX=10090 GN=Igkc PE=1 SV=2 | 84.8 | 105.6 | 148.6 | 95.2 | 99.9 | 95.4 | 105.6 | 104.7 | 70.4 | 89.7 |
| 430 | H-2 class I histocompatibility antigen, Q10 alpha chain OS=Mus musculus OX=10090 GN=H2-Q10 PE=1 SV=3 | 102 | 103 | 111.1 | 94.2 | 107.7 | 115.8 | 112 | 88 | 85.8 | 80.3 |
| 431 | H-2 class I histocompatibility antigen, K-B alpha chain OS=Mus musculus OX=10090 GN=H2-K1 PE=1 SV=1 | 55.9 | 107.5 | 74.8 | 92.5 | 102.4 | 171.5 | 153.5 | 81.4 | 68.6 | 91.9 |
| 432 | Hemoglobin subunit alpha OS=Mus musculus OX=10090 GN=Hba PE=1 SV=2 | 98.4 | 106.6 | 190.8 | 82.3 | 70.9 | 86 | 48.2 | 110.8 | 113.2 | 92.8 |
| 433 | Histone H3.3C OS=Mus musculus OX=10090 GN=H3-5 PE=3 SV=3 | 80.6 | 131.9 | 107.1 | 62.3 | 197.6 | 59.8 | 73.3 | 88.3 | 52.8 | 146.3 |
| 434 | NADH-ubiquinone oxidoreductase chain 1 OS=Mus musculus OX=10090 GN=Mtnd1 PE=1 SV=3 | 106.5 | 81 | 103.9 | 131.6 | 75.9 | 66.5 | 83.1 | 136.1 | 121.3 | 94.2 |
| 435 | NADH-ubiquinone oxidoreductase chain 4 OS=Mus musculus OX=10090 GN=Mtnd4 PE=1 SV=1 | 77.4 | 89.5 | 117.8 | 90.1 | 171.7 | 99.3 | 86.5 | 109.4 | 84.5 | 73.9 |
| 436 | ATP synthase protein 8 OS=Mus musculus OX=10090 GN=Mtatp8 PE=1 SV=1 | 83.1 | 97.3 | 90.2 | 88 | 114.8 | 86.2 | 171.8 | 99.1 | 78 | 91.5 |
| 437 | Fatty acid-binding protein, adipocyte OS=Mus musculus OX=10090 GN=Fabp4 PE=1 SV=3 | 58.9 | 61.7 | 65.3 | 79.9 | 176.4 | 70.4 | 196.7 | 89.4 | 68.8 | 132.4 |
| 438 | Complement factor B OS=Mus musculus OX=10090 GN=Cfb PE=1 SV=2 | 122.1 | 119.9 | 137.9 | 86.7 | 106.5 | 108.5 | 87.4 | 76.7 | 78.1 | 76.2 |
| 439 | Band 3 anion transport protein OS=Mus musculus OX=10090 GN=Slc4a1 PE=1 SV=1 | 85.4 | 86.8 | 112.6 | 89.7 | 84.1 | 89.4 | 104.7 | 96.6 | 122.1 | 128.7 |
| 440 | Fructose-bisphosphate aldolase C OS=Mus musculus OX=10090 GN=Aldoc PE=1 SV=4 | 96.9 | 92.9 | 91.7 | 107.9 | 92.4 | 94.9 | 120.6 | 106.4 | 87 | 109.3 |
| 441 | Fructose-bisphosphate aldolase A OS=Mus musculus OX=10090 GN=Aldoa PE=1 SV=2 | 15.9 | 22.6 | 14.7 | 47.5 | 84.2 | 44.6 | 638.2 | 21.7 | 33 | 77.7 |
| 442 | Aspartate aminotransferase, cytoplasmic OS=Mus musculus OX=10090 GN=Got1 PE=1 SV=3 | 37.4 | 33.7 | 36.1 | 157.9 | 64.7 | 91 | 167 | 61.3 | 184.2 | 166.5 |
| 443 | Aspartate aminotransferase, mitochondrial OS=Mus musculus OX=10090 GN=Got2 PE=1 SV=1 | 99.6 | 91.2 | 97.8 | 101.4 | 95.8 | 97.7 | 104.4 | 99.7 | 104.9 | 107.5 |
| 444 | Myosin light chain 1/3, skeletal muscle isoform OS=Mus musculus OX=10090 GN=Myl1 PE=1 SV=2 | 35.6 | 44.6 | 27.8 | 36.4 | 388 | 33.9 | 271.8 | 65.4 | 30.7 | 65.9 |
| 445 | Hemoglobin subunit zeta OS=Mus musculus OX=10090 GN=Hbz PE=2 SV=2 | 118.2 | 102.8 | 197.1 | 78.2 | 66 | 83.2 | 46.9 | 102.7 | 113.8 | 91.2 |
| 446 | Apolipoprotein A-IV OS=Mus musculus OX=10090 GN=Apoa4 PE=1 SV=3 | 71.6 | 98.7 | 89.7 | 78.9 | 104.8 | 96.9 | 87.5 | 141.9 | 80 | 149.9 |
| 447 | Glucose-6-phosphate isomerase OS=Mus musculus OX=10090 GN=Gpi PE=1 SV=4 | 110.9 | 95.3 | 120.7 | 94.7 | 118.9 | 97.3 | 113.3 | 94.5 | 81.7 | 72.6 |
| 448 | ATP-dependent translocase ABCB1 OS=Mus musculus OX=10090 GN=Abcb1b PE=1 SV=1 | 89.5 | 99.7 | 98.9 | 104.2 | 86.6 | 82.4 | 81 | 117.6 | 131.4 | 108.6 |
| 449 | NADP-dependent malic enzyme OS=Mus musculus OX=10090 GN=Me1 PE=1 SV=2 | 217.4 | 163 | 168.8 | 61.2 | 83.7 | 93.9 | 80.6 | 41.8 | 46.6 | 43.1 |
| 450 | Creatine kinase M-type OS=Mus musculus OX=10090 GN=Ckm PE=1 SV=1 | 35.7 | 53.2 | 19.4 | 27.4 | 339.3 | 19.4 | 413.3 | 23.5 | 12.8 | 55.9 |
| 451 | Annexin A2 OS=Mus musculus OX=10090 GN=Anxa2 PE=1 SV=2 | 93.5 | 93.3 | 97.6 | 86.2 | 128.2 | 115 | 141.7 | 74.4 | 77.2 | 92.9 |

| | | | | | | | | | | | |
|---|---|---|---|---|---|---|---|---|---|---|---|
| 452 | Albumin OS=Mus musculus OX=10090 GN=Alb PE=1 SV=3 | 123.7 | 117.8 | 128.5 | 73.8 | 122.2 | 87.1 | 127.9 | 70.8 | 66.2 | 82 |
| 453 | Alpha-1-antitrypsin 1-1 OS=Mus musculus OX=10090 GN=Serpina1a PE=1 SV=4 | 101.5 | 123.8 | 131.1 | 96.2 | 88.4 | 82.8 | 98.4 | 88 | 86.1 | 103.7 |
| 454 | Heat shock protein HSP 90-alpha OS=Mus musculus OX=10090 GN=Hsp90aa1 PE=1 SV=4 | 134.9 | 113.9 | 141.3 | 82.7 | 104.8 | 119.9 | 81.5 | 85.9 | 73.6 | 61.6 |
| 455 | Phosphorylase b kinase gamma catalytic chain, skeletal muscle/heart isoform OS=Mus musculus OX=10090 GN=Phkg1 PE=1 SV=3 | 127.8 | 91.8 | 108.1 | 91.3 | 95.9 | 106.9 | 104.1 | 104 | 70.1 | 99.9 |
| 456 | Carbonyl reductase [NADPH] 2 OS=Mus musculus OX=10090 GN=Cbr2 PE=1 SV=1 | 96.1 | 101.8 | 140 | 140.8 | 74.4 | 99.1 | 53.5 | 98.9 | 111.1 | 84.3 |
| 457 | Endoplasmin OS=Mus musculus OX=10090 GN=Hsp90b1 PE=1 SV=2 | 118.2 | 106.3 | 128.7 | 78.6 | 117.9 | 127.7 | 109.1 | 76.6 | 73.6 | 63.3 |
| 458 | Collagen alpha-1(III) chain OS=Mus musculus OX=10090 GN=Col3a1 PE=1 SV=4 | 66.3 | 88.1 | 77.6 | 76.9 | 137.5 | 87 | 96.1 | 106.4 | 86.8 | 177.3 |
| 459 | Collagen alpha-2(IV) chain OS=Mus musculus OX=10090 GN=Col4a2 PE=1 SV=4 | 77.8 | 97.6 | 92.9 | 88.8 | 111.8 | 84.8 | 143.9 | 85.3 | 102.2 | 114.9 |
| 460 | Apolipoprotein E OS=Mus musculus OX=10090 GN=Apoe PE=1 SV=2 | 112.3 | 122.8 | 119.8 | 91.3 | 115.8 | 115.9 | 89.7 | 81.7 | 63.3 | 87.5 |
| 461 | Malate dehydrogenase, mitochondrial OS=Mus musculus OX=10090 GN=Mdh2 PE=1 SV=3 | 76.9 | 79.5 | 73.9 | 106.4 | 114 | 99 | 110 | 128.9 | 113.7 | 97.8 |
| 462 | Phosphoglycerate kinase 2 OS=Mus musculus OX=10090 GN=Pgk2 PE=1 SV=4 | 112 | 104 | 98 | 99.8 | 85.3 | 103.6 | 89.9 | 111.3 | 99.1 | 97 |
| 463 | Integrin beta-1 OS=Mus musculus OX=10090 GN=Itgb1 PE=1 SV=1 | 74.2 | 105.4 | 79.7 | 96.7 | 72.8 | 123.3 | 104.4 | 102.8 | 109.5 | 131.3 |
| 464 | Nucleolin OS=Mus musculus OX=10090 GN=Ncl PE=1 SV=2 | 69.9 | 87.1 | 71.3 | 68.3 | 93.3 | 108 | 99.2 | 120.5 | 79.5 | 202.9 |
| 465 | Phosphoglycerate kinase 1 OS=Mus musculus OX=10090 GN=Pgk1 PE=1 SV=4 | 109.5 | 99.8 | 96.6 | 102.9 | 103.6 | 95 | 99.2 | 106.4 | 93.4 | 93.5 |
| 466 | Ferritin heavy chain OS=Mus musculus OX=10090 GN=Fth1 PE=1 SV=2 | 66 | 92.8 | 97.6 | 66.3 | 64.8 | 99.8 | 70.3 | 119.8 | 81.6 | 240.9 |
| 467 | Myosin light chain 3 OS=Mus musculus OX=10090 GN=Myl3 PE=1 SV=4 | 13.5 | 32.7 | 14 | 20.1 | 426.2 | 16.2 | 350.2 | 19.5 | 15.6 | 92 |
| 468 | Polyubiquitin-C OS=Mus musculus OX=10090 GN=Ubc PE=1 SV=2 | 96.5 | 118.7 | 103 | 82.6 | 117.6 | 107.7 | 81.4 | 121.1 | 98.3 | 73.1 |
| 469 | Calmodulin-2 OS=Mus musculus OX=10090 GN=Calm2 PE=1 SV=1 | 112.6 | 114.1 | 67.1 | 78.4 | 131.1 | 95.7 | 79 | 145.1 | 50 | 127 |
| 470 | Annexin A1 OS=Mus musculus OX=10090 GN=Anxa1 PE=1 SV=2 | | 113.2 | 100 | 88.6 | 159.9 | 90.1 | 164 | 98 | 80.2 | 106 |
| 471 | Elongation factor 1-alpha 1 OS=Mus musculus OX=10090 GN=Eef1a1 PE=1 SV=3 | 136.1 | 119.4 | 127 | 89 | 98.8 | 122.5 | 71.1 | 90.3 | 80 | 65.7 |
| 472 | Nidogen-1 OS=Mus musculus OX=10090 GN=Nid1 PE=1 SV=2 | 64.2 | 89.8 | 84.9 | 72.9 | 159.8 | 61.7 | 174.7 | 108.3 | 78.9 | 104.8 |
| 473 | Delta-aminolevulinic acid dehydratase OS=Mus musculus OX=10090 GN=Alad PE=1 SV=1 | 189.5 | 125.7 | 169.2 | 120.8 | 52.3 | 125.6 | 40.8 | 71.4 | 63.9 | 40.6 |
| 474 | Cathepsin B OS=Mus musculus OX=10090 GN=Ctsb PE=1 SV=2 | 121.4 | 101.8 | 122.1 | 98.5 | 88.1 | 127 | 55.1 | 92.9 | 100.4 | 92.7 |
| 475 | Eukaryotic initiation factor 4A-II OS=Mus musculus OX=10090 GN=Eif4a2 PE=1 SV=2 | 124.5 | 92.4 | 108 | 90.6 | 95.4 | 111.8 | 76 | 111.7 | 83.7 | 106 |
| 476 | Glutathione S-transferase A2 OS=Mus musculus OX=10090 GN=Gsta2 PE=1 SV=3 | 53.8 | 30 | 36.9 | 171.8 | 104.1 | 76.4 | 60.6 | 168.9 | 207.3 | 90.2 |
| 477 | Glutathione S-transferase Mu 1 OS=Mus musculus OX=10090 GN=Gstm1 PE=1 SV=2 | 87.3 | 80.4 | 85.6 | 129.5 | 66.9 | 124.5 | 65.4 | 103.6 | 136 | 120.7 |
| 478 | Histone H1.0 OS=Mus musculus OX=10090 GN=H1-0 PE=2 SV=4 | 195.6 | 161.6 | 103.8 | 72.8 | 45.8 | 72.7 | 70.8 | 89.2 | 98.1 | 89.6 |
| 479 | Collagen alpha-1(I) chain OS=Mus musculus OX=10090 GN=Col1a1 PE=1 SV=4 | 71.6 | 136.8 | 88 | 63 | 186.6 | 36.9 | 201.6 | 83.5 | 59.9 | 72 |
| 480 | Myeloperoxidase OS=Mus musculus OX=10090 GN=Mpo PE=1 SV=2 | 146.8 | 34 | 20.8 | 276 | 40 | 169 | 147.4 | 91 | 47.4 | 27.7 |
| 481 | Glutathione peroxidase 1 OS=Mus musculus OX=10090 GN=Gpx1 PE=1 SV=2 | 119.4 | 77.8 | 120.5 | 89.4 | 56.3 | 129.3 | 65.7 | 125.8 | 125.1 | 90.8 |
| 482 | Fatty acid-binding protein, heart OS=Mus musculus OX=10090 GN=Fabp3 PE=1 SV=5 | 41.2 | 49.2 | 40.6 | 46.3 | 407.7 | 38.3 | 221.5 | 38.8 | 35.3 | 81.1 |
| 483 | Keratin, type II cytoskeletal 8 OS=Mus musculus OX=10090 GN=Krt8 PE=1 SV=4 | 94.8 | 81.8 | 90.7 | 85.4 | 118.3 | 108.7 | 92.3 | 88.5 | 107.1 | 132.3 |
| 484 | Ornithine transcarbamylase, mitochondrial OS=Mus musculus OX=10090 GN=Otc PE=1 SV=1 | 118 | 101.6 | 114.8 | 108.5 | 91.1 | 112.2 | 55.8 | 101.1 | 108.6 | 88.3 |
| 485 | Fatty acid-binding protein, liver OS=Mus musculus OX=10090 GN=Fabp1 PE=1 SV=2 | 115.9 | 261.8 | 103.7 | 53.1 | 63.3 | 144.6 | 121.4 | 54 | 32.7 | 49.5 |
| 486 | Cytochrome c oxidase subunit 5A, mitochondrial OS=Mus musculus OX=10090 GN=Cox5a PE=1 SV=2 | 85 | 101.7 | 73.4 | 117.2 | 101.8 | 75.1 | 115.4 | 114.3 | 104.5 | 111.7 |
| 487 | Cytochrome P450 2B9 OS=Mus musculus OX=10090 GN=Cyp2b9 PE=1 SV=2 | 145.5 | 126 | 141.9 | 104.4 | 75 | 103.2 | 50.5 | 87.2 | 104.1 | 62.3 |

| | | | | | | | | | | | |
|---|---|---|---|---|---|---|---|---|---|---|---|
| 488 | Cytochrome P450 2B10 OS=Mus musculus OX=10090 GN=Cyp2b10 PE=1 SV=1 | 78.6 | 78.4 | 86.9 | 92.5 | 96.5 | 86.4 | 120.5 | 125.6 | 113.8 | 120.8 |
| 489 | 60S ribosomal protein L7a OS=Mus musculus OX=10090 GN=Rpl7a PE=1 SV=2 | 103.4 | 95.3 | 97.5 | 94.3 | 71.1 | 135.6 | 99.8 | 83.2 | 101.4 | 118.5 |
| 490 | Carbonic anhydrase 1 OS=Mus musculus OX=10090 GN=Ca1 PE=1 SV=4 | 75.9 | 84.4 | 116.8 | 79.3 | 105.8 | 71.2 | 181.8 | 99.2 | 95.6 | 90.1 |
| 491 | Glycerol-3-phosphate dehydrogenase [NAD(+)], cytoplasmic OS=Mus musculus OX=10090 GN=Gpd1 PE=1 SV=3 | 140.2 | 114.5 | 118.5 | 105 | 92.1 | 113.8 | 56.7 | 86.8 | 91.6 | 80.9 |
| 492 | 60S ribosomal protein L27a OS=Mus musculus OX=10090 GN=Rpl27a PE=1 SV=5 | 123.4 | 107.2 | 109.8 | 86.3 | 107.1 | 118.2 | 84.3 | 97.8 | 81.1 | 85 |
| 493 | 40S ribosomal protein S16 OS=Mus musculus OX=10090 GN=Rps16 PE=1 SV=4 | 126.1 | 105.6 | 107.8 | 90.9 | 106 | 122.2 | 74.7 | 95.3 | 83.8 | 87.6 |
| 494 | 60S ribosomal protein L7 OS=Mus musculus OX=10090 GN=Rpl7 PE=1 SV=2 | 114.4 | 118.1 | 110.6 | 83.6 | 91.4 | 125.5 | 103.5 | 96.5 | 79.1 | 77.1 |
| 495 | Malate dehydrogenase, cytoplasmic OS=Mus musculus OX=10090 GN=Mdh1 PE=1 SV=3 | 108.3 | 98 | 116.9 | 111.4 | 99.5 | 93.7 | 107.4 | 90.7 | 94.4 | 79.7 |
| 496 | 40S ribosomal protein SA OS=Mus musculus OX=10090 GN=Rpsa PE=1 SV=4 | 139.7 | 114.5 | 124.5 | 83.6 | 92.3 | 122.5 | 75 | 94.6 | 83.6 | 69.8 |
| 497 | Calreticulin OS=Mus musculus OX=10090 GN=Calr PE=1 SV=1 | 123.4 | 121.8 | 118.2 | 83 | 107.1 | 131.1 | 95.5 | 77.8 | 75 | 67.1 |
| 498 | Lamin-B1 OS=Mus musculus OX=10090 GN=Lmnb1 PE=1 SV=3 | 90.6 | 93.7 | 98.6 | 74.5 | 146.8 | 121.4 | 112.4 | 78.1 | 76.2 | 107.8 |
| 499 | 60S acidic ribosomal protein P0 OS=Mus musculus OX=10090 GN=Rplp0 PE=1 SV=3 | 133.9 | 108.4 | 131 | 93.2 | 90.7 | 125.8 | 52.7 | 90.5 | 84.2 | 89.6 |
| 500 | Glutamine synthetase OS=Mus musculus OX=10090 GN=Glul PE=1 SV=6 | 209 | 168.9 | 156.9 | 100.7 | 48.3 | 97.3 | 48.9 | 64.1 | 62.4 | 43.5 |
| 501 | Bisphosphoglycerate mutase OS=Mus musculus OX=10090 GN=Bpgm PE=1 SV=2 | 108 | 110.4 | 139.7 | 84.7 | 84.4 | 92.8 | 90.8 | 113.4 | 83.5 | 92.2 |
| 502 | Cytochrome P450 2A4 OS=Mus musculus OX=10090 GN=Cyp2a4 PE=2 SV=3 | 126.6 | 133.5 | 121.9 | 108.6 | 73.3 | 107.5 | 46.4 | 99.5 | 102.5 | 80 |
| 503 | Glutathione S-transferase Mu 2 OS=Mus musculus OX=10090 GN=Gstm2 PE=1 SV=2 | 89.7 | 72.8 | 107 | 119.1 | 158.1 | 90.8 | 53.8 | 121.8 | 110.9 | 75.9 |
| 504 | Histone H1.2 OS=Mus musculus OX=10090 GN=H1-2 PE=1 SV=2 | 49.8 | 54 | 43 | 129 | 21.4 | 151.4 | 113.3 | 47.9 | 159.8 | 230.4 |
| 505 | Carbonic anhydrase 3 OS=Mus musculus OX=10090 GN=Ca3 PE=1 SV=3 | 116.3 | 159.2 | 171.7 | 65.4 | 108.9 | 102.5 | 157.1 | 42.4 | 38.7 | 38 |
| 506 | Coagulation factor IX OS=Mus musculus OX=10090 GN=F9 PE=2 SV=3 | 95.1 | 85.3 | 117.3 | 119.1 | 68.5 | 90 | 66 | 106.5 | 144.8 | 107.4 |
| 507 | Phenylalanine-4-hydroxylase OS=Mus musculus OX=10090 GN=Pah PE=1 SV=4 | 108 | 93.5 | 98.9 | 116.4 | 76.6 | 118.2 | 52.2 | 124.4 | 119.8 | 92.1 |
| 508 | Methylmalonyl-CoA mutase, mitochondrial OS=Mus musculus OX=10090 GN=Mmut PE=1 SV=2 | 84.7 | 79.1 | 87 | 119.7 | 70.5 | 83.3 | 65.6 | 133.1 | 128.7 | 148.2 |
| 509 | Glutamyl aminopeptidase OS=Mus musculus OX=10090 GN=Enpep PE=1 SV=1 | 105.3 | 94.3 | 102.8 | 96.9 | 77.3 | 91.2 | 77.3 | 101.5 | 119.1 | 134.2 |
| 510 | Argininosuccinate synthase OS=Mus musculus OX=10090 GN=Ass1 PE=1 SV=1 | 102 | 93 | 91.5 | 132.5 | 58.4 | 108.9 | 65.5 | 129.1 | 128.4 | 90.5 |
| 511 | Lysosomal protective protein OS=Mus musculus OX=10090 GN=Ctsa PE=1 SV=1 | 91.7 | 93.5 | 89.8 | 78.3 | 140.9 | 104 | 126.5 | 94.4 | 86 | 95 |
| 512 | Lysosome-associated membrane glycoprotein 2 OS=Mus musculus OX=10090 GN=Lamp2 PE=1 SV=2 | 90.4 | 99.6 | 92 | 77.4 | 138.6 | 92.1 | 119.7 | 96.8 | 80.8 | 112.5 |
| 513 | Heat shock-related 70 kDa protein 2 OS=Mus musculus OX=10090 GN=Hspa2 PE=1 SV=2 | 124.9 | 110.9 | 116.5 | 86.5 | 102.9 | 111.2 | 85.8 | 93.4 | 88.9 | 79 |
| 514 | Alpha-enolase OS=Mus musculus OX=10090 GN=Eno1 PE=1 SV=3 | 122.8 | 106.5 | 105.2 | 114.3 | 97.1 | 106.1 | 62.9 | 99.9 | 96.4 | 88.8 |
| 515 | AP-2 complex subunit alpha-1 OS=Mus musculus OX=10090 GN=Ap2a1 PE=1 SV=1 | 46.5 | 111.9 | 106.7 | 100.5 | 91 | 111.7 | 109.7 | 108.5 | 101.1 | 112.2 |
| 516 | AP-2 complex subunit alpha-2 OS=Mus musculus OX=10090 GN=Ap2a2 PE=1 SV=2 | 162.2 | 104.8 | 141.1 | 96 | 47 | 110.6 | 67.2 | 90.7 | 89.5 | 91 |
| 517 | Lysosomal acid glucosylceramidase OS=Mus musculus OX=10090 GN=Gba PE=1 SV=1 | | 112.7 | 123.9 | 116.4 | 54.6 | 88.3 | 63.9 | 145.6 | 120.2 | 174.4 |
| 518 | Methanethiol oxidase OS=Mus musculus OX=10090 GN=Selenbp1 PE=1 SV=2 | 138.1 | 92.7 | 85.8 | 118.4 | 63.1 | 90.4 | 64.7 | 118.3 | 108.4 | 120.2 |
| 519 | Peptidyl-prolyl cis-trans isomerase A OS=Mus musculus OX=10090 GN=Ppia PE=1 SV=2 | 119.8 | 109.9 | 99.9 | 88.4 | 108 | 114.6 | 85.5 | 101.2 | 84 | 88.6 |
| 520 | Cathepsin D OS=Mus musculus OX=10090 GN=Ctsd PE=1 SV=1 | 115.3 | 106.9 | 104.9 | 76.3 | 137.2 | 130.5 | 84.8 | 77.8 | 87.4 | 79 |
| 521 | Basigin OS=Mus musculus OX=10090 GN=Bsg PE=1 SV=2 | 99 | 91.7 | 104.5 | 90.5 | 115.6 | 98.5 | 94.3 | 97.2 | 94.6 | 114.1 |
| 522 | Cofilin-1 OS=Mus musculus OX=10090 GN=Cfl1 PE=1 SV=3 | 106.1 | 108.6 | 108 | 99.9 | 74.8 | 120.7 | 74 | 128.6 | 83 | 96.4 |
| 523 | Glutathione S-transferase P 1 OS=Mus musculus OX=10090 GN=Gstp1 PE=1 SV=2 | 75.6 | 75 | 93 | 87.7 | 175 | 118.1 | 74.2 | 111.9 | 89.1 | 100.5 |

| | | | | | | | | | | | |
|---|---|---|---|---|---|---|---|---|---|---|---|
| 524 | Glutathione S-transferase Mu 3 OS=Mus musculus OX=10090 GN=Gstm3 PE=1 SV=2 | 47.9 | 53.1 | 67.6 | 125.5 | 191.3 | 124.2 | 61.6 | 129.8 | 130.7 | 68.4 |
| 525 | Cytochrome c oxidase subunit 4 isoform 1, mitochondrial OS=Mus musculus OX=10090 GN=Cox4i1 PE=1 SV=2 | 108.9 | 74.9 | 97.5 | 97.8 | 103.7 | 93.7 | 116 | 100 | 90.2 | 117.2 |
| 526 | Endoplasmic reticulum chaperone BiP OS=Mus musculus OX=10090 GN=Hspa5 PE=1 SV=3 | 118.8 | 117.9 | 114.8 | 73.7 | 125.5 | 128.3 | 80.2 | 85.2 | 74.2 | 81.5 |
| 527 | Beta-hexosaminidase subunit beta OS=Mus musculus OX=10090 GN=Hexb PE=1 SV=2 | 109.3 | 111.6 | 121.3 | 93.7 | 90.3 | 107.1 | 76.2 | 102.6 | 97.6 | 90.3 |
| 528 | Cytochrome P450 2A5 OS=Mus musculus OX=10090 GN=Cyp2a5 PE=2 SV=1 | 87.3 | 83 | 112.8 | 148.1 | 61.1 | 84.6 | 49.1 | 131.7 | 132.5 | 109.6 |
| 529 | Plasminogen OS=Mus musculus OX=10090 GN=Plg PE=1 SV=3 | 98.6 | 111.3 | 114.1 | 64.2 | 134.6 | 122.1 | 107.4 | 116.8 | 66 | 64.9 |
| 530 | Phosphatidylcholine translocator ABCB4 OS=Mus musculus OX=10090 GN=Abcb4 PE=1 SV=2 | 79.7 | 110.5 | 88 | 106.6 | 77 | 97.8 | 81.2 | 106 | 141.1 | 112 |
| 531 | Beta-enolase OS=Mus musculus OX=10090 GN=Eno3 PE=1 SV=3 | 56.5 | 69.5 | 54.5 | 53.5 | 228.8 | 59 | 323.5 | 49.5 | 42.8 | 62.3 |
| 532 | Ferrochelatase, mitochondrial OS=Mus musculus OX=10090 GN=Fech PE=1 SV=3 | 113.8 | 85.2 | 102.6 | 112.8 | 76.1 | 99.5 | 79.6 | 108.7 | 111.8 | 109.9 |
| 533 | Alpha-1-antitrypsin 1-2 OS=Mus musculus OX=10090 GN=Serpina1b PE=1 SV=2 | 74.3 | 94.7 | 139.7 | 102 | 78.6 | 82.5 | 60.4 | 92 | 102.5 | 173.4 |
| 534 | AP-1 complex subunit gamma-1 OS=Mus musculus OX=10090 GN=Ap1g1 PE=1 SV=3 | 129.9 | 104.4 | 114.2 | 93.6 | 62.9 | 120.7 | 91 | 102.6 | 91.8 | 89 |
| 535 | Eukaryotic translation initiation factor 3 subunit A OS=Mus musculus OX=10090 GN=Eif3a PE=1 SV=5 | 105.3 | 89.1 | 107.2 | 81.9 | 122.8 | 108.2 | 106.2 | 93.9 | 80.2 | 105.1 |
| 536 | Carbonic anhydrase 5A, mitochondrial OS=Mus musculus OX=10090 GN=Ca5a PE=1 SV=2 | 95.1 | 89.4 | 96.5 | 94.2 | 115.2 | 95.7 | 163 | 88.7 | 81.1 | 81 |
| 537 | Alpha-crystallin B chain OS=Mus musculus OX=10090 GN=Cryab PE=1 SV=2 | 45.2 | 63.6 | 49.1 | 50.2 | 222.1 | 41.7 | 378.1 | 43 | 37 | 70 |
| 538 | Carboxylesterase 1C OS=Mus musculus OX=10090 GN=Ces1c PE=1 SV=4 | | 202.8 | | 123.1 | | 167.5 | 150.7 | 114.4 | 115.8 | 125.8 |
| 539 | Peptidyl-prolyl cis-trans isomerase B OS=Mus musculus OX=10090 GN=Ppib PE=1 SV=2 | 128.4 | 117.4 | 113.4 | 88.4 | 111.7 | 141.4 | 64.2 | 84.3 | 87.1 | 63.7 |
| 540 | Cytochrome P450 2D10 OS=Mus musculus OX=10090 GN=Cyp2d10 PE=1 SV=2 | 98.9 | 100.1 | 92.4 | 93.1 | 141.7 | 79.8 | 147.4 | 84.8 | 92.9 | 69.1 |
| 541 | Glutathione S-transferase A4 OS=Mus musculus OX=10090 GN=Gsta4 PE=1 SV=3 | 98.7 | 78.9 | 99.1 | 144.7 | 93.3 | 87.6 | 59.3 | 137.9 | 125.7 | 74.8 |
| 542 | Leukotriene A-4 hydrolase OS=Mus musculus OX=10090 GN=Lta4h PE=1 SV=4 | 114 | 118.7 | 119.4 | 97.2 | 78.9 | 129.9 | 76.3 | 92.4 | 92.7 | 80.6 |
| 543 | Aldehyde dehydrogenase 1A1 OS=Mus musculus OX=10090 GN=Aldh1a1 PE=1 SV=5 | 109.6 | 110.7 | 130.6 | 114.7 | 82.9 | 107.6 | 64.3 | 100.9 | 105.3 | 73.4 |
| 544 | Moesin OS=Mus musculus OX=10090 GN=Msn PE=1 SV=3 | 113.3 | 92.7 | 106.6 | 93.5 | 111 | 132.5 | 87.1 | 74 | 96.6 | 92.5 |
| 545 | Catenin alpha-1 OS=Mus musculus OX=10090 GN=Ctnna1 PE=1 SV=1 | 124.4 | 132.2 | 128.8 | 85.9 | 82.7 | 119.8 | 86 | 80.7 | 85.4 | 74.2 |
| 546 | Glutamate dehydrogenase 1, mitochondrial OS=Mus musculus OX=10090 GN=Glud1 PE=1 SV=1 | 121.8 | 105.7 | 103.1 | 108.3 | 98.4 | 118.8 | 70.7 | 92.2 | 85.4 | 95.5 |
| 547 | Alpha-mannosidase 2 OS=Mus musculus OX=10090 GN=Man2a1 PE=1 SV=2 | 103.5 | 104.9 | 109.1 | 87.6 | 96.5 | 101.8 | 116 | 105.4 | 95.5 | 79.6 |
| 548 | Guanine nucleotide-binding protein subunit alpha-13 OS=Mus musculus OX=10090 GN=Gna13 PE=1 SV=1 | | 158.1 | | 132.4 | | 183.8 | 119 | 125.9 | 144.8 | 136.1 |
| 549 | Phospholipase A-2-activating protein OS=Mus musculus OX=10090 GN=Plaa PE=1 SV=4 | 124.7 | 98.4 | 102.6 | 80.3 | 103.8 | 162.3 | 64.2 | 100.3 | 84.2 | 79.2 |
| 550 | Histone H2AX OS=Mus musculus OX=10090 GN=H2ax PE=1 SV=2 | 116.8 | 98.1 | 117.2 | 79.1 | 116.9 | 133.6 | 79.2 | 79.8 | 89.9 | 89.3 |
| 551 | Cytoplasmic aconitate hydratase OS=Mus musculus OX=10090 GN=Aco1 PE=1 SV=3 | 78.3 | 138.9 | 65.2 | 133.7 | 47.6 | 138.3 | 85.3 | 96.4 | 114 | 102.2 |
| 552 | DNA-(apurinic or apyrimidinic site) endonuclease OS=Mus musculus OX=10090 GN=Apex1 PE=1 SV=2 | 120.6 | 87 | 94.8 | 96.9 | 75.6 | 96.7 | 94.1 | 116.2 | 94.7 | 123.5 |
| 553 | Alcohol dehydrogenase class-3 OS=Mus musculus OX=10090 GN=Adh5 PE=1 SV=3 | 133.1 | 108.1 | 121.8 | 110 | 72.9 | 102.8 | 66.7 | 107.3 | 90.2 | 87 |
| 554 | Adenylosuccinate synthetase isozyme 1 OS=Mus musculus OX=10090 GN=Adss1 PE=1 SV=2 | 86.1 | 90.3 | 84.4 | 80.7 | 167.7 | 87.5 | 137.6 | 98.6 | 77.7 | 89.4 |
| 555 | Biglycan OS=Mus musculus OX=10090 GN=Bgn PE=1 SV=1 | 105.5 | 81.9 | 103 | 96 | 94 | 99.9 | 124.6 | 98.6 | 91.6 | 104.9 |
| 556 | Murinoglobulin-1 OS=Mus musculus OX=10090 GN=Mug1 PE=1 SV=3 | 156.8 | 142.7 | 137.6 | 86.9 | 97.5 | 112.7 | 60.8 | 67.4 | 90.2 | 47.5 |
| 557 | Dipeptidyl peptidase 4 OS=Mus musculus OX=10090 GN=Dpp4 PE=1 SV=3 | 128.6 | 106.3 | 115.3 | 102.3 | 66.9 | 97.6 | 74.1 | 102 | 105.9 | 101.2 |
| 558 | Ferritin light chain 1 OS=Mus musculus OX=10090 GN=Ftl1 PE=1 SV=2 | 82.1 | 101.5 | 126.3 | 58.2 | 88.6 | 116.4 | 94.6 | 94.5 | 65.6 | 172.1 |
| 559 | Ornithine aminotransferase, mitochondrial OS=Mus musculus OX=10090 GN=Oat PE=1 SV=1 | 64.5 | 63.3 | 88.6 | 207 | 27.3 | 44.3 | 34.4 | 164.1 | 206.5 | 100 |

| | | | | | | | | | | | |
|---|---|---|---|---|---|---|---|---|---|---|---|
| 560 | Glutathione S-transferase A3 OS=Mus musculus OX=10090 GN=Gsta3 PE=1 SV=2 | 101.7 | 141.9 | 91.7 | 114.5 | 39.2 | 112.3 | 78.6 | 123.8 | 106 | 90.3 |
| 561 | Peptidyl-prolyl cis-trans isomerase FKBP4 OS=Mus musculus OX=10090 GN=Fkbp4 PE=1 SV=5 | 134.7 | 114.8 | 120.6 | 82.5 | 101.8 | 119 | 93.1 | 83.8 | 74.8 | 74.8 |
| 562 | Desmin OS=Mus musculus OX=10090 GN=Des PE=1 SV=3 | 23.6 | 91.9 | 48.9 | 45.8 | 240.1 | 52.6 | 354.9 | 45 | 31.9 | 65.2 |
| 563 | Lupus La protein homolog OS=Mus musculus OX=10090 GN=Ssb PE=1 SV=1 | 96.3 | 76.1 | 83.3 | 72.1 | 57.6 | 97.4 | 129.1 | 93.7 | 118.5 | 175.8 |
| 564 | Cytochrome P450 2F2 OS=Mus musculus OX=10090 GN=Cyp2f2 PE=1 SV=1 | 134.4 | 121.4 | 154.5 | 81.7 | 72.9 | 97.2 | 61.3 | 83.3 | 110.8 | 82.5 |
| 565 | Macrophage migration inhibitory factor OS=Mus musculus OX=10090 GN=Mif PE=1 SV=2 | 108.5 | 86 | 73.9 | 110 | 90.6 | 98.8 | 59 | 177 | 72.1 | 124 |
| 566 | Bifunctional epoxide hydrolase 2 OS=Mus musculus OX=10090 GN=Ephx2 PE=1 SV=2 | 135.3 | 112.6 | 131.7 | 110.7 | 67.8 | 113.5 | 45.5 | 99.9 | 110.1 | 73.1 |
| 567 | Asialoglycoprotein receptor 1 OS=Mus musculus OX=10090 GN=Asgr1 PE=1 SV=4 | 112.5 | 110.3 | 101.7 | 90.5 | 93.1 | 107.2 | 91.5 | 103.4 | 88.4 | 101.5 |
| 568 | Apolipoprotein C-I OS=Mus musculus OX=10090 GN=Apoc1 PE=1 SV=1 | | 157.9 | 80.7 | 97.5 | 111 | 155.5 | 75.1 | 101.7 | 88.5 | 131.9 |
| 569 | Histidine ammonia-lyase OS=Mus musculus OX=10090 GN=Hal PE=1 SV=1 | 98.3 | 91.6 | 107.3 | 137.7 | 70.4 | 101 | 69.6 | 115.5 | 130 | 78.6 |
| 570 | Fumarylacetoacetase OS=Mus musculus OX=10090 GN=Fah PE=1 SV=2 | 113.9 | 112.1 | 95.4 | 119.8 | 60.6 | 143.3 | 85.6 | 70.1 | 107.3 | 91.9 |
| 571 | Calnexin OS=Mus musculus OX=10090 GN=Canx PE=1 SV=1 | 143 | 137.5 | 134.2 | 86.2 | 79.7 | 123 | 74.9 | 82.7 | 81.8 | 57.1 |
| 572 | Glucose-6-phosphatase catalytic subunit 1 OS=Mus musculus OX=10090 GN=G6pc1 PE=1 SV=2 | 129.2 | 125.9 | 100.2 | 102.7 | 75.5 | 90 | 47.8 | 115.9 | 108.5 | 104.2 |
| 573 | AP-1 complex subunit mu-1 OS=Mus musculus OX=10090 GN=Ap1m1 PE=1 SV=3 | 118.8 | 102.9 | 114.1 | 94.7 | 73 | 112.6 | 83.8 | 107.9 | 87.9 | 104.2 |
| 574 | Peroxiredoxin-1 OS=Mus musculus OX=10090 GN=Prdx1 PE=1 SV=1 | 136.1 | 116.8 | 117.6 | 98.4 | 87.6 | 123.3 | 87.8 | 87.6 | 82.4 | 62.4 |
| 575 | 60S ribosomal protein L12 OS=Mus musculus OX=10090 GN=Rpl12 PE=1 SV=2 | 72 | 77.4 | 78.1 | 120.3 | 78.4 | 157.7 | 116 | 61.1 | 119.8 | 119.4 |
| 576 | Oxygen-dependent coproporphyrinogen-III oxidase, mitochondrial OS=Mus musculus OX=10090 GN=Cpox PE=1 SV=2 | 99.1 | 101.2 | 127.3 | 135.2 | 62.5 | 110.4 | 51.3 | 105.3 | 124.1 | 83.8 |
| 577 | NADPH--cytochrome P450 reductase OS=Mus musculus OX=10090 GN=Por PE=1 SV=2 | 88.2 | 85.5 | 90.4 | 109.8 | 66.5 | 101.7 | 95.6 | 147.8 | 106.7 | 107.7 |
| 578 | Hydroxymethylglutaryl-CoA lyase, mitochondrial OS=Mus musculus OX=10090 GN=Hmgcl PE=1 SV=2 | 94 | 93.3 | 84.6 | 97.8 | 106.2 | 110.8 | 79.4 | 119 | 92.9 | 122.1 |
| 579 | Adenylyl cyclase-associated protein 1 OS=Mus musculus OX=10090 GN=Cap1 PE=1 SV=4 | 106 | 107.3 | 110.4 | 88.9 | 103.9 | 124 | 93.4 | 89.6 | 88.9 | 87.7 |
| 580 | Indolethylamine N-methyltransferase OS=Mus musculus OX=10090 GN=Inmt PE=1 SV=1 | 160.7 | 123.2 | 132.7 | 86.7 | 79.4 | 153.4 | 48 | 73.6 | 63 | 79.3 |
| 581 | 60S ribosomal protein L28 OS=Mus musculus OX=10090 GN=Rpl28 PE=1 SV=2 | 145.7 | 125.3 | 129 | 92.3 | 89.3 | 108.1 | 66.7 | 109.2 | 80.9 | 53.4 |
| 582 | Long-chain-fatty-acid--CoA ligase 1 OS=Mus musculus OX=10090 GN=Acsl1 PE=1 SV=2 | 123 | 115.2 | 111.2 | 97.3 | 96.7 | 114.5 | 119.1 | 83.2 | 81.6 | 58.2 |
| 583 | Mannose-binding protein C OS=Mus musculus OX=10090 GN=Mbl2 PE=1 SV=2 | 120.7 | 120.2 | 141 | 93.2 | 94.3 | 101.4 | 70.6 | 82.4 | 90.2 | 86 |
| 584 | Histone H1.4 OS=Mus musculus OX=10090 GN=H1-4 PE=1 SV=2 | 118.2 | 159.8 | 121.4 | 69.2 | 83.2 | 85.5 | 82.2 | 93.8 | 80 | 106.6 |
| 585 | Histone H1.1 OS=Mus musculus OX=10090 GN=H1-1 PE=1 SV=2 | 118.6 | 140.3 | 107.7 | 77.1 | 60.5 | 82.1 | 82.6 | 110.1 | 90.2 | 130.8 |
| 586 | Histone H1.5 OS=Mus musculus OX=10090 GN=H1-5 PE=1 SV=2 | 122.8 | 169.1 | 138.2 | 69.3 | 59 | 72.9 | 71.5 | 91.2 | 87.3 | 118.6 |
| 587 | Histone H1.3 OS=Mus musculus OX=10090 GN=H1-3 PE=1 SV=2 | 96.4 | 130.4 | 99 | 86 | 67.9 | 106.3 | 91.5 | 76.5 | 103.4 | 142.5 |
| 588 | Medium-chain specific acyl-CoA dehydrogenase, mitochondrial OS=Mus musculus OX=10090 GN=Acadm PE=1 SV=1 | 88.9 | 78.5 | 99.6 | 100.6 | 127.7 | 100.3 | 101.3 | 113.9 | 95 | 94.3 |
| 589 | Glutathione peroxidase 3 OS=Mus musculus OX=10090 GN=Gpx3 PE=1 SV=2 | 57.9 | 57.4 | 76.8 | 102 | 106.7 | 68.8 | 128.1 | 80.7 | 152.5 | 169.2 |
| 590 | E3 ubiquitin-protein ligase NEDD4 OS=Mus musculus OX=10090 GN=Nedd4 PE=1 SV=3 | 110.6 | 104.1 | 103.2 | 99 | 83.5 | 100.9 | 90.4 | 104.8 | 95.7 | 107.9 |
| 591 | Dolichyl-diphosphooligosaccharide--protein glycosyltransferase subunit STT3A OS=Mus musculus OX=10090 GN=Stt3a PE=1 SV=1 | 129.6 | 120.6 | 124.2 | 71.2 | 128.7 | 110.9 | 78 | 84.9 | 69.6 | 82.3 |
| 592 | Aldehyde dehydrogenase, mitochondrial OS=Mus musculus OX=10090 GN=Aldh2 PE=1 SV=1 | 90.4 | 90.4 | 106.5 | 123.9 | 63.1 | 103.1 | 56.4 | 102.5 | 126.2 | 137.4 |
| 593 | F-actin-capping protein subunit alpha-2 OS=Mus musculus OX=10090 GN=Capza2 PE=1 SV=3 | 115.2 | 101.8 | 113.8 | 90.1 | 111.7 | 119.9 | 106.8 | 81.7 | 78.1 | 80.9 |
| 594 | Glutathione reductase, mitochondrial OS=Mus musculus OX=10090 GN=Gsr PE=1 SV=3 | 111.1 | 110.2 | 107.5 | 97.4 | 100.5 | 107.2 | 54.6 | 110.6 | 100.3 | 100.8 |

| | | | | | | | | | | | |
|---|---|---|---|---|---|---|---|---|---|---|---|
| 595 | Glutamine--fructose-6-phosphate aminotransferase [isomerizing] 1 OS=Mus musculus OX=10090 GN=Gfpt1 PE=1 SV=3 | 101.1 | 83.3 | 80.2 | 65.8 | 129.5 | 103.2 | 176.4 | 86.4 | 82.8 | 91.4 |
| 596 | 60S ribosomal protein L6 OS=Mus musculus OX=10090 GN=Rpl6 PE=1 SV=3 | 109.8 | 117.5 | 110.8 | 89.5 | 77.5 | 102.7 | 74.3 | 116.8 | 104.7 | 96.4 |
| 597 | Carnitine O-acetyltransferase OS=Mus musculus OX=10090 GN=Crat PE=1 SV=3 | 71 | 80.7 | 96.3 | 86.6 | 128.7 | 88.8 | 195.4 | 86.9 | 82.8 | 82.8 |
| 598 | Crk-like protein OS=Mus musculus OX=10090 GN=Crkl PE=1 SV=2 | | 138.9 | | 122.6 | | 142.1 | 107.3 | 179.5 | 109.6 | 200 |
| 599 | 60S ribosomal protein L5 OS=Mus musculus OX=10090 GN=Rpl5 PE=1 SV=3 | 77.3 | 74.7 | 74.2 | 58.5 | 81.3 | 85.8 | 202.6 | 57.6 | 138.5 | 149.5 |
| 600 | 60S ribosomal protein L13 OS=Mus musculus OX=10090 GN=Rpl13 PE=1 SV=3 | 110 | 109.5 | 111.5 | 93.2 | 105.6 | 105.8 | 93.3 | 95.7 | 85.6 | 89.8 |
| 601 | Annexin A5 OS=Mus musculus OX=10090 GN=Anxa5 PE=1 SV=1 | 124.7 | 98.2 | 121.6 | 105.9 | 108.7 | 139.6 | 74.3 | 75.1 | 85.9 | 66 |
| 602 | Prelamin-A/C OS=Mus musculus OX=10090 GN=Lmna PE=1 SV=2 | 87.4 | 87.7 | 99.5 | 81 | 139.4 | 113.5 | 110 | 98.5 | 75.5 | 107.3 |
| 603 | Heat shock 70 kDa protein 4L OS=Mus musculus OX=10090 GN=Hspa4l PE=1 SV=2 | 126 | 115.6 | 162.6 | 89.9 | 74.9 | 123.7 | 64.9 | 96.9 | 74.7 | 70.8 |
| 604 | Cytochrome c oxidase subunit 7A2, mitochondrial OS=Mus musculus OX=10090 GN=Cox7a2 PE=1 SV=2 | 88.4 | 86.1 | 94.1 | 106.6 | 106.1 | 89.8 | 101.7 | 119 | 102.4 | 105.8 |
| 605 | Glutathione S-transferase Mu 5 OS=Mus musculus OX=10090 GN=Gstm5 PE=1 SV=1 | 116.1 | 108.3 | 106.7 | 102.8 | 84.6 | 100.8 | 80.7 | 107.8 | 92.4 | 99.8 |
| 606 | ADP/ATP translocase 1 OS=Mus musculus OX=10090 GN=Slc25a4 PE=1 SV=4 | 25.6 | 46.5 | 26.6 | 24.8 | 248.3 | 30.4 | 536.7 | 15.6 | 13.6 | 31.8 |
| 607 | Heterogeneous nuclear ribonucleoprotein A1 OS=Mus musculus OX=10090 GN=Hnrnpa1 PE=1 SV=2 | | 128.2 | | 123.6 | | 159.6 | 118.5 | 144 | 130.3 | 195.9 |
| 608 | 4-hydroxyphenylpyruvate dioxygenase OS=Mus musculus OX=10090 GN=Hpd PE=1 SV=3 | 114.6 | 99.8 | 102 | 119.1 | 85.4 | 105.8 | 70 | 112.7 | 99.9 | 90.7 |
| 609 | Inositol polyphosphate 1-phosphatase OS=Mus musculus OX=10090 GN=Inpp1 PE=1 SV=2 | 111.5 | 112.5 | 108.2 | 88.8 | 99.9 | 113 | 94 | 103.6 | 81.9 | 86.5 |
| 610 | Pro-cathepsin H OS=Mus musculus OX=10090 GN=Ctsh PE=1 SV=2 | 117.3 | 120 | 97.1 | 71.2 | 118.8 | 144.8 | 71.5 | 82.1 | 82.8 | 94.3 |
| 611 | Adenosylhomocysteinase OS=Mus musculus OX=10090 GN=Ahcy PE=1 SV=3 | 103.1 | 91.9 | 90.4 | 117.1 | 63.9 | 93.3 | 67.1 | 134.6 | 120.3 | 118.2 |
| 612 | Dimethylaniline monooxygenase [N-oxide-forming] 1 OS=Mus musculus OX=10090 GN=Fmo1 PE=1 SV=1 | 127.6 | 117 | 145.4 | 129.1 | 60.9 | 94.4 | 68.8 | 83.7 | 113.3 | 59.8 |
| 613 | Arylamine N-acetyltransferase 2 OS=Mus musculus OX=10090 GN=Nat2 PE=1 SV=1 | 130.3 | 104.7 | 125.8 | 100.8 | 78.1 | 114.8 | 87.9 | 85.7 | 89.2 | 82.8 |
| 614 | Arylsulfatase B OS=Mus musculus OX=10090 GN=Arsb PE=1 SV=3 | 115.3 | 105.8 | 101.5 | 72.1 | 50.6 | 83.4 | 101.8 | 86.4 | 149 | 133.9 |
| 615 | Proliferation-associated protein 2G4 OS=Mus musculus OX=10090 GN=Pa2g4 PE=1 SV=3 | 60.7 | 130.1 | 69.8 | 124.8 | 48.5 | 141.1 | 95 | 106.4 | 118.1 | 105.4 |
| 616 | Long-chain specific acyl-CoA dehydrogenase, mitochondrial OS=Mus musculus OX=10090 GN=Acadl PE=1 SV=2 | 92.3 | 87.1 | 105.9 | 107.1 | 102.8 | 101.3 | 118.4 | 110.1 | 99.2 | 75.8 |
| 617 | Estradiol 17-beta-dehydrogenase 2 OS=Mus musculus OX=10090 GN=Hsd17b2 PE=1 SV=2 | 150.4 | 175.9 | 150.5 | 66.5 | 80.9 | 141 | 69.5 | 56.2 | 64.4 | 44.5 |
| 618 | Peroxisomal multifunctional enzyme type 2 OS=Mus musculus OX=10090 GN=Hsd17b4 PE=1 SV=3 | 136.9 | 106 | 121 | 100 | 68.6 | 107 | 59.4 | 108.2 | 104.2 | 88.6 |
| 619 | Glutathione synthetase OS=Mus musculus OX=10090 GN=Gss PE=1 SV=1 | 98.6 | 103.2 | 101.1 | 98.7 | 118.4 | 85.9 | 98.5 | 101.9 | 89.6 | 104.1 |
| 620 | Hepatoma-derived growth factor OS=Mus musculus OX=10090 GN=Hdgf PE=1 SV=2 | 96.4 | 97.7 | 89.8 | 77.1 | 111.8 | 105.4 | 115.4 | 106 | 80 | 120.4 |
| 621 | ADP/ATP translocase 2 OS=Mus musculus OX=10090 GN=Slc25a5 PE=1 SV=3 | 94.8 | 93.5 | 86.4 | 100.6 | 103 | 87.2 | 118.5 | 111.7 | 103.3 | 100.9 |
| 622 | Lumican OS=Mus musculus OX=10090 GN=Lum PE=1 SV=2 | 71 | 88.7 | 82.4 | 74.3 | 170.8 | 51.9 | 214.1 | 62.7 | 61.4 | 122.8 |
| 623 | Hexokinase-4 OS=Mus musculus OX=10090 GN=Gck PE=1 SV=1 | 187.4 | 240.3 | 188.2 | 53.1 | 35.8 | 95.7 | 65.9 | 38.9 | 52.9 | 41.8 |
| 624 | Carnitine O-palmitoyltransferase 2, mitochondrial OS=Mus musculus OX=10090 GN=Cpt2 PE=1 SV=2 | 95.5 | 86.7 | 96.5 | 107.9 | 91 | 100.2 | 97.7 | 109 | 111.3 | 104.3 |
| 625 | Lipoamide acyltransferase component of branched-chain alpha-keto acid dehydrogenase complex, mitochondrial OS=Mus musculus OX=10090 GN=Dbt PE=1 SV=2 | 94.9 | 85.3 | 90.2 | 115.5 | 76.1 | 97.1 | 67.3 | 131.8 | 128.6 | 113.3 |
| 626 | Monocarboxylate transporter 1 OS=Mus musculus OX=10090 GN=Slc16a1 PE=1 SV=1 | | 159 | | 142.8 | | 159.1 | 95 | 138.6 | 144.3 | 161.3 |
| 627 | CCHC-type zinc finger nucleic acid binding protein OS=Mus musculus OX=10090 GN=Cnbp PE=1 SV=2 | 76.6 | 63.2 | 102.4 | 44.6 | 123.7 | 164.1 | 93 | 146.8 | 50.7 | 134.9 |
| 628 | Hydroxymethylglutaryl-CoA synthase, mitochondrial OS=Mus musculus OX=10090 GN=Hmgcs2 PE=1 SV=2 | 119.4 | 96.8 | 101.6 | 110 | 69.1 | 111.7 | 51.9 | 113.6 | 117.2 | 108.9 |
| 629 | Fatty acid-binding protein, intestinal OS=Mus musculus OX=10090 GN=Fabp2 PE=1 SV=2 | 129.3 | 171.6 | 115.3 | 43.6 | 183.5 | 140.1 | 76 | 51.7 | 36.6 | 52.3 |

| | | | | | | | | | | | |
|---|---|---|---|---|---|---|---|---|---|---|---|
| 630 | Adenosine kinase OS=Mus musculus OX=10090 GN=Adk PE=1 SV=2 | 128.7 | 119.2 | 112.5 | 100 | 77.2 | 105.3 | 81.1 | 93.8 | 89.3 | 92.9 |
| 631 | Alpha-2-macroglobulin receptor-associated protein OS=Mus musculus OX=10090 GN=Lrpap1 PE=1 SV=1 | 102.4 | 110.7 | 97.4 | 71.2 | 177 | 112.3 | 90.5 | 81.1 | 66.6 | 90.7 |
| 632 | ATP synthase subunit f, mitochondrial OS=Mus musculus OX=10090 GN=Atp5mf PE=1 SV=3 | 99.5 | 83.8 | 86 | 120.8 | 90.7 | 98.5 | 110.7 | 104 | 112.7 | 93.4 |
| 633 | FAD-linked sulfhydryl oxidase ALR OS=Mus musculus OX=10090 GN=Gfer PE=1 SV=2 | 121.9 | 108.2 | 116.2 | 111.2 | 72.9 | 106.1 | 60.9 | 106.5 | 94.3 | 101.8 |
| 634 | ATP synthase subunit ATP5MJ, mitochondrial OS=Mus musculus OX=10090 GN=Atp5mj PE=1 SV=1 | 99.2 | 86.3 | 86.8 | 125.2 | 92.9 | 76.5 | 92.5 | 122.1 | 116.2 | 102.3 |
| 635 | ATP synthase subunit epsilon, mitochondrial OS=Mus musculus OX=10090 GN=Atp5f1e PE=1 SV=2 | 101.4 | 86.2 | 101.1 | 111.7 | 109.9 | 84.4 | 115.5 | 105.4 | 95.8 | 88.6 |
| 636 | ATP synthase subunit beta, mitochondrial OS=Mus musculus OX=10090 GN=Atp5f1b PE=1 SV=2 | 94.8 | 97.1 | 98.2 | 110 | 108 | 94.6 | 102 | 98.2 | 95.5 | 101.5 |
| 637 | ADP-ribosyl cyclase/cyclic ADP-ribose hydrolase 1 OS=Mus musculus OX=10090 GN=Cd38 PE=1 SV=2 | 144.2 | 106.9 | 98.5 | 96.2 | 62.8 | 94.1 | 66.5 | 132.7 | 105.3 | 92.9 |
| 638 | Cytochrome P450 2A12 OS=Mus musculus OX=10090 GN=Cyp2a12 PE=1 SV=2 | 131.3 | 104.1 | 147 | 102.6 | 78.8 | 91.3 | 59 | 103.6 | 108.3 | 74 |
| 639 | Cytochrome P450 2C37 OS=Mus musculus OX=10090 GN=Cyp2c37 PE=1 SV=2 | 117.4 | 106.9 | 100.5 | 131.7 | 71.1 | 67.1 | 42.1 | 122.5 | 166.1 | 74.6 |
| 640 | Nicastrin OS=Mus musculus OX=10090 GN=Ncstn PE=1 SV=3 | 104.5 | 88.6 | 105.5 | 99.1 | 97.8 | 101.3 | 100.9 | 93.5 | 99.1 | 109.8 |
| 641 | Endoplasmic reticulum resident protein 29 OS=Mus musculus OX=10090 GN=Erp29 PE=1 SV=2 | 122.3 | 118.9 | 138.2 | 84.2 | 117.9 | 139.6 | 79.4 | 69.1 | 73.3 | 57.1 |
| 642 | Elongation factor 2 OS=Mus musculus OX=10090 GN=Eef2 PE=1 SV=2 | 110.2 | 109.7 | 113 | 90.6 | 101.9 | 118.6 | 86.4 | 109 | 72.7 | 87.8 |
| 643 | L-gulonolactone oxidase OS=Mus musculus OX=10090 GN=Gulo PE=1 SV=3 | 135.2 | 122.1 | 117.7 | 94.9 | 99.4 | 109.9 | 87.5 | 83.3 | 83.2 | 66.8 |
| 644 | Eukaryotic translation initiation factor 3 subunit E OS=Mus musculus OX=10090 GN=Eif3e PE=1 SV=1 | 124.3 | 107.9 | 117.2 | 96.5 | 80.3 | 112.9 | 74.7 | 99.7 | 96.8 | 89.6 |
| 645 | Poly(rC)-binding protein 1 OS=Mus musculus OX=10090 GN=Pcbp1 PE=1 SV=1 | 119.3 | 103.4 | 109 | 84 | 99.6 | 118 | 67.7 | 109.3 | 91.4 | 98.4 |
| 646 | Gamma-aminobutyric acid receptor-associated protein-like 2 OS=Mus musculus OX=10090 GN=Gabarapl2 PE=1 SV=1 | 139.8 | 124.7 | 114.8 | 95.9 | 113.1 | 122.2 | 64.2 | 89 | 72.9 | 63.3 |
| 647 | Eukaryotic initiation factor 4A-I OS=Mus musculus OX=10090 GN=Eif4a1 PE=1 SV=1 | 121.4 | 103.9 | 111 | 85.4 | 90.7 | 112.4 | 105.2 | 101.6 | 83.7 | 84.8 |
| 648 | 40S ribosomal protein S20 OS=Mus musculus OX=10090 GN=Rps20 PE=1 SV=1 | 114.1 | 99.2 | 105.7 | 99.5 | 86.9 | 118.5 | 57.9 | 123.8 | 102.9 | 91.4 |
| 649 | NEDD8-conjugating enzyme Ubc12 OS=Mus musculus OX=10090 GN=Ube2m PE=1 SV=1 | 149.8 | 104.6 | 106.1 | 105.9 | 101.4 | 106.3 | 37.9 | 129.6 | 95.3 | 63.2 |
| 650 | ADP-ribosylation factor 3 OS=Mus musculus OX=10090 GN=Arf3 PE=2 SV=2 | 95.5 | 87.6 | 114.4 | 70.3 | 67.8 | 101.2 | 123.6 | 56.6 | 158.4 | 124.6 |
| 651 | ATP-binding cassette sub-family E member 1 OS=Mus musculus OX=10090 GN=Abce1 PE=1 SV=1 | 135 | 105.3 | 100.1 | 92.8 | 92.2 | 116.3 | 76.2 | 101.2 | 86.3 | 94.7 |
| 652 | 60S ribosomal protein L27 OS=Mus musculus OX=10090 GN=Rpl27 PE=1 SV=2 | 59.5 | 143.7 | 96.6 | 95.5 | 126.4 | 119.3 | 78.7 | 104.9 | 96.2 | 79.2 |
| 653 | 60S ribosomal protein L37a OS=Mus musculus OX=10090 GN=Rpl37a PE=1 SV=2 | 124.5 | 121.5 | 110.9 | 87.5 | 122.1 | 118.1 | 64.4 | 89.5 | 81.7 | 79.8 |
| 654 | Dolichyl-diphosphooligosaccharide--protein glycosyltransferase subunit DAD1 OS=Mus musculus OX=10090 GN=Dad1 PE=1 SV=3 | 120.9 | 117.2 | 124.8 | 82.4 | 97 | 118.2 | 94.2 | 91 | 77.5 | 76.9 |
| 655 | 4-aminobutyrate aminotransferase, mitochondrial OS=Mus musculus OX=10090 GN=Abat PE=1 SV=1 | 41.9 | 75.7 | 32.4 | 185.5 | 28.7 | 104.9 | 72.1 | 130.7 | 180.6 | 147.5 |
| 656 | Nuclear transport factor 2 OS=Mus musculus OX=10090 GN=Nutf2 PE=1 SV=1 | 108.9 | 97.5 | 105.2 | 91.1 | 107.1 | 106.3 | 109.6 | 95.2 | 88.3 | 90.9 |
| 657 | 40S ribosomal protein S7 OS=Mus musculus OX=10090 GN=Rps7 PE=2 SV=1 | 90.4 | 106.7 | 137.6 | 92.3 | 91.7 | 126 | 90.2 | 85.7 | 98.9 | 80.4 |
| 658 | 40S ribosomal protein S8 OS=Mus musculus OX=10090 GN=Rps8 PE=1 SV=2 | 104.1 | 104.8 | 93.6 | 76.6 | 116.9 | 119.9 | 78.5 | 117.6 | 88.8 | 99.3 |
| 659 | 40S ribosomal protein S15a OS=Mus musculus OX=10090 GN=Rps15a PE=1 SV=2 | 134 | 103.2 | 114.6 | 94.5 | 93.1 | 118.8 | 69.8 | 99.8 | 92 | 80.1 |
| 660 | 40S ribosomal protein S14 OS=Mus musculus OX=10090 GN=Rps14 PE=1 SV=3 | 84.3 | 70.6 | 79.8 | 62.7 | 75.8 | 85.5 | 117.6 | 60.9 | 189.5 | 173.2 |
| 661 | 40S ribosomal protein S23 OS=Mus musculus OX=10090 GN=Rps23 PE=1 SV=3 | 72.3 | 116.3 | 68.1 | 81.9 | 123.6 | 111 | 142.3 | 115.8 | 80 | 88.7 |
| 662 | Elongation factor 1-alpha 2 OS=Mus musculus OX=10090 GN=Eef1a2 PE=1 SV=1 | 134.4 | 116.5 | 123.2 | 89.7 | 102.5 | 124.4 | 78.5 | 89.9 | 77.6 | 63.4 |
| 663 | 40S ribosomal protein S4, X isoform OS=Mus musculus OX=10090 GN=Rps4x PE=1 SV=2 | 128.3 | 104.5 | 118 | 91 | 96 | 113 | 89.9 | 92.5 | 85.5 | 81.3 |
| 664 | 60S ribosomal protein L23a OS=Mus musculus OX=10090 GN=Rpl23a PE=1 SV=1 | 106 | 92.9 | 98 | 83.6 | 111.5 | 128.2 | 70.6 | 130.3 | 83.3 | 95.6 |

| | | | | | | | | | | | |
|---|---|---|---|---|---|---|---|---|---|---|---|
| 665 | 40S ribosomal protein S6 OS=Mus musculus OX=10090 GN=Rps6 PE=1 SV=1 | 187.4 | 103.1 | 120.1 | 94.5 | 111.4 | 122 | 62.7 | 68.2 | 69.1 | 61.3 |
| 666 | Myotrophin OS=Mus musculus OX=10090 GN=Mtpn PE=1 SV=2 | 96.7 | 124.3 | 96.7 | 92.9 | 102 | 91.7 | 129.2 | 84.9 | 81 | 100.5 |
| 667 | Histone H4 OS=Mus musculus OX=10090 GN=H4c1 PE=1 SV=2 | 79.1 | 129.8 | 99.8 | 64.4 | 193.7 | 65.5 | 73.2 | 90.9 | 56.9 | 146.6 |
| 668 | 60S ribosomal protein L23 OS=Mus musculus OX=10090 GN=Rpl23 PE=1 SV=1 | 133.8 | 101.7 | 137.5 | 104.9 | 67.4 | 132.8 | 36.7 | 117.9 | 95.3 | 71.9 |
| 669 | 40S ribosomal protein S24 OS=Mus musculus OX=10090 GN=Rps24 PE=1 SV=1 | 104.8 | 109.9 | 109.7 | 85.9 | 124.6 | 135.8 | 62.2 | 97.5 | 86 | 83.7 |
| 670 | 40S ribosomal protein S26 OS=Mus musculus OX=10090 GN=Rps26 PE=1 SV=3 | 117.2 | 105.1 | 119.1 | 90.4 | 97.9 | 132.2 | 59.8 | 106.1 | 87.9 | 84.2 |
| 671 | Elongin-B OS=Mus musculus OX=10090 GN=Elob PE=1 SV=1 | 99.4 | 93.7 | 99.9 | 95.9 | 109.8 | 110 | 125.5 | 91.4 | 86.9 | 87.4 |
| 672 | 60S ribosomal protein L30 OS=Mus musculus OX=10090 GN=Rpl30 PE=1 SV=2 | 134.7 | 106.9 | 112 | 86.4 | 100 | 123 | 75.7 | 102 | 79.6 | 79.5 |
| 673 | Cytochrome c, somatic OS=Mus musculus OX=10090 GN=Cycs PE=1 SV=2 | 40.6 | 78.7 | 62.3 | 86.8 | 198.7 | 79 | 100.3 | 132.5 | 80 | 141.2 |
| 674 | 60S ribosomal protein L32 OS=Mus musculus OX=10090 GN=Rpl32 PE=1 SV=2 | 145 | 119.1 | 125.8 | 90.3 | 74.4 | 123.8 | 67.7 | 93.4 | 82.1 | 78.2 |
| 675 | 60S ribosomal protein L8 OS=Mus musculus OX=10090 GN=Rpl8 PE=1 SV=2 | 105.8 | 145.5 | 111.7 | 81.2 | 94.6 | 131.7 | 65.4 | 104.2 | 88.5 | 71.4 |
| 676 | Profilin-1 OS=Mus musculus OX=10090 GN=Pfn1 PE=1 SV=2 | 145.8 | 120.3 | 108.5 | 93.3 | 79.8 | 144 | 63.7 | 98.5 | 72.8 | 73.3 |
| 677 | Platelet-activating factor acetylhydrolase IB subunit beta OS=Mus musculus OX=10090 GN=Pafah1b1 PE=1 SV=2 | 118.2 | 98.6 | 115.1 | 106.5 | 100.9 | 114.9 | 75 | 94.2 | 88.7 | 88 |
| 678 | 60 kDa heat shock protein, mitochondrial OS=Mus musculus OX=10090 GN=Hspd1 PE=1 SV=1 | 108.8 | 104 | 106.4 | 103 | 97.2 | 101.8 | 84.6 | 104 | 101.1 | 88.9 |
| 679 | Prohibitin 1 OS=Mus musculus OX=10090 GN=Phb1 PE=1 SV=1 | 100.8 | 100.7 | 96.1 | 112.8 | 80.1 | 114.2 | 73.1 | 110 | 105.8 | 106.5 |
| 680 | 60S ribosomal protein L22 OS=Mus musculus OX=10090 GN=Rpl22 PE=1 SV=2 | 119.1 | 108 | 110.6 | 84.4 | 96.8 | 127.7 | 67.4 | 109.4 | 78.6 | 98 |
| 681 | Actin, alpha skeletal muscle OS=Mus musculus OX=10090 GN=Acta1 PE=1 SV=1 | 59.4 | 61.2 | 54 | 51.1 | 192.1 | 63.6 | 354.4 | 47.4 | 55.8 | 61 |
| 682 | Histone H3.1 OS=Mus musculus OX=10090 GN=H3c1 PE=1 SV=2 | 80.6 | 131.9 | 107.1 | 62.3 | 197.6 | 59.8 | 73.3 | 88.3 | 52.8 | 146.3 |
| 683 | Importin subunit beta-1 OS=Mus musculus OX=10090 GN=Kpnb1 PE=1 SV=2 | 118 | 101.9 | 115.6 | 95.5 | 97.9 | 111.6 | 83.6 | 99 | 90 | 87.1 |
| 684 | 3-beta-hydroxysteroid-Delta(8),Delta(7)-isomerase OS=Mus musculus OX=10090 GN=Ebp PE=1 SV=3 | 138 | 120.1 | 118.1 | 85.9 | 80.7 | 106.2 | 97.3 | 101.8 | 81.8 | 69.9 |
| 685 | ELAV-like protein 1 OS=Mus musculus OX=10090 GN=Elavl1 PE=1 SV=2 | 114.5 | 103.2 | 111.3 | 89 | 100.3 | 114.8 | 88.9 | 89.2 | 89.7 | 99.2 |
| 686 | Na(+)/H(+) exchange regulatory cofactor NHE-RF1 OS=Mus musculus OX=10090 GN=Slc9a3r1 PE=1 SV=3 | 76.6 | 95.6 | 83.2 | 77.9 | 124.2 | 106.9 | 54.7 | 132.9 | 80.2 | 167.8 |
| 687 | Caspase-3 OS=Mus musculus OX=10090 GN=Casp3 PE=1 SV=1 | 175.8 | 112 | 118.5 | 87.2 | 90.7 | 131 | 59.9 | 84.7 | 66.5 | 73.8 |
| 688 | PHD finger-like domain-containing protein 5A OS=Mus musculus OX=10090 GN=Phf5a PE=1 SV=1 | 99.9 | 93.2 | 90.7 | 87.6 | 103.3 | 104.1 | 69.3 | 104 | 92.7 | 155.2 |
| 689 | AP-2 complex subunit mu OS=Mus musculus OX=10090 GN=Ap2m1 PE=1 SV=1 | 137.2 | 115.1 | 114.9 | 85.9 | 91 | 113.1 | 72.2 | 93.5 | 88 | 89.2 |
| 690 | Histone H3.3 OS=Mus musculus OX=10090 GN=H3-3a PE=1 SV=2 | 80.6 | 131.9 | 107.1 | 62.3 | 197.6 | 59.8 | 73.3 | 88.3 | 52.8 | 146.3 |
| 691 | Isochorismatase domain-containing protein 2A OS=Mus musculus OX=10090 GN=Isoc2a PE=1 SV=1 | 144.4 | 111.2 | 118.8 | 116.4 | 54.3 | 94.7 | 52.4 | 91.9 | 127.8 | 88.2 |
| 692 | 40S ribosomal protein S3a OS=Mus musculus OX=10090 GN=Rps3a PE=1 SV=3 | 145.4 | 115.4 | 110.3 | 89.4 | 87 | 128.9 | 68.4 | 102 | 74.3 | 79 |
| 693 | Annexin A11 OS=Mus musculus OX=10090 GN=Anxa11 PE=1 SV=2 | 124.3 | 92.1 | 106.9 | 77.5 | 106.1 | 111.6 | 114.3 | 98.7 | 72.8 | 95.6 |
| 694 | Annexin A4 OS=Mus musculus OX=10090 GN=Anxa4 PE=1 SV=4 | 158.7 | 104.6 | 115.1 | 93 | 86.9 | 136.3 | 64 | 96.2 | 81 | 64.2 |
| 695 | Aminopeptidase N OS=Mus musculus OX=10090 GN=Anpep PE=1 SV=4 | 75.5 | 97.5 | 105.9 | 68.1 | 163.7 | 88.1 | 139.1 | 90.8 | 74 | 97.4 |
| 696 | 40S ribosomal protein S5 OS=Mus musculus OX=10090 GN=Rps5 PE=1 SV=3 | 111.7 | 106.8 | 78.4 | 85.7 | 141.8 | 122.6 | 92.1 | 92.9 | 72.9 | 95.2 |
| 697 | 5-demethoxyubiquinone hydroxylase, mitochondrial OS=Mus musculus OX=10090 GN=Coq7 PE=1 SV=3 | 80.2 | 75.6 | 86.9 | 103.3 | 112.4 | 81.2 | 113.3 | 117.5 | 113.9 | 115.7 |
| 698 | Glutamate--cysteine ligase catalytic subunit OS=Mus musculus OX=10090 GN=Gclc PE=1 SV=4 | 120.6 | 108.5 | 133.8 | 99.5 | 94 | 135.9 | 65.2 | 93.2 | 76.4 | 72.9 |
| 699 | Dimethylaniline monooxygenase [N-oxide-forming] 3 OS=Mus musculus OX=10090 GN=Fmo3 PE=1 SV=1 | 108.3 | 99.8 | 104.2 | 115.1 | 35 | 74.2 | 43.5 | 154.9 | 169.7 | 95.2 |
| 700 | Carnitine O-palmitoyltransferase 1, liver isoform OS=Mus musculus OX=10090 GN=Cpt1a PE=1 SV=4 | 97.4 | 106.6 | 110.4 | 106.9 | 95.3 | 111.1 | 66.3 | 94.7 | 119.1 | 92.2 |

| | | | | | | | | | | | |
|---|---|---|---|---|---|---|---|---|---|---|---|
| 701 | Fumarate hydratase, mitochondrial OS=Mus musculus OX=10090 GN=Fh PE=1 SV=3 | 81.4 | 73 | 82.2 | 134.5 | 90.2 | 90.1 | 92.5 | 136.1 | 119.3 | 100.7 |
| 702 | Caspase-7 OS=Mus musculus OX=10090 GN=Casp7 PE=1 SV=2 | | 130.3 | | 150.7 | | 192.2 | 139 | 133.2 | 138.2 | 116.5 |
| 703 | Peroxisomal targeting signal 2 receptor OS=Mus musculus OX=10090 GN=Pex7 PE=1 SV=1 | 122.9 | 109.4 | 130.2 | 95 | 99.5 | 111.9 | 103.3 | 87.7 | 80.7 | 59.3 |
| 704 | Flavin-containing monooxygenase 5 OS=Mus musculus OX=10090 GN=Fmo5 PE=1 SV=4 | 102.9 | 81.7 | 103.8 | 130.1 | 68.9 | 102.4 | 56.3 | 132.3 | 133 | 88.5 |
| 705 | Cytochrome b-c1 complex subunit 6, mitochondrial OS=Mus musculus OX=10090 GN=Uqcrh PE=1 SV=2 | 68.9 | 75.9 | 59.8 | 71.8 | 178.3 | 55.2 | 229.3 | 92.4 | 64.1 | 104.4 |
| 706 | Peroxiredoxin-5, mitochondrial OS=Mus musculus OX=10090 GN=Prdx5 PE=1 SV=2 | 69.1 | 65.9 | 72.6 | 76.7 | 58 | 68.8 | 83.3 | 78.6 | 219.1 | 207.9 |
| 707 | Glucose-6-phosphate 1-dehydrogenase X OS=Mus musculus OX=10090 GN=G6pdx PE=1 SV=3 | 196.4 | 107.5 | 142 | 72.2 | 76 | 147.7 | 39.3 | 118.7 | 54.7 | 45.6 |
| 708 | Apolipoprotein A-I OS=Mus musculus OX=10090 GN=Apoa1 PE=1 SV=2 | 74.9 | 102.7 | 106 | 94.7 | 102.8 | 113.8 | 111.1 | 70.4 | 90 | 133.6 |
| 709 | Alpha-1-antitrypsin 1-3 OS=Mus musculus OX=10090 GN=Serpina1c PE=1 SV=2 | 96.5 | 125.9 | 133.6 | 90.3 | 89.7 | 82.2 | 100.3 | 90.7 | 85.2 | 105.7 |
| 710 | Alpha-1-antitrypsin 1-4 OS=Mus musculus OX=10090 GN=Serpina1d PE=1 SV=1 | 96 | 127.4 | 112.7 | 95.7 | 84.8 | 101.9 | 61.7 | 82.4 | 120.9 | 116.5 |
| 711 | Heterogeneous nuclear ribonucleoprotein U-like protein 2 OS=Mus musculus OX=10090 GN=Hnrnpul2 PE=1 SV=2 | 91.1 | 90.1 | 85.1 | 75.9 | 127.1 | 95.2 | 160.4 | 90.5 | 80.4 | 104.4 |
| 712 | Collagen alpha-2(I) chain OS=Mus musculus OX=10090 GN=Col1a2 PE=1 SV=2 | 68.4 | 133.8 | 81.2 | 62.8 | 194.5 | 39.6 | 204.9 | 77.3 | 59.2 | 78.2 |
| 713 | Beta-2-glycoprotein 1 OS=Mus musculus OX=10090 GN=Apoh PE=1 SV=1 | 99.5 | 89.4 | 89.2 | 106.1 | 76.6 | 75.9 | 111.4 | 115.4 | 103.2 | 133.2 |
| 714 | Aquaporin-1 OS=Mus musculus OX=10090 GN=Aqp1 PE=1 SV=3 | 121.5 | 108.2 | 131.8 | 95 | 80.1 | 115.9 | 76.2 | 89.9 | 96.7 | 84.7 |
| 715 | Junction plakoglobin OS=Mus musculus OX=10090 GN=Jup PE=1 SV=3 | 118 | 101.3 | 101.1 | 111.7 | 68 | 102.3 | 72 | 98.4 | 111.9 | 115.4 |
| 716 | ATP synthase subunit alpha, mitochondrial OS=Mus musculus OX=10090 GN=Atp5f1a PE=1 SV=1 | 96.6 | 89.6 | 92.5 | 105.4 | 107.7 | 92.4 | 127.9 | 100.6 | 102.1 | 85.3 |
| 717 | Cholinesterase OS=Mus musculus OX=10090 GN=Bche PE=1 SV=2 | 110.4 | 95.3 | 108.9 | 104 | 134.2 | 105.2 | 114.7 | 80.6 | 89.3 | 57.5 |
| 718 | Collagen alpha-1(VI) chain OS=Mus musculus OX=10090 GN=Col6a1 PE=1 SV=1 | 61 | 93 | 92 | 72.4 | 155.9 | 61.8 | 247.2 | 65.4 | 65.1 | 86.2 |
| 719 | Basement membrane-specific heparan sulfate proteoglycan core protein OS=Mus musculus OX=10090 GN=Hspg2 PE=1 SV=1 | 99.8 | 98.2 | 92.9 | 85.1 | 116.5 | 95.9 | 129.1 | 93 | 85.2 | 104.2 |
| 720 | Fatty acid-binding protein 5 OS=Mus musculus OX=10090 GN=Fabp5 PE=1 SV=3 | 217.7 | 162.2 | 129.4 | 40 | 132.4 | 112.5 | 70 | 48.1 | 35 | 52.8 |
| 721 | GTP cyclohydrolase 1 OS=Mus musculus OX=10090 GN=Gch1 PE=1 SV=1 | 123.8 | 87.9 | 116.7 | 120.1 | 68.2 | 109.6 | 67.4 | 113.1 | 110.4 | 82.8 |
| 722 | Echinoderm microtubule-associated protein-like 1 OS=Mus musculus OX=10090 GN=Eml1 PE=1 SV=1 | 69.7 | 67.7 | 71.2 | 55 | 228 | 89.5 | 214.1 | 83.7 | 50.5 | 70.6 |
| 723 | Eukaryotic translation initiation factor 5B OS=Mus musculus OX=10090 GN=Eif5b PE=1 SV=2 | 137.7 | 89.8 | 97.8 | 93.5 | 121.8 | 114.4 | 115.8 | 63.3 | 75.8 | 90 |
| 724 | NADPH--hemoprotein reductase OS=Mus musculus OX=10090 GN=Por PE=1 SV=1 | 88 | 88.4 | 91.4 | 112.9 | 43.5 | 111 | 78.7 | 145.5 | 124.3 | 116.2 |
| 725 | Calcium-binding protein 39 OS=Mus musculus OX=10090 GN=Cab39 PE=1 SV=2 | 110.9 | 91.7 | 114.9 | 92.8 | 92.3 | 135.8 | 112.9 | 96.6 | 81.1 | 71 |
| 726 | ATP synthase subunit e, mitochondrial OS=Mus musculus OX=10090 GN=Atp5me PE=1 SV=2 | 104.7 | 86.7 | 93.8 | 113.1 | 89.5 | 92.3 | 109.4 | 113.3 | 104.2 | 93 |
| 727 | Annexin A7 OS=Mus musculus OX=10090 GN=Anxa7 PE=1 SV=2 | | 109.5 | 132.6 | 119.3 | 87.2 | 128.4 | 103 | 116.9 | 108.3 | 94.9 |
| 728 | Histone H1t OS=Mus musculus OX=10090 GN=H1-6 PE=1 SV=4 | 134.8 | 160.8 | 141.6 | 68.6 | 48.5 | 69.3 | 60 | 103.1 | 87 | 126.3 |
| 729 | Galectin-3-binding protein OS=Mus musculus OX=10090 GN=Lgals3bp PE=1 SV=1 | 108.5 | 94.5 | 140.1 | 80.7 | 116.7 | 110 | 92.6 | 48.4 | 80.2 | 128.3 |
| 730 | Platelet glycoprotein 4 OS=Mus musculus OX=10090 GN=Cd36 PE=1 SV=2 | 88.7 | 82.7 | 97.9 | 101.3 | 109.7 | 94.1 | 130.9 | 94.8 | 99.1 | 100.8 |
| 731 | Cytochrome P450 2E1 OS=Mus musculus OX=10090 GN=Cyp2e1 PE=1 SV=1 | 125.8 | 124.8 | 130.9 | 105.5 | 61.7 | 92.7 | 63.9 | 130.5 | 88.6 | 75.6 |
| 732 | Pentatricopeptide repeat domain-containing protein 3, mitochondrial OS=Mus musculus OX=10090 GN=Ptcd3 PE=1 SV=2 | 104.2 | 87.5 | 100.4 | 119.8 | 70.6 | 92.3 | 54.3 | 122.4 | 129.6 | 119 |
| 733 | Acyl-CoA synthetase short-chain family member 3, mitochondrial OS=Mus musculus OX=10090 GN=Acss3 PE=1 SV=2 | 115.4 | 93.7 | 96.8 | 111.9 | 82.3 | 106.1 | 88 | 118.6 | 107.3 | 79.9 |
| 734 | Collagen alpha-1(XXIV) chain OS=Mus musculus OX=10090 GN=Col24a1 PE=2 SV=2 | 53.9 | 139.6 | 74.2 | 65.2 | 202.9 | 24.1 | 236.5 | 79.1 | 47.2 | 77.3 |

| | | | | | | | | | | | |
|---|---|---|---|---|---|---|---|---|---|---|---|
| 735 | Fumarylacetoacetate hydrolase domain-containing protein 2A OS=Mus musculus OX=10090 GN=Fahd2 PE=1 SV=1 | 43.1 | 139 | 46.3 | 126 | 38.1 | 125.8 | 129.7 | 125.2 | 117.6 | 109 |
| 736 | Enoyl-CoA delta isomerase 2 OS=Mus musculus OX=10090 GN=Eci2 PE=1 SV=1 | 107.3 | 98.7 | 107.3 | 117.1 | 66 | 89.8 | 61.4 | 122.1 | 115.7 | 114.5 |
| 737 | Heterochromatin protein 1-binding protein 3 OS=Mus musculus OX=10090 GN=Hp1bp3 PE=1 SV=1 | 121.5 | 97.4 | 117.1 | 95.5 | 82.4 | 107.6 | 78.7 | 119.8 | 89.2 | 90.9 |
| 738 | Myosin regulatory light chain 12B OS=Mus musculus OX=10090 GN=Myl12b PE=1 SV=2 | | 105.6 | 113.8 | 107.7 | 119.9 | 119.8 | 71.1 | 113.8 | 112.7 | 135.7 |
| 739 | GMP synthase [glutamine-hydrolyzing] OS=Mus musculus OX=10090 GN=Gmps PE=1 SV=2 | 134.9 | 86.3 | 103.2 | 83.4 | 135.7 | 113.8 | 71.2 | 99.9 | 74.6 | 97 |
| 740 | Corticosteroid 11-beta-dehydrogenase isozyme 1 OS=Mus musculus OX=10090 GN=Hsd11b1 PE=1 SV=1 | 148.7 | 111.5 | 135 | 88.1 | 80 | 114.1 | 85.9 | 94.4 | 88 | 54.3 |
| 741 | Enoyl-CoA hydratase domain-containing protein 2, mitochondrial OS=Mus musculus OX=10090 GN=Echdc2 PE=1 SV=2 | 103 | 87.4 | 91.6 | 102.6 | 113.5 | 123.3 | 84 | 113.9 | 89.6 | 91 |
| 742 | Ectonucleoside triphosphate diphosphohydrolase 5 OS=Mus musculus OX=10090 GN=Entpd5 PE=1 SV=1 | 78.5 | 80.9 | 115.3 | 135.3 | 74.6 | 91.9 | 49.5 | 134.7 | 145.3 | 93.9 |
| 743 | Malic enzyme OS=Mus musculus OX=10090 GN=Me1 PE=1 SV=1 | 216.6 | 163.3 | 168.9 | 61.1 | 83.3 | 94 | 81.1 | 41.8 | 46.6 | 43.3 |
| 744 | Adapter molecule crk OS=Mus musculus OX=10090 GN=Crk PE=1 SV=1 | 121.1 | 100.8 | 120.4 | 94.4 | 89.5 | 115.8 | 72 | 95.7 | 86.7 | 103.5 |
| 745 | Actin-related protein 2/3 complex subunit 4 OS=Mus musculus OX=10090 GN=Arpc4 PE=1 SV=1 | 121.8 | 109.9 | 110.6 | 92.3 | 87.9 | 133.2 | 77.5 | 86.3 | 90.2 | 90.4 |
| 746 | Peregrin OS=Mus musculus OX=10090 GN=Brpf1 PE=1 SV=1 | | 96 | | 106.5 | | 110.9 | 289.8 | 112.8 | 125 | 159 |
| 747 | DNA damage-binding protein 1 OS=Mus musculus OX=10090 GN=Ddb1 PE=1 SV=2 | 67.5 | 99.5 | 97.9 | 89.1 | 75.3 | 93.1 | 109.9 | 76.7 | 149.1 | 141.9 |
| 748 | Polymerase delta-interacting protein 3 OS=Mus musculus OX=10090 GN=Poldip3 PE=1 SV=1 | 92.9 | 92.7 | 110.3 | 68.2 | 131.2 | 121.3 | 144.5 | 83.7 | 75.4 | 79.7 |
| 749 | Interferon-gamma-inducible GTPase Ifgga2 protein OS=Mus musculus OX=10090 GN=Gm4951 PE=1 SV=1 | 102.3 | 81.4 | 106 | 70.2 | 160.4 | 180.4 | 124.4 | 64.1 | 58.6 | 52.2 |
| 750 | Imidazolonepropionate hydrolase OS=Mus musculus OX=10090 GN=Uroc1 PE=1 SV=1 | 118.2 | 106.3 | 112.1 | 95.4 | 86.6 | 124.3 | 82.5 | 103.4 | 90.8 | 80.3 |
| 751 | Fibrinogen gamma chain OS=Mus musculus OX=10090 GN=Fgg PE=1 SV=1 | 106.6 | 85.8 | 129.3 | 72.6 | 84.6 | 94.6 | 81 | 130.9 | 94 | 120.5 |
| 752 | Monoglyceride lipase OS=Mus musculus OX=10090 GN=Mgll PE=1 SV=1 | 117.5 | 112.5 | 115.1 | 88.9 | 101.7 | 101.5 | 105.2 | 99.6 | 78.8 | 79.1 |
| 753 | NADH dehydrogenase [ubiquinone] 1 beta subcomplex subunit 6 OS=Mus musculus OX=10090 GN=Ndufb6 PE=1 SV=3 | 92.7 | 80 | 87.9 | 109.8 | 93.9 | 95.2 | 101.5 | 131.5 | 101.1 | 106.4 |
| 754 | Methylcrotonoyl-CoA carboxylase beta chain, mitochondrial OS=Mus musculus OX=10090 GN=Mccc2 PE=1 SV=1 | 104.3 | 94.5 | 93.9 | 102.8 | 96 | 92.3 | 100.4 | 115.5 | 103.9 | 96.5 |
| 755 | Glycerol-3-phosphate dehydrogenase 1-like protein OS=Mus musculus OX=10090 GN=Gpd1l PE=1 SV=2 | 116.5 | 101.7 | 108.9 | 129.6 | 67.9 | 109 | 59.4 | 99.2 | 113.9 | 93.9 |
| 756 | Plasminogen activator inhibitor 1 RNA-binding protein OS=Mus musculus OX=10090 GN=Serbp1 PE=1 SV=1 | | 153.9 | | 102 | | 166.5 | 134.9 | 149.7 | 98.7 | 194.3 |
| 757 | Hepatoma-derived growth factor-related protein 2 OS=Mus musculus OX=10090 GN=Hdgfl2 PE=1 SV=1 | 98.7 | 99 | 92 | 79.2 | 107.6 | 106.1 | 113.3 | 105.7 | 80.1 | 118.2 |
| 758 | Acyl-coenzyme A synthetase ACSM3, mitochondrial OS=Mus musculus OX=10090 GN=Acsm3 PE=1 SV=2 | 93.9 | 84.2 | 90.7 | 123.3 | 72.2 | 92.7 | 68.1 | 119.8 | 129.1 | 125.9 |
| 759 | Inactive phospholipase C-like protein 1 OS=Mus musculus OX=10090 GN=Plcl1 PE=1 SV=3 | 7.9 | 32.2 | 13.8 | 14.9 | 513.5 | 5.4 | 333.8 | 11.7 | 9.9 | 56.9 |
| 760 | Cytochrome P450 2C29 OS=Mus musculus OX=10090 GN=Cyp2c29 PE=1 SV=1 | 128 | 98.6 | 119.5 | 110 | 69.4 | 96.4 | 42.3 | 125 | 126.9 | 83.9 |
| 761 | Bifunctional ATP-dependent dihydroxyacetone kinase/FAD-AMP lyase (cyclizing) OS=Mus musculus OX=10090 GN=Tkfc PE=1 SV=1 | 121.2 | 102.7 | 148.5 | 105.5 | 90.8 | 104.9 | 54.2 | 99.5 | 81.4 | 91.2 |
| 762 | 40S ribosomal protein S10 OS=Mus musculus OX=10090 GN=Rps10 PE=1 SV=1 | 63.9 | 117.8 | 63.4 | 123.5 | 51.4 | 131.3 | 86.5 | 119.9 | 116.7 | 125.6 |
| 763 | Aldo-keto reductase family 1, member C-like OS=Mus musculus OX=10090 GN=Akr1cl PE=1 SV=1 | 146.1 | 130.9 | 142.9 | 87.9 | 101.6 | 122.6 | 74.1 | 68.3 | 64.8 | 60.8 |
| 764 | Aminoacyl tRNA synthase complex-interacting multifunctional protein 1 OS=Mus musculus OX=10090 GN=Aimp1 PE=1 SV=1 | 98.8 | 110.6 | 103 | 73.9 | 143.6 | 132.5 | 83.6 | 90.9 | 67.8 | 95.5 |
| 765 | Kynurenine 3-monooxygenase OS=Mus musculus OX=10090 GN=Kmo PE=1 SV=1 | 104.9 | 96 | 109.3 | 113 | 78.1 | 86.5 | 66.3 | 122 | 120.2 | 103.7 |
| 766 | Polypyrimidine tract-binding protein 2 OS=Mus musculus OX=10090 GN=Ptbp2 PE=1 SV=1 | 127.6 | 83.7 | 134.8 | 85.2 | 96.9 | 117.4 | 28.4 | 77.9 | 97.9 | 150.1 |
| 767 | Integrin alpha-1 OS=Mus musculus OX=10090 GN=Itga1 PE=1 SV=2 | 87.3 | 87.1 | 99.6 | 66.7 | 88.7 | 175 | 109.7 | 108.2 | 73 | 104.7 |

| | | | | | | | | | | | |
|---|---|---|---|---|---|---|---|---|---|---|---|
| 768 | Interleukin enhancer-binding factor 3 OS=Mus musculus OX=10090 GN=Ilf3 PE=1 SV=1 | 145.5 | 118.7 | 111.5 | 88.7 | 92.9 | 106.7 | 67.7 | 104.7 | 82 | 81.7 |
| 769 | Fibronectin OS=Mus musculus OX=10090 GN=Fn1 PE=1 SV=1 | 93.7 | 93 | 106 | 83.3 | 116.7 | 111.5 | 122.7 | 96.4 | 79.5 | 97.1 |
| 770 | Heat shock cognate 71 kDa protein OS=Mus musculus OX=10090 GN=Hspa8 PE=1 SV=1 | 121.5 | 110.6 | 114.8 | 87.1 | 102.7 | 112.3 | 84.5 | 93.4 | 93.3 | 79.8 |
| 771 | Chaperonin containing Tcp1, subunit 6a (Zeta) OS=Mus musculus OX=10090 GN=Cct6a PE=1 SV=1 | 124.9 | 103.6 | 126.3 | 93.9 | 101.4 | 109.8 | 107.9 | 68.2 | 76.7 | 87.4 |
| 772 | Cytochrome P450, family 2, subfamily c, polypeptide 67 OS=Mus musculus OX=10090 GN=Cyp2c67 PE=1 SV=1 | 117.2 | 139.2 | 141.2 | 108.6 | 57.3 | 93.1 | 78.9 | 96.9 | 107.9 | 59.6 |
| 773 | Major urinary protein 1 OS=Mus musculus OX=10090 GN=Mup7 PE=2 SV=1 | 181.7 | 189.7 | 149.5 | 64.1 | 64.8 | 77.3 | 64.1 | 81.6 | 69 | 58.1 |
| 774 | Glycine N-acyltransferase-like protein OS=Mus musculus OX=10090 GN=Gm4952 PE=1 SV=3 | 139 | 124.2 | 117 | 105 | 105.3 | 109.5 | 83.8 | 66.1 | 84.3 | 66 |
| 775 | Alpha-2-antiplasmin (Fragment) OS=Mus musculus OX=10090 GN=Serpinf2 PE=1 SV=1 | 104.6 | 122.6 | 146 | 99.2 | 92.9 | 108.5 | 91.4 | 74.4 | 78.1 | 82.1 |
| 776 | Phosphatidylcholine transfer protein OS=Mus musculus OX=10090 GN=Pctp PE=1 SV=1 | 117.4 | 114.6 | 132.3 | 118.8 | 53.2 | 105.7 | 62.3 | 104.6 | 110.7 | 80.4 |
| 777 | Acetyl-CoA carboxylase 1 OS=Mus musculus OX=10090 GN=Acaca PE=1 SV=1 | 136.9 | 116.3 | 133.5 | 95.8 | 88.8 | 121.5 | 63.6 | 94.2 | 74.7 | 74.9 |
| 778 | Myosin-4 OS=Mus musculus OX=10090 GN=Myh4 PE=1 SV=1 | 59.2 | 84.6 | 57.5 | 67.5 | 118.4 | 59 | 373.1 | 58 | 47.4 | 75.3 |
| 779 | Myosin-1 OS=Mus musculus OX=10090 GN=Myh1 PE=1 SV=1 | 15.4 | 38.9 | 12.6 | 17.1 | 301.6 | 11.9 | 544.6 | 17.6 | 7.1 | 33.3 |
| 780 | Clathrin heavy chain OS=Mus musculus OX=10090 GN=Cltc PE=1 SV=1 | 138.8 | 93.1 | 118.5 | 92.4 | 82.5 | 109 | 71.8 | 101 | 87.9 | 104.9 |
| 781 | Coatomer subunit delta OS=Mus musculus OX=10090 GN=Arcn1 PE=1 SV=2 | 105.7 | 121 | 98.4 | 96.6 | 69.5 | 136.6 | 73.1 | 101.6 | 98.3 | 99.1 |
| 782 | Myosin-binding protein C, fast-type OS=Mus musculus OX=10090 GN=Mybpc2 PE=1 SV=1 | 70.5 | 87.6 | 115 | 72.1 | 176.8 | 92.3 | 185.5 | 67.1 | 52.3 | 80.9 |
| 783 | Myosin light polypeptide 6 OS=Mus musculus OX=10090 GN=Myl6 PE=1 SV=3 | 97.6 | 108.4 | 94.7 | 87.6 | 130.2 | 90.7 | 105.2 | 97.5 | 77 | 111.2 |
| 784 | Growth factor receptor-bound protein 2 OS=Mus musculus OX=10090 GN=Grb2 PE=1 SV=1 | 83 | 98.2 | 82.1 | 67.5 | 162.1 | 98.1 | 167.4 | 81.8 | 65.1 | 94.8 |
| 785 | Ganglioside GM2 activator OS=Mus musculus OX=10090 GN=Gm2a PE=1 SV=2 | 44.5 | 120 | 121.5 | 98 | 104.6 | 108.2 | 87.6 | 96.7 | 95.3 | 123.6 |
| 786 | Deoxynucleoside triphosphate triphosphohydrolase SAMHD1 OS=Mus musculus OX=10090 GN=Samhd1 PE=1 SV=3 | 120.5 | 106.9 | 115.2 | 84.1 | 94.3 | 139.7 | 83.7 | 79.9 | 86 | 89.8 |
| 787 | Immunity-related GTPase family M protein 1 OS=Mus musculus OX=10090 GN=Irgm1 PE=1 SV=1 | 99.9 | 87.8 | 138 | 100.7 | 98.5 | 122.8 | 116.2 | 75.9 | 81.3 | 78.9 |
| 788 | Phosphotriesterase-related protein OS=Mus musculus OX=10090 GN=Pter PE=1 SV=1 | 162.7 | 146.1 | 147.9 | 72 | 106.3 | 141.1 | 72.8 | 55.2 | 42.6 | 53.5 |
| 789 | Atypical kinase COQ8A, mitochondrial OS=Mus musculus OX=10090 GN=Coq8a PE=1 SV=2 | 62.3 | 98.9 | 109.9 | 107.8 | 120.3 | 119.3 | 143.3 | 82.9 | 85.4 | 70.1 |
| 790 | Histone-binding protein RBBP7 OS=Mus musculus OX=10090 GN=Rbbp7 PE=1 SV=1 | 127.8 | 116.7 | 109.5 | 81.2 | 91 | 122.5 | 82.1 | 93.1 | 84.4 | 91.7 |
| 791 | Histidine--tRNA ligase, cytoplasmic OS=Mus musculus OX=10090 GN=Hars1 PE=1 SV=2 | 138.4 | 117.1 | 111.6 | 101.2 | 79.2 | 126 | 66.4 | 102.6 | 81.3 | 76.3 |
| 792 | Hsp90 co-chaperone Cdc37 OS=Mus musculus OX=10090 GN=Cdc37 PE=1 SV=1 | 129.3 | 115.9 | 122.5 | 93.4 | 94.3 | 126.8 | 73.6 | 82.4 | 83.8 | 78.1 |
| 793 | Iron-sulfur clusters transporter ABCB7, mitochondrial OS=Mus musculus OX=10090 GN=Abcb7 PE=1 SV=3 | 112.9 | 97.2 | 99.4 | 101.8 | 90.4 | 110.8 | 85.5 | 104.3 | 97.8 | 100 |
| 794 | Peroxiredoxin-2 OS=Mus musculus OX=10090 GN=Prdx2 PE=1 SV=3 | 109.1 | 104.3 | 122.4 | 102.6 | 112.6 | 111.2 | 79.9 | 96.9 | 90.4 | 70.7 |
| 795 | Arginase-1 OS=Mus musculus OX=10090 GN=Arg1 PE=1 SV=1 | 113.9 | 100.9 | 99.2 | 104.5 | 74.4 | 93.6 | 73.2 | 130.5 | 102.9 | 107.1 |
| 796 | Platelet-activating factor acetylhydrolase IB subunit alpha2 OS=Mus musculus OX=10090 GN=Pafah1b2 PE=1 SV=2 | 129.5 | 122.3 | 127 | 94.1 | 84 | 121.1 | 42.8 | 90 | 88.4 | 100.8 |
| 797 | Plastin-2 OS=Mus musculus OX=10090 GN=Lcp1 PE=1 SV=4 | 112.5 | 104.6 | 121.3 | 91.5 | 103.9 | 162.3 | 64.2 | 78.8 | 80.4 | 80.4 |
| 798 | Heat shock 70 kDa protein 4 OS=Mus musculus OX=10090 GN=Hspa4 PE=1 SV=1 | 110.5 | 104.8 | 109.3 | 89.2 | 90.9 | 114.6 | 90.9 | 85.7 | 104.7 | 99.3 |
| 799 | B-cell receptor-associated protein 31 OS=Mus musculus OX=10090 GN=Bcap31 PE=1 SV=4 | 104.5 | 102.6 | 120.1 | 79.4 | 122.3 | 115.9 | 132.3 | 80.6 | 73.9 | 68.4 |
| 800 | Hydroxyacyl-coenzyme A dehydrogenase, mitochondrial OS=Mus musculus OX=10090 GN=Hadh PE=1 SV=2 | 120.5 | 99.6 | 109 | 104.3 | 113 | 94.6 | 94.9 | 99.5 | 89.2 | 75.3 |
| 801 | Fibrillin-1 OS=Mus musculus OX=10090 GN=Fbn1 PE=1 SV=2 | 68.8 | 84 | 83.4 | 62.2 | 177.9 | 64.4 | 238.6 | 66.6 | 61.5 | 92.7 |
| 802 | NADPH:adrenodoxin oxidoreductase, mitochondrial OS=Mus musculus OX=10090 GN=Fdxr PE=1 SV=1 | 121.7 | 104.1 | 120.3 | 111.7 | 85.5 | 114 | 60.6 | 85.6 | 114.2 | 82.4 |

| | | | | | | | | | | | |
|---|---|---|---|---|---|---|---|---|---|---|---|
| 803 | G0/G1 switch protein 2 OS=Mus musculus OX=10090 GN=G0s2 PE=1 SV=1 | 110 | 100.7 | 110.7 | 108 | 73.1 | 100.9 | 69.4 | 121.8 | 105.2 | 100.2 |
| 804 | GTP-binding protein OS=Mus musculus OX=10090 GN=Ifi47 PE=1 SV=1 | 110.2 | 100.5 | 127 | 74.7 | 132.7 | 182.3 | 102.8 | 60.3 | 55.2 | 54.1 |
| 805 | Haptoglobin OS=Mus musculus OX=10090 GN=Hp PE=1 SV=1 | | 107.3 | 98.5 | 67.7 | 127.6 | 119.7 | 87.9 | 156 | 68.3 | 167 |
| 806 | Probable ATP-dependent RNA helicase DDX5 OS=Mus musculus OX=10090 GN=Ddx5 PE=1 SV=2 | 87.9 | 104.5 | 88.9 | 91.5 | 93.5 | 108 | 118.3 | 96 | 87.7 | 123.7 |
| 807 | Heat shock 70 kDa protein 1A OS=Mus musculus OX=10090 GN=Hspa1a PE=1 SV=2 | 95.5 | 92.1 | 106.3 | 87.3 | 123.7 | 106 | 121.1 | 85 | 83.6 | 99.4 |
| 808 | Inter-alpha-trypsin inhibitor heavy chain H2 OS=Mus musculus OX=10090 GN=Itih2 PE=1 SV=1 | 108 | 108.3 | 102 | 109.7 | 84.5 | 97.2 | 87.6 | 100.8 | 97.5 | 104.5 |
| 809 | LIM and SH3 domain protein 1 OS=Mus musculus OX=10090 GN=Lasp1 PE=1 SV=1 | 43.9 | 114.6 | 50 | 103.7 | 72.9 | 159.8 | 83.9 | 107 | 98.3 | 165.9 |
| 810 | Macrophage mannose receptor 1 OS=Mus musculus OX=10090 GN=Mrc1 PE=1 SV=2 | 94.9 | 134.2 | 94.1 | 103.8 | 72.6 | 134.9 | 93.8 | 86.4 | 95.7 | 89.7 |
| 811 | Mitogen-activated protein kinase 10 OS=Mus musculus OX=10090 GN=Mapk10 PE=1 SV=2 | 121.5 | 92.7 | 113.7 | 102.4 | 102.7 | 112.3 | 83.7 | 101.2 | 89.1 | 80.7 |
| 812 | Pregnancy zone protein OS=Mus musculus OX=10090 GN=Pzp PE=1 SV=3 | 75.3 | 80.9 | 108 | 62.7 | 80.7 | 85.9 | 127.4 | 58.6 | 124.6 | 195.9 |
| 813 | Myosin regulatory light chain 10 OS=Mus musculus OX=10090 GN=Myl10 PE=2 SV=1 | 59.3 | 71.4 | 59 | 59.3 | 210.6 | 59.8 | 276.1 | 60.6 | 45.4 | 98.3 |
| 814 | ATP-dependent RNA helicase DDX3X OS=Mus musculus OX=10090 GN=Ddx3x PE=1 SV=3 | 62.2 | 51.8 | 68.8 | 47.6 | 45.1 | 56.3 | 158.3 | 56.9 | 192.6 | 260.3 |
| 815 | Carboxylesterase 3A OS=Mus musculus OX=10090 GN=Ces3a PE=1 SV=2 | 157.3 | 205.4 | 160.7 | 24.9 | 73.7 | 165.7 | 100.1 | 20.9 | 45.6 | 45.8 |
| 816 | Caveolae-associated protein 2 OS=Mus musculus OX=10090 GN=Cavin2 PE=1 SV=3 | 78.4 | 94.9 | 97.1 | 71.7 | 173.6 | 73.4 | 158.6 | 71.7 | 60 | 120.5 |
| 817 | Carboxylesterase 1E OS=Mus musculus OX=10090 GN=Ces1e PE=1 SV=1 | | 138.4 | | 158.3 | | 157.3 | 112.7 | 140.2 | 157 | 136.2 |
| 818 | Interferon-induced protein with tetratricopeptide repeats 1 OS=Mus musculus OX=10090 GN=Ifit1 PE=1 SV=2 | 60.9 | 40 | 183 | 92.2 | 80.5 | 116.1 | 142.3 | 54.6 | 110.2 | 120.1 |
| 819 | Calcium/calmodulin-dependent 3',5'-cyclic nucleotide phosphodiesterase 1C OS=Mus musculus OX=10090 GN=Pde1c PE=1 SV=2 | 111 | 86.2 | 75.2 | 111.5 | 137 | 97.6 | 105.5 | 89.2 | 104.7 | 82.1 |
| 820 | 10 kDa heat shock protein, mitochondrial OS=Mus musculus OX=10090 GN=Hspe1 PE=1 SV=2 | 75.1 | 90.1 | 77.1 | 79.3 | 154.8 | 94 | 63.1 | 116.3 | 86.8 | 163.5 |
| 821 | Potassium-transporting ATPase alpha chain 1 OS=Mus musculus OX=10090 GN=Atp4a PE=1 SV=4 | 96.5 | 89 | 98.6 | 104.9 | 68.8 | 89.9 | 65.7 | 125.2 | 120.4 | 141 |
| 822 | All-trans-retinol dehydrogenase [NAD(+)] ADH7 OS=Mus musculus OX=10090 GN=Adh7 PE=2 SV=2 | 135.1 | 120.8 | 132.3 | 107.9 | 60.6 | 85.9 | 53.9 | 107.3 | 102 | 94.1 |
| 823 | Copper-transporting ATPase 2 OS=Mus musculus OX=10090 GN=Atp7b PE=1 SV=2 | | 150.4 | 142.6 | 98.3 | 81.2 | 124.5 | 87.5 | 131.6 | 96.8 | 87.1 |
| 824 | Cytochrome P450 3A11 OS=Mus musculus OX=10090 GN=Cyp3a11 PE=1 SV=1 | 85 | 87.1 | 137.2 | 100.8 | 53.1 | 76.8 | 30.5 | 182.1 | 158.5 | 88.9 |
| 825 | Cytochrome P450 3A13 OS=Mus musculus OX=10090 GN=Cyp3a13 PE=1 SV=1 | 104.6 | 87 | 98.5 | 84.9 | 95.9 | 125 | 45.4 | 152.1 | 101.7 | 104.8 |
| 826 | Glyceraldehyde-3-phosphate dehydrogenase, testis-specific OS=Mus musculus OX=10090 GN=Gapdhs PE=1 SV=1 | 109.6 | 91.8 | 120.6 | 98.3 | 116 | 96.9 | 117.9 | 86.5 | 93.8 | 68.6 |
| 827 | Glutathione S-transferase theta-1 OS=Mus musculus OX=10090 GN=Gstt1 PE=1 SV=4 | 127 | 118.4 | 132.4 | 107.2 | 63.4 | 107.8 | 58.3 | 107 | 102.6 | 76.1 |
| 828 | Cytochrome P450 3A16 OS=Mus musculus OX=10090 GN=Cyp3a16 PE=2 SV=2 | 88.2 | 83.6 | 111.8 | 107.2 | 53.9 | 78.4 | 52.2 | 194.9 | 146 | 83.9 |
| 829 | Glycerol kinase OS=Mus musculus OX=10090 GN=Gk PE=1 SV=2 | 142.4 | 139.2 | 137.4 | 98.6 | 71.6 | 132.5 | 54.8 | 79 | 79 | 65.6 |
| 830 | Methionine--tRNA ligase, cytoplasmic OS=Mus musculus OX=10090 GN=Mars1 PE=1 SV=1 | 122.5 | 109.6 | 126.3 | 85.2 | 86 | 115.3 | 92.2 | 98.3 | 79.2 | 85.2 |
| 831 | Cell division cycle 5-like protein OS=Mus musculus OX=10090 GN=Cdc5l PE=1 SV=2 | 86.8 | 87.3 | 84 | 67.7 | 209.6 | 84.8 | 135.3 | 89.2 | 64.8 | 90.5 |
| 832 | Probable aminopeptidase NPEPL1 OS=Mus musculus OX=10090 GN=Npepl1 PE=1 SV=1 | 118.9 | 104.7 | 117.3 | 74.3 | 69.7 | 96.9 | 114 | 73.6 | 112.7 | 117.8 |
| 833 | Acsm5 protein OS=Mus musculus OX=10090 GN=Acsm5 PE=1 SV=1 | 112.7 | 90.4 | 98.7 | 132.1 | 62.1 | 90.1 | 52.7 | 116.8 | 135.8 | 108.6 |
| 834 | Creatine kinase S-type, mitochondrial OS=Mus musculus OX=10090 GN=Ckmt2 PE=1 SV=1 | 43.8 | 59.1 | 48.7 | 46.4 | 251.1 | 43.4 | 361.7 | 46.1 | 40.1 | 59.5 |
| 835 | Leucine-rich PPR motif-containing protein, mitochondrial OS=Mus musculus OX=10090 GN=Lrpprc PE=1 SV=2 | 113.7 | 89.9 | 109.2 | 106 | 68.5 | 82.7 | 91.9 | 130.2 | 123.4 | 84.5 |
| 836 | Mccc2 protein (Fragment) OS=Mus musculus OX=10090 GN=Mccc2 PE=1 SV=1 | 104.3 | 94.5 | 93.9 | 102.8 | 96 | 92.3 | 100.4 | 115.5 | 103.9 | 96.5 |
| 837 | Antiviral innate immune response receptor RIG-I OS=Mus musculus OX=10090 GN=Ddx58 PE=1 SV=2 | 125.1 | 100 | 124.6 | 91.7 | 102 | 134.6 | 104.8 | 75.9 | 67.6 | 73.6 |

| | | | | | | | | | | | |
|---|---|---|---|---|---|---|---|---|---|---|---|
| 838 | HMW kininogen-II OS=Mus musculus OX=10090 GN=Kng2 PE=1 SV=1 | 69.2 | 82.6 | 105.2 | 82.9 | 111 | 77 | 111.9 | 114.2 | 109.1 | 136.7 |
| 839 | Myosin-14 OS=Mus musculus OX=10090 GN=Myh14 PE=1 SV=1 | 92.3 | 82.8 | 99.4 | 84.3 | 99.6 | 112.3 | 137.7 | 99.4 | 99.8 | 92.4 |
| 840 | Cytochrome P450 2C54 OS=Mus musculus OX=10090 GN=Cyp2c54 PE=1 SV=1 | 95.8 | 93.9 | 107.7 | 81.6 | 66.4 | 88.1 | 129.4 | 139.6 | 93.1 | 104.3 |
| 841 | Fetuin-B OS=Mus musculus OX=10090 GN=Fetub PE=1 SV=1 | 125 | 114.4 | 132 | 91.3 | 121.2 | 110.8 | 70.9 | 78.5 | 78.4 | 77.6 |
| 842 | Cullin-associated NEDD8-dissociated protein 1 OS=Mus musculus OX=10090 GN=Cand1 PE=1 SV=2 | 105.4 | 99.1 | 97.4 | 95.7 | 91 | 116.5 | 116.3 | 98 | 85.9 | 94.7 |
| 843 | Eukaryotic translation initiation factor 2 subunit 1 OS=Mus musculus OX=10090 GN=Eif2s1 PE=1 SV=3 | 121.9 | 96.6 | 115 | 93.5 | 92.5 | 115.7 | 81.1 | 101.9 | 87.7 | 94.1 |
| 844 | Histone H2B type 1-C/E/G OS=Mus musculus OX=10090 GN=H2bc4 PE=1 SV=3 | 109.9 | 99.3 | 104 | 78 | 99.2 | 132.1 | 85.9 | 87.2 | 89.7 | 114.7 |
| 845 | 60S ribosomal protein L36 OS=Mus musculus OX=10090 GN=Rpl36 PE=1 SV=1 | 133.8 | 111.5 | 104.7 | 79.3 | 114.6 | 125.3 | 61.1 | 120.9 | 71.7 | 77.2 |
| 846 | Kynurenine--oxoglutarate transaminase 3 OS=Mus musculus OX=10090 GN=Kyat3 PE=1 SV=1 | 92.5 | 76.7 | 85.6 | 160.4 | 53.7 | 69.1 | 54.1 | 163.2 | 144.2 | 100.5 |
| 847 | ATP synthase membrane subunit K, mitochondrial OS=Mus musculus OX=10090 GN=Atp5mk PE=1 SV=1 | 67.6 | 70.3 | 59.7 | 67.2 | 144.4 | 58.2 | 316.6 | 76.6 | 58.9 | 80.4 |
| 848 | MICOS complex subunit Mic27 OS=Mus musculus OX=10090 GN=Apool PE=1 SV=1 | | 99.5 | | 79.2 | | 77.9 | 486.6 | 87.5 | 77.2 | 92.1 |
| 849 | Enoyl-CoA delta isomerase 3, peroxisomal OS=Mus musculus OX=10090 GN=Eci3 PE=1 SV=1 | 107.1 | 99.1 | 107.2 | 117.1 | 66.5 | 89.7 | 61.9 | 121.8 | 114.7 | 114.9 |
| 850 | L-fucose kinase OS=Mus musculus OX=10090 GN=Fcsk PE=1 SV=1 | 102.5 | 93.7 | 116.2 | 89.2 | 96.3 | 97.2 | 101.8 | 102.8 | 99.7 | 100.6 |
| 851 | E3 ubiquitin-protein ligase HUWE1 OS=Mus musculus OX=10090 GN=Huwe1 PE=1 SV=5 | 115.2 | 106 | 119.4 | 112.2 | 73.6 | 110 | 57.9 | 107.9 | 106.4 | 91.3 |
| 852 | Probable D-lactate dehydrogenase, mitochondrial OS=Mus musculus OX=10090 GN=Ldhd PE=1 SV=1 | 80.7 | 99 | 101.4 | 101.8 | 93.2 | 102.1 | 76.3 | 121.3 | 98.4 | 125.8 |
| 853 | Myo1b protein OS=Mus musculus OX=10090 GN=Myo1b PE=1 SV=1 | 61.8 | 66.6 | 61.2 | 59 | 57.8 | 62.1 | 48.6 | 68.6 | 222.6 | 291.7 |
| 854 | N-alpha-acetyltransferase 15, NatA auxiliary subunit OS=Mus musculus OX=10090 GN=Naa15 PE=1 SV=1 | 111.8 | 90.6 | 102.4 | 94.3 | 108.3 | 100.5 | 118.7 | 90.4 | 84.5 | 98.5 |
| 855 | Mannosyl-oligosaccharide glucosidase OS=Mus musculus OX=10090 GN=Mogs PE=1 SV=1 | 128.3 | 108.2 | 119.6 | 93.2 | 100.9 | 109.5 | 118.8 | 79.4 | 76.2 | 65.9 |
| 856 | Citrate synthase OS=Mus musculus OX=10090 GN=Csl PE=1 SV=1 | 112.9 | 107.6 | 108.7 | 87.3 | 103.4 | 101 | 125.2 | 88.1 | 84.8 | 80.9 |
| 857 | Acetyl-Coenzyme A acetyltransferase 3 OS=Mus musculus OX=10090 GN=Acat3 PE=1 SV=1 | 153.2 | 130.6 | 135 | 105.6 | 73.6 | 110.2 | 68.6 | 83.3 | 87.3 | 52.6 |
| 858 | Filamin-B OS=Mus musculus OX=10090 GN=Flnb PE=1 SV=3 | 68.2 | 96.6 | 77.1 | 92 | 94.5 | 100.3 | 174.7 | 103.5 | 93.6 | 99.5 |
| 859 | Bromodomain and PHD finger containing, 1 OS=Mus musculus OX=10090 GN=Brpf1 PE=1 SV=1 | | 96 | | 106.5 | | 110.9 | 289.8 | 112.8 | 125 | 159 |
| 860 | D-beta-hydroxybutyrate dehydrogenase, mitochondrial OS=Mus musculus OX=10090 GN=Bdh1 PE=1 SV=2 | 118 | 107.8 | 125.5 | 114 | 93.8 | 124 | 91.8 | 52.5 | 96.5 | 76.2 |
| 861 | Nuclear ubiquitous casein and cyclin-dependent kinase substrate 1 OS=Mus musculus OX=10090 GN=Nucks1 PE=1 SV=1 | 50.9 | 96.7 | 61.8 | 52.6 | 170 | 101 | 77 | 110.2 | 58.6 | 221.3 |
| 862 | ATP-dependent RNA helicase SUPV3L1, mitochondrial OS=Mus musculus OX=10090 GN=Supv3l1 PE=1 SV=1 | | 96.3 | 118.8 | 105 | 138 | 106.9 | 100.5 | 167 | 84.6 | 83 |
| 863 | Armet protein OS=Mus musculus OX=10090 GN=Manf PE=1 SV=1 | 103.9 | 104.4 | 102.8 | 85.6 | 120.9 | 131.2 | 70.9 | 96 | 78.4 | 105.9 |
| 864 | Proline synthetase co-transcribed OS=Mus musculus OX=10090 GN=Plpbp PE=1 SV=1 | 98.9 | 104.7 | 103.9 | 107.8 | 92.5 | 106 | 93.4 | 98.9 | 101.4 | 92.4 |
| 865 | Iron-responsive element-binding protein 2 OS=Mus musculus OX=10090 GN=Ireb2 PE=1 SV=2 | 134.1 | 124.6 | 126.4 | 99.1 | 67.9 | 117.4 | 54.7 | 103.8 | 81.1 | 90.9 |
| 866 | N-acetylglucosamine-6-sulfatase OS=Mus musculus OX=10090 GN=Gns PE=1 SV=1 | 94 | 109.5 | 98.1 | 97.5 | 106.2 | 109.2 | 92.9 | 106.1 | 94.9 | 91.7 |
| 867 | Elongation factor Tu, mitochondrial OS=Mus musculus OX=10090 GN=Tufm PE=1 SV=1 | 94.1 | 86.4 | 93 | 102.1 | 95.9 | 94.8 | 90.4 | 105.2 | 105.9 | 132.2 |
| 868 | Erlin-2 OS=Mus musculus OX=10090 GN=Erlin2 PE=1 SV=1 | 124.3 | 117.9 | 119.2 | 94 | 103.6 | 119 | 86 | 81.4 | 84.3 | 70.2 |
| 869 | Heterogeneous nuclear ribonucleoprotein A3 OS=Mus musculus OX=10090 GN=Hnrnpa3 PE=1 SV=1 | 101.2 | 98.6 | 96.1 | 79.3 | 95.8 | 110.4 | 61.2 | 103.4 | 96.1 | 158 |
| 870 | Peroxiredoxin-6 OS=Mus musculus OX=10090 GN=Prdx6b PE=1 SV=1 | 133.1 | 131.2 | 122.3 | 107 | 95.4 | 101.5 | 72.6 | 89.1 | 78.1 | 69.6 |
| 871 | Mitochondrial Rho GTPase 1 OS=Mus musculus OX=10090 GN=Rhot1 PE=1 SV=1 | 134.5 | 114.9 | 121.5 | 108.9 | 68.4 | 99.4 | 60.8 | 99.9 | 100.7 | 91 |
| 872 | Acyl-coenzyme A synthetase ACSM5, mitochondrial OS=Mus musculus OX=10090 GN=Acsm5 PE=1 SV=1 | 109.7 | 92.6 | 102.8 | 129.7 | 69.4 | 89 | 50.4 | 114.7 | 132.7 | 109 |
| 873 | Eukaryotic translation initiation factor 4B OS=Mus musculus OX=10090 GN=Eif4b PE=1 SV=1 | 96.6 | 117.9 | 97.5 | 73.7 | 137 | 107.8 | 77.5 | 104.1 | 67.9 | 120 |

| | | | | | | | | | | | |
|---|---|---|---|---|---|---|---|---|---|---|---|
| 874 | Alanine--tRNA ligase, cytoplasmic OS=Mus musculus OX=10090 GN=Aars1 PE=1 SV=1 | 115.7 | 112.6 | 113.6 | 101.4 | 86.5 | 111.1 | 76.3 | 99 | 87.5 | 96.4 |
| 875 | Alanine aminotransferase 2 OS=Mus musculus OX=10090 GN=Gpt2 PE=1 SV=1 | 92.1 | 81.3 | 98.3 | 110 | 93.8 | 89.3 | 58.8 | 119 | 122.3 | 135.2 |
| 876 | Eukaryotic translation initiation factor 5A-2 OS=Mus musculus OX=10090 GN=Eif5a2 PE=1 SV=3 | 88.7 | 89.5 | 96.2 | 88.4 | 161.9 | 98 | 103.7 | 98.4 | 72 | 103.1 |
| 877 | Enoyl-CoA hydratase, mitochondrial OS=Mus musculus OX=10090 GN=Echs1 PE=1 SV=1 | 102.3 | 93.1 | 99.8 | 118.7 | 84.9 | 107.1 | 79.1 | 109 | 103.5 | 102.5 |
| 878 | Elongation of very long chain fatty acids protein 5 OS=Mus musculus OX=10090 GN=Elovl5 PE=1 SV=1 | 179.3 | 146.2 | 154.6 | 77.3 | 124.8 | 130.2 | 50.4 | 52.2 | 48.3 | 36.7 |
| 879 | Neutral alpha-glucosidase AB OS=Mus musculus OX=10090 GN=Ganab PE=1 SV=1 | 133.7 | 116.5 | 131.7 | 84.9 | 96.2 | 120.4 | 90 | 89.1 | 82.3 | 55.2 |
| 880 | Isoleucine--tRNA ligase, mitochondrial OS=Mus musculus OX=10090 GN=Iars2 PE=1 SV=1 | 102.3 | 91.5 | 93.6 | 108.3 | 62.2 | 96.2 | 93.1 | 108.3 | 128.7 | 115.7 |
| 881 | Choline dehydrogenase, mitochondrial OS=Mus musculus OX=10090 GN=Chdh PE=1 SV=1 | 124 | 104.4 | 124.7 | 122 | 79.2 | 96.2 | 47.7 | 115.1 | 113 | 73.7 |
| 882 | Charged multivesicular body protein 2b OS=Mus musculus OX=10090 GN=Chmp2b PE=1 SV=1 | 84.2 | 85.1 | 82.2 | 72.1 | 169 | 88.9 | 176.6 | 85.3 | 63.1 | 93.5 |
| 883 | Eukaryotic translation initiation factor 2A OS=Mus musculus OX=10090 GN=Eif2a PE=1 SV=2 | 81.3 | 93.3 | 93.3 | 70.8 | 65 | 94.9 | 159.5 | 82 | 123.7 | 136.2 |
| 884 | Lon protease homolog 2, peroxisomal OS=Mus musculus OX=10090 GN=Lonp2 PE=1 SV=1 | 139 | 127.2 | 131.5 | 101.9 | 71 | 116.2 | 51.2 | 89 | 95.1 | 77.9 |
| 885 | Early endosome antigen 1 OS=Mus musculus OX=10090 GN=Eea1 PE=1 SV=2 | 50.8 | 106.4 | 53.1 | 99.5 | 81.1 | 121.3 | 137 | 127.1 | 96 | 127.9 |
| 886 | Lanosterol synthase OS=Mus musculus OX=10090 GN=Lss PE=1 SV=2 | 106.2 | 82.6 | 94.7 | 98.5 | 79.9 | 88.6 | 66.7 | 154 | 103.3 | 125.4 |
| 887 | Dihydrolipoyllysine-residue acetyltransferase component of pyruvate dehydrogenase complex, mitochondrial OS=Mus musculus OX=10090 GN=Dlat PE=1 SV=2 | 99.5 | 102.1 | 105.6 | 91.1 | 123 | 104.8 | 132.9 | 85.7 | 74.9 | 80.4 |
| 888 | Glutamine--tRNA ligase OS=Mus musculus OX=10090 GN=Qars1 PE=1 SV=1 | 138.4 | 105.4 | 123.5 | 93.4 | 87.4 | 129.7 | 63.9 | 97.5 | 87.5 | 73.3 |
| 889 | C-type lectin domain family 4 member G OS=Mus musculus OX=10090 GN=Clec4g PE=1 SV=1 | 171.7 | 159.1 | 155.2 | 67.6 | 98 | 112.5 | 78.2 | 47.3 | 75.8 | 34.7 |
| 890 | 60S ribosomal protein L24 OS=Mus musculus OX=10090 GN=Rpl24 PE=1 SV=2 | 168.1 | 85 | 144.1 | 86.8 | 55.9 | 94.3 | 56.9 | 107.8 | 97 | 104 |
| 891 | Alkylglycerol monooxygenase OS=Mus musculus OX=10090 GN=Agmo PE=1 SV=1 | 140.6 | 113.7 | 110.1 | 112.7 | 71.3 | 104.6 | 81.2 | 96.8 | 92.2 | 76.9 |
| 892 | Copine-3 OS=Mus musculus OX=10090 GN=Cpne3 PE=1 SV=2 | 145 | 108.6 | 121.2 | 97.9 | 83.2 | 113.8 | 62 | 95.9 | 86.8 | 85.7 |
| 893 | Filamin-A OS=Mus musculus OX=10090 GN=Flna PE=1 SV=5 | | 168.9 | | 118.7 | | 159.8 | 164.2 | 123.4 | 124.5 | 140.6 |
| 894 | DEAD box protein 5 OS=Mus musculus OX=10090 GN=Ddx5 PE=1 SV=1 | 87.9 | 104.5 | 88.9 | 91.5 | 93.5 | 108 | 118.3 | 96 | 87.7 | 123.7 |
| 895 | Isoleucine--tRNA ligase, cytoplasmic OS=Mus musculus OX=10090 GN=Iars1 PE=1 SV=2 | 149.6 | 113.8 | 124.2 | 102.1 | 78.3 | 111.5 | 61.5 | 90.9 | 87.6 | 80.4 |
| 896 | Complex I assembly factor TIMMDC1, mitochondrial OS=Mus musculus OX=10090 GN=Timmdc1 PE=1 SV=1 | 99.2 | 93.5 | 109.9 | 112.1 | 72.9 | 94.4 | 81.5 | 117.2 | 104.3 | 114.9 |
| 897 | Eukaryotic translation initiation factor 5 OS=Mus musculus OX=10090 GN=Eif5 PE=1 SV=1 | 124.2 | 105 | 107.5 | 97.9 | 78.5 | 120.1 | 63.7 | 113.9 | 89.5 | 99.9 |
| 898 | 3-ketoacyl-CoA thiolase, mitochondrial OS=Mus musculus OX=10090 GN=Acaa2 PE=1 SV=3 | 117.5 | 106.4 | 123.7 | 106.4 | 71.1 | 102.7 | 75.3 | 92.1 | 107.1 | 97.8 |
| 899 | Eukaryotic peptide chain release factor subunit 1 OS=Mus musculus OX=10090 GN=Etf1 PE=1 SV=4 | 131.9 | 88.5 | 114.6 | 98.3 | 87.6 | 111.1 | 95.3 | 80.4 | 99.7 | 92.6 |
| 900 | Dynamin-3 OS=Mus musculus OX=10090 GN=Dnm3 PE=1 SV=1 | 135.8 | 109.4 | 124 | 84.5 | 87.1 | 115.8 | 63.7 | 102.9 | 80.3 | 96.6 |
| 901 | Isoaspartyl peptidase/L-asparaginase OS=Mus musculus OX=10090 GN=Asrgl1 PE=1 SV=1 | 108.7 | 124.7 | 114.8 | 99.6 | 102.5 | 103.6 | 90.1 | 81.5 | 80.6 | 94 |
| 902 | N-fatty-acyl-amino acid synthase/hydrolase PM20D1 OS=Mus musculus OX=10090 GN=Pm20d1 PE=1 SV=1 | 142.3 | 100.6 | 105.4 | 112.4 | 80.1 | 124.3 | 72.8 | 96.6 | 87.4 | 77.9 |
| 903 | Carbamoyl-phosphate synthase [ammonia], mitochondrial OS=Mus musculus OX=10090 GN=Cps1 PE=1 SV=2 | 98.8 | 86.9 | 82.6 | 144.5 | 60.9 | 92.7 | 65.8 | 135.2 | 122.8 | 109.9 |
| 904 | NAD kinase 2, mitochondrial OS=Mus musculus OX=10090 GN=Nadk2 PE=1 SV=2 | 115.5 | 94.9 | 111.4 | 103.7 | 82.4 | 104.5 | 73.4 | 96.6 | 98.1 | 119.6 |
| 905 | Flavin-containing monooxygenase OS=Mus musculus OX=10090 GN=Fmo1 PE=1 SV=1 | 90.9 | 114.4 | 133.4 | 116.2 | 66.6 | 100.9 | 71.5 | 100.9 | 112.3 | 92.9 |
| 906 | N-acetylgalactosamine-6-sulfatase OS=Mus musculus OX=10090 GN=Galns PE=1 SV=1 | | 133.1 | | 123.8 | | 158 | 96.5 | 154 | 152 | 182.5 |
| 907 | Nicotinate phosphoribosyltransferase OS=Mus musculus OX=10090 GN=Naprt PE=1 SV=1 | 123.4 | 106.2 | 115.7 | 99 | 92.1 | 115.2 | 86.5 | 98.8 | 82.6 | 80.5 |
| 908 | E3 UFM1-protein ligase 1 OS=Mus musculus OX=10090 GN=Ufl1 PE=1 SV=2 | 107.9 | 104.5 | 106.4 | 86.2 | 103.4 | 110.1 | 118.9 | 99.8 | 78.3 | 84.4 |
| 909 | Core histone macro-H2A.2 OS=Mus musculus OX=10090 GN=Macroh2a2 PE=1 SV=3 | 125.5 | 107.7 | 125.1 | 89.8 | 59.7 | 114.6 | 70.1 | 106.4 | 94.9 | 106.2 |

| | | | | | | | | | | | |
|---|---|---|---|---|---|---|---|---|---|---|---|
| 910 | Aldo-keto reductase family 1, member E1 OS=Mus musculus OX=10090 GN=Akr1e1 PE=1 SV=1 | 114.2 | 119.4 | 124.8 | 100.4 | 90.4 | 130.6 | 79.3 | 78.5 | 91.1 | 71.4 |
| 911 | BolA-like protein 3 OS=Mus musculus OX=10090 GN=Bola3 PE=1 SV=1 | 72.7 | 83 | 67.9 | 82.9 | 163.3 | 84.4 | 92.2 | 130.9 | 79.3 | 143.5 |
| 912 | Aminomethyltransferase, mitochondrial OS=Mus musculus OX=10090 GN=Amt PE=1 SV=1 | 102.9 | 98.3 | 94.7 | 127 | 70.1 | 101.3 | 49.6 | 120.5 | 132.9 | 102.8 |
| 913 | GDH/6PGL endoplasmic bifunctional protein OS=Mus musculus OX=10090 GN=H6pd PE=1 SV=2 | 113.7 | 93.9 | 104.4 | 122.6 | 64.1 | 117.7 | 59.6 | 128.1 | 119.9 | 76.1 |
| 914 | Aflatoxin B1 aldehyde reductase member 2 OS=Mus musculus OX=10090 GN=Akr7a2 PE=1 SV=3 | 118 | 111.3 | 102.3 | 115.1 | 82 | 92.8 | 75.9 | 96.3 | 106.2 | 100 |
| 915 | Bifunctional glutamate/proline--tRNA ligase OS=Mus musculus OX=10090 GN=Eprs1 PE=1 SV=4 | 117.5 | 100.2 | 100.6 | 103.6 | 78.6 | 134.5 | 83.1 | 90.1 | 91.5 | 100.3 |
| 916 | Lon protease homolog, mitochondrial OS=Mus musculus OX=10090 GN=Lonp1 PE=1 SV=2 | 98.5 | 104 | 99.3 | 98 | 101.9 | 120.6 | 70.2 | 104.1 | 102.6 | 101 |
| 917 | Histone H2B type 3-B OS=Mus musculus OX=10090 GN=H2bu1 PE=1 SV=3 | 82.8 | 82.2 | 87.5 | 62.9 | 170.9 | 182.5 | 109.9 | 68.2 | 63.5 | 89.7 |
| 918 | Cell division cycle and apoptosis regulator protein 1 OS=Mus musculus OX=10090 GN=Ccar1 PE=1 SV=1 | 106.6 | 99.8 | 110.9 | 86.4 | 97.3 | 100.4 | 83.6 | 88.3 | 90.9 | 135.9 |
| 919 | N-acetyltransferase family 8 member 2 OS=Mus musculus OX=10090 GN=Nat8f2 PE=1 SV=1 | 120.6 | 115.8 | 126.2 | 106.5 | 76.7 | 111.2 | 67.7 | 105.4 | 97.8 | 72.3 |
| 920 | Dihydropyrimidine dehydrogenase [NADP(+)] OS=Mus musculus OX=10090 GN=Dpyd PE=1 SV=1 | 99.3 | 97.1 | 104.9 | 123.2 | 86 | 135.2 | 88.4 | 100.1 | 82.5 | 83.3 |
| 921 | Delta-1-pyrroline-5-carboxylate dehydrogenase, mitochondrial OS=Mus musculus OX=10090 GN=Aldh4a1 PE=1 SV=3 | 100.7 | 93 | 106 | 124 | 82.5 | 98.6 | 77.9 | 116.3 | 108.4 | 92.6 |
| 922 | Hydroxymethylglutaryl-CoA synthase, cytoplasmic OS=Mus musculus OX=10090 GN=Hmgcs1 PE=1 SV=1 | 103.1 | 94.1 | 97.2 | 110.7 | 84.3 | 99.9 | 102.3 | 115.7 | 102.2 | 90.6 |
| 923 | Cold shock domain-containing protein E1 OS=Mus musculus OX=10090 GN=Csde1 PE=1 SV=1 | 21 | 140.3 | 18.2 | 123.7 | 12.9 | 134.9 | 88 | 142.2 | 135.9 | 183 |
| 924 | Complex I assembly factor ACAD9, mitochondrial OS=Mus musculus OX=10090 GN=Acad9 PE=1 SV=2 | 111.7 | 93.3 | 113.5 | 104.4 | 86.3 | 103.3 | 78.6 | 111.5 | 100.3 | 97 |
| 925 | AFG3-like protein 2 OS=Mus musculus OX=10090 GN=Afg3l2 PE=1 SV=1 | 91.9 | 68.3 | 79.5 | 142.7 | 58.4 | 114.8 | 116.2 | 78.5 | 127 | 122.7 |
| 926 | Eukaryotic translation initiation factor 3 subunit B OS=Mus musculus OX=10090 GN=Eif3b PE=1 SV=1 | 101.3 | 93.1 | 80.7 | 89.2 | 110.1 | 107.2 | 140.7 | 109.5 | 86.1 | 82 |
| 927 | Long-chain-fatty-acid--CoA ligase 5 OS=Mus musculus OX=10090 GN=Acsl5 PE=1 SV=1 | 109.6 | 101.1 | 110.3 | 86.3 | 92.7 | 89.6 | 81.5 | 84.1 | 68.7 | 176.2 |
| 928 | Phosphatidylinositol transfer protein beta isoform OS=Mus musculus OX=10090 GN=Pitpnb PE=1 SV=1 | 106.7 | 109.5 | 115.9 | 97.2 | 91.8 | 110.3 | 106.2 | 95.3 | 86.7 | 80.3 |
| 929 | Mitochondrial 10-formyltetrahydrofolate dehydrogenase OS=Mus musculus OX=10090 GN=Aldh1l2 PE=1 SV=2 | 104.8 | 93.9 | 92.2 | 137 | 75.2 | 111 | 65.3 | 132.3 | 105 | 83.4 |
| 930 | 5-oxoprolinase OS=Mus musculus OX=10090 GN=Oplah PE=1 SV=1 | 113 | 108.9 | 117.4 | 99.7 | 92.2 | 107.7 | 84.9 | 96.6 | 92.6 | 86.8 |
| 931 | Lanosterol 14-alpha demethylase OS=Mus musculus OX=10090 GN=Cyp51a1 PE=1 SV=1 | 115.2 | 105.7 | 105.3 | 111.5 | 88.2 | 90.7 | 86.3 | 108.8 | 108.5 | 79.9 |
| 932 | Elongation factor G, mitochondrial OS=Mus musculus OX=10090 GN=Gfm1 PE=1 SV=1 | 80.8 | 93.9 | 79.8 | 79.6 | 89.8 | 89.1 | 80.5 | 135.9 | 105 | 165.6 |
| 933 | Fibrinogen beta chain OS=Mus musculus OX=10090 GN=Fgb PE=1 SV=1 | 85.3 | 91.7 | 115.4 | 81.9 | 88.1 | 90.2 | 97.7 | 122.3 | 99.2 | 128.2 |
| 934 | Galactose mutarotase OS=Mus musculus OX=10090 GN=Galm PE=1 SV=1 | 121.8 | 112.1 | 122.9 | 84.4 | 99 | 109.4 | 89.2 | 106.4 | 79.1 | 75.7 |
| 935 | Dynamin-1-like protein OS=Mus musculus OX=10090 GN=Dnm1l PE=1 SV=2 | 101.7 | 91.5 | 95.4 | 67.3 | 84.7 | 94.2 | 130.4 | 76 | 138 | 120.8 |
| 936 | Polyribonucleotide nucleotidyltransferase 1, mitochondrial OS=Mus musculus OX=10090 GN=Pnpt1 PE=1 SV=1 | 69.1 | 47.9 | 66.7 | 141.6 | 38.6 | 135.8 | 132 | 57.8 | 140.6 | 169.9 |
| 937 | LYR motif-containing protein 4 OS=Mus musculus OX=10090 GN=Lyrm4 PE=1 SV=1 | 121.8 | 90.1 | 111 | 106.2 | 65.1 | 103.2 | 75 | 113.2 | 111.8 | 102.6 |
| 938 | Matrin-3 OS=Mus musculus OX=10090 GN=Matr3 PE=1 SV=1 | 91.5 | 94.3 | 88.2 | 74.2 | 122.9 | 106.7 | 109.8 | 105.7 | 79.7 | 127 |
| 939 | Carbonyl reductase [NADPH] 3 OS=Mus musculus OX=10090 GN=Cbr3 PE=1 SV=1 | 110.5 | 92.2 | 83.1 | 116.3 | 84.8 | 86.9 | 33.4 | 138.5 | 121.2 | 133.1 |
| 940 | Acyl-CoA dehydrogenase family member 10 OS=Mus musculus OX=10090 GN=Acad10 PE=1 SV=1 | 98.2 | 103.2 | 91.2 | 104.1 | 89.1 | 102.6 | 95.8 | 116.6 | 106.9 | 92.3 |
| 941 | NADH dehydrogenase [ubiquinone] iron-sulfur protein 8, mitochondrial OS=Mus musculus OX=10090 GN=Ndufs8 PE=1 SV=1 | 29.3 | 25.8 | 26.4 | 36 | 25.3 | 33.2 | 293.8 | 34.1 | 231.7 | 264.5 |
| 942 | Cis-retinol/3alpha hydroxysterol short-chain dehydrogenase-like protein OS=Mus musculus OX=10090 GN=Rdh16f2 PE=1 SV=1 | 144.1 | 128.1 | 136.1 | 113.9 | 55.2 | 94.9 | 56.2 | 101.3 | 113.2 | 57 |
| 943 | Presequence protease, mitochondrial OS=Mus musculus OX=10090 GN=Pitrm1 PE=1 SV=1 | 84.8 | 71.8 | 92.6 | 92.6 | 63 | 86.5 | 118.2 | 85.5 | 155.5 | 149.5 |
| 944 | ATP-binding cassette sub-family A member 6 OS=Mus musculus OX=10090 GN=Abca6 PE=1 SV=2 | 109 | 98.9 | 102.7 | 93.8 | 87.3 | 104.3 | 108.9 | 99.7 | 88.4 | 107 |

| | | | | | | | | | | | |
|---|---|---|---|---|---|---|---|---|---|---|---|
| 945 | Kynurenine formamidase OS=Mus musculus OX=10090 GN=Afmid PE=1 SV=1 | | 146.2 | | 171.4 | | 165.8 | 100.9 | 130.2 | 148.5 | 136.9 |
| 946 | Cytoplasmic phosphatidylinositol transfer protein 1 OS=Mus musculus OX=10090 GN=Pitpnc1 PE=1 SV=1 | | 157.5 | | 143.4 | | 167 | 144.4 | 137.7 | 128.1 | 121.9 |
| 947 | Activator of 90 kDa heat shock protein ATPase homolog 2 OS=Mus musculus OX=10090 GN=Ahsa2 PE=1 SV=2 | 112.4 | 103.1 | 107.1 | 91 | 112.1 | 92.8 | 125.7 | 93.1 | 80.1 | 82.5 |
| 948 | Alanine aminotransferase 1 OS=Mus musculus OX=10090 GN=Gpt PE=1 SV=3 | 101.9 | 112.7 | 103.5 | 121 | 69.8 | 108.3 | 87.9 | 94.9 | 103.2 | 96.6 |
| 949 | 3-hydroxyisobutyryl-CoA hydrolase, mitochondrial OS=Mus musculus OX=10090 GN=Hibch PE=1 SV=1 | 86.5 | 67 | 73.5 | 71.2 | 58 | 63 | 116.7 | 80.4 | 202.5 | 181.3 |
| 950 | Acetyl-CoA acetyltransferase, mitochondrial OS=Mus musculus OX=10090 GN=Acat1 PE=1 SV=1 | 131 | 129.2 | 122.1 | 90.4 | 95.1 | 103.9 | 99.9 | 69.8 | 89.5 | 69.1 |
| 951 | Eukaryotic translation initiation factor 3 subunit L OS=Mus musculus OX=10090 GN=Eif3l PE=1 SV=1 | 114.2 | 105.7 | 100.4 | 77.5 | 118.9 | 108.2 | 101.2 | 97.4 | 86.1 | 90.3 |
| 952 | Glycerate kinase OS=Mus musculus OX=10090 GN=Glyctk PE=1 SV=1 | 133.1 | 122.2 | 129.2 | 106.2 | 72.5 | 121.2 | 55.5 | 90.8 | 93.3 | 75.9 |
| 953 | Eukaryotic peptide chain release factor GTP-binding subunit ERF3A OS=Mus musculus OX=10090 GN=Gspt1 PE=1 SV=2 | 112.4 | 109.5 | 116.9 | 87.5 | 92.8 | 114.4 | 85.6 | 99.5 | 92.2 | 89.3 |
| 954 | Heterogeneous nuclear ribonucleoprotein L OS=Mus musculus OX=10090 GN=Hnrnpl PE=1 SV=2 | 77.9 | 108 | 97.9 | 74.1 | 143.5 | 143.4 | 69.9 | 88.3 | 77.2 | 119.8 |
| 955 | Acylpyruvase FAHD1, mitochondrial OS=Mus musculus OX=10090 GN=Fahd1 PE=1 SV=2 | 120.3 | 111.8 | 109.1 | 103 | 80.5 | 106.2 | 76.3 | 102.4 | 91.2 | 99.3 |
| 956 | Hydroxyacid-oxoacid transhydrogenase, mitochondrial OS=Mus musculus OX=10090 GN=Adhfe1 PE=1 SV=2 | 99.1 | 99.2 | 103.7 | 117.8 | 88.3 | 94.4 | 84.9 | 109.1 | 95.5 | 108.1 |
| 957 | Epiplakin OS=Mus musculus OX=10090 GN=Eppk1 PE=1 SV=2 | 66.4 | 66.6 | 70.1 | 98.9 | 89.3 | 89.3 | 75.6 | 117.2 | 121.1 | 205.5 |
| 958 | Cytosolic 10-formyltetrahydrofolate dehydrogenase OS=Mus musculus OX=10090 GN=Aldh1l1 PE=1 SV=1 | 118.1 | 94.9 | 107.8 | 123.4 | 78.6 | 111.4 | 70.9 | 117.8 | 102.3 | 74.9 |
| 959 | Mitochondrial coenzyme A transporter SLC25A42 OS=Mus musculus OX=10090 GN=Slc25a42 PE=1 SV=1 | 94.1 | 76.7 | 107.9 | 76.7 | 90.4 | 113.1 | 95 | 92.5 | 113.3 | 140.3 |
| 960 | Acylamino-acid-releasing enzyme OS=Mus musculus OX=10090 GN=Apeh PE=1 SV=3 | 128.5 | 112.5 | 121.8 | 86 | 110.9 | 122 | 90.2 | 79.5 | 78.9 | 69.6 |
| 961 | Eukaryotic translation initiation factor 3 subunit C OS=Mus musculus OX=10090 GN=Eif3c PE=1 SV=1 | 129.1 | 104.7 | 121.8 | 89.6 | 80.4 | 111.4 | 100.3 | 90.3 | 96.9 | 75.4 |
| 962 | Carboxymethylenebutenolidase homolog OS=Mus musculus OX=10090 GN=Cmbl PE=1 SV=1 | 84.3 | 110.1 | 130.5 | 142.2 | 62.1 | 87.2 | 74.4 | 89.5 | 138 | 81.7 |
| 963 | 5-formyltetrahydrofolate cyclo-ligase OS=Mus musculus OX=10090 GN=Mthfsl PE=1 SV=1 | 99.7 | 107.1 | 89.7 | 112.3 | 61.4 | 99 | 62.4 | 135.4 | 111.2 | 121.9 |
| 964 | Peptidyl-tRNA hydrolase 2, mitochondrial OS=Mus musculus OX=10090 GN=Ptrh2 PE=1 SV=1 | 132.8 | 109.6 | 121.7 | 109 | 74 | 102.2 | 59.3 | 101.3 | 102 | 88.2 |
| 965 | Malonyl-CoA-acyl carrier protein transacylase, mitochondrial OS=Mus musculus OX=10090 GN=Mcat PE=1 SV=3 | 109.3 | 95.3 | 105.1 | 103.1 | 86.7 | 99.8 | 72 | 110.3 | 104.1 | 114.3 |
| 966 | MICOS complex subunit MIC13 OS=Mus musculus OX=10090 GN=Micos13 PE=1 SV=1 | 88.3 | 87 | 91 | 102 | 102.8 | 91.2 | 124.9 | 104.6 | 102.8 | 105.3 |
| 967 | N-acetylglutamate synthase, mitochondrial OS=Mus musculus OX=10090 GN=Nags PE=1 SV=2 | 126 | 112.4 | 108.4 | 101.7 | 83.4 | 111.2 | 70.3 | 104.3 | 89.7 | 92.7 |
| 968 | Citramalyl-CoA lyase, mitochondrial OS=Mus musculus OX=10090 GN=Clybl PE=1 SV=2 | 99 | 95.8 | 81.1 | 105.2 | 80.3 | 90.2 | 79.5 | 129.7 | 125.8 | 113.5 |
| 969 | Glutathione S-transferase Mu 4 OS=Mus musculus OX=10090 GN=Gstm4 PE=1 SV=1 | 109.3 | 80.2 | 108.2 | 141.4 | 66.8 | 92.6 | 38.9 | 136.1 | 137.9 | 88.5 |
| 970 | PRA1 family protein 3 OS=Mus musculus OX=10090 GN=Arl6ip5 PE=1 SV=2 | 102.6 | 100.6 | 96.4 | 79.3 | 121 | 99.5 | 115.9 | 107 | 86.2 | 91.6 |
| 971 | L-serine dehydratase/L-threonine deaminase OS=Mus musculus OX=10090 GN=Sds PE=1 SV=3 | 99.7 | 93.4 | 98.6 | 108.7 | 80.4 | 106.6 | 77.8 | 115.4 | 106 | 113.4 |
| 972 | Aldo-keto reductase family 1 member C13 OS=Mus musculus OX=10090 GN=Akr1c13 PE=1 SV=2 | 136 | 126.5 | 126.3 | 95.1 | 78.5 | 108.5 | 65.9 | 84.5 | 94.1 | 84.6 |
| 973 | Beta-ureidopropionase OS=Mus musculus OX=10090 GN=Upb1 PE=1 SV=1 | 90.3 | 78.9 | 77.2 | 141.4 | 65 | 77.2 | 18.8 | 157.4 | 155.2 | 138.6 |
| 974 | Glycogen [starch] synthase, liver OS=Mus musculus OX=10090 GN=Gys2 PE=1 SV=2 | 115 | 94.7 | 122.2 | 129.2 | 78.6 | 93.7 | 57.8 | 106.5 | 86.6 | 115.5 |
| 975 | Liver carboxylesterase 1 OS=Mus musculus OX=10090 GN=Ces1 PE=1 SV=1 | 124 | 99.3 | 134.1 | 115.9 | 97 | 110.2 | 55.7 | 92.6 | 102.1 | 69.1 |
| 976 | 3-ketoacyl-CoA thiolase B, peroxisomal OS=Mus musculus OX=10090 GN=Acaa1b PE=1 SV=1 | 139.1 | 101.8 | 116.1 | 97.1 | 113.7 | 120.8 | 44.6 | 109.1 | 84.2 | 73.5 |
| 977 | Delta(24)-sterol reductase OS=Mus musculus OX=10090 GN=Dhcr24 PE=1 SV=1 | 127.9 | 109.6 | 121.2 | 107.9 | 81.2 | 114.5 | 72.4 | 98 | 95.9 | 71.3 |
| 978 | Cystathionine gamma-lyase OS=Mus musculus OX=10090 GN=Cth PE=1 SV=1 | 69.7 | 66 | 71.1 | 164.3 | 52.9 | 83 | 50.7 | 163.4 | 158.2 | 120.8 |
| 979 | Carboxylesterase 1D OS=Mus musculus OX=10090 GN=Ces1d PE=1 SV=1 | 162.9 | 137.4 | 159.9 | 90.8 | 83.6 | 122.5 | 59.7 | 69.2 | 76.2 | 37.8 |
| 980 | Medium-chain acyl-CoA ligase ACSF2, mitochondrial OS=Mus musculus OX=10090 GN=Acsf2 PE=1 SV=1 | 101.9 | 90.8 | 100.9 | 96.5 | 103.8 | 89.1 | 103.8 | 110.1 | 107.4 | 95.7 |

| | | | | | | | | | | | |
|---|---|---|---|---|---|---|---|---|---|---|---|
| 981 | Aldo-keto reductase family 1 member D1 OS=Mus musculus OX=10090 GN=Akr1d1 PE=1 SV=1 | 110.7 | 92.4 | 110 | 127.8 | 95 | 106 | 102.5 | 97.4 | 80.3 | 77.9 |
| 982 | Hydroxyproline dehydrogenase OS=Mus musculus OX=10090 GN=Prodh2 PE=1 SV=1 | 115.1 | 102.9 | 102.8 | 116.6 | 69.8 | 105.2 | 75 | 114.3 | 112 | 86.4 |
| 983 | Probable leucine--tRNA ligase, mitochondrial OS=Mus musculus OX=10090 GN=Lars2 PE=1 SV=1 | 101.8 | 93 | 103.7 | 108 | 48.2 | 96 | 87.9 | 124.9 | 114.7 | 121.7 |
| 984 | Myosin-9 OS=Mus musculus OX=10090 GN=Myh9 PE=1 SV=4 | 24.7 | 75.7 | 82.4 | 126 | 52.9 | 172.4 | 120.8 | 60.4 | 141.1 | 143.6 |
| 985 | Deaminated glutathione amidase OS=Mus musculus OX=10090 GN=Nit1 PE=1 SV=2 | 87.5 | 80.7 | 101.5 | 92.4 | 89.2 | 107.5 | 166.5 | 93.6 | 84 | 97.2 |
| 986 | Mitochondrial intermembrane space import and assembly protein 40 OS=Mus musculus OX=10090 GN=Chchd4 PE=1 SV=1 | 81.7 | 76.7 | 78.8 | 97.8 | 104 | 83.4 | 86.1 | 132.2 | 147.4 | 111.9 |
| 987 | Heterogeneous nuclear ribonucleoprotein U OS=Mus musculus OX=10090 GN=Hnrnpu PE=1 SV=1 | 135.1 | 100.5 | 104.9 | 92 | 73.7 | 123 | 67.9 | 110.3 | 96 | 96.6 |
| 988 | Phosphate carrier protein, mitochondrial OS=Mus musculus OX=10090 GN=Slc25a3 PE=1 SV=1 | 98 | 71.2 | 106.4 | 107 | 85 | 78.8 | 142.6 | 106 | 118.4 | 86.6 |
| 989 | Dimethylaniline monooxygenase [N-oxide-forming] 4 OS=Mus musculus OX=10090 GN=Fmo4 PE=1 SV=3 | 81.7 | 94.6 | 93.1 | 101 | 60.5 | 67.4 | 118.4 | 149.8 | 126.8 | 106.8 |
| 990 | ATP-binding cassette sub-family C member 2 OS=Mus musculus OX=10090 GN=Abcc2 PE=1 SV=2 | 97.2 | 90 | 94.4 | 103.6 | 89.1 | 87.6 | 74.1 | 116.4 | 110.2 | 137.4 |
| 991 | Lysophospholipid acyltransferase 5 OS=Mus musculus OX=10090 GN=Lpcat3 PE=1 SV=1 | 131.2 | 106.8 | 122.2 | 104.6 | 76.1 | 103.1 | 81.6 | 112.4 | 86.1 | 76 |
| 992 | Isochorismatase domain-containing protein 1 OS=Mus musculus OX=10090 GN=Isoc1 PE=1 SV=1 | 127.1 | 110.5 | 132.7 | 94.7 | 89.9 | 118.9 | 76.3 | 99 | 75.3 | 75.6 |
| 993 | Ester hydrolase C11orf54 homolog OS=Mus musculus OX=10090 PE=1 SV=1 | 110.6 | 109.5 | 115.9 | 101.1 | 84.7 | 88.7 | 91.1 | 101.5 | 104.7 | 92.3 |
| 994 | ATP-citrate synthase OS=Mus musculus OX=10090 GN=Acly PE=1 SV=1 | 197.8 | 113.8 | 151.4 | 69.9 | 111.7 | 125.1 | 79.8 | 54.2 | 50.9 | 45.5 |
| 995 | Acyl-coenzyme A synthetase ACSM1, mitochondrial OS=Mus musculus OX=10090 GN=Acsm1 PE=1 SV=1 | 128 | 119.3 | 112.6 | 98.1 | 78.9 | 92.8 | 58.8 | 120.5 | 100.2 | 90.7 |
| 996 | Polymerase delta-interacting protein 2 OS=Mus musculus OX=10090 GN=Poldip2 PE=1 SV=1 | 77.2 | 81.1 | 71.3 | 79.5 | 128.2 | 69.1 | 248.2 | 80.7 | 76.4 | 88.4 |
| 997 | Alpha globin 1 OS=Mus musculus OX=10090 GN=Hba-a1 PE=1 SV=1 | 98.4 | 106.6 | 190.8 | 82.3 | 70.9 | 86 | 48.2 | 110.8 | 113.2 | 92.8 |
| 998 | NADH-ubiquinone oxidoreductase 75 kDa subunit, mitochondrial OS=Mus musculus OX=10090 GN=Ndufs1 PE=1 SV=2 | 107 | 80.6 | 77.3 | 102.9 | 87.9 | 86.1 | 131.5 | 112.5 | 105.2 | 108.9 |
| 999 | Histamine N-methyltransferase OS=Mus musculus OX=10090 GN=Hnmt PE=1 SV=1 | 100.4 | 92.6 | 104.9 | 102 | 110.3 | 97.8 | 127.7 | 91.1 | 82 | 91.2 |
| 1000 | Eef1d protein OS=Mus musculus OX=10090 GN=Eef1d PE=1 SV=1 | 110.1 | 110.3 | 107.5 | 79.9 | 136.5 | 127.9 | 66.6 | 91 | 76.1 | 94.2 |
| 1001 | ATP synthase subunit gamma, mitochondrial OS=Mus musculus OX=10090 GN=Atp5f1c PE=1 SV=1 | 89.2 | 97.3 | 92.3 | 113.1 | 99.3 | 104.3 | 99.1 | 116.8 | 94 | 94.7 |
| 1002 | 3-oxoacyl-[acyl-carrier-protein] reductase OS=Mus musculus OX=10090 GN=Cbr4 PE=1 SV=2 | 105 | 89.7 | 97.3 | 108.7 | 76.7 | 95.7 | 85.1 | 109.6 | 113.8 | 118.4 |
| 1003 | DDB1- and CUL4-associated factor 11 OS=Mus musculus OX=10090 GN=Dcaf11 PE=1 SV=1 | 95.1 | 104.8 | 102 | 86.7 | 129.9 | 105.9 | 110.7 | 100.8 | 82.9 | 81.2 |
| 1004 | Glycine dehydrogenase (decarboxylating), mitochondrial OS=Mus musculus OX=10090 GN=Gldc PE=1 SV=1 | 95.8 | 100.6 | 97.5 | 111.1 | 84.4 | 91.8 | 55.3 | 146.8 | 121.7 | 95 |
| 1005 | Cytochrome P450 2C70 OS=Mus musculus OX=10090 GN=Cyp2c70 PE=1 SV=2 | 145 | 123.3 | 138.5 | 66.1 | 91.2 | 132 | 65.5 | 74.2 | 84.6 | 79.7 |
| 1006 | NADH dehydrogenase [ubiquinone] iron-sulfur protein 2, mitochondrial OS=Mus musculus OX=10090 GN=Ndufs2 PE=1 SV=1 | 95 | 84.8 | 94.8 | 109.4 | 86.1 | 88.1 | 129.5 | 108.3 | 105.4 | 98.7 |
| 1007 | Far upstream element-binding protein 1 OS=Mus musculus OX=10090 GN=Fubp1 PE=1 SV=1 | 91 | 102.3 | 86.7 | 72.1 | 138.9 | 109.4 | 81 | 109.2 | 72.3 | 137.1 |
| 1008 | Eukaryotic translation initiation factor 3 subunit H OS=Mus musculus OX=10090 GN=Eif3h PE=1 SV=1 | 124.7 | 109 | 110.3 | 86.7 | 80.4 | 111.7 | 72 | 111.3 | 89.5 | 104.4 |
| 1009 | Haloacid dehalogenase-like hydrolase domain-containing 5 OS=Mus musculus OX=10090 GN=Hdhd5 PE=1 SV=1 | 99.8 | 109.8 | 110.7 | 100.8 | 82.7 | 97.2 | 99.4 | 106.5 | 89.8 | 103.3 |
| 1010 | Cystathionine beta-synthase OS=Mus musculus OX=10090 GN=Cbs PE=1 SV=3 | 127.7 | 114.3 | 123.3 | 102.3 | 73.3 | 110 | 78.4 | 92.8 | 92.7 | 85 |
| 1011 | Carboxylesterase 1F OS=Mus musculus OX=10090 GN=Ces1f PE=1 SV=1 | 138.6 | 140.7 | 168.5 | 92.1 | 87.9 | 104.2 | 71.7 | 78.5 | 72.4 | 45.5 |
| 1012 | Arsenite methyltransferase OS=Mus musculus OX=10090 GN=As3mt PE=1 SV=2 | 135 | 111 | 122.7 | 91.7 | 90.4 | 128.9 | 65 | 97.5 | 80.9 | 76.9 |
| 1013 | Bile acid-CoA:amino acid N-acyltransferase OS=Mus musculus OX=10090 GN=Baat PE=1 SV=1 | 121 | 95 | 86.3 | 96.5 | 75.9 | 106.4 | 59.4 | 129.6 | 114.8 | 115.2 |
| 1014 | Glucokinase regulatory protein OS=Mus musculus OX=10090 GN=Gckr PE=1 SV=3 | 130.5 | 114.3 | 117.8 | 82.9 | 113 | 110.5 | 72.1 | 91.3 | 86.5 | 81 |

| | | | | | | | | | | | |
|---|---|---|---|---|---|---|---|---|---|---|---|
| 1015 | Hemopexin OS=Mus musculus OX=10090 GN=Hpx PE=1 SV=2 | 82 | 75.5 | 104.5 | 71.7 | 101.5 | 110.8 | 119.1 | 73.1 | 81.8 | 180.1 |
| 1016 | Cytochrome P450 2C50 OS=Mus musculus OX=10090 GN=Cyp2c50 PE=1 SV=2 | 126.8 | 128.7 | 120.7 | 120.3 | 70.8 | 93.7 | 61.3 | 120.2 | 111 | 46.6 |
| 1017 | Nicotinate-nucleotide pyrophosphorylase [carboxylating] OS=Mus musculus OX=10090 GN=Qprt PE=1 SV=1 | 139.1 | 121 | 134.5 | 87 | 98.1 | 157.8 | 63.8 | 71.9 | 69.7 | 57 |
| 1018 | Formimidoyltransferase-cyclodeaminase OS=Mus musculus OX=10090 GN=Ftcd PE=1 SV=1 | 108.9 | 87.1 | 114.7 | 125 | 66.3 | 105.6 | 55.8 | 102.1 | 124.5 | 110.1 |
| 1019 | Glycine N-acyltransferase OS=Mus musculus OX=10090 GN=Glyat PE=1 SV=1 | 83.2 | 91.3 | 111.8 | 128.3 | 64.4 | 96.6 | 60.8 | 128.6 | 130.5 | 104.5 |
| 1020 | N-acyl-aromatic-L-amino acid amidohydrolase (carboxylate-forming) OS=Mus musculus OX=10090 GN=Acy3 PE=1 SV=1 | 171.6 | 125 | 143.6 | 101.2 | 66.5 | 126.7 | 61.1 | 74 | 80.4 | 49.9 |
| 1021 | Oxysterol-binding protein-related protein 1 OS=Mus musculus OX=10090 GN=Osbpl1a PE=1 SV=2 | 116.7 | 101.3 | 106.7 | 122.1 | 67.1 | 81.4 | 87.1 | 116.3 | 101.5 | 99.8 |
| 1022 | Fructose-bisphosphate aldolase B OS=Mus musculus OX=10090 GN=Aldob PE=1 SV=3 | 95.8 | 82.4 | 81.7 | 109.8 | 70.8 | 97.1 | 94.4 | 136.9 | 113.5 | 117.7 |
| 1023 | Argininosuccinate lyase OS=Mus musculus OX=10090 GN=Asl PE=1 SV=1 | 107.3 | 82.9 | 84.9 | 114.9 | 65.6 | 81.6 | 67.6 | 152.5 | 129.7 | 113 |
| 1024 | Deoxyribose-phosphate aldolase OS=Mus musculus OX=10090 GN=Dera PE=1 SV=1 | 130.7 | 108.5 | 110 | 113.6 | 86.5 | 117.9 | 71.3 | 90.2 | 101.8 | 69.5 |
| 1025 | Dolichyl-diphosphooligosaccharide--protein glycosyltransferase subunit 1 OS=Mus musculus OX=10090 GN=Rpn1 PE=1 SV=1 | 121.6 | 113.8 | 114.9 | 95.1 | 111.8 | 122.5 | 77.1 | 86.9 | 79.7 | 76.5 |
| 1026 | Dual-specificity mitogen-activated protein kinase kinase 2 OS=Mus musculus OX=10090 GN=Map2k2 PE=1 SV=1 | 116.2 | 99.1 | 111.8 | 111.2 | 71.3 | 107.4 | 73.2 | 110.7 | 99.3 | 99.8 |
| 1027 | DnaJ homolog subfamily C member 3 OS=Mus musculus OX=10090 GN=Dnajc3 PE=1 SV=1 | 93.6 | 89.7 | 87.1 | 115.8 | 54.9 | 169.8 | 107.8 | 63.1 | 113.8 | 104.4 |
| 1028 | Myosin-7 OS=Mus musculus OX=10090 GN=Myh7 PE=2 SV=1 | 47.9 | 62.8 | 49.8 | 42.8 | 206.2 | 49.8 | 397.8 | 50.1 | 33.3 | 59.5 |
| 1029 | Phospholipase ABHD3 OS=Mus musculus OX=10090 GN=Abhd3 PE=1 SV=1 | 106.3 | 92.8 | 103.8 | 134.6 | 77 | 89.2 | 93.2 | 106.4 | 102.9 | 93.7 |
| 1030 | Farnesyl pyrophosphate synthase OS=Mus musculus OX=10090 GN=Fdps PE=1 SV=1 | 180.6 | 144.1 | 137.1 | 74.5 | 81.6 | 104.2 | 68.5 | 79.6 | 56.3 | 73.3 |
| 1031 | Heterogeneous nuclear ribonucleoprotein L-like OS=Mus musculus OX=10090 GN=Hnrnpll PE=1 SV=3 | 118.8 | 91.3 | 88.6 | 74.9 | 111.7 | 107.5 | 119.8 | 107.4 | 87.2 | 92.7 |
| 1032 | Electron transfer flavoprotein-ubiquinone oxidoreductase, mitochondrial OS=Mus musculus OX=10090 GN=Etfdh PE=1 SV=1 | 91.1 | 103.9 | 110.3 | 113.5 | 85.5 | 100.3 | 100.2 | 102.8 | 101.2 | 91.1 |
| 1033 | Cytochrome c oxidase assembly factor 7 OS=Mus musculus OX=10090 GN=Coa7 PE=1 SV=1 | 98.6 | 81.5 | 81.9 | 94.7 | 111.1 | 95 | 107.8 | 123.8 | 90.4 | 115.4 |
| 1034 | GTP-binding protein Rheb OS=Mus musculus OX=10090 GN=Rheb PE=1 SV=1 | 119.3 | 102.3 | 103.5 | 102.4 | 76.9 | 112.3 | 83.1 | 107 | 99.9 | 93.4 |
| 1035 | ADP-ribose glycohydrolase MACROD1 OS=Mus musculus OX=10090 GN=Macrod1 PE=1 SV=2 | 122.1 | 99.8 | 92.4 | 98.7 | 111.7 | 100.9 | 134.8 | 91.3 | 83 | 65.4 |
| 1036 | Aspartate--tRNA ligase, cytoplasmic OS=Mus musculus OX=10090 GN=Dars1 PE=1 SV=2 | 121.7 | 107.8 | 106.8 | 97.6 | 81.7 | 113.8 | 96.2 | 99.4 | 85.5 | 89.5 |
| 1037 | C-1-tetrahydrofolate synthase, cytoplasmic OS=Mus musculus OX=10090 GN=Mthfd1 PE=1 SV=4 | 130.3 | 106.3 | 120.1 | 110 | 93.6 | 118.5 | 82.5 | 90.1 | 84.7 | 63.8 |
| 1038 | Ethanolamine-phosphate cytidylyltransferase OS=Mus musculus OX=10090 GN=Pcyt2 PE=1 SV=1 | 150.7 | 119 | 126.8 | 93.9 | 97 | 109 | 65.5 | 88.9 | 78.5 | 70.7 |
| 1039 | Mitochondrial amidoxime reducing component 2 OS=Mus musculus OX=10090 GN=Mtarc2 PE=1 SV=1 | 112.1 | 91.5 | 128.5 | 143.6 | 62.6 | 97 | 42 | 132 | 113.3 | 77.3 |
| 1040 | Leucine-rich repeat-containing protein 59 OS=Mus musculus OX=10090 GN=Lrrc59 PE=1 SV=1 | 131.7 | 114.8 | 128.8 | 87.3 | 88.3 | 116.2 | 73.4 | 90.6 | 76.9 | 92 |
| 1041 | Mannose-6-phosphate isomerase OS=Mus musculus OX=10090 GN=Mpi PE=1 SV=1 | 117.9 | 113.3 | 101.8 | 90.1 | 109.9 | 101.8 | 101.4 | 91.9 | 88 | 83.8 |
| 1042 | Gamma-butyrobetaine dioxygenase OS=Mus musculus OX=10090 GN=Bbox1 PE=1 SV=1 | 84.7 | 106.8 | 143.5 | 121.2 | 71.9 | 109.7 | 48.9 | 105.2 | 123.5 | 84.5 |
| 1043 | Ceramide synthase 2 OS=Mus musculus OX=10090 GN=Cers2 PE=1 SV=1 | 107.1 | 108.7 | 102.6 | 96.9 | 109.1 | 104.9 | 84.7 | 101.3 | 87.6 | 97 |
| 1044 | ATPase family AAA domain-containing protein 3 OS=Mus musculus OX=10090 GN=Atad3 PE=1 SV=1 | 97.7 | 88.2 | 101.4 | 112.8 | 84 | 87.5 | 105.4 | 112.4 | 118.6 | 91.9 |
| 1045 | Malonyl-CoA decarboxylase, mitochondrial OS=Mus musculus OX=10090 GN=Mlycd PE=1 SV=1 | 104.7 | 89.9 | 81.4 | 88.7 | 76.5 | 89.4 | 68.3 | 98 | 90.4 | 212.6 |
| 1046 | PC4 and SFRS1-interacting protein OS=Mus musculus OX=10090 GN=Psip1 PE=1 SV=1 | 98.7 | 99 | 92 | 79.2 | 107.6 | 106.1 | 113.3 | 105.7 | 80.1 | 118.2 |
| 1047 | Aminoacylase-1 OS=Mus musculus OX=10090 GN=Acy1 PE=1 SV=1 | 108.8 | 107.8 | 117.7 | 98.8 | 80.8 | 113.7 | 90.6 | 94.2 | 89.8 | 97.8 |
| 1048 | GTPase IMAP family member 4 OS=Mus musculus OX=10090 GN=Gimap4 PE=1 SV=2 | 156.3 | 107.3 | 128.1 | 89.9 | 86.9 | 132.2 | 53.8 | 89.5 | 80.9 | 75.1 |
| 1049 | Actin-related protein 3 OS=Mus musculus OX=10090 GN=Actr3 PE=1 SV=3 | | 115 | 83.3 | 99.9 | 55.8 | 133.7 | 159.9 | 87.8 | 134.6 | 130 |

| | | | | | | | | | | | |
|---|---|---|---|---|---|---|---|---|---|---|---|
| 1050 | ADP-ribosylation factor GTPase-activating protein 2 OS=Mus musculus OX=10090 GN=Arfgap2 PE=1 SV=1 | 115.7 | 100.4 | 108.9 | 82.8 | 82.8 | 94.9 | 121.9 | 79 | 107.4 | 106.1 |
| 1051 | Non-POU domain-containing octamer-binding protein OS=Mus musculus OX=10090 GN=Nono PE=1 SV=3 | 108.5 | 97.6 | 93.2 | 81 | 98.6 | 109.3 | 106.5 | 104.7 | 87.8 | 112.9 |
| 1052 | Plastin-3 OS=Mus musculus OX=10090 GN=Pls3 PE=1 SV=3 | 110.2 | 109.6 | 108.9 | 94.2 | 88.4 | 123.3 | 77.1 | 101.1 | 93 | 94.2 |
| 1053 | D16H22S680E protein OS=Mus musculus OX=10090 GN=Tango2 PE=1 SV=1 | 117.7 | 127.4 | 143.1 | 101.8 | 58.7 | 109.2 | 60.5 | 101 | 99.7 | 80.8 |
| 1054 | Alpha-aminoadipic semialdehyde synthase, mitochondrial OS=Mus musculus OX=10090 GN=Aass PE=1 SV=1 | 57.2 | 55.8 | 57.8 | 128.9 | 73.2 | 70.8 | 68.2 | 173.7 | 158.3 | 156.1 |
| 1055 | Beta-glucuronidase OS=Mus musculus OX=10090 GN=Gusb PE=1 SV=1 | 124.6 | 120.3 | 123.9 | 99.4 | 83.3 | 98.8 | 72 | 92.2 | 99.3 | 86.2 |
| 1056 | N-acylneuraminate cytidylyltransferase OS=Mus musculus OX=10090 GN=Cmas PE=1 SV=2 | 114.6 | 94.8 | 119.2 | 92.7 | 80.5 | 110.7 | 81.1 | 105 | 96.1 | 105.5 |
| 1057 | Lambda-crystallin homolog OS=Mus musculus OX=10090 GN=Cryl1 PE=1 SV=3 | 64.7 | 79.3 | 82.4 | 109.6 | 123.1 | 77.5 | 191.8 | 86.1 | 103.2 | 82.2 |
| 1058 | Nicotinamide phosphoribosyltransferase OS=Mus musculus OX=10090 GN=Nampt PE=1 SV=1 | 121 | 106.7 | 125 | 95.9 | 80.3 | 125 | 73.7 | 95.3 | 95 | 82.2 |
| 1059 | Peptidyl-prolyl cis-trans isomerase F, mitochondrial OS=Mus musculus OX=10090 GN=Ppif PE=1 SV=1 | 57.4 | 58.7 | 52.4 | 46.5 | 128.5 | 58.9 | 234.5 | 83.2 | 56.8 | 223 |
| 1060 | DnaJ homolog subfamily B member 11 OS=Mus musculus OX=10090 GN=Dnajb11 PE=1 SV=1 | 109 | 114.1 | 97.1 | 92.3 | 109 | 105.4 | 74.3 | 108.7 | 92.9 | 97.2 |
| 1061 | Dehydrogenase/reductase SDR family member 1 OS=Mus musculus OX=10090 GN=Dhrs1 PE=1 SV=1 | 126.1 | 113.2 | 108.9 | 80.3 | 129.9 | 98.2 | 102 | 88.1 | 72.2 | 81 |
| 1062 | 3-hydroxyisobutyrate dehydrogenase, mitochondrial OS=Mus musculus OX=10090 GN=Hibadh PE=1 SV=1 | 77.9 | 84 | 91 | 139.5 | 80.1 | 81.4 | 76.9 | 129.8 | 132.2 | 107.1 |
| 1063 | Glutathione S-transferase theta-3 OS=Mus musculus OX=10090 GN=Gstt3 PE=1 SV=1 | 73.4 | 65.6 | 69.9 | 169.9 | 31.5 | 97.6 | 65.9 | 91.9 | 193.5 | 140.9 |
| 1064 | Hsc70-interacting protein OS=Mus musculus OX=10090 GN=St13 PE=1 SV=1 | 121.1 | 109.2 | 117.8 | 91.4 | 119.8 | 122.8 | 79.8 | 88.9 | 69.3 | 79.8 |
| 1065 | Beta-1-syntrophin OS=Mus musculus OX=10090 GN=Sntb1 PE=1 SV=4 | 104.7 | 96.8 | 96.4 | 91.4 | 96.3 | 90.7 | 99.9 | 99 | 107.4 | 117.2 |
| 1066 | NADH dehydrogenase [ubiquinone] 1 alpha subcomplex subunit 10, mitochondrial OS=Mus musculus OX=10090 GN=Ndufa10 PE=1 SV=1 | 102.4 | 80.3 | 103.9 | 121.4 | 78.5 | 79.6 | 109.2 | 127.5 | 109.3 | 87.8 |
| 1067 | Electron transfer flavoprotein subunit alpha, mitochondrial OS=Mus musculus OX=10090 GN=Etfa PE=1 SV=2 | 109.3 | 98.7 | 91.2 | 108.7 | 74.4 | 111.8 | 80.9 | 107.7 | 112 | 105.4 |
| 1068 | N(G),N(G)-dimethylarginine dimethylaminohydrolase 2 OS=Mus musculus OX=10090 GN=Ddah2 PE=1 SV=1 | 116.9 | 103.2 | 102.7 | 116.7 | 91.3 | 110.1 | 73 | 102.4 | 106.9 | 76.7 |
| 1069 | CDK5 regulatory subunit-associated protein 3 OS=Mus musculus OX=10090 GN=Cdk5rap3 PE=1 SV=1 | 131.9 | 117.9 | 133.1 | 93.3 | 85 | 114.6 | 74.5 | 87.3 | 85.9 | 76.4 |
| 1070 | GrpE protein homolog 1, mitochondrial OS=Mus musculus OX=10090 GN=Grpel1 PE=1 SV=1 | 92.7 | 90.1 | 97.3 | 91.7 | 115.7 | 77.6 | 115.2 | 103.1 | 104.6 | 112.1 |
| 1071 | Parkinson disease protein 7 homolog OS=Mus musculus OX=10090 GN=Park7 PE=1 SV=1 | 108.9 | 127.8 | 103.4 | 103.7 | 98.2 | 100.2 | 63.6 | 103.2 | 93.9 | 97.2 |
| 1072 | DnaJ homolog subfamily A member 3, mitochondrial OS=Mus musculus OX=10090 GN=Dnaja3 PE=1 SV=1 | 91.4 | 96.7 | 97.2 | 94.3 | 107.7 | 96.1 | 95.5 | 111 | 94.3 | 115.8 |
| 1073 | Lysine--tRNA ligase OS=Mus musculus OX=10090 GN=Kars1 PE=1 SV=1 | 90.8 | 119.2 | 96.2 | 98.9 | 74.3 | 138.6 | 97.8 | 99.8 | 96.9 | 87.4 |
| 1074 | Peroxisomal coenzyme A diphosphatase NUDT7 OS=Mus musculus OX=10090 GN=Nudt7 PE=1 SV=2 | 149.4 | 145.3 | 116.4 | 69 | 103.1 | 126 | 77.5 | 82.7 | 64.5 | 65.9 |
| 1075 | Arylacetamide deacetylase OS=Mus musculus OX=10090 GN=Aadac PE=1 SV=3 | 151.1 | 118.8 | 133.9 | 100.4 | 86.1 | 150.4 | 66.2 | 63.7 | 78.8 | 50.5 |
| 1076 | Pre-mRNA-processing-splicing factor 8 OS=Mus musculus OX=10090 GN=Prpf8 PE=1 SV=2 | 112 | 102.1 | 103.6 | 93.4 | 87.1 | 108.3 | 95.8 | 102 | 94.7 | 101 |
| 1077 | NADH dehydrogenase [ubiquinone] 1 alpha subcomplex subunit 5 OS=Mus musculus OX=10090 GN=Ndufa5 PE=1 SV=3 | 90.5 | 77.8 | 84.6 | 121.2 | 96.1 | 77.7 | 106.1 | 116.9 | 119.5 | 109.6 |
| 1078 | Cytochrome c oxidase subunit 6C OS=Mus musculus OX=10090 GN=Cox6c PE=1 SV=3 | 73.8 | 87.2 | 77.4 | 107.7 | 122.4 | 79.7 | 96.2 | 126.9 | 108.5 | 120.2 |
| 1079 | ATP synthase subunit g, mitochondrial OS=Mus musculus OX=10090 GN=Atp5mg PE=1 SV=1 | 94 | 87.8 | 98.6 | 115.3 | 79.4 | 94 | 106.9 | 97.4 | 114.5 | 112.1 |
| 1080 | Cytosol aminopeptidase OS=Mus musculus OX=10090 GN=Lap3 PE=1 SV=3 | 114 | 104.3 | 117 | 105.2 | 90.7 | 112.6 | 84.2 | 95.7 | 95.4 | 80.9 |
| 1081 | 6-phosphogluconolactonase OS=Mus musculus OX=10090 GN=Pgls PE=1 SV=1 | 104.5 | 111.8 | 115.6 | 104.7 | 97.7 | 109.3 | 75.5 | 95.7 | 98.4 | 86.7 |
| 1082 | Cytochrome b-c1 complex subunit 8 OS=Mus musculus OX=10090 GN=Uqcrq PE=1 SV=3 | 95.5 | 87.5 | 85.8 | 117.6 | 102 | 84.2 | 82.4 | 108.3 | 127.8 | 108.7 |
| 1083 | Plakophilin 2 OS=Mus musculus OX=10090 GN=Pkp2 PE=1 SV=1 | 98.5 | 115.7 | 94.4 | 84.4 | 87.3 | 106.2 | 90.9 | 81.3 | 99.8 | 141.5 |
| 1084 | NADH dehydrogenase [ubiquinone] 1 alpha subcomplex subunit 2 OS=Mus musculus OX=10090 GN=Ndufa2 PE=1 SV=3 | 88.2 | 79.9 | 85.2 | 100.8 | 125.4 | 84.1 | 118.4 | 128.2 | 83.5 | 106.3 |

| | | | | | | | | | | | |
|---|---|---|---|---|---|---|---|---|---|---|---|
| 1085 | NADH dehydrogenase [ubiquinone] 1 alpha subcomplex subunit 3 OS=Mus musculus OX=10090 GN=Ndufa3 PE=1 SV=1 | 88.2 | 76.5 | 95.6 | 134.6 | 78.5 | 86 | 108.7 | 140.7 | 112.7 | 78.5 |
| 1086 | Cytochrome b-c1 complex subunit 7 OS=Mus musculus OX=10090 GN=Uqcrb PE=1 SV=1 | 65.9 | 70.3 | 64.9 | 118.7 | 96.8 | 106 | 147 | 80.1 | 118 | 132.3 |
| 1087 | GTP-binding protein SAR1b OS=Mus musculus OX=10090 GN=Sar1b PE=1 SV=1 | 125.3 | 112.3 | 123.9 | 89.7 | 90.6 | 101.1 | 104 | 87.4 | 96 | 69.8 |
| 1088 | Heat shock protein 75 kDa, mitochondrial OS=Mus musculus OX=10090 GN=Trap1 PE=1 SV=1 | 96.1 | 96 | 99.3 | 102.3 | 73.7 | 101.5 | 94.3 | 122.4 | 114.3 | 100.1 |
| 1089 | ATP synthase F(0) complex subunit B1, mitochondrial OS=Mus musculus OX=10090 GN=Atp5pb PE=1 SV=1 | 124.3 | 89.4 | 105.2 | 98.7 | 106.5 | 90.8 | 106.4 | 94.6 | 100.8 | 83.4 |
| 1090 | 40S ribosomal protein S21 OS=Mus musculus OX=10090 GN=Rps21 PE=1 SV=1 | 105.3 | 102.7 | 95.2 | 76.7 | 156.5 | 106.6 | 66.8 | 96.3 | 78.7 | 115.2 |
| 1091 | Methylthioribose-1-phosphate isomerase OS=Mus musculus OX=10090 GN=Mri1 PE=1 SV=1 | 113.3 | 110.8 | 112.7 | 100.2 | 84.1 | 112.4 | 86.9 | 97.7 | 88 | 93.8 |
| 1092 | Mitochondrial import inner membrane translocase subunit TIM14 OS=Mus musculus OX=10090 GN=Dnajc19 PE=1 SV=3 | 94.9 | 100.6 | 115.4 | 91 | 101.3 | 95.5 | 108.3 | 97.7 | 90.8 | 104.5 |
| 1093 | Peptidyl-prolyl cis-trans isomerase D OS=Mus musculus OX=10090 GN=Ppid PE=1 SV=3 | 123.7 | 115.6 | 118.1 | 92.4 | 98.3 | 114.8 | 67.5 | 104.1 | 79.7 | 85.8 |
| 1094 | Mitochondrial 2-oxoglutarate/malate carrier protein OS=Mus musculus OX=10090 GN=Slc25a11 PE=1 SV=3 | 109 | 99.5 | 97.8 | 110.8 | 102.6 | 94.8 | 122.7 | 92.6 | 93.4 | 76.8 |
| 1095 | Cytochrome b-c1 complex subunit Rieske, mitochondrial OS=Mus musculus OX=10090 GN=Uqcrfs1 PE=1 SV=1 | 93.9 | 87 | 87.6 | 104.8 | 111.4 | 90 | 116.4 | 101.6 | 105.7 | 101.6 |
| 1096 | ATP synthase F(0) complex subunit C1, mitochondrial OS=Mus musculus OX=10090 GN=Atp5mc1 PE=1 SV=1 | 95.8 | 125.6 | 85.5 | 100.9 | 100.3 | 95.3 | 121.5 | 100.1 | 91.2 | 83.9 |
| 1097 | Calcium-regulated heat stable protein 1 OS=Mus musculus OX=10090 GN=Carhsp1 PE=1 SV=1 | 80.3 | 96.1 | 77.4 | 67 | 165.2 | 88.3 | 206.3 | 76.1 | 54.4 | 88.9 |
| 1098 | Methylsterol monooxygenase 1 OS=Mus musculus OX=10090 GN=Msmo1 PE=1 SV=1 | 196.8 | 116.4 | 106.6 | 117 | 63 | 64.8 | 21.6 | 157.7 | 102.2 | 53.8 |
| 1099 | 5-hydroxyisourate hydrolase OS=Mus musculus OX=10090 GN=Urah PE=1 SV=1 | 138.1 | 128.2 | 124.8 | 100.2 | 59.8 | 104.6 | 52.3 | 110.6 | 93.8 | 87.5 |
| 1100 | Actin-related protein 2/3 complex subunit 2 OS=Mus musculus OX=10090 GN=Arpc2 PE=1 SV=3 | 115.1 | 92.4 | 101.3 | 86.3 | 85.9 | 113.2 | 61.1 | 88.3 | 83.4 | 172.9 |
| 1101 | Bifunctional purine biosynthesis protein ATIC OS=Mus musculus OX=10090 GN=Atic PE=1 SV=2 | 137.4 | 113.4 | 128.2 | 102.5 | 95.8 | 120.8 | 72.8 | 90.8 | 73.9 | 64.3 |
| 1102 | Beta-catenin-like protein 1 OS=Mus musculus OX=10090 GN=Ctnnbl1 PE=1 SV=1 | 115.1 | 104.9 | 94.3 | 79.4 | 85 | 92.3 | 114.3 | 82.2 | 95.7 | 136.9 |
| 1103 | Motile sperm domain-containing protein 2 OS=Mus musculus OX=10090 GN=Mospd2 PE=1 SV=2 | | 181.8 | | 169.1 | | 196.9 | 80.4 | 142.2 | 130.1 | 99.5 |
| 1104 | N(G),N(G)-dimethylarginine dimethylaminohydrolase 1 OS=Mus musculus OX=10090 GN=Ddah1 PE=1 SV=3 | 84 | 75.5 | 74 | 124 | 70.7 | 126 | 98.1 | 76.1 | 157.2 | 114.4 |
| 1105 | IST1 homolog OS=Mus musculus OX=10090 GN=Ist1 PE=1 SV=1 | 116.3 | 110 | 109.3 | 95.5 | 86.3 | 107.6 | 82.6 | 100.7 | 94.7 | 97.1 |
| 1106 | Centromere protein V OS=Mus musculus OX=10090 GN=Cenpv PE=1 SV=2 | 118.7 | 103.4 | 106.5 | 95.6 | 83.9 | 105 | 73.2 | 106.5 | 92.4 | 114.7 |
| 1107 | Mitochondrial-processing peptidase subunit beta OS=Mus musculus OX=10090 GN=Pmpcb PE=1 SV=1 | 128.5 | 99.6 | 98.6 | 101.5 | 91.1 | 102.4 | 87.4 | 113 | 87.8 | 90.1 |
| 1108 | 60S ribosomal protein L11 OS=Mus musculus OX=10090 GN=Rpl11 PE=1 SV=4 | 118.9 | 114 | 121.4 | 85.2 | 103.2 | 126.3 | 62.8 | 92.1 | 87.2 | 88.9 |
| 1109 | NADH dehydrogenase [ubiquinone] iron-sulfur protein 4, mitochondrial OS=Mus musculus OX=10090 GN=Ndufs4 PE=1 SV=3 | 77.3 | 70.8 | 71.6 | 94.5 | 144.9 | 76.1 | 107.9 | 127.5 | 97.8 | 131.6 |
| 1110 | Peroxiredoxin-like 2A OS=Mus musculus OX=10090 GN=Prxl2a PE=1 SV=2 | 141.2 | 94.9 | 96.2 | 98.8 | 79.8 | 118.4 | 77.5 | 98 | 116.7 | 78.5 |
| 1111 | Haloacid dehalogenase-like hydrolase domain-containing protein 3 OS=Mus musculus OX=10090 GN=Hdhd3 PE=1 SV=1 | 125.1 | 111.3 | 116.4 | 98.4 | 92.5 | 115.9 | 73.8 | 87.8 | 87.3 | 91.5 |
| 1112 | Cytochrome b-c1 complex subunit 1, mitochondrial OS=Mus musculus OX=10090 GN=Uqcrc1 PE=1 SV=2 | 87.6 | 89.1 | 89.2 | 98.4 | 103.5 | 91.7 | 101.2 | 112.8 | 111.8 | 114.7 |
| 1113 | Obg-like ATPase 1 OS=Mus musculus OX=10090 GN=Ola1 PE=1 SV=1 | 81.2 | 82.7 | 88.3 | 119.6 | 92.4 | 132.7 | 129.2 | 76.9 | 104.8 | 92.3 |
| 1114 | Aldehyde dehydrogenase X, mitochondrial OS=Mus musculus OX=10090 GN=Aldh1b1 PE=1 SV=1 | 82.4 | 75.8 | 100.1 | 130.8 | 104.5 | 113.8 | 76.7 | 110.3 | 123.3 | 82.5 |
| 1115 | Citrate synthase, mitochondrial OS=Mus musculus OX=10090 GN=Cs PE=1 SV=1 | 110 | 106.7 | 107 | 89.2 | 112.7 | 98.9 | 125.5 | 86.8 | 83.8 | 79.6 |
| 1116 | Mitochondrial import receptor subunit TOM70 OS=Mus musculus OX=10090 GN=Tomm70 PE=1 SV=2 | 123.2 | 100.7 | 116.3 | 97.1 | 86.5 | 107.1 | 80.4 | 109.8 | 89.8 | 89 |
| 1117 | ER membrane protein complex subunit 4 OS=Mus musculus OX=10090 GN=Emc4 PE=1 SV=1 | 15.2 | 137.4 | 16.9 | 136.6 | 14.4 | 167.2 | 111.5 | 126.3 | 126.7 | 147.7 |
| 1118 | General transcription factor IIE subunit 1 OS=Mus musculus OX=10090 GN=Gtf2e1 PE=1 SV=1 | | 109.5 | | 131.9 | | 162.7 | 137 | 174.3 | 150.1 | 134.5 |

| | | | | | | | | | | | |
|---|---|---|---|---|---|---|---|---|---|---|---|
| 1119 | Heterogeneous nuclear ribonucleoprotein M OS=Mus musculus OX=10090 GN=Hnrnpm PE=1 SV=3 | 114.6 | 90.5 | 94.8 | 80.6 | 120 | 112.9 | 110.3 | 96.9 | 78.4 | 100.9 |
| 1120 | Arginine--tRNA ligase, cytoplasmic OS=Mus musculus OX=10090 GN=Rars1 PE=1 SV=2 | 115.1 | 114.3 | 118 | 98.2 | 84.8 | 122 | 80.3 | 95.7 | 87.7 | 83.9 |
| 1121 | Cytochrome c1, heme protein, mitochondrial OS=Mus musculus OX=10090 GN=Cyc1 PE=1 SV=1 | 101.8 | 90.3 | 93.3 | 112.5 | 86 | 93.1 | 121.6 | 108 | 105.4 | 88 |
| 1122 | Dynein light chain 2, cytoplasmic OS=Mus musculus OX=10090 GN=Dynll2 PE=1 SV=1 | 138.3 | 131.7 | 121.2 | 89.4 | 84.8 | 108.4 | 75.3 | 84.2 | 83 | 83.7 |
| 1123 | Adenosine 5'-monophosphoramidase HINT2 OS=Mus musculus OX=10090 GN=Hint2 PE=1 SV=1 | 100.3 | 87.8 | 95.2 | 105 | 80.2 | 92.1 | 66.6 | 122.4 | 111 | 139.4 |
| 1124 | Glutamine amidotransferase-like class 1 domain-containing protein 3, mitochondrial OS=Mus musculus OX=10090 GN=Gatd3 PE=1 SV=1 | 68.3 | 78.2 | 73.1 | 87 | 94.5 | 80.1 | 135.2 | 89.6 | 137.3 | 156.8 |
| 1125 | Cytosolic non-specific dipeptidase OS=Mus musculus OX=10090 GN=Cndp2 PE=1 SV=1 | 113.8 | 106.6 | 133.3 | 91.6 | 104.5 | 125.3 | 88 | 87 | 71.5 | 78.4 |
| 1126 | Coiled-coil-helix-coiled-coil-helix domain-containing protein 2 OS=Mus musculus OX=10090 GN=Chchd2 PE=2 SV=1 | 97.8 | 96.8 | 99.5 | 91.1 | 142.5 | 98.8 | 88.2 | 106.2 | 74.5 | 104.4 |
| 1127 | Eukaryotic translation elongation factor 1 epsilon-1 OS=Mus musculus OX=10090 GN=Eef1e1 PE=1 SV=1 | 119.5 | 106.4 | 110.1 | 90.8 | 103.2 | 128.1 | 88.8 | 80.9 | 91.3 | 81.1 |
| 1128 | Endoplasmic reticulum resident protein 44 OS=Mus musculus OX=10090 GN=Erp44 PE=1 SV=1 | 134.3 | 114.3 | 129.2 | 90 | 85.2 | 130.3 | 59.5 | 93.6 | 86.6 | 76.9 |
| 1129 | 60S ribosomal protein L34 OS=Mus musculus OX=10090 GN=Rpl34 PE=1 SV=2 | 136 | 107.6 | 134.5 | 93.1 | 72.7 | 112.5 | 70.5 | 102.3 | 95.1 | 75.6 |
| 1130 | Dihydrolipoyllysine-residue succinyltransferase component of 2-oxoglutarate dehydrogenase complex, mitochondrial OS=Mus musculus OX=10090 GN=Dlst PE=1 SV=1 | 85.9 | 73.8 | 105.2 | 115.2 | 87.9 | 78.1 | 80.2 | 131 | 135.7 | 107 |
| 1131 | Acetoacetyl-CoA synthetase OS=Mus musculus OX=10090 GN=Aacs PE=1 SV=1 | 133.7 | 121.3 | 110.9 | 70.2 | 89.8 | 116.5 | 86.6 | 87.5 | 72.5 | 111.1 |
| 1132 | Dehydrogenase/reductase SDR family member 4 OS=Mus musculus OX=10090 GN=Dhrs4 PE=1 SV=1 | 106.7 | 104.9 | 103.8 | 102.2 | 82.2 | 98.9 | 79.2 | 95.8 | 118.4 | 108 |
| 1133 | Low molecular weight phosphotyrosine protein phosphatase OS=Mus musculus OX=10090 GN=Acp1 PE=1 SV=3 | 134.2 | 116.4 | 112.6 | 98.9 | 86.9 | 113.2 | 72.2 | 92.8 | 84.7 | 88.1 |
| 1134 | Epoxide hydrolase 1 OS=Mus musculus OX=10090 GN=Ephx1 PE=1 SV=2 | 87.4 | 75.1 | 134.3 | 151.6 | 58 | 77.3 | 44.9 | 127.7 | 161.6 | 82 |
| 1135 | 3-oxoacyl-[acyl-carrier-protein] synthase, mitochondrial OS=Mus musculus OX=10090 GN=Oxsm PE=1 SV=1 | 118.8 | 100.5 | 118.5 | 123 | 47.3 | 82.5 | 58.7 | 116.9 | 113.4 | 120.5 |
| 1136 | Polyadenylate-binding protein OS=Mus musculus OX=10090 GN=Pabpc6 PE=1 SV=1 | 143.9 | 117.7 | 152.2 | 86.2 | 85.8 | 111.6 | 59 | 85.6 | 79.7 | 78.3 |
| 1137 | Cullin-2 OS=Mus musculus OX=10090 GN=Cul2 PE=1 SV=2 | 125.9 | 111.5 | 115.2 | 88.1 | 84.4 | 114.8 | 88.8 | 96.6 | 86.1 | 88.8 |
| 1138 | NADH dehydrogenase [ubiquinone] flavoprotein 2, mitochondrial OS=Mus musculus OX=10090 GN=Ndufv2 PE=1 SV=2 | 50.3 | 89.2 | 53.8 | 83 | 97.9 | 79.3 | 213.2 | 114.8 | 84.9 | 133.6 |
| 1139 | Calmodulin-like protein 3 OS=Mus musculus OX=10090 GN=Calml3 PE=2 SV=1 | 144.4 | 115.7 | 87 | 89.5 | 132.5 | 86.6 | 62.8 | 116.2 | 43.7 | 121.5 |
| 1140 | Isocitrate dehydrogenase [NAD] subunit alpha, mitochondrial OS=Mus musculus OX=10090 GN=Idh3a PE=1 SV=1 | 49.5 | 78 | 70.2 | 90.3 | 149.8 | 78.7 | 208.1 | 90.6 | 83.7 | 101 |
| 1141 | Mitochondrial peptide methionine sulfoxide reductase OS=Mus musculus OX=10090 GN=Msra PE=1 SV=1 | 108.9 | 133.2 | 91.5 | 114.2 | 85.8 | 116.8 | 73.5 | 98.6 | 97.5 | 80.1 |
| 1142 | HAUS augmin-like complex subunit 5 OS=Mus musculus OX=10090 GN=Haus5 PE=1 SV=1 | 114 | 82.3 | 118.9 | 128.4 | 75.9 | 105.6 | 80.4 | 120.3 | 99.1 | 75.1 |
| 1143 | Enoyl-CoA hydratase domain-containing protein 3, mitochondrial OS=Mus musculus OX=10090 GN=Echdc3 PE=1 SV=1 | 115.6 | 102.5 | 115.9 | 103 | 91.4 | 99.8 | 71.2 | 105.3 | 94.8 | 100.5 |
| 1144 | Cytochrome P450 2C55 OS=Mus musculus OX=10090 GN=Cyp2c55 PE=1 SV=1 | 119.9 | 101.5 | 98.7 | 152.1 | 53.8 | 65.4 | 28.4 | 113.2 | 203.2 | 63.8 |
| 1145 | Inorganic pyrophosphatase OS=Mus musculus OX=10090 GN=Ppa1 PE=1 SV=1 | 103.6 | 106.1 | 102.3 | 106.5 | 66.3 | 129 | 71.4 | 116.7 | 92 | 106.1 |
| 1146 | 60S ribosomal protein L37 OS=Mus musculus OX=10090 GN=Rpl37 PE=1 SV=3 | 106.8 | 94.8 | 97.7 | 80.1 | 124.7 | 127.2 | 64 | 125.2 | 75.6 | 103.9 |
| 1147 | Peroxisomal sarcosine oxidase OS=Mus musculus OX=10090 GN=Pipox PE=1 SV=1 | 129.6 | 105 | 102.9 | 101.9 | 89.9 | 122.9 | 76.7 | 100.9 | 97.1 | 73.1 |
| 1148 | Mitochondrial import inner membrane translocase subunit TIM50 OS=Mus musculus OX=10090 GN=Timm50 PE=1 SV=1 | 98.6 | 93 | 106 | 94.7 | 98.5 | 107.5 | 82.2 | 108.1 | 95.6 | 115.6 |
| 1149 | Cytochrome c oxidase polypeptide Vb OS=Mus musculus OX=10090 GN=Cox5b-ps PE=1 SV=1 | 84.5 | 97.2 | 88.5 | 99.1 | 87.9 | 101.4 | 98.6 | 103.5 | 107.7 | 131.6 |
| 1150 | Inosine triphosphate pyrophosphatase OS=Mus musculus OX=10090 GN=Itpa PE=1 SV=2 | 122.2 | 114.2 | 119.8 | 94.6 | 95.1 | 101.5 | 96.6 | 78.6 | 76.8 | 100.6 |
| 1151 | Interferon-induced 35 kDa protein homolog OS=Mus musculus OX=10090 GN=Ifi35 PE=1 SV=3 | 105.5 | 100.9 | 128.8 | 95.2 | 95.8 | 114.7 | 87.5 | 101 | 85.1 | 85.5 |

| | | | | | | | | | | | |
|---|---|---|---|---|---|---|---|---|---|---|---|
| 1152 | 60S ribosomal protein L4 OS=Mus musculus OX=10090 GN=Rpl4 PE=1 SV=3 | 145.8 | 75 | 109.2 | 81.6 | 79.2 | 114.8 | 77.5 | 119.3 | 84.6 | 112.8 |
| 1153 | Elongation factor 1-gamma OS=Mus musculus OX=10090 GN=Eef1g PE=1 SV=3 | 116.3 | 100.8 | 109.7 | 82.8 | 111.6 | 122.9 | 81.2 | 100.3 | 90.4 | 83.8 |
| 1154 | Minor histocompatibility antigen H13 OS=Mus musculus OX=10090 GN=Hm13 PE=1 SV=1 | 141.8 | 118.2 | 100.3 | 96 | 69.5 | 99 | 60.7 | 121.8 | 95.3 | 97.5 |
| 1155 | Ethylmalonyl-CoA decarboxylase OS=Mus musculus OX=10090 GN=Echdc1 PE=1 SV=2 | 127.2 | 107.8 | 119.6 | 100.7 | 78 | 109.3 | 48.4 | 105.4 | 107.4 | 96.1 |
| 1156 | m7GpppX diphosphatase OS=Mus musculus OX=10090 GN=Dcps PE=1 SV=1 | 136.6 | 128.8 | 121.1 | 87.7 | 73.8 | 103.4 | 68.1 | 88.5 | 88.2 | 103.9 |
| 1157 | Calponin-3 OS=Mus musculus OX=10090 GN=Cnn3 PE=1 SV=1 | 55.8 | 110.1 | 90.5 | 98.5 | 104.2 | 105.6 | 77.4 | 102.6 | 83.7 | 171.4 |
| 1158 | Nucleophosmin OS=Mus musculus OX=10090 GN=Npm1 PE=1 SV=1 | 95.5 | 88.6 | 90.2 | 80.8 | 148 | 94.1 | 70.8 | 106.7 | 82.3 | 142.9 |
| 1159 | ATP synthase subunit O, mitochondrial OS=Mus musculus OX=10090 GN=Atp5po PE=1 SV=1 | 92.6 | 84.7 | 95.7 | 110.4 | 109.1 | 85.4 | 110.8 | 98.3 | 114.8 | 98.1 |
| 1160 | Mitochondrial glutamate carrier 2 OS=Mus musculus OX=10090 GN=Slc25a18 PE=1 SV=4 | 118.1 | 76.2 | 100.8 | 147.7 | 68.3 | 94 | 63.8 | 114.7 | 115.5 | 100.9 |
| 1161 | Cytochrome b-c1 complex subunit 2, mitochondrial OS=Mus musculus OX=10090 GN=Uqcrc2 PE=1 SV=1 | 116.6 | 88.5 | 94.1 | 102.3 | 88.3 | 103.5 | 115.7 | 103.5 | 93.8 | 93.7 |
| 1162 | Probable imidazolonepropionase OS=Mus musculus OX=10090 GN=Amdhd1 PE=1 SV=1 | 131.5 | 98.8 | 106.7 | 112 | 83.6 | 110.6 | 63.4 | 117.8 | 102.2 | 73.4 |
| 1163 | Cap-specific mRNA (nucleoside-2'-O-)-methyltransferase 1 OS=Mus musculus OX=10090 GN=Cmtr1 PE=1 SV=1 | | 77.4 | | 208.8 | | 129 | 134 | 146 | 129.1 | 175.6 |
| 1164 | Double-stranded RNA-binding protein Staufen homolog 1 OS=Mus musculus OX=10090 GN=Stau1 PE=1 SV=1 | 111.5 | 106.6 | 111.8 | 96.5 | 99.9 | 115.8 | 81.5 | 99.7 | 83.3 | 93.4 |
| 1165 | Alpha-aminoadipic semialdehyde dehydrogenase OS=Mus musculus OX=10090 GN=Aldh7a1 PE=1 SV=4 | 96.7 | 88.9 | 95.3 | 131.4 | 97.4 | 93.3 | 59.3 | 117.3 | 112.4 | 108 |
| 1166 | AP-2 complex subunit beta OS=Mus musculus OX=10090 GN=Ap2b1 PE=1 SV=1 | 56.7 | 110.1 | 116.8 | 100.4 | 91.5 | 119.7 | 87.9 | 103.8 | 99.9 | 113.2 |
| 1167 | Phosphoglycerate mutase 1 OS=Mus musculus OX=10090 GN=Pgam1 PE=1 SV=3 | 44.5 | 103.7 | 45.3 | 103.5 | 88.5 | 104.1 | 195.5 | 97.7 | 105.7 | 111.7 |
| 1168 | Acetyl-coenzyme A thioesterase OS=Mus musculus OX=10090 GN=Acot12 PE=1 SV=1 | 140.1 | 112.4 | 115.5 | 95.4 | 64.1 | 111.1 | 53.5 | 111.6 | 106.4 | 89.8 |
| 1169 | Bifunctional coenzyme A synthase OS=Mus musculus OX=10090 GN=Coasy PE=1 SV=2 | 116.5 | 103.2 | 136.1 | 114.7 | 59.3 | 100 | 65.1 | 129.6 | 113.2 | 62.3 |
| 1170 | Peroxisomal bifunctional enzyme OS=Mus musculus OX=10090 GN=Ehhadh PE=1 SV=4 | 119.8 | 99.1 | 120.8 | 113.9 | 69.8 | 97.8 | 60.3 | 116.5 | 120 | 82.1 |
| 1171 | Dimethylglycine dehydrogenase, mitochondrial OS=Mus musculus OX=10090 GN=Dmgdh PE=1 SV=1 | 91.5 | 94.1 | 110.8 | 156.1 | 53.7 | 71.3 | 55.4 | 109.8 | 143 | 114.2 |
| 1172 | Peroxisomal carnitine O-octanoyltransferase OS=Mus musculus OX=10090 GN=Crot PE=1 SV=1 | 95 | 128.2 | 131.8 | 89.5 | 104.5 | 112 | 88 | 109.8 | 75.5 | 65.7 |
| 1173 | Mitochondrial-processing peptidase subunit alpha OS=Mus musculus OX=10090 GN=Pmpca PE=1 SV=1 | 115.1 | 105.4 | 113.2 | 91.2 | 101.5 | 100 | 85.7 | 99.9 | 84.4 | 103.7 |
| 1174 | F-box only protein 3 OS=Mus musculus OX=10090 GN=Fbxo3 PE=1 SV=1 | 119 | 115.5 | 94.4 | 78.7 | 86.2 | 87.6 | 153.1 | 110.2 | 76.5 | 78.8 |
| 1175 | NADH dehydrogenase [ubiquinone] 1 alpha subcomplex subunit 9, mitochondrial OS=Mus musculus OX=10090 GN=Ndufa9 PE=1 SV=2 | 105 | 90.1 | 104.2 | 97.1 | 97.4 | 101.7 | 73.5 | 126.6 | 99.5 | 105 |
| 1176 | NADH dehydrogenase [ubiquinone] iron-sulfur protein 7, mitochondrial OS=Mus musculus OX=10090 GN=Ndufs7 PE=1 SV=1 | 94.8 | 84.3 | 91.3 | 105.3 | 92.5 | 90.2 | 110.4 | 126.8 | 105.4 | 99 |
| 1177 | 6-phosphogluconate dehydrogenase, decarboxylating OS=Mus musculus OX=10090 GN=Pgd PE=1 SV=3 | 122.3 | 98 | 123.2 | 82.1 | 132.5 | 133.9 | 97.2 | 64.3 | 72.3 | 74.3 |
| 1178 | Eukaryotic translation initiation factor 3 subunit F OS=Mus musculus OX=10090 GN=Eif3f PE=1 SV=2 | 113.4 | 101 | 109.1 | 96.9 | 47.3 | 112.2 | 82.5 | 128.2 | 90.8 | 118.7 |
| 1179 | NADH dehydrogenase [ubiquinone] 1 alpha subcomplex subunit 8 OS=Mus musculus OX=10090 GN=Ndufa8 PE=1 SV=3 | 101 | 90.8 | 92 | 107.3 | 115.1 | 83.9 | 90.7 | 123.9 | 95.9 | 99.5 |
| 1180 | Persulfide dioxygenase ETHE1, mitochondrial OS=Mus musculus OX=10090 GN=Ethe1 PE=1 SV=2 | | 81.3 | | 163.2 | | 94.7 | 111.9 | 156.5 | 195.9 | 196.5 |
| 1181 | Glutathione S-transferase kappa 1 OS=Mus musculus OX=10090 GN=Gstk1 PE=1 SV=3 | 99 | 97.5 | 81.9 | 127.1 | 64.9 | 95.6 | 72.2 | 121.5 | 135.1 | 105.1 |
| 1182 | NAD-capped RNA hydrolase NUDT12 OS=Mus musculus OX=10090 GN=Nudt12 PE=1 SV=1 | 94.7 | 99.4 | 99.3 | 102.3 | 89.1 | 86.9 | 88.5 | 122.5 | 101.2 | 116 |
| 1183 | NADH-cytochrome b5 reductase 3 OS=Mus musculus OX=10090 GN=Cyb5r3 PE=1 SV=3 | 145.6 | 133.1 | 140.2 | 82.6 | 89.6 | 111.2 | 88.1 | 74 | 85.6 | 50 |
| 1184 | NADH dehydrogenase [ubiquinone] 1 beta subcomplex subunit 10 OS=Mus musculus OX=10090 GN=Ndufb10 PE=1 SV=3 | 87.1 | 92 | 90.9 | 79.8 | 157.8 | 98.4 | 168.7 | 64.8 | 88 | 72.6 |
| 1185 | 1,5-anhydro-D-fructose reductase OS=Mus musculus OX=10090 GN=Akr1e2 PE=1 SV=1 | 114.2 | 119.4 | 124.8 | 100.4 | 90.4 | 130.6 | 79.3 | 78.5 | 91.1 | 71.4 |

| | | | | | | | | | | | |
|---|---|---|---|---|---|---|---|---|---|---|---|
| 1186 | ATP synthase subunit d, mitochondrial OS=Mus musculus OX=10090 GN=Atp5pd PE=1 SV=3 | 91.4 | 88.7 | 97.9 | 102 | 119.4 | 84.2 | 134.8 | 102.8 | 100.1 | 78.8 |
| 1187 | Iodotyrosine deiodinase 1 OS=Mus musculus OX=10090 GN=Iyd PE=1 SV=1 | 86.6 | 99.5 | 112.8 | 73.7 | 188.5 | 112 | 53.1 | 120.5 | 74.1 | 79.3 |
| 1188 | Glycine N-acyltransferase-like protein Keg1 OS=Mus musculus OX=10090 GN=Keg1 PE=1 SV=1 | 127 | 119.2 | 115.4 | 90.5 | 89.7 | 105.7 | 91.9 | 80 | 90 | 90.6 |
| 1189 | Exportin-7 OS=Mus musculus OX=10090 GN=Xpo7 PE=1 SV=3 | 102 | 100.4 | 92 | 80.8 | 131.9 | 88.9 | 148.7 | 90.5 | 75 | 89.8 |
| 1190 | Estradiol 17-beta-dehydrogenase 11 OS=Mus musculus OX=10090 GN=Hsd17b11 PE=1 SV=1 | 126.8 | 117.6 | 114.7 | 98.4 | 73.7 | 109.7 | 72.5 | 105.2 | 107.6 | 73.7 |
| 1191 | Heparan N-sulfatase OS=Mus musculus OX=10090 GN=Sgsh PE=1 SV=1 | 95.5 | 72.8 | 75.6 | 111.1 | 96.9 | 100.1 | 61.4 | 149.4 | 80.2 | 157.2 |
| 1192 | Methylmalonate-semialdehyde dehydrogenase [acylating], mitochondrial OS=Mus musculus OX=10090 GN=Aldh6a1 PE=1 SV=1 | 106.1 | 93.5 | 105.6 | 107.7 | 80.9 | 100.7 | 74.2 | 122.2 | 101.5 | 107.7 |
| 1193 | Dihydropyrimidinase OS=Mus musculus OX=10090 GN=Dpys PE=1 SV=2 | 171.2 | 133.9 | 124.9 | 101.1 | 75 | 130.6 | 60.8 | 80.6 | 75.4 | 46.6 |
| 1194 | Endoplasmic reticulum aminopeptidase 1 OS=Mus musculus OX=10090 GN=Erap1 PE=1 SV=2 | 149 | 106.1 | 117.4 | 96.4 | 98.9 | 129.9 | 71.2 | 98.1 | 86.9 | 46.1 |
| 1195 | EH domain-containing protein 4 OS=Mus musculus OX=10090 GN=Ehd4 PE=1 SV=1 | 95.2 | 97.6 | 91.9 | 86.3 | 116.3 | 101.3 | 97.7 | 107.1 | 86.7 | 119.9 |
| 1196 | LIM domain and actin-binding protein 1 OS=Mus musculus OX=10090 GN=Lima1 PE=1 SV=3 | 102.4 | 108.3 | 113 | 82.6 | 107.1 | 108.6 | 99.3 | 86.7 | 89.6 | 102.5 |
| 1197 | H/ACA ribonucleoprotein complex subunit DKC1 OS=Mus musculus OX=10090 GN=Dkc1 PE=1 SV=4 | 124.7 | 106.3 | 113.1 | 93.6 | 84.9 | 110 | 68 | 97.1 | 104.4 | 97.9 |
| 1198 | Glycogen phosphorylase, liver form OS=Mus musculus OX=10090 GN=Pygl PE=1 SV=4 | 152.8 | 110.5 | 135 | 96.2 | 85.3 | 100.6 | 75.7 | 89 | 92.7 | 62.3 |
| 1199 | Dipeptidyl peptidase 2 OS=Mus musculus OX=10090 GN=Dpp7 PE=1 SV=2 | 137.5 | 133.4 | 120 | 59.9 | 116.2 | 138.5 | 62.7 | 80 | 75.1 | 76.7 |
| 1200 | Isovaleryl-CoA dehydrogenase, mitochondrial OS=Mus musculus OX=10090 GN=Ivd PE=1 SV=1 | 94.2 | 92.3 | 97.5 | 109.9 | 90.2 | 95.6 | 77 | 120.8 | 114.2 | 108.3 |
| 1201 | Insulin-degrading enzyme OS=Mus musculus OX=10090 GN=Ide PE=1 SV=1 | 148.6 | 120.9 | 113.4 | 99.7 | 82 | 96.7 | 56.2 | 101.1 | 97.4 | 84 |
| 1202 | ATP-dependent Clp protease ATP-binding subunit clpX-like, mitochondrial OS=Mus musculus OX=10090 GN=Clpx PE=1 SV=2 | 90.6 | 92 | 81 | 72.7 | 132.9 | 96.2 | 147.9 | 109.7 | 74 | 103 |
| 1203 | Cytoplasmic dynein 1 heavy chain 1 OS=Mus musculus OX=10090 GN=Dync1h1 PE=1 SV=2 | 83.8 | 90.3 | 92 | 84.2 | 126.2 | 102.9 | 165.6 | 94.2 | 79.4 | 81.5 |
| 1204 | Alpha-actinin-2 OS=Mus musculus OX=10090 GN=Actn2 PE=1 SV=2 | 20.2 | 40.2 | 21.9 | 23.5 | 267.5 | 23.6 | 527.1 | 22.4 | 11.5 | 42.2 |
| 1205 | CPN10-like protein OS=Mus musculus OX=10090 GN=Hspe1-rs1 PE=3 SV=1 | 79.6 | 90.5 | 77.4 | 80.6 | 151.4 | 94.2 | 65.7 | 116.7 | 88.5 | 155.5 |
| 1206 | Coatomer subunit beta OS=Mus musculus OX=10090 GN=Copb1 PE=1 SV=1 | 93.9 | 70.5 | 85.5 | 58.6 | 51.6 | 68.6 | 131.9 | 77.6 | 167.6 | 194.1 |
| 1207 | Aldo-keto reductase family 1 member A1 OS=Mus musculus OX=10090 GN=Akr1a1 PE=1 SV=3 | 110.8 | 103 | 105.5 | 99.3 | 95 | 101 | 90.7 | 94 | 105.8 | 94.9 |
| 1208 | Nucleolar RNA helicase 2 OS=Mus musculus OX=10090 GN=Ddx21 PE=1 SV=3 | 98.4 | 81.2 | 107.9 | 85.5 | 103.3 | 90.6 | 119.9 | 108.7 | 96.7 | 107.7 |
| 1209 | Na(+)/H(+) exchange regulatory cofactor NHE-RF3 OS=Mus musculus OX=10090 GN=Pdzk1 PE=1 SV=1 | 118 | 118.1 | 113.5 | 76.1 | 93.9 | 105.7 | 81.4 | 80.1 | 92.4 | 120.8 |
| 1210 | Probable N-acetyltransferase CML1 OS=Mus musculus OX=10090 GN=Cml1 PE=1 SV=1 | 139 | 125.8 | 100.4 | 112.4 | 84.3 | 162.9 | 46.1 | 89.2 | 90.5 | 49.4 |
| 1211 | 60S ribosomal protein L38 OS=Mus musculus OX=10090 GN=Rpl38 PE=1 SV=3 | 106.9 | 106.1 | 98.7 | 83.2 | 130.1 | 102 | 127.6 | 97 | 75.3 | 73.2 |
| 1212 | Myozenin-1 OS=Mus musculus OX=10090 GN=Myoz1 PE=1 SV=1 | 23.6 | 44.1 | 27.5 | 30 | 300.7 | 27.9 | 450 | 28.3 | 18.5 | 49.4 |
| 1213 | MYG1 exonuclease OS=Mus musculus OX=10090 GN=Myg1 PE=1 SV=1 | 87.8 | 94.2 | 85.5 | 75.5 | 128.9 | 90.8 | 173.9 | 95.1 | 73.7 | 94.6 |
| 1214 | High mobility group nucleosome-binding domain-containing protein 5 OS=Mus musculus OX=10090 GN=Hmgn5 PE=1 SV=2 | 73.5 | 109.6 | 91.5 | 78.3 | 114.7 | 112.8 | 65.5 | 116.8 | 77.5 | 159.9 |
| 1215 | 4-trimethylaminobutyraldehyde dehydrogenase OS=Mus musculus OX=10090 GN=Aldh9a1 PE=1 SV=1 | 151.9 | 111.4 | 131.5 | 113.6 | 67.8 | 114.2 | 62.7 | 96 | 93.1 | 57.7 |
| 1216 | CD2-associated protein OS=Mus musculus OX=10090 GN=Cd2ap PE=1 SV=3 | 86.8 | 93.8 | 107.8 | 84.9 | 125.8 | 113 | 98.3 | 89.3 | 82.6 | 117.8 |
| 1217 | BAG family molecular chaperone regulator 3 OS=Mus musculus OX=10090 GN=Bag3 PE=1 SV=2 | 92.7 | 105.3 | 89.5 | 74.7 | 160.8 | 105.6 | 97.5 | 89 | 74.7 | 110.2 |
| 1218 | Methylglutaconyl-CoA hydratase, mitochondrial OS=Mus musculus OX=10090 GN=Auh PE=1 SV=1 | 93.3 | 92.7 | 103.3 | 111.4 | 91.6 | 97.1 | 51.1 | 113 | 125.1 | 121.3 |
| 1219 | Cytochrome P450 3A41 OS=Mus musculus OX=10090 GN=Cyp3a41a PE=1 SV=2 | 89.8 | 95.9 | 226.6 | 127.7 | 25.7 | 125.3 | 67.5 | 46.8 | 115.3 | 79.6 |
| 1220 | Endothelial differentiation-related factor 1 OS=Mus musculus OX=10090 GN=Edf1 PE=1 SV=1 | | 150.2 | | 110.6 | | 152.8 | 154.7 | 143.3 | 129.8 | 158.6 |

| | | | | | | | | | | | |
|---|---|---|---|---|---|---|---|---|---|---|---|
| 1221 | Glutaredoxin-1 OS=Mus musculus OX=10090 GN=Glrx PE=1 SV=3 | 111 | 105.3 | 106.3 | 75.3 | 121.7 | 81.2 | 143.5 | 89.6 | 85.3 | 80.6 |
| 1222 | Peroxisomal acyl-coenzyme A oxidase 2 OS=Mus musculus OX=10090 GN=Acox2 PE=1 SV=2 | 94 | 83.4 | 87.1 | 99 | 75.4 | 88.6 | 74.7 | 133.9 | 128.3 | 135.5 |
| 1223 | Fructose-1,6-bisphosphatase 1 OS=Mus musculus OX=10090 GN=Fbp1 PE=1 SV=3 | 114.7 | 106.1 | 106.1 | 112 | 96.3 | 111.2 | 82.9 | 104 | 100.5 | 66.2 |
| 1224 | Glycine N-methyltransferase OS=Mus musculus OX=10090 GN=Gnmt PE=1 SV=3 | 106.5 | 94.9 | 93.7 | 136.3 | 69.2 | 97.4 | 57.5 | 126.5 | 126.7 | 91.2 |
| 1225 | Acetyl-coenzyme A synthetase, cytoplasmic OS=Mus musculus OX=10090 GN=Acss2 PE=1 SV=2 | 159.8 | 127.5 | 136 | 81.4 | 108.7 | 103.3 | 102.6 | 68.2 | 56.9 | 55.6 |
| 1226 | Coatomer subunit gamma-2 OS=Mus musculus OX=10090 GN=Copg2 PE=1 SV=1 | 157.4 | 139.2 | 149.9 | 92 | | 160.6 | 51.2 | 89.5 | 98.2 | 62 |
| 1227 | Plectin OS=Mus musculus OX=10090 GN=Plec PE=1 SV=3 | 81.8 | 83.1 | 77.9 | 79.8 | 133.5 | 84.7 | 188 | 90.6 | 87.1 | 93.5 |
| 1228 | Calcium-binding mitochondrial carrier protein Aralar2 OS=Mus musculus OX=10090 GN=Slc25a13 PE=1 SV=1 | 110.2 | 98.6 | 106.6 | 111.9 | 74.6 | 103.5 | 63.9 | 113 | 112.2 | 105.5 |
| 1229 | EH domain-containing protein 3 OS=Mus musculus OX=10090 GN=Ehd3 PE=1 SV=2 | 110.8 | 100.8 | 100.2 | 112.3 | 72.9 | 94 | 57.6 | 124.4 | 111.6 | 115.4 |
| 1230 | Mitochondrial import receptor subunit TOM40 homolog OS=Mus musculus OX=10090 GN=Tomm40 PE=1 SV=3 | 102.2 | 91.5 | 102.2 | 111.5 | 85.2 | 87.3 | 82.4 | 120.5 | 113.6 | 103.6 |
| 1231 | DnaJ homolog subfamily A member 2 OS=Mus musculus OX=10090 GN=Dnaja2 PE=1 SV=1 | 110.9 | 106.7 | 113.6 | 94.1 | 98.7 | 103.1 | 88.7 | 100.9 | 86.9 | 96.3 |
| 1232 | Acyl-coenzyme A thioesterase 2, mitochondrial OS=Mus musculus OX=10090 GN=Acot2 PE=1 SV=2 | 107.9 | 97 | 135.4 | 133.7 | 62.1 | 96.2 | 47.2 | 119.4 | 121.3 | 79.7 |
| 1233 | Interferon-inducible GTPase 1 OS=Mus musculus OX=10090 GN=Iigp1 PE=1 SV=2 | 72.2 | 81.8 | 74.8 | 84.1 | 113.2 | 236.4 | 153.9 | 57.2 | 69.7 | 56.8 |
| 1234 | Mitochondrial dicarboxylate carrier OS=Mus musculus OX=10090 GN=Slc25a10 PE=1 SV=2 | 118.2 | 90.7 | 108.2 | 133.1 | 78.7 | 98.1 | 61.2 | 115.5 | 130.8 | 65.5 |
| 1235 | Coatomer subunit gamma-1 OS=Mus musculus OX=10090 GN=Copg1 PE=1 SV=1 | 163.5 | 141.3 | 115 | 97.4 | 59.7 | 106.2 | 62.2 | 90.5 | 88.7 | 75.6 |
| 1236 | Core histone macro-H2A.1 OS=Mus musculus OX=10090 GN=Macroh2a1 PE=1 SV=3 | 125.6 | 106.1 | 125.2 | 89.9 | 61.9 | 114 | 70.1 | 108.3 | 95 | 103.9 |
| 1237 | Cathepsin F OS=Mus musculus OX=10090 GN=Ctsf PE=1 SV=1 | 116.1 | 104.7 | 93.2 | 70.8 | 62.4 | 92.3 | 72.6 | 75.8 | 147.2 | 164.9 |
| 1238 | Glycogenin-1 OS=Mus musculus OX=10090 GN=Gyg1 PE=1 SV=3 | 83.9 | 84.7 | 93 | 86.9 | 149.8 | 86.6 | 205.4 | 70 | 69.9 | 69.9 |
| 1239 | Peroxisomal acyl-coenzyme A oxidase 1 OS=Mus musculus OX=10090 GN=Acox1 PE=1 SV=5 | 116 | 108.2 | 115.8 | 102.4 | 75.1 | 97.8 | 71.1 | 114.3 | 105.6 | 93.8 |
| 1240 | Galactokinase OS=Mus musculus OX=10090 GN=Galk1 PE=1 SV=2 | 127.7 | 104.3 | 114.8 | 109.9 | 100.6 | 123.2 | 62 | 90.6 | 85.9 | 81.1 |
| 1241 | Destrin OS=Mus musculus OX=10090 GN=Dstn PE=1 SV=3 | 117.4 | 110.1 | 96.8 | 104.9 | 89.1 | 106.8 | 66.4 | 108.6 | 89 | 110.9 |
| 1242 | Adenylate kinase isoenzyme 1 OS=Mus musculus OX=10090 GN=Ak1 PE=1 SV=1 | 17.5 | 43.5 | 25.5 | 33.8 | 317 | 19.3 | 436.1 | 19 | 25.9 | 62.3 |
| 1243 | Angiopoietin-related protein 3 OS=Mus musculus OX=10090 GN=Angptl3 PE=1 SV=1 | 105.9 | 100.6 | 123.6 | 109.6 | 90 | 114.5 | 86.3 | 74.2 | 78.1 | 117.2 |
| 1244 | Metastasis-associated protein MTA2 OS=Mus musculus OX=10090 GN=Mta2 PE=1 SV=1 | 109.7 | 104.9 | 101.6 | 80.8 | 96.9 | 110.8 | 111.7 | 102.4 | 81.8 | 99.4 |
| 1245 | ATP-binding cassette sub-family C member 6 OS=Mus musculus OX=10090 GN=Abcc6 PE=1 SV=3 | 158.4 | 121.7 | 139.8 | 87.5 | 73.5 | 122 | 58.4 | 77.1 | 87 | 74.5 |
| 1246 | Adenylate kinase 2, mitochondrial OS=Mus musculus OX=10090 GN=Ak2 PE=1 SV=5 | 88.6 | 101.4 | 80.3 | 107 | 120.1 | 96 | 105.2 | 110.5 | 95.2 | 95.7 |
| 1247 | GTP:AMP phosphotransferase AK3, mitochondrial OS=Mus musculus OX=10090 GN=Ak3 PE=1 SV=3 | 99.9 | 99 | 94.3 | 104.7 | 101.3 | 98.9 | 83.6 | 99.3 | 103.9 | 114.9 |
| 1248 | Cullin-1 OS=Mus musculus OX=10090 GN=Cul1 PE=1 SV=1 | 118.7 | 104.3 | 99.9 | 90.3 | 89.6 | 112.3 | 69.3 | 102 | 85.9 | 127.6 |
| 1249 | Glycerol kinase 2 OS=Mus musculus OX=10090 GN=Gk2 PE=1 SV=1 | 144.5 | 146.5 | 140.6 | 100.1 | 66.7 | 135.9 | 45.8 | 76.4 | 78.2 | 65.3 |
| 1250 | Programmed cell death 6-interacting protein OS=Mus musculus OX=10090 GN=Pdcd6ip PE=1 SV=3 | 69.7 | 72.2 | 70.4 | 90.5 | 161.4 | 101.3 | 170.9 | 72.7 | 86.8 | 104 |
| 1251 | Proline dehydrogenase 1, mitochondrial OS=Mus musculus OX=10090 GN=Prodh PE=1 SV=2 | 101.7 | 86.1 | 88.2 | 113.4 | 70.6 | 90.8 | 52.5 | 120 | 120.7 | 156.1 |
| 1252 | Phenylalanine--tRNA ligase beta subunit OS=Mus musculus OX=10090 GN=Farsb PE=1 SV=2 | 97.3 | 126.3 | 97.5 | 110.3 | 67.1 | 139.9 | 71.8 | 83.9 | 120.6 | 85.4 |
| 1253 | Glycogen phosphorylase, muscle form OS=Mus musculus OX=10090 GN=Pygm PE=1 SV=3 | 25.3 | 60.7 | 23.2 | 72 | 81.3 | 59.5 | 462.6 | 53.5 | 59.3 | 102.6 |
| 1254 | Cathepsin Z OS=Mus musculus OX=10090 GN=Ctsz PE=1 SV=1 | 104.1 | 94.4 | 111.1 | 89.2 | 80.7 | 111.5 | 64.9 | 105.6 | 98 | 140.3 |
| 1255 | Actin-related protein 2/3 complex subunit 1B OS=Mus musculus OX=10090 GN=Arpc1b PE=1 SV=4 | 129.8 | 107.7 | 119.9 | 84.5 | 90.3 | 144.2 | 72.4 | 78.5 | 81.7 | 91.1 |
| 1256 | Acid ceramidase OS=Mus musculus OX=10090 GN=Asah1 PE=1 SV=1 | 104.8 | 98 | 106.4 | 92.4 | 106.9 | 116.6 | 80.8 | 96.2 | 90.2 | 107.8 |

| | | | | | | | | | | | |
|---|---|---|---|---|---|---|---|---|---|---|---|
| 1257 | Peroxisomal 2,4-dienoyl-CoA reductase [(3E)-enoyl-CoA-producing] OS=Mus musculus OX=10090 GN=Decr2 PE=1 SV=1 | 112.3 | 103.6 | 106 | 112.3 | 69.6 | 97.9 | 65.6 | 105.4 | 121.7 | 105.7 |
| 1258 | Nucleoside diphosphate kinase 3 OS=Mus musculus OX=10090 GN=Nme3 PE=1 SV=3 | 101.6 | 105.8 | 101.2 | 80.1 | 105.2 | 97.9 | 130.4 | 86.9 | 82.7 | 108.2 |
| 1259 | Mitochondrial ornithine transporter 1 OS=Mus musculus OX=10090 GN=Slc25a15 PE=1 SV=1 | 90.7 | 80.2 | 86.1 | 158.5 | 48.4 | 72.7 | 50.5 | 148.7 | 147.7 | 116.7 |
| 1260 | Carboxypeptidase Q OS=Mus musculus OX=10090 GN=Cpq PE=1 SV=1 | 97.5 | 111.3 | 95.5 | 84.1 | 117.8 | 98 | 118.2 | 78.8 | 91.3 | 107.6 |
| 1261 | Kynurenine/alpha-aminoadipate aminotransferase, mitochondrial OS=Mus musculus OX=10090 GN=Aadat PE=1 SV=1 | 83.5 | 76.3 | 81.3 | 138.8 | 46.7 | 76.2 | 46.3 | 149.5 | 158 | 143.5 |
| 1262 | NPC intracellular cholesterol transporter 2 OS=Mus musculus OX=10090 GN=Npc2 PE=1 SV=1 | 114.4 | 114.1 | 108.4 | 67.3 | 152.3 | 97.3 | 75 | 89.8 | 76.1 | 105.3 |
| 1263 | Lysosomal acid lipase/cholesteryl ester hydrolase OS=Mus musculus OX=10090 GN=Lipa PE=1 SV=2 | 153 | 138.6 | 131.5 | 86.6 | 87.2 | 115.7 | 86.5 | 66.4 | 73.8 | 60.7 |
| 1264 | Eukaryotic translation initiation factor 2 subunit 3, X-linked OS=Mus musculus OX=10090 GN=Eif2s3x PE=1 SV=2 | 119.3 | 107.4 | 112.4 | 89.8 | 94.2 | 124.1 | 74.4 | 109.3 | 81.2 | 87.9 |
| 1265 | Acyl-CoA 6-desaturase OS=Mus musculus OX=10090 GN=Fads2 PE=1 SV=1 | 147.1 | 120.1 | 134.4 | 81.7 | 96 | 119.2 | 80.5 | 73.1 | 81.3 | 66.6 |
| 1266 | Eukaryotic translation initiation factor 3 subunit G OS=Mus musculus OX=10090 GN=Eif3g PE=1 SV=2 | 87 | 99.1 | 89 | 71.7 | 131 | 98.9 | 166.5 | 90.8 | 71.3 | 94.6 |
| 1267 | Cysteine desulfurase, mitochondrial OS=Mus musculus OX=10090 GN=Nfs1 PE=1 SV=3 | 106.1 | 93.9 | 104.1 | 99.5 | 96.6 | 95.7 | 96.5 | 106.4 | 103.6 | 97.6 |
| 1268 | NADH dehydrogenase [ubiquinone] 1 alpha subcomplex subunit 7 OS=Mus musculus OX=10090 GN=Ndufa7 PE=1 SV=3 | 86 | 81.7 | 66.9 | 102.7 | 132.1 | 80.2 | 119.6 | 117.6 | 91.4 | 121.8 |
| 1269 | Chloride intracellular channel protein 1 OS=Mus musculus OX=10090 GN=Clic1 PE=1 SV=3 | 126.8 | 103.2 | 115.4 | 95.1 | 95.3 | 137.9 | 69.9 | 90.7 | 84.7 | 81 |
| 1270 | AP-3 complex subunit beta-1 OS=Mus musculus OX=10090 GN=Ap3b1 PE=1 SV=2 | 101.5 | 93.8 | 104.4 | 80.4 | 122.1 | 99.5 | 154.9 | 76.7 | 83.5 | 83.4 |
| 1271 | Potassium-transporting ATPase alpha chain 2 OS=Mus musculus OX=10090 GN=Atp12a PE=1 SV=3 | 97.5 | 88.3 | 93.4 | 102.5 | 76.2 | 88 | 86.5 | 123.7 | 114.5 | 129.3 |
| 1272 | Oxidoreductase HTATIP2 OS=Mus musculus OX=10090 GN=Htatip2 PE=1 SV=3 | 91.1 | 88 | 93 | 98.7 | 113.8 | 94.5 | 88.6 | 122.5 | 105.9 | 104 |
| 1273 | Mitochondrial proton/calcium exchanger protein OS=Mus musculus OX=10090 GN=Letm1 PE=1 SV=1 | 106.2 | 99.3 | 93.6 | 106.2 | 83.7 | 88.8 | 74.3 | 121.2 | 117.5 | 109.3 |
| 1274 | Phosphomannomutase 2 OS=Mus musculus OX=10090 GN=Pmm2 PE=1 SV=1 | | 154.5 | | 135.4 | | 193.9 | 130.3 | 135.8 | 114.5 | 135.6 |
| 1275 | Actin-like protein 6A OS=Mus musculus OX=10090 GN=Actl6a PE=1 SV=2 | 118.4 | 103.2 | 106.2 | 92.8 | 97.5 | 108.2 | 98.8 | 93.9 | 68.4 | 112.6 |
| 1276 | Phosphoenolpyruvate carboxykinase, cytosolic [GTP] OS=Mus musculus OX=10090 GN=Pck1 PE=1 SV=1 | 75.7 | 79.7 | 72.8 | 117.7 | 69.9 | 69.6 | 43.7 | 171.3 | 145.1 | 154.6 |
| 1277 | Aspartyl aminopeptidase OS=Mus musculus OX=10090 GN=Dnpep PE=1 SV=2 | 128 | 118.2 | 127 | 96.6 | 75.8 | 116.5 | 62.4 | 93.7 | 87.2 | 94.5 |
| 1278 | Heterogeneous nuclear ribonucleoprotein F OS=Mus musculus OX=10090 GN=Hnrnpf PE=1 SV=3 | 117.2 | 103.1 | 112.1 | 85 | 99.7 | 116.4 | 75.5 | 100.2 | 85.2 | 105.7 |
| 1279 | Mitochondrial carnitine/acylcarnitine carrier protein OS=Mus musculus OX=10090 GN=Slc25a20 PE=1 SV=1 | 109.7 | 102 | 112.4 | 101.2 | 96.1 | 93.5 | 89.8 | 107.8 | 98.5 | 89 |
| 1280 | 40S ribosomal protein S18 OS=Mus musculus OX=10090 GN=Rps18 PE=3 SV=1 | 121 | 105.7 | 122.9 | 94.1 | 97.4 | 122.6 | 87 | 97.5 | 81.9 | 70 |
| 1281 | Phosphoglycerate kinase OS=Mus musculus OX=10090 GN=Pgk1 PE=1 SV=1 | 102.5 | 98.2 | 96.2 | 100.9 | 101.4 | 92.8 | 100.8 | 105.8 | 99.1 | 102.4 |
| 1282 | ELKS/Rab6-interacting/CAST family member 1 OS=Mus musculus OX=10090 GN=Erc1 PE=1 SV=1 | | 113.3 | 112.6 | 99.4 | 106.7 | 135.6 | 116.7 | 84.7 | 91.6 | 139.4 |
| 1283 | Isocitrate dehydrogenase 3 (NAD+) beta (Fragment) OS=Mus musculus OX=10090 GN=Idh3b PE=1 SV=1 | 84.5 | 88.5 | 74.3 | 88.1 | 126.2 | 81.3 | 169.6 | 98.1 | 86.6 | 102.8 |
| 1284 | Myomesin-1 OS=Mus musculus OX=10090 GN=Myom1 PE=1 SV=1 | 76.7 | 89 | 84.3 | 79.9 | 145.4 | 80.2 | 219.7 | 82.4 | 64.2 | 78.2 |
| 1285 | Oxoglutarate dehydrogenase (succinyl-transferring) OS=Mus musculus OX=10090 GN=Ogdh PE=1 SV=1 | 48.1 | 75 | 43 | 121.6 | 46.5 | 91.8 | 185 | 106.9 | 139.9 | 142.2 |
| 1286 | Clustered mitochondria protein homolog OS=Mus musculus OX=10090 GN=Cluh PE=1 SV=1 | 106 | 98.2 | 100.4 | 102.6 | 73.2 | 116.6 | 83.8 | 104.5 | 103.7 | 110.9 |
